# Next Generation Opto-Jasplakinolides Enable Local Remodeling of Actin Networks

**DOI:** 10.1101/2022.02.21.480923

**Authors:** Florian Küllmer, Nynke A. Vepřek, Malgorzata Borowiak, Veselin Nasufović, Sebastian Barutzki, Oliver Thorn-Seshold, Hans-Dieter Arndt, Dirk Trauner

## Abstract

The natural product jasplakinolide is a widely used tool compound to stabilize F-actin and influence actin dynamics. We have previously introduced photoswitchable jasplakinolides (optojasps) that are activated with violet light and deactivated with blue light. Based on insights from cryo-electron microscopy and structure-activity relationship (SAR) studies, we now developed a new generation of functionally superior optojasps that are better suited for biological investigations. These compounds are procured through chemical total synthesis and feature rationally designed red-shifted azobenzene photoswitches. Our new optojasps can be activated with longer wavelengths in the visible range (e.g. 440-477 nm) and rapidly return to their inactive state through thermal relaxation. This has enabled the reversible control of F-actin dynamics, as shown through live-cell imaging and cell migration, as well as cell proliferation assays. Brief sub-cellular activation with blue-green light resulted in highly localized F-actin clusters that gradually dissolved in the dark. Our light-responsive tools can be useful in diverse fields to study actin dynamics with outstanding spatiotemporal precision.

**Graphical Abstract:** 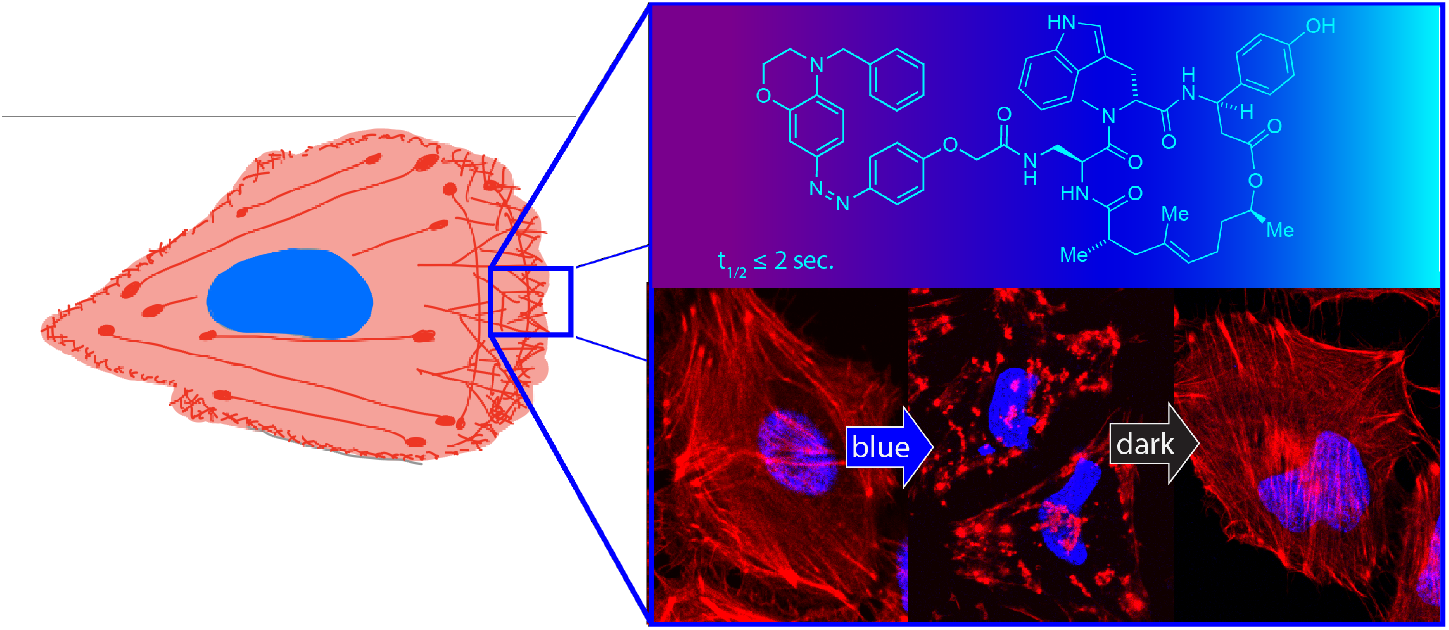

## Introduction

Actin is the most abundant protein in eukaryotic cells and a key element of the cytoskeleton. It is involved in a wide range of cellular functions: e.g. it serves as a regulator for cell shape and cell cycle progression, is implicated in various cellular signaling cascades, and indirectly regulates transcription.^1,2^ The actin cytoskeleton is formed by reversible assembly of monomeric G-actin to filamentous F-actin. The dynamics of this process and the large number of interactions of G- and F-actin with other proteins, such as profilin, thymosin β-4, tropomyosin, and cofilin, makes the actin cytoskeleton challenging to study mechanistically.^2,3^

Natural products have evolved that interfere with the dynamic instability of the actin cytoskeleton in various ways.^4^ These molecules not only affect the interaction between actin monomers but also interfere with actin binding proteins.^4,5^ Jasplakinolides and phalloidin disrupt actin dynamics by stabilizing F-actin as molecular glues that stabilize protomer assemblies.^6^ Kabiramide C and related structures can sever actin filaments,^7–9^ whereas cytochalasin D and pectenotoxin cap actin filaments at the barbed end and so prevent further polymerization.^10–12^ Latrunculin A sequesters actin monomers by blocking its ATP binding site,^13,14^ and swinholide A removes actin monomers from the G-actin pool as a 1:2 complex.^15,16^

Each of these sophisticated natural products, whose binding sites have been well characterized by structural biology, could become a basis for actin-targeting probes that impart additional functionalities. Of particular interest is optical control, which could be achieved by the introduction of a small molecule photoswitch, that changes the efficacy of the probe upon irradiation.^17–20^ We have recently reported on the optical control of F-actin with a first generation photoswitchable jasplakinolide (optojasp, **OJ**) that could be activated with violet and deactivated with blue light (Fig. 1A).^21^ We now present a new generation of optojasps that were developed based on structure-activity relationship (SAR) analysis and examination of the pose of optojasps inside the binding cleft. This new generation can be efficiently activated with blue to cyan light and shows a large change in potency between the active and inactive state. Importantly, these new optojasps thermally relax within seconds to the inactive form once the light source is turned off. With these improved biological, photophysical and thermal properties, we demonstrate temporally reversible optical control of the actin cytoskeleton with sub-cellular spatial resolution.

**Figure 1:**
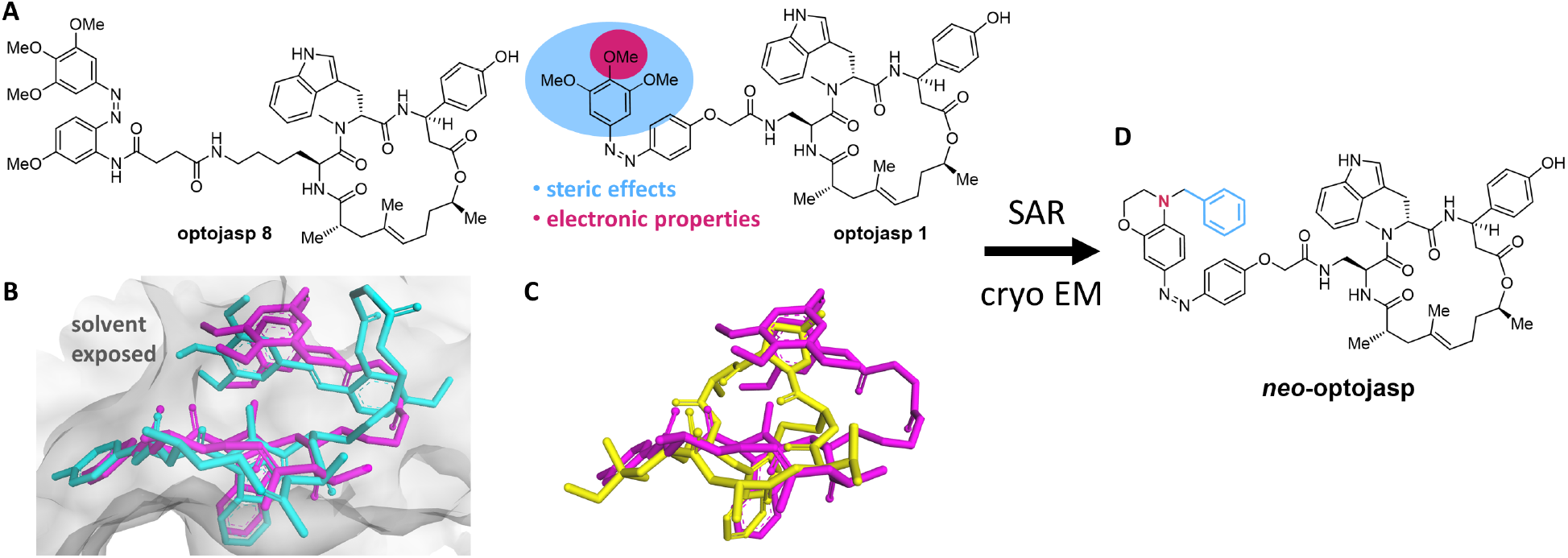
A) First generation optojasps for light dependent manipulation of F-actin. SAR analysis of optojasp targeting steric effects and electronic properties of the distal azobenzene ring (detailed information see text). B) cryo-EM structures of **optojasp 8** explain different binding affinities for the *cis* (magenta, PDB 7AHN) and *trans* (cyan, PDB 7AHQ) isomers. C) Mimic of phalloidin (yellow, PDB: 6T1Y) by *cis* **optojasp 8** (magenta) inside the actin binding cleft. D) Structure of ***neo*-optojasp** designed on the basis of SAR and cryo-EM analysis.

### Design, Synthesis, and Preliminary Evaluation

Our improved optojasp design was guided by three considerations: a) red-shifting the activation wavelength for biological experimentation under milder conditions, b) thermally destabilizing the bioactive *cis* isomer to suppress diffusion-based inhibition in biological studies, and c) enhancing the difference in potency between the two isomeric forms, with ideally only one isomer showing activity, and the other one being fully inactive in the dark. Optojasps feature the jasplakinolide macrocycle, which enables selective binding to the target, a linker, and the azobenzene switch that regulates the activity and specifies the photophysical properties. Our recent cryo-EM structure of **optojasp-8** bound to F-actin showed that the macrocyclic core adopts a pose that is very similar to jasplakinolide itself (Fig. 1B).^22^ The photoswitch binds in a pocket that is also occupied by phalloidin according to X-ray and cryo-EM structures.^6^ The *cis* azobenzene mimics the phalloidin-actin interaction better than the corresponding *trans* isomer, explaining the increased activity of **optojasp-8** upon irradiation (Fig. 1C). The linker was not resolved in the cryo-EM structures and seemed to be inconsequencial for binding. Upon further inspection of the photoswitch binding pocket, we reasoned that an increase in steric demand at the distal arene should further enhance differences in activity between the isomers. According to our analysis, the *trans* isomer should not be able to bind at all if sufficiently bulky derivatives were employed. By contrast, a sterically demanding substituent of the *cis* isomer could be accommodated. Aiming at redshifted and thermally destabilized photoswitches, we therefore decided to explore *para* amino azobenzenes that are larger than the original optojasp series. Electron rich aniline-derived azobenzenes are well known to be both red-shifted and fast relaxing.^23,24^

The new series of photoswitchable jasplakinolide derivatives was synthesized following previously developed methodology (Fig. 2).^25,26^ Easily obtained peptide carboxylic acid **2** underwent esterification with secondary alcohol **3** to yield diene **4**. A subsequent ring closing metathesis, followed by deprotection, furnished a primary amine, which underwent amide coupling with a variety of azobenzene carboxylic acids.^27^ This gave a library of depsipeptides, exemplified by compounds **5-14**, which feature a benzomorpholine moiety with different *N*-substituents. Derivatives with an extended linker, such as **15**, were explored as well (Fig 2D, Table **S6**). In addition, we explored sterically less demanding azobenzene derivatives, including simple anilines and phenol ethers (Table **S3, S4**). Modifications of the macrocyclic core, such as additional methyl groups and amide bonds (**16**), were also investigated (Table **S1, S2**). Substitutions on the proximal arene, as well as extended arenes and additional polar substituents were briefly investigated, too (Table **S5**).

**Figure 2:**
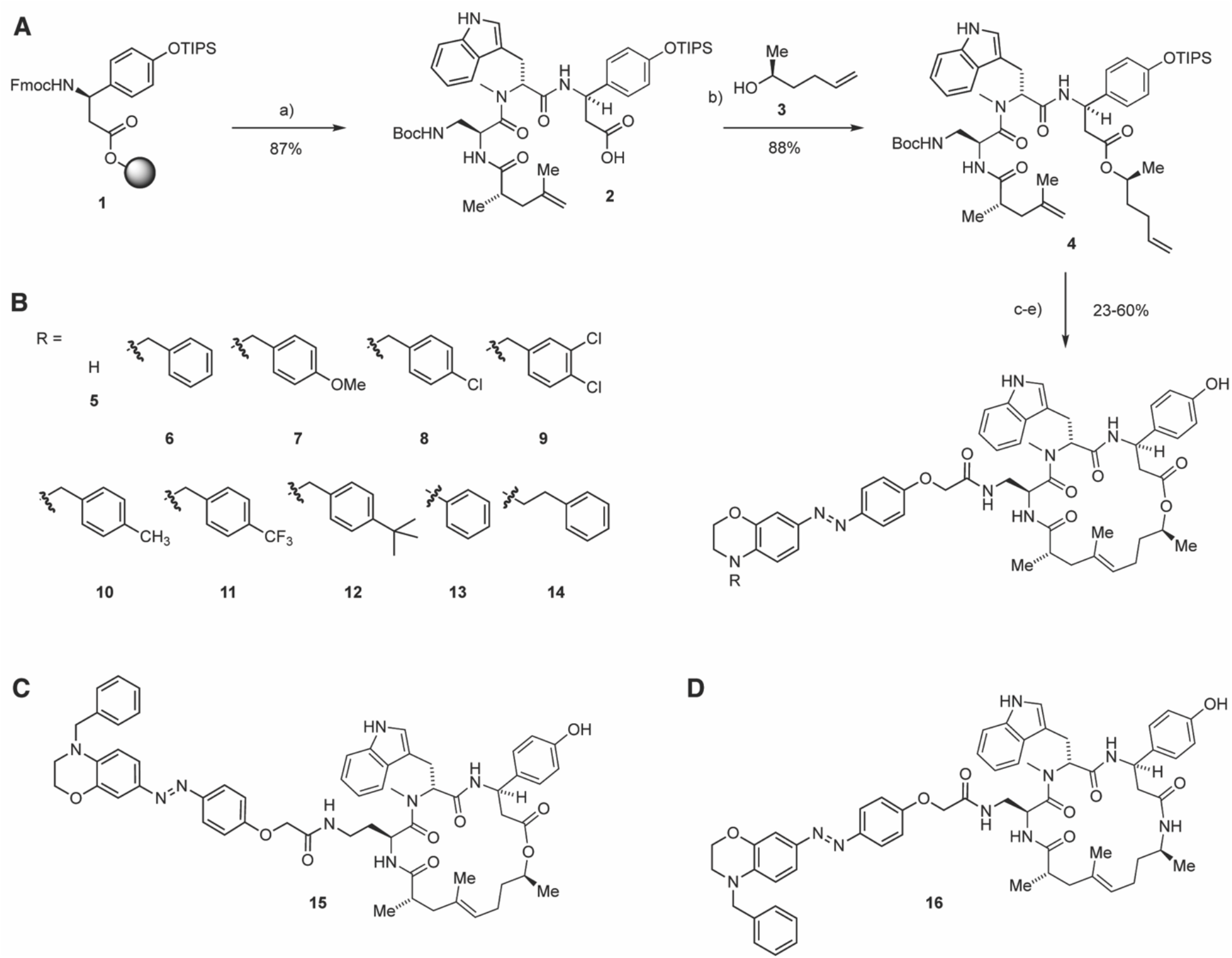
Synthesis of new generation of optojasps. a) 1. piperidine, DMF; 2. HATU, HOAt, *N*-Fmoc-*N*-methyl-L-tryptophan, DIPEA, CH_2_Cl_2_/DMF; 3. piperidine, DMF; 4. HATU, HOAt, Fmoc-Dap(Boc)-OH, DIPEA, CH_2_Cl_2_/DMF; 5. piperidine, DMF; 6. HATU, HOAt, (*S*)-2,4-dimethylpent-4-enoic acid, DIPEA, CH_2_Cl_2_/DMF; 7. HFIP, CH_2_Cl_2_; b) EDCI, DMAP, DIPEA, **3**, CH_2_Cl_2_/DMF; c) Grubbs 2nd gen., CH_2_Cl_2_, reflux.; d) 4 M HCl in dioxane, CH_2_Cl_2_, 0 °C; e) azobenzene, HATU, DIPEA, THF; 2. HF-pyridine, THF.

The newly synthesized optojasps were first evaluated in light-dependent cell proliferation assays using a cell DISCO-system (Table 1 and Supporting Information).^28^ We found that compound 6 (R = Bn) was completely inactive in the dark, but became highly cytotoxic upon irradiation with 410 nm light. Introduction of *para* substituents on the benzyl residue was tolerated for a trifluoromethyl substituent, but methyl, methoxy or *tert*-butyl groups reduced activity of the *cis* isomer while remaining inactive in the dark. The introduction of halogen substituents did not improve activity of *cis* **nOJ**. Unsubstituted benzomorpholine **7** only showed negligible light dependence, whereas the phenethyl derivative **14** was inactive in both the *cis* and *trans* form.

**Table 1:** photophysical properties and antiproliferative activities (at 410 nm (25 ms every 0.5 s) or dark) of new optojasps.

| # | EC<sub>50</sub> [μM]<br>light | EC<sub>50</sub> [μM]<br>dark | Δ light/<br>dark | t<sub>1/2</sub><br>[sec.]<sup>a</sup> | λ(Abs<sub>max</sub>)<sup>a,b</sup><br>[nm] |
| --- | --- | --- | --- | --- | --- |
| **5** | 0.37 | 0.54 | – | < 1 | 414 |
| **6** | 0.74 | >10 | ++ | 2 | 448 |
| **7** | >10 | >10 | – | < 1 | 448 |
| **8** | >10 | >10 | – | 3 | 443 |
| **9** | 0.59 | >10 | ++ | 8 | 438 |
| **10** | 0.99 | >10 | ++ | 2 | 448 |
| **11** | 0.57 | >10 | ++ | 6 | 437 |
| **12** | >10 | >10 | – | 2 | 449 |
| **13** | 0.45 | >10 | ++ | 22 | 438 |
| **14** | >10 | >10 | – | < 1 | 454 |
| **15** | 0.18 | 4.94 | + | 2 | 448 |

<sup>a</sup>measured at 37 °C in PBS: MeCN = 2:1; <sup>b</sup>*trans* isomer

Extending the linker length in **15** rendered *cis* **15** even more active, but *trans* 15 also showed increased activity, resulting in a smaller photopharmacological window. This was also observed for other derivatives with extended linkers (table **S6**) and sterically less demanding derivatives (table **S3, S4**). Modifications to the macrocyclic core did not result in improved light-dependence of activity, and amide derivatives such as **16** showed no cellular activity at all. Additional polar groups retained biological activity, albeit without light dependence (table **S5**). Modifications to the proximal arene of the photoswitch were not well tolerated (table **S5**). Overall, these observations support our hypothesis that an increased steric demand at the distal arene of the photoswitch is crucial for optimizing the difference in activity between the *cis* and *trans* isomers.

Having established that compound **6**, which we term ***neo*-optojasp** (**nOJ**), showed the most promising activity in these initial assays, we proceeded to investigate it in more detail (Fig. 3). We first evaluated the antiproliferative activity of **nOJ** as a function of irradiation wavelength (chromo-dosing, Fig, 3B). The highest antiproliferative activity was observed at 440 nm (EC_50_ = 0.37 μM) while wavelengths above 505 nm did not lead to a biological response. Antiproliferative activities at 410 and 477 nm irradiation were comparable, suiting **nOJ** to microscopy applications. The chromo-dosing results are in accordance with UV-VIS spectroscopic measurements, which show that irradiation with 460 and 415 nm led to most *cis* isomerization, while 365 and 520 nm gave less (Fig 3C). The half-life of *cis* **nOJ** was determined to be approximately 2 seconds in aqueous buffer at 37 °C (see Supporting Information).

**Figure 3:**
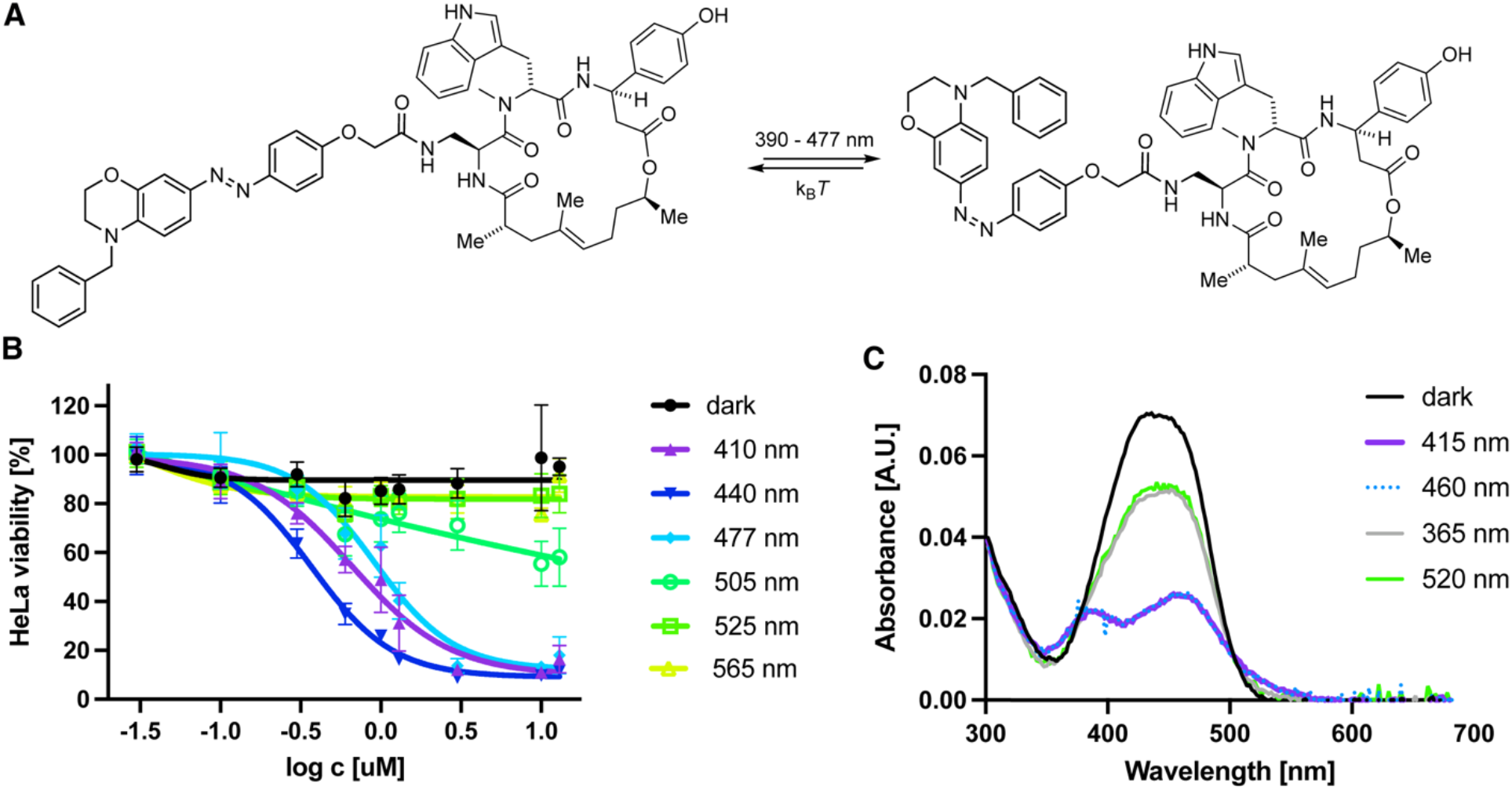
A) ***neo*-optojasp** B) wavelength dependent antiproliferative activity. C) UV-Vis spectra of **nOJ** under different irradiation conditions.

### Evaluation of neo Optojasp Using Cellular Imaging

Having established the advantageous photophysical properties of **nOJ** in cell proliferation assays, we proceeded to establish its function in different biological settings. In addition to visualization by fluorescent labelling in fixed cells, we monitored the activity of **nOJ** using live-cell imaging. Furthermore, we investigated the effect of our lead compound on cellular migration in wound healing assays.

Fig. 4A shows the effect of **nOJ** on HeLa cells using fluorescent labelling after fixation. In the activated (irradiated) form, **nOJ** induced aggregation of actin in a concentration dependent manner (Fig 4Ab, S1). The tubulin network remained unaffected under these conditions (Fig. 4Af). Treatment with **nOJ** in the inactive (dark-adapted) form resulted in no change in actin phenotype (Fig. 4Ae). Notably, the aggregates formed under irradiation gradually disappeared once the light was turned off, indicating the thermal deactivation of the filament stabilizer (16 h dark after 5 h 410 nm; Fig 4Ac). Reversibility of jasplakinolide induced F-actin aggregation upon washout has been described.^29^ Our first generation of optojasps, which did not thermally relax quickly, required irradiation with a second wavelength to enforce the inactive form and their reversibility was difficult to demonstrate.^21^ By contract **nOJ**’s rapid *cis→trans* relaxation makes for full reversibility.

**Figure 4:**
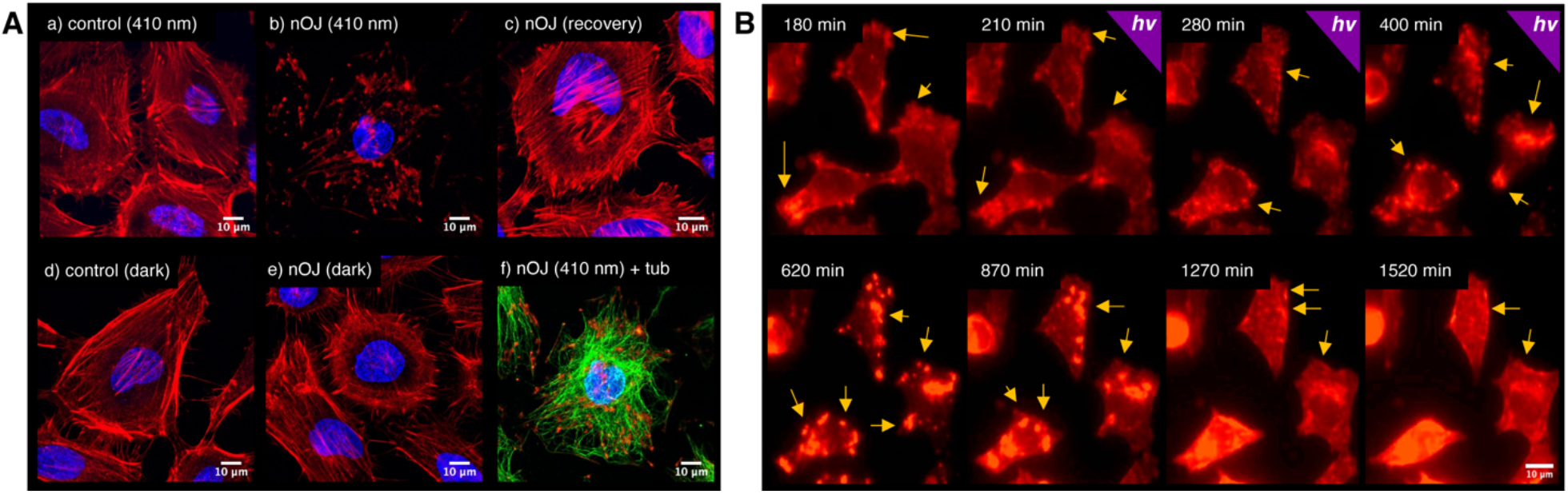
Light-dependent effect of optojasp **nOJ** on human cancer cells under global irradiation. **A**) HeLa cells were treated with **nOJ**, 3 μM or control (cosolvent) and exposed to either dark (5 h), 410 nm pulsed irradiation (5 h, 25 ms/ 0.5 s) conditions or first irradiation (5 h) followed by darkness (16 h). Upon treatment, cells were fixed, permeabilized and stained; nucleus: blue, actin: red, tubulin: green. (40×, confocal microscope). **B**) live-cell imaging of MDA-MB-231 mCherry:LifeAct treated with **nOJ** first in the dark (0-180 min), then under pulsed irradiation with 395 nm (25 ms/ 0.5 s) (181 – 420 min) followed by dark. (40×, widefield microscope).

The full reversibility of nOJ could also be shown using live-cell imaging. For this experiment we used MDA-MB-231 cells stably expressing mCherry:LifeAct, which is commonly employed to visualize F-actin.^30^ Notably, the onset of aggregation was delayed following light administration. However, within 30 minutes of irradiation, lamellipodium protrusions were reduced, followed by retraction (Fig 4B, t = 210 min). Subsequently, aggregate formation was observed (t = 280, 400 min). After the light was turned off, these aggregates first continued to grow in size and fused to form larger structures (t = 620 min). Subsequently, they migrated inwards and began to disassemble (t = 870 min). Approximately 20 h after irradiation stopped, the aggregates appeared to be largely dissolved (t = 1270, 1520 min). Under control conditions, i.e. in the absence of compound, the cell motility and protrusion dynamics remained unaffected during irradiation (Figure S2).

Our observations are in agreement with the effect of jasplakinolide on the inhibition of actin turnover and retrograde flow in lamellipodia.^31,32^ Actin polymerization and depolymerization in lamellipodia are essential for efficient treadmilling and filament turnover takes place within seconds to minutes.^32,33^ Inhibition of F-actin depolymerization leads to attenuated lamellipodium protrusion dynamics. Further, jasplakinolide has been reported to induce nucleation, which leads to a fast depletion of sequestered G-actin and also prevents filament elongation by monomer sequestration.^34^ Jasplakinolide treatment leads to the formation of F-actin puncta, i.e. amorphous aggregates that coalesce into large aggresomes and inclusion bodies.^35,36,37^ The cellular formation of these aggregates has been reported to be highly cell type dependent and to be linked to polymerization-competent G-actin concentrations. In cells with strongly pronounced actin cables and stress fibers, significant aggregate formation is observed only after the G-actin concentration has been augmented by remodeling of stress fibers.^34^ The delayed observation of aggregate formation with respect to light administration can be explained by a sudden drop in the local concentration of G-actin capable of polymerization upon **nOJ** binding. Before filament elongation and aggregation can occur, a reorganization of the actin network is required. The prolonged growth and coalescence of actin aggregates after deactivation of **nOJ** (light off), as well as retrograde flow are linked to myosin II and microtubule dynamics.^35,38^ Such a delay may be expected, as neither myosin II nor tubulin are affected by jasplakinolide (and hence, optojasps).^32,35^ Disaggregation depends on chaperones, microtubules, proteasomes and autophagy,^29^ which can explain the observed slowness of actin aggregate disassembly that we observed.

The advantage of a photoswitchable jasplakinolide is the possibility of dosing the probe with high local and spatial precision, without background activity outside an irradiated area. Therefore, we next set out to study local activation of **nOJ** by irradiation of a small region outside of the cell. We observed that this mode of activation led to strongly localized responses (Fig. 5A). Initially, actin turnover slowed down in the lamellipodium protrusions proximal to the region of irradiation (ROI). Several minutes after light administration, this was followed by retraction of this cellular region, while protrusion dynamics remained unaltered or even accelerated in areas of the cell that were not in close proximity of the ROI. This effect can be illustrated by using a kymograph, which shows time dependent changes along a specified axis (Fig. 5Aa-c). As shown in Fig 5Ab, an increase in actin protrusion dynamics can be observed in the lamellipodium region distant to the ROI. On the other hand, in the lamellipodium region close to the ROI, actin dynamics initially slowed upon irradiation before the lamellipodium was retracted (Fig. 5Aa). Our results demonstrate that the active *cis* form of **nOJ** can be locally generated within a specified area outside of the cell and then diffuse through the cell membrane to reach the actin networks inside the cell.

**Figure 5:**
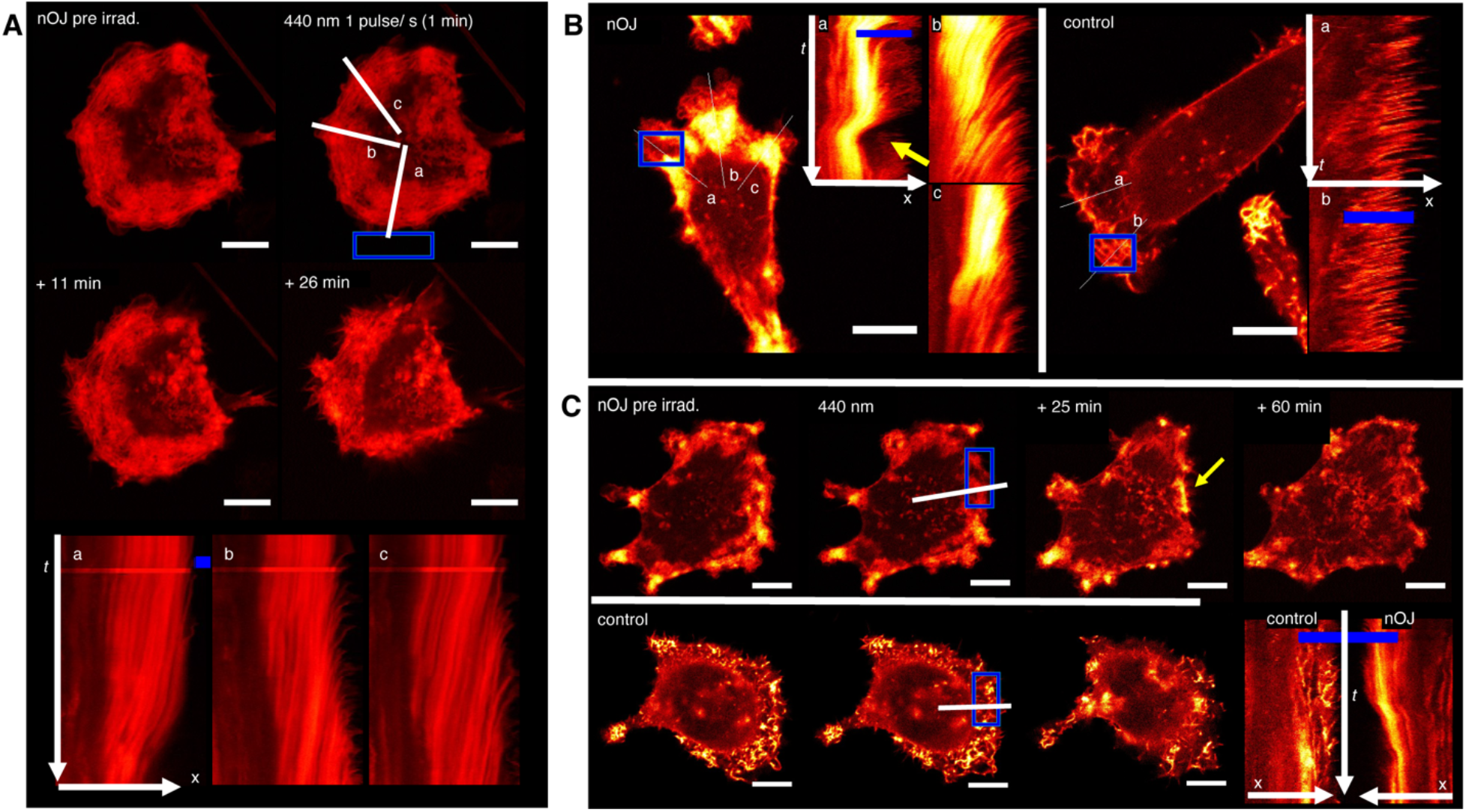
Live-cell imaging under local activation of **nOJ: A**) A small region (ROI, blue) outside of the MDA-MB-231 mCherry:LifeAct cell was irradiated with 440 nm pulses (1 min, ca. 1 pulse/s) lines and a/b/c indicate axes represented in kymographs. **B**) in the presence of **nOJ**, MDA-MB-231 mCherry:LifeAct cells react strongly to ROI with 440 nm light pulses (1 puls/ sec.): control: no change in actin dynamics. **C**) MDA-MB-231 mCherry:LifeAct cells with irradiation in the lamellipodium. Top row in presence of **nOJ**, bottom row control (cosolvent). Bottom right direct comparison of kymographs of treated and untreated cell; all scale bars: 10 μm; C-E: confocal microscope, 63× objective. mCherry:LifeAct signal graded from red (lower intensity) to yellow (high intensity), intensities between images/ cells are not comparable (expression level differences)

Notably, the decline of the effective concentration of *cis* **nOJ** through dilution is overlayed by the thermal inactivation of the photoswitch.

Next, we investigated local activation of **nOJ** in a region inside of the cell. We selected an area within a lamellipodium that was irradiated with the same light regime as previously applied. Again, protrusion dynamics initially slowed down before the lamellipodium was retracted. Aggregates were dissolved over time and the lamellipodium forefront was reestablished (Fig. 5 B/C, Supporting Video). As observed before, actin dynamics in regions that were more distant from the activation area, and therefore not in immediate contact with *cis* **nOJ**, did not show this effect. While retraction and reversion are processes that are often observed in migrating cells, this process is fast in unirradiated areas that were not affected by **nOJ** or in control cells. These dynamics appeared to be much slower when induced by *cis* **nOJ** (see above).

It has been reported that the speed with which cells respond to jasplakinolide itself, can vary strongly between differently polarized cells and different cell types.^31,34^ In polarized cells, the actin networks are organized such that fast filament turnover can be ensured to drive directional membrane extension.^39^ In nonpolarized cells, the barbed ends of actin filaments are capped and filament turnover is much slower.^40^ It is therefore not surprising, that the effects of *cis* **nOJ** take place on different time scales depending on the actin network organization of a particular cell.

Collectively, our experiments demonstrate that ***neo*-optojasp** can be activated with comparable efficiency outside and inside of the cell and that the photophysical properties of **nOJ** are not greatly affected by the extra or intracellular milieu.

### Cell Migration Assays

Next, we monitored the effect of ***neo*-optojasp** on the migratory behavior of the invasive human breast cancer cell line MDA-MB-231 in *in vitro* wound healing assays.^41,42^ To ensure that cytotoxicity did not interfere with migration, we chose low concentrations of **nOJ**. Under control conditions, full wound closure occurred between 24 and 48 h. Therefore, quantification of the effects of **nOJ** was based on relative wound closure after 24 h. While wound closure in the presence of inactive *trans* **nOJ** (i.e. in the dark) was comparable to the control, activation of **nOJ** with light resulted in a strongly concentration dependent reduction of wound closure (Fig. 6B). Cell viability after 48 h of treatment indicated only low cytotoxicity of light-activated **nOJ** at 0.5 μM. Lower concentrations appeared not to be cytotoxic at all.

**Figure 6:**
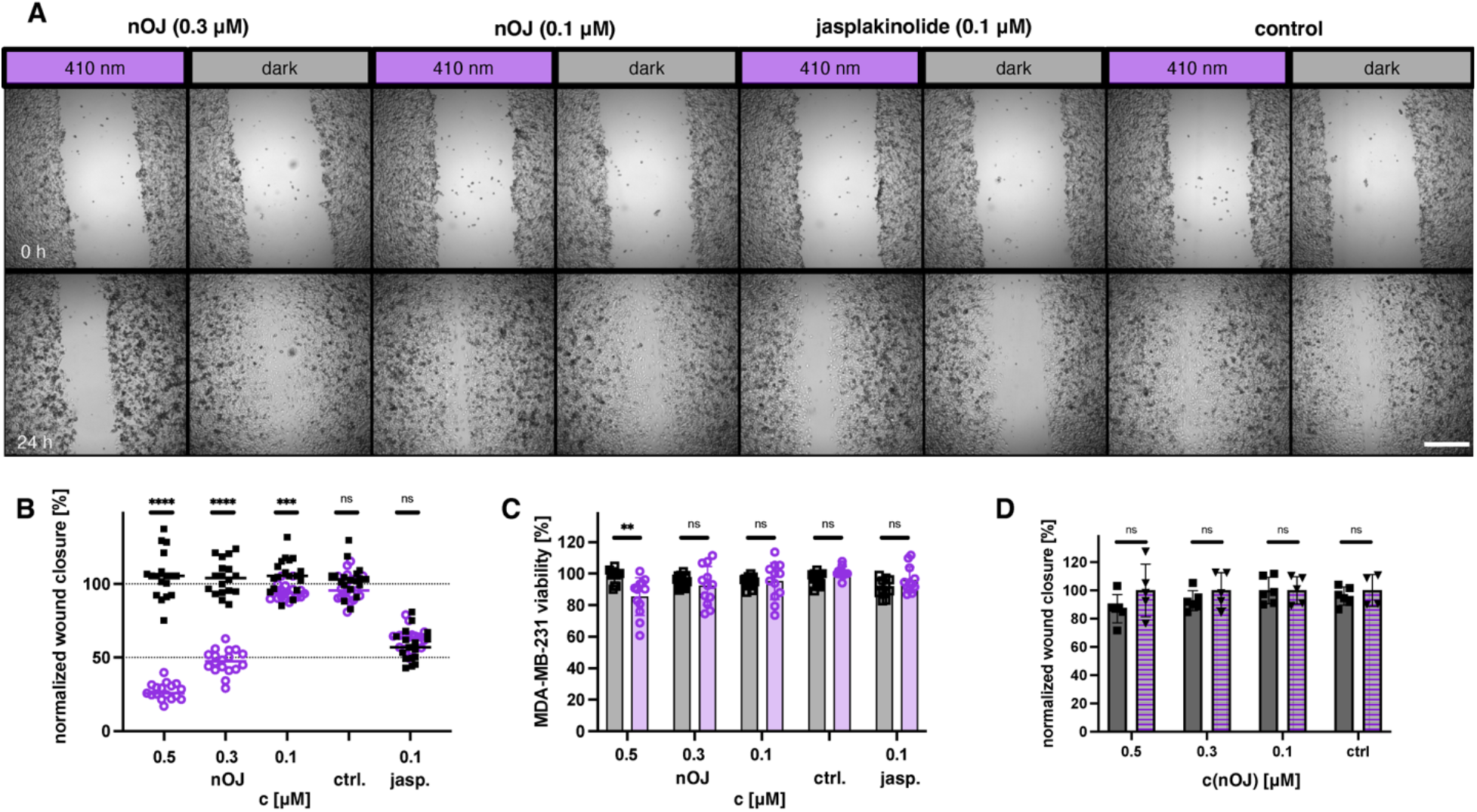
Wound healing assay on MDA-MB-231 cells. **A**) representative images of wound closure before treatment (t = 0 h, top row) and after treatment (t = 24 h, bottom row) under dark or pulsed irradiation (410 nm, 25 ms/ 1 s) conditions for selected concentrations and control. (2.5x, widefield microscope), scale bar = 500 μm. **B**) normalized wound closure from three independent experiments n = 17-18, round: light, squares: dark; line indicates median. **C**) MDA-MB-231 viability (PrestoBlue) after 48 h treatment with optojasp or jasplakinolide light/dark. **D**) comparison of wound closure between rescue (2 h 410 nm (25 ms/ 1 s), then dark) and dark. squares: dark, triangle: rescue; statistics: p ≤ 0.00001 ****, p ≤ 0.0001 ***,p ≤ 0.001 **, ns = not significant; multiple Mann-Whitney test for comparison between light/dark.

We then investigated whether cell migration could be recovered following thermal relaxation after 2 h of pulsed light activation. Indeed, the degree of wound closure in these rescue experiments was comparable to the extent observed in the dark (Fig. 6D). While reports of an increase in migration of cells treated with very low doses of jasplakinolide exist,^43^ we have not observed this effect in our study.

## Discussion

We have introduced a new generation of photoswitchable jasplakinolides that are optimized for biological experimentation in terms of activation wavelength, thermal relaxation, reversibility, and differences in potency between the two isomers. Our lead compound, **nOJ**, is inactive in the dark and becomes almost half as potent as jasplakinolide itself upon irradiation with blue light. **nOJ** shows superior qualities in comparison to our first generation of compounds, quite apart from its significantly improved potency. The efficient activation at 477 nm allows for its use in cells and tissues that are sensitive to deep violet or UV-A light. The low thermal stability of *cis* **nOJ**, combined with the possibility of applying high concentrations of the inactive isomer, enables fast activation and deactivation in a confined area of interest. As such, it can be used to stabilize F-actin with subcellular resolution. This thermal inactivation through fast relaxation can distinguish photoswitchable compounds from caged compounds, which, once released, only inactivate through diffusion on the time scales of interest.

Our chemical probe complements optogenetic tools that have been developed to control the actin cytoskeleton.^44–46^ While **nOJ** forgoes the power of genetic targeting of individual cell types, it does not require the overexpression of a factor that could perturb native protein networks and lead to unphysiological responses. As such, **nOJ** presents a powerful probe that allows for the study of systems in which local actin networks are formed or remodeled. For example, **nOJ** could enable the investigation of receptor mediated endocytosis, exocytosis, or fusion processes, as well as cell adhesion, with sub-cellular control. As an optical precision tool, **nOJ** could be used to study the local reorganization of actin networks and its effect on the redistribution of organelles.^47,48^ The inactivity of **nOJ** in the dark makes it suitable to study actin dependent processes in confined systems, tissues, organoids, and organisms where the localized application of generally toxic drugs may be challenging. Compounds such as **nOJ** could be envisioned to reduce the invasiveness of cancer cells and localize cytotoxicity in malignant tissues, potentially opening an avenue for adapting actin targeting natural products towards cancer chemotherapy.

## Acknowledgements

The authors thank Prof. Anna Akhmanova (Utrecht University) for a generous gift of the MDA-MB-231 mCherry:LifeAct cell line, and Dr. Peter Bellstedt (NMR platform Jena) for excellent support. We thank the Studienstiftung des deutschen Volkes for a PhD fellowship (to N.A.V.) and the Deutsche Akademischen Austauschdienst for a PhD fellowship (to V.N.). We thank the Deutsche Forschungsgemeinschaft (SFB 1032 project B09 number 201269156 and Emmy Noether number 400324123 to O.T.S.; Cluster of Excellence EXC2051 Balance of the Microverse— Project-ID 390713860, equipment grant INST 275/442-1 FUGG to H.D.A.), and the National Institutes of Health (Grant R01GM126228, to D.T.) for financial support.

## Data Availability Statement

The data that support the findings of this study are available from the corresponding authors upon reasonable request.

