## Supplementary Information for "Next Generation Opto-Jasplakinolides Enable Local Remodeling of Actin Networks"

- Supporting Information -

### Supplementary Tables and Figures

**Table S1:**

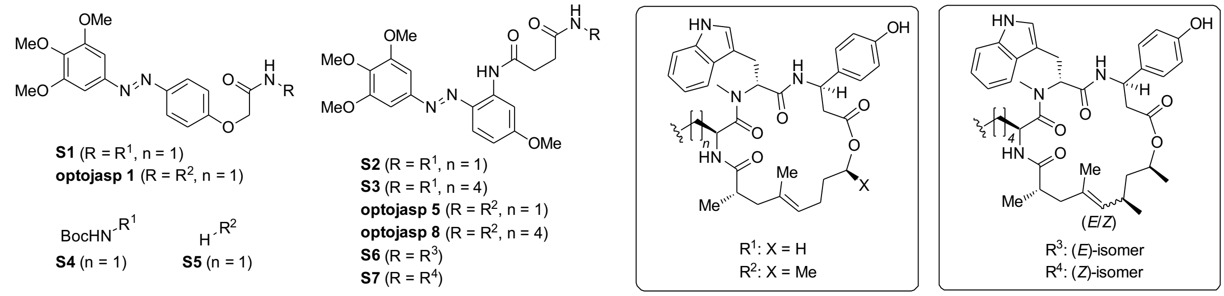

| Entry | $t_{1/2}$[a] [min] | $\lambda_{LED}$[b] [nm] | pulse[c] | EC50 (*dark*) [µM] | EC50 (*irrad.*) [µM] |
|---|---|---|---|---|---|
| **S1** | 360 | 390 | 75 ms every 15 s | > 50 | > 50 |
| **S2** | 385 | 390 | 75 ms every 15 s | > 50 | 1.6 |
| **S3** | 387 | 390 | 75 ms every 15 s | > 10 | > 10 |
| **S4** | - | - | - | 1.0 | 1.4 |
| **optojasp 1**[1] | 361 | 390 | 75 ms every 15 s | > 15 | 1.5 |
| **optojasp 5**[1] | 386 | 390 | 75 ms every 15 s | > 5 | 3.1 |
| **optojasp 8**[1] | 386 | 390 | 75 ms every 15 s | > 10 | 1.4 |
| **S5** | - | - | - | 0.1 | 0.1 |
| **S6** | 385 | 390 | 75 ms every 15 s | 0.2 | 0.2 |
| **S7** | 385 | 390 | 75 ms every 15 s | > 10 | > 10 |
| **jasplakinolide**[1] | - | 390 | 75 ms every 15 s | 0.13 | 0.15 |

[a] measured at 37 °C in PBS-buffer/$CH_3CN$ (2:1) [b] LEDs see supplementary information [c] Irradiation conditions during MTT-assay.

**Table S2:**

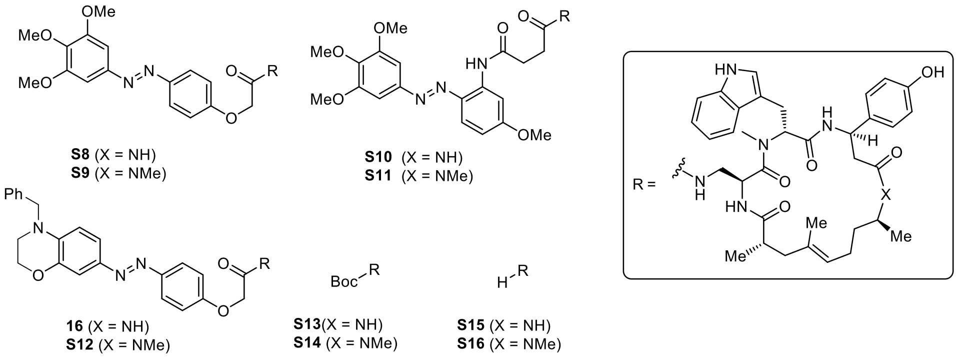

| Entry | $t_{1/2}$[a] [min] | $\lambda_{LED}$[b] [nm] | pulse[c] | EC50 (*dark*) [µM] | EC50 (*irrad.*) [µM] |
|---|---|---|---|---|---|
| **S8** | 298 | 370 | 75 ms every 15 s | > 15 | > 15 |
| **S9** | 293 | 370 | 75 ms every 15 s | > 15 | > 15 |
| **S10** | 15.6 | 390 | 75 ms every 15 s | > 10 | > 10 |
| **S11** | 16.6 | 390 | 75 ms every 15 s | > 15 | > 15 |
| **16** | 0.03 | 410 | 25 ms every 0.5 s | > 10 | > 10 |
| **S12** | 0.03 | 410 | 25 ms every 0.5 s | > 10 | > 10 |
| **S13** | - | - | - | > 10 | > 10 |
| **S14** | - | - | - | > 10 | > 10 |
| **S15** | - | - | - | > 10 | > 10 |
| **S16** | - | - | - | > 10 | > 10 |

[a] measured at 37 °C in PBS-buffer/$CH_3CN$ (2:1) [b] LEDs see supplementary information [c] Irradiation conditions during MTT-assay.

[1] M. Borowiak, F. Küllmer, F. Gegenfurtner, S. Peil, V. Nasufovic, S. Zahler, O. Thorn-Seshold, D. Trauner, H.-D. Arndt, *Journal of the American Chemical Society* **2020**, *142*, 9240-9249.

**Table S3:**

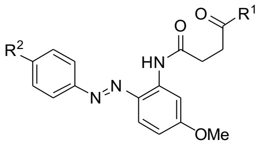

**S17** ($R^2$ = $NMe_2$, n = 1)
**S18** ($R^2$ = $NMe_2$, n = 2)
**S19** ($R^2$ = $NMe_2$, n = 3)
**S20** ($R^2$ = $NMe_2$, n = 4)
**S21** ($R^2$ = morp, n = 1)
**S22** ($R^2$ = nmp, n = 1)

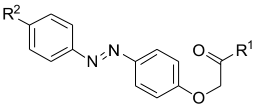

**S23** ($R^2$ = $NMe_2$, n = 1
**S24** ($R^2$ = morp, n = 1)
**S25** ($R^2$ = nmp, n = 1)

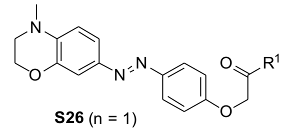

**S26** (n = 1)

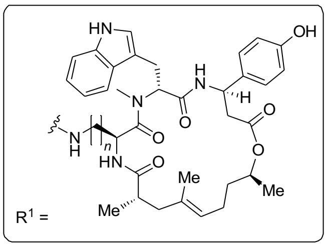

| Entry | $t_{1/2}$[a] [min] | $\lambda_{LED}$[b] [nm] | pulse[c] | EC50 (*dark*) [µM] | EC50 (*irrad.*) [µM] |
|---|---|---|---|---|---|
| **S17** | < 0.02 | 450 | 25 ms every 500 ms | 1.4 | 1.2 |
| **S18** | < 0.02 | 450 | 25 ms every 500 ms | 0.3 | 0.3 |
| **S19** | < 0.02 | 450 | 25 ms every 500 ms | 2.0 | 2.0 |
| **S20** | < 0.02 | 450 | 25 ms every 500 ms | 0.6 | 0.5 |
| **S21** | 0.21 | 425 | 25 ms every 500 ms | 3.5 | 2.3 |
| **S22** | 0.09 | 410 | 25 ms every 500 ms | 1.0 | 1.2 |
| **S23** | < 0.02 | 425 | 25 ms every 500 ms | 0.8 | 0.7 |
| **S24** | 0.08 | 390 | 25 ms every 2 s | 0.3 | 0.2 |
| **S25** | 0.05 | 390 | 25 ms every 2 s | 0.1 | 0.1 |
| **S26** | < 0.02 | 425 | 25 ms every 500 ms | 0.4 | 0.4 |

[a] measured at 37 °C in PBS-buffer/$CH_3CN$ (2:1) [b] LEDs see supplementary information [c] Irradiation conditions during MTT-assay.

**Table S4:**

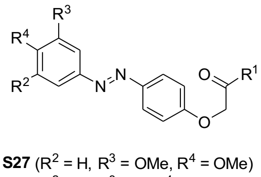

**S27** ($R^2$ = H, $R^3$ = OMe, $R^4$ = OMe)
**S28** ($R^2$ = H, $R^3$ = H, $R^4$ = OMe)
**S29** ($R^2$ = OMe, $R^3$ = OMe, $R^4$ = H)

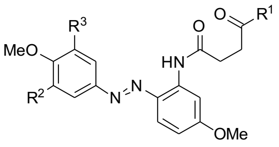

**S30** ($R^2$ = H, $R^3$ = OMe)
**S31** ($R^2$ = H, $R^3$ = H)

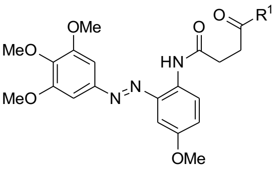

**S32**

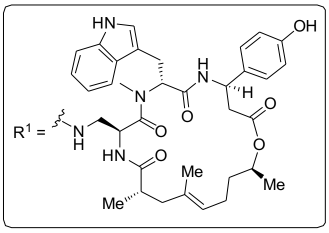

| Entry | $t_{1/2}$[a] [min] | $\lambda_{LED}$[b] [nm] | pulse[c] | EC50 (*dark*) [µM] | EC50 (*irrad.*) [µM] |
|---|---|---|---|---|---|
| **S27** | 68 | 370 | 75 ms every 15 s | 0.7 | 0.5 |
| **S28** | 254 | 360 | 75 ms every 15 s | 1.2 | 0.4 |
| **S29** | 4610 | 360 | 75 ms every 15 s | 3.9 | 1.7 |
| **S30** | 8.1 | 390 | 75 ms every 15 s | 1.2 | 0.8 |
| **S31** | 17.3 | 390 | 75 ms every 15 s | 0.9 | 0.6 |
| **S32** | 110 | 390 | 75 ms every 15 s | 0.9 | 0.8 |

[a] measured at 37 °C in PBS-buffer/$CH_3CN$ (2:1) [b] LEDs see supplementary information [c] Irradiation conditions during MTT-assay.

**Table S5:**

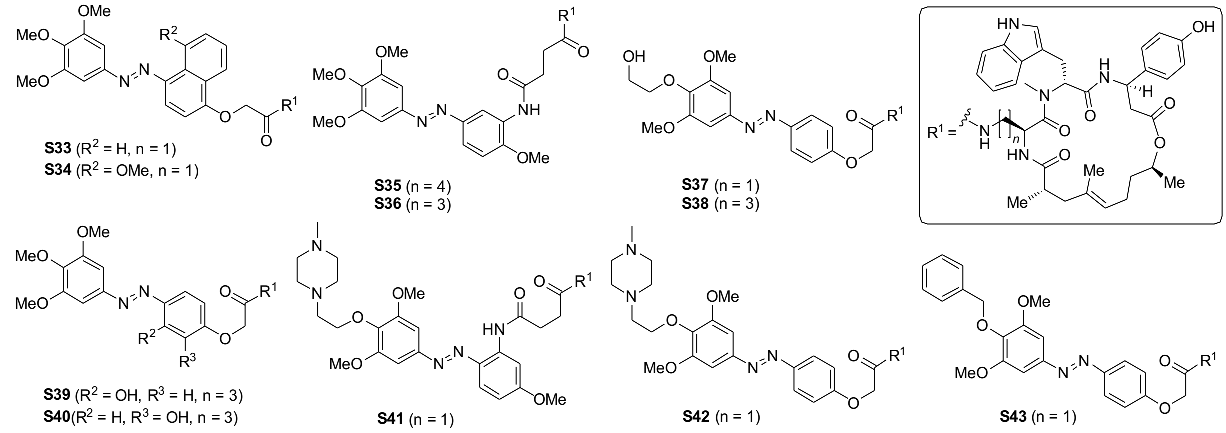

| Entry | $t_{1/2}$[a] [min] | $\lambda_{LED}$[b] [nm] | pulse[c] | EC50 (*dark*) [µM] | EC50 (*irrad.*) [µM] |
|---|---|---|---|---|---|
| **S33** | 6.2 | 390 | 75 ms every 15 s | >10 | >10 |
| **S34** | 399 | 390 | 75 ms every 15 s | >10 | >10 |
| **S35** | 145 | 390 | 75 ms every 15 s | 1.0 | 1.7 |
| **S36** | 149 | 390 | 75 ms every 15 s | >20 | >20 |
| **S37** | 271 | 360 | 75 ms every 15 s | 0.6 | 0.8 |
| **S38** | 275 | 360 | 75 ms every 15 s | 0.1 | 0.1 |
| **S39** | < 0.02 | 400 | 25 ms every 500 ms | 0.1 | 0.2 |
| **S40** | 77.7 | 370 | 75 ms every 15 s | >10 | >10 |
| **S41** | 7.8 | 390 | 75 ms every 15 s | 1.6 | 1.1 |
| **S42** | 23.0 | 370 | 75 ms every 15 s | 0.1 | 0.1 |
| **S43** | 337 | 380 | 75 ms every 15 s | >10 | >10 |

[a] measured at 37 °C in PBS-buffer/$CH_3CN$ (2:1) [b] LEDs see supplementary information [c] Irradiation conditions during MTT-assay.

**Table S6:**

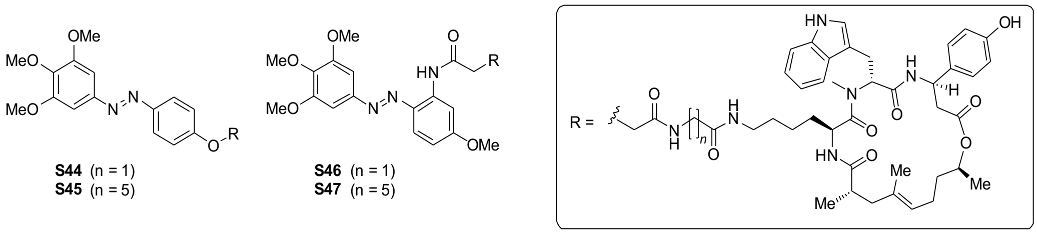

| Entry | $t_{1/2}$[a] [min] | $\lambda_{LED}$[b] [nm] | pulse[c] | EC50 (*dark*) [µM] | EC50 (*irrad.*) [µM] |
|---|---|---|---|---|---|
| **S44** | 253 | 390 | 75 ms every 15 s | 0.6 | 0.8 |
| **S45** | 258 | 390 | 75 ms every 15 s | 0.7 | 0.6 |
| **S46** | 16.9 | 390 | 75 ms every 15 s | 8.5 | 3.6 |
| **S47** | 17.4 | 390 | 75 ms every 15 s | 0.2 | 0.1 |

[a] measured at 37 °C in PBS-buffer/$CH_3CN$ (2:1) [b] LEDs see supplementary information [c] Irradiation conditions during MTT-assay.

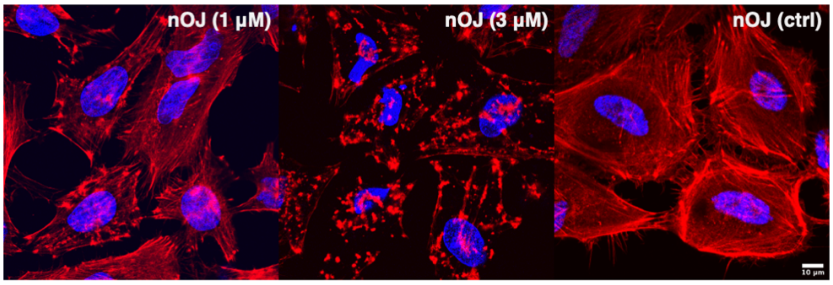

**Figure S1:** concentration dependent actin aggregation with **nOJ** (410 nm, 25 ms/ 0.5 s) versus control. Red: Phalloidin-iFluor594 (actin), blue: Hoechst (nucleus). Scale bar = 10 µm.

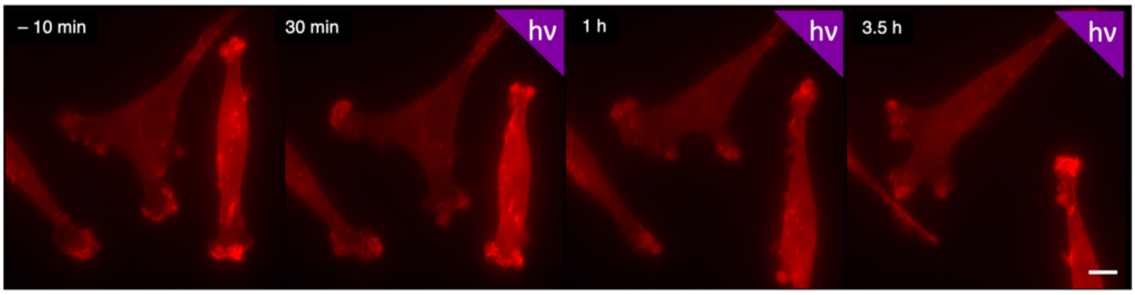

**Figure S2**: Live-cell imaging of MDA-MB-231 mCherry LifeAct cells under control conditions: cells were incubated with cosolvent only, imaged 3 h in the dark, then 4 h under light: λ = 395 nm, 25 ms/ 0.5 s. Scale bar = 10 µm.

#### Supplementary Video Material

##### Video for Fig 5c.

### Materials and Methods - Biology

#### UV-Vis-Spectroscopy

UV-Vis spectrometry was performed using a Varian Cary 60 UV-Visible Spectrophotometer using disposable UV-cuvettes (BRAND UV-Cuvette Disposable Spectrophotometer/ Photometer Cuvettes, BrandTech; Ultra-Micro Cuvettes Vol. 70 – 850 µl, window height 8.5 mm, pathlength 10 mm; VWR cat# 47743-834)). Sample temperatures were controlled using an Agilent Technologies PCB 1500 Water Peltier system and samples were irradiated orthogonally using prizmatix (U)HP-LEDs: 365 nm, 415 nm, 460 nm, 520 nm.

Thermal half-lives were determined by first order exponential decay fit using GraphPad Prism 9 for macOS (San Diego, CA, USA).

#### Cell Culture

HeLa, MDA-MB-231 (purchased from ATCC) and mCherry-LifeAct MDA-MB-231 cells (gift from Prof. A. Akhmanova, Utrecht University) were cultured in antibiotic free Dulbecco's Modified Eagle Medium (DMEM) (Gibco, Thermo Fisher cat# 10566024) supplemented with 10% fetal bovine serum (FBS) (Gibco, Thermo Fischer cat# 10437036) for the first 2 passages after thawing. Thereafter they were conditioned to phenol-red free DMEM (Thermo Fisher cat# 31053036), supplemented with 10% FBS (Gibco, Thermo Fischer cat# 10437036), 1% penicillin-streptomycin-glutamine (Gibco, Thermo Fischer cat# 10378016) and a final concentration of 4 mM L-glutamine (Gibco, Thermo Fischer cat# 25030081) for at least 1 passage before use in assays. Cells were grown in a cell culture incubator at 37 °C in a 5% $CO_2$ atmosphere and passaged at 70-90% confluency every 2-4 days. Cells were used for up to 25 passages for cell proliferation assays and up to 15 passages for imaging studies.

#### Handling of photoswitchable compounds

Test compounds were dissolved in DMSO (sterile filtered) to the desired stock-concentration (e.g. 10 mM) and stored at –20 °C. In case of long half-lives, the compound stock was left in the dark at room temperature for an appropriate time to ensure full thermal relaxation of the photoswitch. Compounds were protected from light and only handled in the dark or under red-light to avoid isomerization.

##### MTT Cell Proliferation Assay

Cells (HeLa: 5000, MDA-MB-231 mCherry LifeAct: 4000 cells) were seeded in 96 well-plates using 90 µl phenolred-free DMEM supplemented with 10% FBS 1% penicillin-streptomycin and a final concentration of 4 mM L-glutamine. After 24 h, cells were treated with compound stocks which were applied as 10x concentrations in 10 µl medium. As cosolvent, 1% DMSO and 2% MeCN final concentrations were used to ensure full solubility of all compounds at all concentrations and allow for comparability between all experiments. As reported previously, this did not alter cell growth and viability.[2,3] Light-dependent assays were performed as duplicates where one plate was kept in light-proof boxes, shielded from light (dark) and the second one was exposed to the irradiation protocol indicated using a cell DISCO as described previously[3]. After 48 h of treatment, MTT (Invitrogen, Thermo Fischer Cat# M6494; 10 µl, 5 mg/ml in PBS) was added to each well and incubated for 3 h (37 °C, 5% $CO_2$). The wells were emptied and the purple formazan crystals at the bottom of the wells were dissolved in 100 µl DMSO (incubated 10 minutes, 37 °C), followed by colorimetric read-out using a FLUOstar Omega microplate reader (BMG LABTECH) (120 sec. shaking, readout at 570 nm, blank corrected).

The wavelength dependent cell proliferation assay of **nOJ** on HeLa cells was performed as n = 6. $EC_{50}$-values for Wavelength dependent cell proliferation assay are reported in table S7.

**Table S7:**

| conditions (25 ms/0.5s) | **dark** | **410 nm** | **440 nm** | **477 nm** | **505 nm** | **525 nm** | **565 nm** |
|---|---|---|---|---|---|---|---|
| $EC_{50}$ (**nOJ**) [µM] | N/A | 0.66 | 0.37 | 0.91 | N/A | N/A | N/A |

##### Presto Blue viability assay:

Presto Blue reagent (Thermo Fisher #A13262) was added to MDA-MB-231 mCherry Lifeact cells at the end of the wound healing assay (10 µl/well) and incubated for 30 minutes.

[2] M. Borowiak, F. Küllmer, F. Gegenfurtner, S. Peil, V. Nasufovic, S. Zahler, O. Thorn-Seshold, D. Trauner, H.-D. Arndt, *Journal of the American Chemical Society* **2020**, *142*, 9240-9249.
[3] M. Borowiak, W. Nahaboo, M. Reynders, K. Nekolla, P. Jalinot, J. Hasserodt, M. Rehberg, M. Delattre, S. Zahler, A. Vollmaer, D. Trauner, O. Thorn-Seshold, *Cell* **2015**, *162*, 403–411.

Fluorescence readout ($\lambda_{Ex}$ 544 nm/ $\lambda_{Em}$ 590 nm) was performed using a FLUOstar Omega microplate reader (BMG LABTECH). Values were blank corrected and viability is reported as % of control.

**For statistical analysis and graphical representation**, GraphPad Prism 9 for macOS (San Diego, CA, USA) was used. The absorbance values for untreated controls (cosolvent only) were normalized to 100%. Viability was reported as mean percentage of viable cells relative to control. ± standard deviation (SD) was reported from 3 independent experiments, performed in triplicates (3x3). $EC_{50}$-values were determined by four-parameter curve fitting for sigmoidal dose-response with a variable slope.

When 50% cell viability was not reached, $EC_{50}$ values were indicated at >X uM, with X being the highest concentration tested.

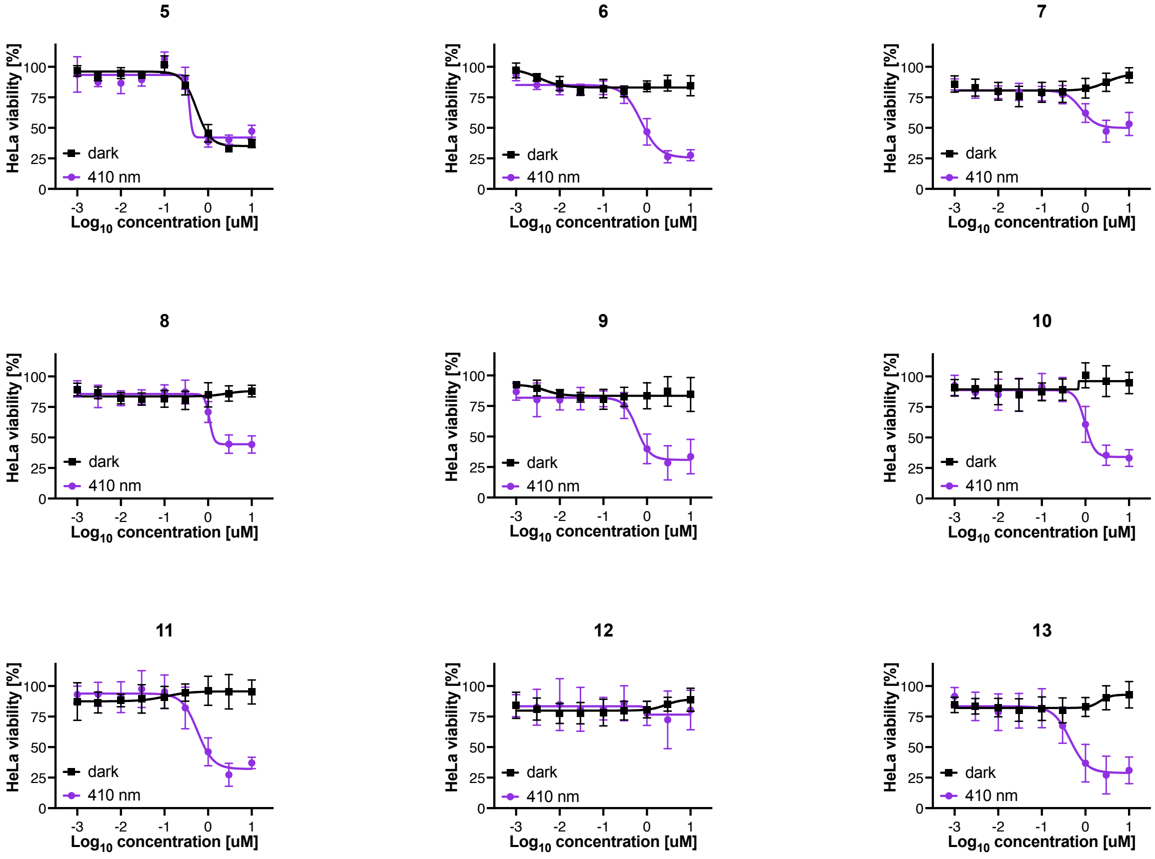

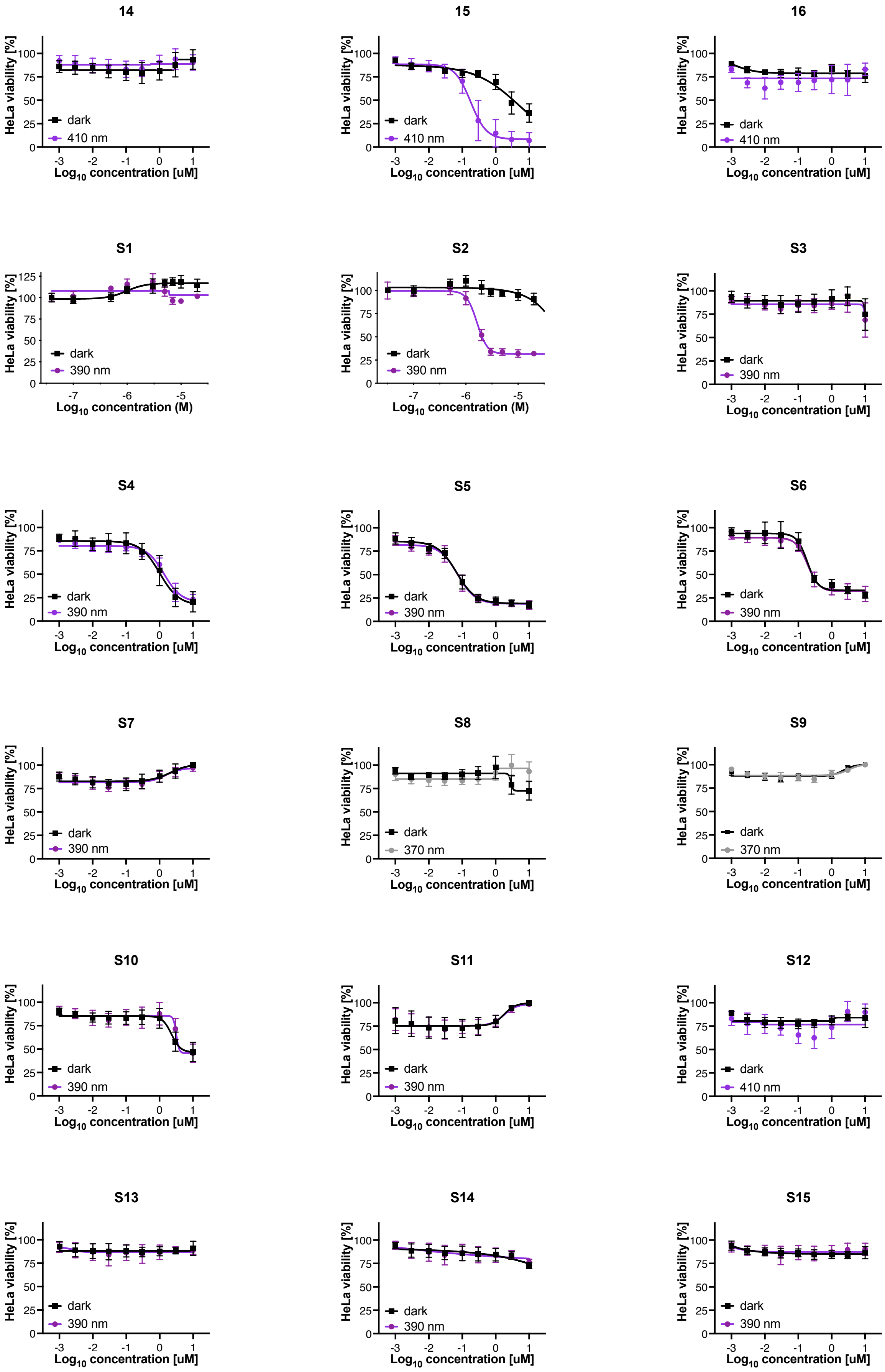
14
15
16
S1
S2
S3
S4
S5
S6
S7
S8
S9
S10
S11
S12
S13
S14
S15
HeLa viability [%]
Log10 concentration [uM]
Log10 concentration (M)
dark
410 nm
390 nm
370 nm

**S16**

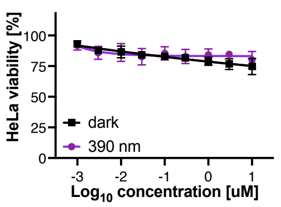

**S17**

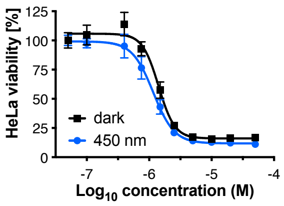

**S18**

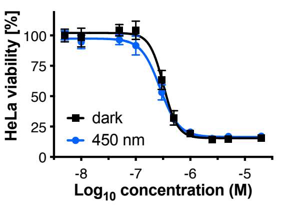

**S19**

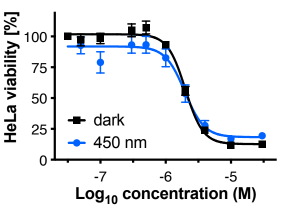

**S20**

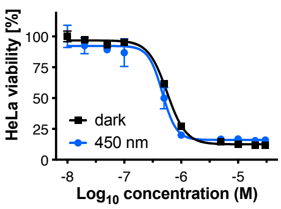

**S21**

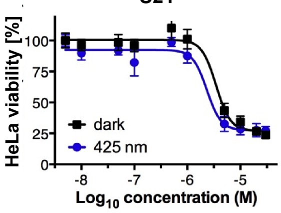

**S22**

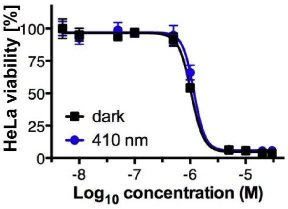

**S23**

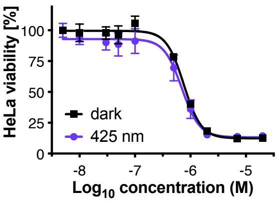

**S24**

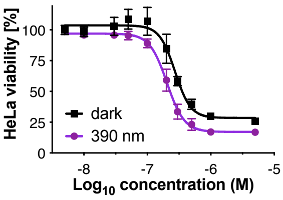

**S25**

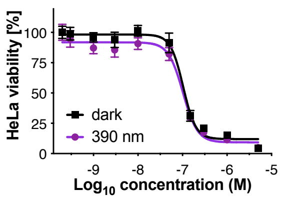

**S26**

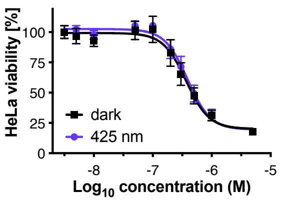

**S27**

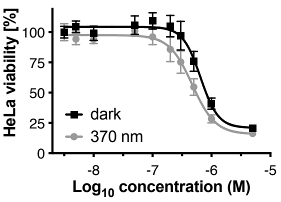

**S28**

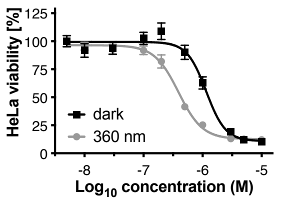

**S29**

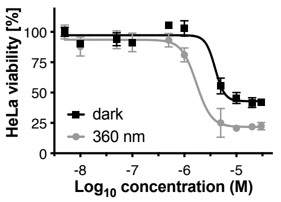

**S30**

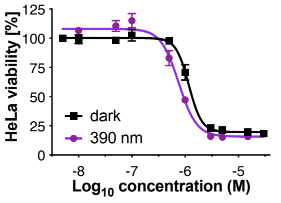

**S31**

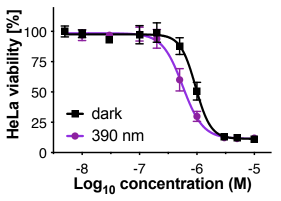

**S32**

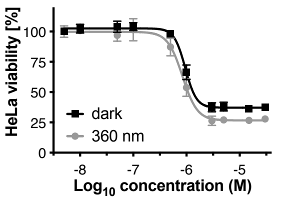

**S33**

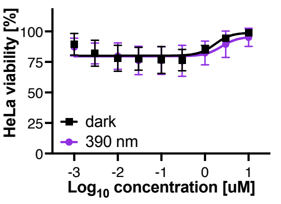

S34

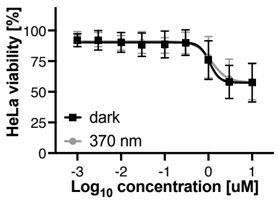

S35

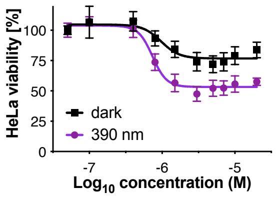

S36

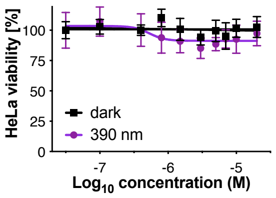

S37

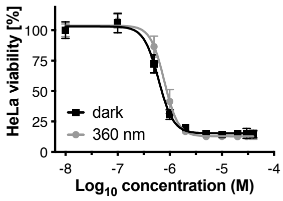

S38

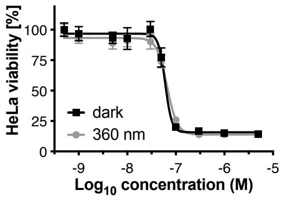

S39

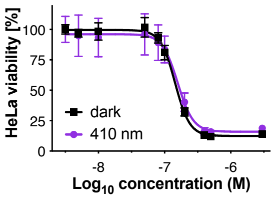

S40

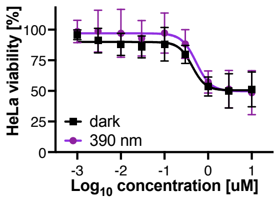

S41

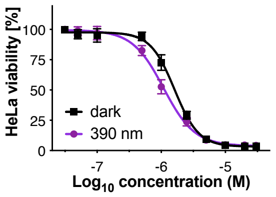

S42

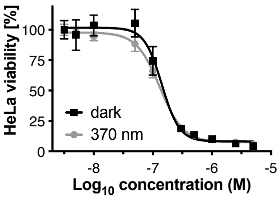

S43

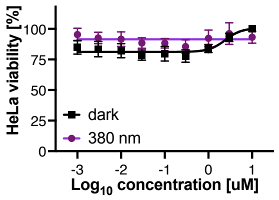

S44

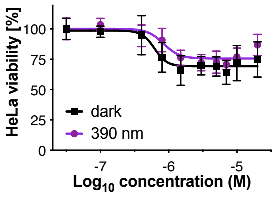

S45

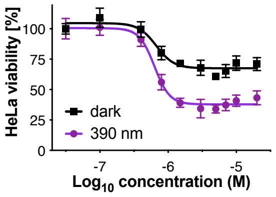

S46

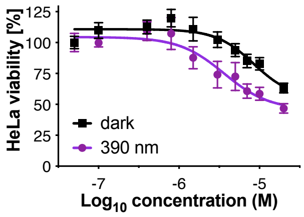

S47

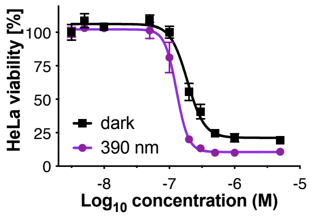

#### *In vitro* Wound Healing Assays

MDA-MB-231 cells were plated in 96 well-plates at a concentration of 40k cells/well in 100 µl phenolred-free DMEM (Thermo Fisher cat# 31053036), supplemented with 10% FBS, 1% penicillin-streptomycin and a final concentration of 4 mM L-glutamine. The next day, cells were starved over night by exchanging the medium for phenolred-free DMEM (Thermo Fisher cat# 31053036), supplemented with 1% FBS (Gibco, Thermo Fischer Cat# 10437036), 1% penicillin-streptomycin-glutamine (Gibco, Thermo Fischer cat# 10378016) and a final concentration of 4 mM L-glutamine (Gibco, Thermo Fischer cat# 25030081). After a fully confluent monolayer was obtained, the monolayer was scratched using a sterilized wooden toothpick, the medium was removed, the cells washed with PBS (pH 7.4, Gibco, Thermo Fischer cat# 10010023; 3 x 100 µl, 37 °C) and 90 µl full growth medium was added. The obtained scratches were imaged using a Leica DMI6000B inverted fluorescent microscope with a Tokai Hit stage-top incubator at 37 °C at 2.5 x magnification, DIC filter (timepoint t=0), before the compounds were added (10 x concentration, 10 µl). Experiments were run in sets of two (or three) duplicates (dark/ light (/ rescue) protocol) using the cell DISCO system with a 24-LED array as previously reported. Quantification of scratch area was performed using a custom macro script for ImageJ (National Institutes of Health, USA). The final selection area was adjusted to fit the wound in every image of the image-stack. Relative wound closure per well was referenced to timepoint $t_0$ and further analyzed using GraphPad Prism 9 for macOS (San Diego, CA, USA). Three independent experiments were combined by internal normalization to the wound closure (dark) after 24 h. Statistical analysis was performed using GraphPad Prism 9 for macOS (San Diego, CA, USA). Error bars are reported as ± standard deviation (SD) from 3 independent experiments (n = 5-6).

The wound closure was monitored for a total of 48 h. After this time period, cell viability was assessed using PrestoBlue viability assay (Thermo Fischer cat# A13261; 10 µl/ well), incubated for 30 minutes and analyzed by fluorescence readout $\lambda_{Ex}$ 544 nm/ $\lambda_{Em}$ 590 nm using a FLUOstar Omega microplate reader (BMG LABTECH, Ortenberg, Germany).

**Macro:**

```
run("Images to Stack", "name=Stack title=[] use");
run("Find Edges", "stack");
setOption("BlackBackground", true);
run("Make Binary", "method=Default background=Default calculate black");
run("Dilate", "stack");
run("Fill Holes", "stack");
run("Dilate", "stack");
```

```
run("Fill Holes", "stack");
run("Invert", "stack");
makeRectangle(657, 753, 768, 576);
```

**Fixed Cell Fluorescent Staining**

Glass coverslips (ø12 mm, thickness #1; VWR cat# 89167-106) were etched with HCl (1 M, aq.) at 60 °C with frequent, gentle agitation for 10 h, rinsed dd $H_2O$ (10 x), 70% EtOH (aq., 3 x) and stored under EtOH (absolut). The coverslips were flamed and upon cooling, treated with 40 µl of poly-L-lysine (Sigma Aldrich cat# P4707-50ML) for 1 h at 37 °C. the coverslips were washed with dd $H_2O$ (3 x) and dried at 37 °C for at least 4 h or over night. The coverslips were stored at +4 °C for up to 3 months.

In a 24 well plate, HeLa cells were seeded on poly-L-lysine coated coverslips at a density of 40 000 cells/ 450 µl phenol red free full growth medium. The next morning, the test compound was added as a 10 x concentration (50 µl, 1% DMSO, 2% MeCN, DMEM) in the dark. Two well-plates were subjected to the respective irradiation conditions for 5 h, while one was kept in the dark. After the irradiation period, one of the irradiated plates was kept in the dark for an additional 16 h, and the two remining plates (one light and one dark) were fixed and blocked immediately under exclusion of ambient light.

Fixing/ Blocking: Under red light, medium was removed and MTSB Buffer (190 µl, 37 °C) was added. After 30 seconds, glutaraldehyde (10% aq., 10 µl) was added and the cells were incubated for 10 minutes. The buffer was removed and $NaBH_4$ (0.1% in PBS, 200 µl) was added and incubated for 7 minutes. $NaBH_4$/PBS was removed and the fixed cells were washed with PBS (3 x 500 µl, 37 °C) and blocked with FBS (10% in PBS, 500 µl, 37 °C) over night at +4 °C or for 3 h at room temperature.

Staining: Coverslips were placed on a drop (40 µl) of a mixture of Phallodin AF 594 (abcam cat# ab176757; dilution 1:1000 – 1:1500.), anti alpha Tubulin AF 488 (abcam cat# ab195887; ca. 1 µg/ml) and Hoechst 33342 (Thermo Fischer cat# 62249; ca. 1 µg/ml) and incubated in the dark (ensure humidity to avoid drying) for 1 h. the coverslips were washed three times with PBS, excess PBS was removed and the coverslip was mounted using fluoroshield aqueous mounting medium (abcam cat# ab104135). The mounted samples were dried at room temperature in the dark and imaged by confocal microscopy on an upright Leica SP8 laser scanning confocal within the next days. Samples were thereafter stored at +4 °C.

Image analysis and processing was performed using ImageJ and Fiji – ImageJ (National Institutes of Health, USA).[4]

#### Live-Cell Imaging

##### General Considerations:

MDA-MB-231 cells that stably express mCherry LifeAct were seeded using phenol-red free DMEM (Thermo Fisher cat# 31053036), supplemented with 10% FBS, 1% penicillin-streptomycin and a final concentration of 4 mM L-glutamine at a density of 40k cells/ 2 ml in a ø35 mm imaging dish (previously coated with poly-L-lysine; see procedure for cover slips) and left to adhere for 16-24 h. Next, the medium was removed and new medium containing optojasp (2.5 µM) was added. The cells were left to incubate for 16-24 h in the dark. When transporting cells, it was ensured that cells were kept in a warm environment and in the dark.

$EC_{50}$ (**nOJ**) in MDA-MB-231 mCherry LifeAct: 0.24 µM (n=3, N=1).

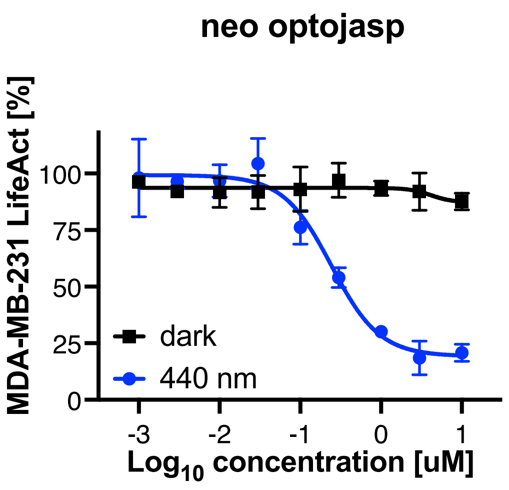

[4] J. Schindelin, I. Arganda-Carreras, E. Frise, V. Kaynig, M. Longair, T. Pietzsch, S. Preibisch, C. Rueden, S. Saalfeld, B. Schmid, J.-Y. Tinevez, D. J. White, V. Hartenstein, K. Eliceiri, P. Tomancak, A. Cardona, *Nat Methods* **2012**, *9* (7), 676–682.

**Live-Cell Imaging with Global Illumination:**

The stage-top incubator was preheated to 37 °C and equilibrated to 5% $CO_2$ using a dummy imaging dish for 60 minutes prior to imaging. The cell containing imaging dish was mounted under minimal ambient light using red head lamps for better vision. When DIC was used for focus, minimal light intensity was used. Cells were imaged using a Leica DMI6000B inverted fluorescent microscope with a TexasRed filter set and a 40x oil immersion objective. Cells were first imaged for 3 h in the dark. Next, 395 nm light pulses were applied orthogonally using a single LED (see setup picture Figure S3 that was connected to the DISCO system (25 ms pulse per 0.5 s) and images acquired every 10 minutes and then every 30 minutes. After light treatment, cells were monitored for an additional 24 h.

Image analysis and processing was performed using ImageJ and Fiji – ImageJ (National Institutes of Health, USA).[5]

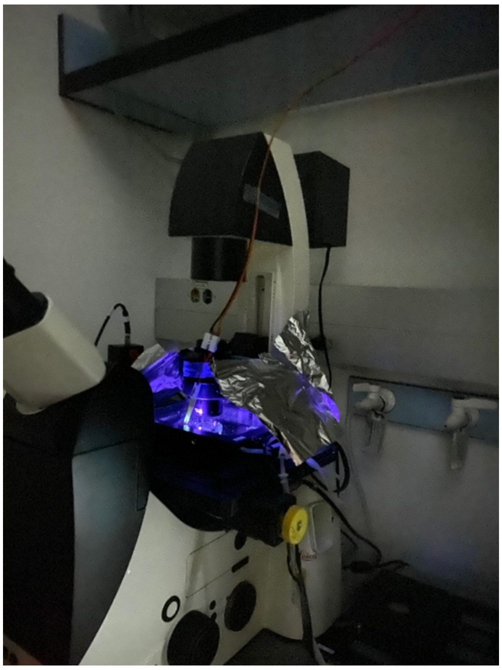

Figure S3: Setup for orthogonal light delivery

[5] J. Schindelin, I. Arganda-Carreras, E. Frise, V. Kaynig, M. Longair, T. Pietzsch, S. Preibisch, C. Rueden, S. Saalfeld, B. Schmid, J.-Y. Tinevez, D. J. White, V. Hartenstein, K. Eliceiri, P. Tomancak, A. Cardona, *Nat Methods* **2012**, *9* (7), 676–682.

**Live-Cell Imaging with Local Illumination:**

The environmental chamber of a Zeiss LSM880 Airyscan microscope system (ZEISS, Oberkochen, Germany) was preheated to 37 °C prior to imaging. Cells were transported as quickly as possible in a tightly sealed styrofoam box with 2 x 500 ml water bottles that had been equilibrated to 37 °C and cells immediately transferred to the environmental chamber of the microscope. The imaging dish was mounted on the stage under minimal ambient light and using red head lamps to allow for better vision. Cells were focused using a red filter foil in the light path oft he microscope to shield blue light from reaching the sample during the light exposure that was required to focus.

Images were acquired using a 63x objective. 440 nm light pulses and ROI were specified using the FRAP tool and different acquisition blocks were programmed in row using the experiment designer in ZEN (ZEISS, Oberkochen, Germany).

Image analysis and processing was performed using Fiji – ImageJ (National Institutes of Health, USA).[6]

[6] J. Schindelin, I. Arganda-Carreras, E. Frise, V. Kaynig, M. Longair, T. Pietzsch, S. Preibisch, C. Rueden, S. Saalfeld, B. Schmid, J.-Y. Tinevez, D. J. White, V. Hartenstein, K. Eliceiri, P. Tomancak, A. Cardona, *Nat Methods* **2012**, *9* (7), 676–682.

### Materials and Methods - Chemistry

#### Reagents and starting materials

All solvents, when not purchased in suitable purity or dryness, were distilled using standard methods,[7] or passed through alumina columns (Innovative Technology, solvent purification system): Methanol (MeOH), ethanol (EtOH) and acetonitrile (MeCN) were dehydrated by adding activated molecular sieve (3 Å) and stored under a protective gas atmosphere. Tetrahydrofuran (THF) and diethyl ether ($Et_2O$) was distilled under a $N_2$ atmosphere from Na/benzophenone before use; dichloromethane ($CH_2Cl_2$) was distilled under a $N_2$ atmosphere from $CaH_2$ before use; and other dry solvents such as toluene and dimethylformamide (DMF) obtained in HPLC quality and passed through a solvent purification system (Pure Solv, Innovative Technology, Inc., USA) by applying $N_2$ overpressure right before use. All other solvents used for work-up or column chromatography were distilled before use. The petroleum ether used had a boiling range of 40-60 ° C. Phosphate buffer (pH = 7) was prepared by dissolving $Na_3PO_4$×12 $H_2O$ (54.8 g, 0.14 mol) and $NaH_2PO_4$ (42.7 g, 0.36 mol) in water (1.0 L). Deionized water was used for all experiments.

All reagents and solvents were purchased from commercial sources (ABCR, Acros Organics, Active Scientific, Alfa Aesar, Apollo Scientific, Carbolution Chemicals, ChemPur, Fluka, Fluorochem, Manchester Organics, Merck, Sigma-Aldrich, TCI Europe and VWR) and were used without further purification unless noted otherwise. Commercial-grade ($i$Pr)$_2$NEt (VWR) was distilled under a $N_2$ atmosphere from $CaH_2$ before use. Salts were dried in a fine vacuum with a heat gun for 10-20 minutes. 2-Chlorotrityl chloride resin (100-200 mesh; 1.6 mmol·g-1) was purchased from Iris Biotech GmbH.

(*S*)-Hex-5-en-2-ol (**3**), Pent-4-en-1-ol, Fmoc-Lys(Boc)-OH and Fmoc-Orn(Boc)-OH were purchased from Sigma Aldrich. Known Fmoc-Dab(Boc)-OH and Fmoc-Dap(Boc)-OH were prepared according to Lau *et al.*[8] followed by Boc-protection using standard methods. (2S,4R)-4-Methylhex-5-en-2-ol (**S48**) was prepared according to Nasufović et al.[9] Fmoc-βTyr(TIPS)-OH, Fmoc-*N*-MeD-Trp-OH, (*S*)-2,4-Dimethyl-4-pentenoic acid and *cyclo*-[(2*S*,4*E*,8*S*)Hdn-*L*-Ala-*D*-*N*Me-Trp-*β*Tyr(OTIPS)] (**S5**) were prepared according to Tannert *et*

[7] W. L. Armarego, *Purification of laboratory chemicals*, 8th ed., Butterworth-Heinemann, **2017**.
[8] Y. H. Lau, D. R. Spring, *Synlett* **2011**, *2011*, 1917-1919.
[9] V. Nasufović, F. Küllmer, J. Bößneck, H.-M. Dahse, H. Görls, P. Bellstedt, P. Stallforth, H.-D. Arndt, *Chemistry – A European Journal* **2021**, *27*, 11633-11642.

*al.*[10] Known peptide acids *L*-Pea-*L*-Lys(Boc)-*D*-*N*Me-Trp-*β*Tyr(OTIPS)-OH (**S49**) and *L*-Pea-*L*-Dap(Boc)-*D*-*N*Me-Trp-*β*Tyr(OTIPS)-OH (**S50**), cyclodepsipeptides *cyclo*-[(2*S*,4*E*,8*S*)Hdn-*L*-Ala-*D*-*N*Me-Trp-*β*Tyr(OTIPS)] (**S5**), *cyclo*-[(2*S*,4*E*,8*S*)Hdn-*L*-Lys(Boc)-*D*-*N*Me-Trp-*β*Tyr(OTIPS)] (**S51**), *cyclo*-[(2*S*,4*E*,8*S*)Hdn-*L*-Orn(Boc)-*D*-*N*Me-Trp-*β*Tyr(OTIPS)] (**S52**), *cyclo*-[(2*S*,4*E*,8*S*)Hdn-*L*-Dab(Boc)-*D*-*N*Me-Trp-*β*Tyr(OTIPS)] (**S53**) and *cyclo*-[(2*S*,4*E*,8*S*)Hdn-*L*-Dap(Boc)-*D*-*N*Me-Trp-*β*Tyr(OTIPS)] (**S54**) were prepared according to Borowiak & Küllmer *et al.*[11] The applied azobenzene acids and azobenzenes were prepared according to Küllmer *et al.*[12] All compounds intended for biological studies were purified using Preparative HPLC to remove solvent residues and salts. Synthetic procedures are described in Supplementary Note 1.

[10] R. Tannert, L.-G. Milroy, B. Ellinger, T.-S. Hu, H.-D. Arndt, H. Waldmann, *Journal of the American Chemical Society* **2010**, *132*, 3063-3077.
[11] M. Borowiak, F. Küllmer, F. Gegenfurtner, S. Peil, V. Nasufovic, S. Zahler, O. Thorn-Seshold, D. Trauner, H.-D. Arndt, *Journal of the American Chemical Society* **2020**, *142*, 9240-9249.
[12] F. Küllmer, L. Gregor, H.-D. Arndt, Property-Selected Asymmetric Azobenzenes for Photoswitchable Ligands. *ChemRxiv Preprint:* https://doi.org/10.26434/chemrxiv-2022-fd950.

#### Chromatography

Reaction progress was monitored by TLC on precoated, Merck Silica gel 60 F254 alumina-backed plates. TLC chromatograms were first visualized by UV-A or UV-B irradiation at 254 or 320 nm, followed by staining with aqueous ninhydrin, $KMnO_4$, or ceric ammonium molybdate solution, and gentle heating. Primary and secondary amines were detected with ninhydrin (6% in EtOH).

Flash silica gel chromatography was performed using silica gel ($SiO_2$, particle size 40–63 µm) purchased from Macherey & Nagel, Düren (Germany) under a pressure of 0.3-0.5 bar.

Analytical HPLC was performed on a Shimadzu machine consisting of a LC-10AT (Pump), DGU-14AT (autosampler), CTO-10AC (column oven), C18 Gravity (precolumn), Nucleodur C18 Gravity 5 µm (column), RF-10A (fluorescence detector), SDP-10A (UV/VIS-detector) und SCL-10A (controller). Evaluation was performed with *Chromeleon* (software). The gradient was started at 10% MeCN (with 0.1% TFA) in $H_2O$ (with 0.1% TFA) with a flow of 1 mL·min-1, and the proportion of organic component was linearly increased after 1 min to 95% over a period of 10 min and then kept at that ratio for a period of 9 min (*method A*). Alternatively the gradient was started at 50% MeCN (*method B*). Thirdly, a gradient was started at 10% MeCN (with 0.1% TFA) in $H_2O$ (with 0.1% TFA) with a flow of 1 mL·min-1, and the proportion of organic component linearly increased to 95% over a period of 10 min and then kept at that ratio for a period of 5 min (*method C*). Alternatively, LC-MS analysis was performed on a Shimadzu (HPLC) consisting of a DGU-14A (degasser), LC-10AT VP (pumps), SIL-20AT VP (autosampler), CTO-10AC VP (column oven), C18 Gravity (precolumn), Nucleodur C18 ISIS 3µm (column), SPD-10A VP (UV/VIS-detector), SCL-10A VP (Controller), Acurate Post-Column Splitter (splitter), fitted to a Finnigan LCQ (ESI-MS). The gradient was started at 10% MeCN (with 1.0% AcOH) in $H_2O$ (with 1.0% AcOH) with a flow of 1 mL·min-1, and the proportion of organic component was linearly increased to 100% over a period of 10 min and then kept at that proportion for a period of 5 min (*method A*). Alternatively the gradient was started at 30% MeCN (*method B*) or was started at 50% MeCN (*method C*). In some cases, *E*- and *Z*-azobenzene isomers were clearly separated. Traces of analytical HPLC analyses of final compounds are reproduced in Supplementary Note 2.

Preparative HPLC purification was performed on a Varian device consisting of a ProStar 215 (pump), ProStar 340 (UV/VIS-detector) and a ProStar 701 (collector). A VP250/21 Nucleodur

C18 Gravity 5µm was used as column. As mobile phase a gradient of water and acetonitrile was applied.

#### NMR spectra

$^{1}$H- and $^{13}$C-NMR spectra were recorded on Bruker Avance I 250 (250 MHz ($^{1}$H) and 63 Hz ($^{13}$C)), Bruker Fourier 300 (300 MHz ($^{1}$H) and 75 Hz ($^{13}$C)), Bruker Avance III 400 (400 MHz ($^{1}$H) and 100 MHz ($^{13}$C)), Bruker Avance III HD 500 (500 MHz ($^{1}$H) and 100 MHz ($^{13}$C)), and Bruker AC 600 (600 MHz ($^{1}$H) and 150 MHz ($^{13}$C)). Chemical shifts are expressed in parts per million (ppm) and the spectra are calibrated to residual solvent signals of $CDCl_3$ (7.26 ppm ($^{1}$H) and 77.0 ppm ($^{13}$C)), DMSO (2.50 ppm ($^{1}$H) and 39.43 ppm ($^{13}$C)), respectively. Coupling constants are given in Hertz (Hz) and the following notations indicate the multiplicity of the signals: s (singlet), d (doublet), t (triplet), q (quartet), qui (quintet), sxt (sextet), m (multiplet), br (broad signal). Owing to the presence of *E* and *Z* isomers of some compounds containing an azobenzene functionality, more signals were observed in the $^{1}$H and $^{13}$C spectra than would be expected for a single isomer, depending on illumination history. Signals for the major *E* isomer are reported. Signal identity and peak assignments were verified by 2D-NMR experiments (COSY, TOCSY, HSQC and HMBC) whenever necessary. NMR spectra are displayed in Supplementary Note 3.

#### UV–Vis spectra and Polarimetry

UV-Vis spectra were recorded using a JASCO V-730 UV–Visible Spectrophotometer with Helma SUPRASIL precision cuvettes (5 mm light path). All compounds were dissolved as 10 mM stock solutions in DMSO and diluted in PBS-Puffer/MeCN (2:1) solutions. Switching was achieved by using a LED at the indicated wavelength. UV-spectra of all compounds are reproduced in Supplementary Note 4.

Optical rotations were measured in a Jasco Polarimeter P 2000 at 589 nm, with values given in $deg \cdot cm^{2} \cdot g^{-1}$ and concentrations c given in g/100mL.

##### Infrared (IR) spectra

Fourier transform infrared spectroscopy (FT-IR) spectra were obtained with an IR-Affinity-1 from Shimadzu (ATR, neat or as a thin film). Wave numbers are reported in $cm^{-1}$.

##### Low- and high-resolution ESI mass spectrometry

Low- and high-resolution ESI mass spectra were obtained on a Thermo-Finnigan LCQ (LR) and Bruker Maxis Impact (HR) spectrometers operating in either positive or negative ionization modes, respectively, fitted to Shimadzu AL-10 (LR-ESI-MS) or Dionex Ultima 3000 HPLC systems (HR-ESI-MS).

### Supplementary Note 1: Chemical Synthesis and Characterisation

Standard Procedures:

Esterification with EDCI (**SP1**):

Under an atmosphere of nitrogen, DIPEA (2.0 – 4.0 equiv.) was added dropwise to a solution of peptide acid (1.0 equiv.), alcohol (4.0 – 8.0 equiv.) and DMAP (4.0 equiv.) in anhydrous $CH_2Cl_2$ (40 mL/mmol) and stirred for 10 minutes at room temperature. Solid EDCI (2.0 equiv.) was added and the reaction solution was stirred for 16 to 38 hours (TLC control). Afterwards, saturated $NH_4Cl$ solution (~ 20 mL/mmol) and EtOAc (~ 30 mL/mmol) were added to the solution. The organic phase was separated off and the aqueous phase was extracted with EtOAc (3 × ~ 20 mL/mmol). The combined organic phases were dehydrated with $Na_2SO_4$, filtered, and concentrated under reduced pressure. Purification by silica gel chromatography gave pure diene.

Ring-closing metathesis (**SP2**):

Under an atmosphere of argon, diene (1.0 equiv.) was dissolved in anhydrous $CH_2Cl_2$ (1000 mL/mmol) and heated to reflux for 30 min. Grubbs catalyst second-generation (0.08 – 0.12 equiv.), dissolved in anhydrous $CH_2Cl_2$ (1.0 mL), was added to the solution and the mixture was stirred under reflux for a further 3 h. After cooling, the solvent was concentrated under reduced pressure. Purification by silica gel chromatography gave pure cyclodepsipeptide.

TIPS deprotection with hydrogen fluoride (**SP3**):

Under an atmosphere of argon, cyclodepsipeptide (1.0 equiv.) was dissolved in a micro reaction vessel (2.0 mL, PP) in a mixture of hydrogen fluoride-pyridine complex (70%) and anhydrous THF (1:17, 40 mL/mmol) and stirred for 18 h at room temperature. $SiO_2$ was added to the solution and the mixture was stirred for a further 30 min before the solvent was evaporated in a stream of nitrogen. The crude product was either converted further directly or was purified by preparative HPLC to give pure cyclodepsipeptide.

Optojasp synthesis (**SP4**):

Under an atmosphere of argon, cyclodepsipeptide (1.0 equiv.) was dissolved in anhydrous $CH_2Cl_2$ (20 mL/mmol) and cooled to 0 °C (ice). HCl in dioxane (4M, 20 mL/mmol) was added and stirring was continued for 2.5 h. The solvent was removed under reduced pressure and the residue was dried in fine vacuum.

The received colourless solid was mixed with azobenzenic acid (1.0 – 1.2 equiv.) and HATU (2.4 equiv.) and dissolved in anhydrous THF (40 mL/mmol) under an atmosphere of argon. After the addition of DIPEA (8.0 equiv.) the reaction mixture was stirred for 4 h at room temperature. The solution was concentrated in a stream of nitrogen and was roughly purified by silica gel chromatography ($CH_2Cl_2$/acetone, 1: 0 → 1: 2, + 0 – 10% MeOH).

The obtained product was transferred to a micro reaction vessel (2.0 mL, PP), dissolved in a mixture of hydrogen fluoride-pyridine complex (70%) and anhydrous THF (1:17, 40 mL/mmol) under an atmosphere of argon and stirred at room temperature for 18 h. $SiO_2$ (5.0 mg/mL) was added to the solution and the mixture was stirred for a further 30 min before the solvent was evaporated in a stream of nitrogen. Purification by silica gel chromatography gave pure optojasp. For the biological studies, all compounds were additionally purified by preparative HPLC.

Reductive amination (**SP5**):

To a solution of amine (1.0 equiv.) in a mixture of AcOH and EtOH (1:9, 25 mL/mmol) aldehyde (3.0 equiv.) was added and stirred for 2 hours at room temperature. The mixture was cooled to 0 °C (icebath) and $NaCNBH_3$ (4.5 equiv.) was added. The icebath was removed and stirring was continued until TLC indicated satisfactory conversion (16 – 38 h). The solution was neutralized with aqueous NaOH (2 M) and most of the EtOH was removed under reduced pressure followed by extraction of the aqueous phase with $CHCl_3$ (3 × 20 mL/mmol). The combined organic extracts were dried with $Na_2SO_4$, filtered, and concentrated under reduced pressure. Purification by silica-gel chromatography provided the pure azobenzene.

Ester cleavage with LiOH (**SP6**):

The ester (1.0 equiv.) was dissolved in THF (20 mL/mmol), treated with aqueous LiOH (2 M, 10.0 equiv.), and stirred for 2 h at room temperature. The solution was neutralized with aqueous HCl (1 M). Most of the THF was removed under reduced pressure and $CHCl_3$ (20 mL/mmol) was added. The organic layer was separated followed by extraction of the aqueous layer with $CHCl_3$ or EtOAc (3 – 12 × 20 mL/mmol). The combined organic extracts were dried with $Na_2SO_4$, filtered, and concentrated under reduced pressure.

**Synthesis of optojasp:**

*cyclo*-[(2*S*,4*E*,8*S*)-Hdn-L-Dap(2-(4´-((3´´,4´´-dihydro-2´´*H*-benzo[*b*][1´´,4´´]oxazin-7´´-yl)diazenyl)phenoxy)acetyl)-D-*N*MeTrp-L-*β*Tyr]; (**5**)

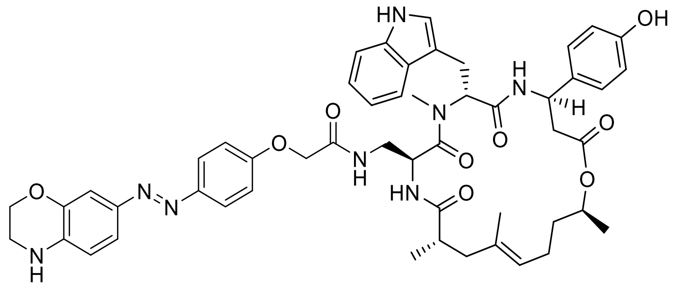

Standard Procedure **SP4** with cyclodepsipeptide **S54** (2.5 mg, 2.8 µmol, 1.0 equiv.), 2-(4´-((3´´,4´´-dihydro-2´´*H*-benzo[b][1´´,4´´]oxazin-7´´-yl)diazenyl)phenoxy) acetic acid (1.5 mg, 3.4 µmol, 1.2 equiv.), HATU (2.6 mg, 6.8 µmol, 2.4 equiv.) and DIPEA (3.8 µl, 22.5 µmol, 8.0 equiv.) gave conjugate **5** (1.4 mg, 54%) as a red solid after purification by preparative HPLC ($H_2O$/MeCN, 60:40 → 0:100). $R_f$ = 0.31 ($CH_2Cl_2$/MeOH, 14:1); $[\alpha]_D^{24}$ = 56.1 (c = 0.05 in MeCN); $^1$H-NMR (400 MHz, DMSO-$d_6$): $\delta$ = 10.78 (d, *J* = 1.5 Hz, 1 H), 9.27 (s, 1 H), 8.54 (d, *J* = 8.8 Hz, 1 H), 7.91 (t, *J* = 6.1 Hz, 1 H), 7.71 (d, *J* = 9.1 Hz, 2 H), 7.67 (d, *J* = 7.9 Hz, 1 H), 7.58 (d, *J* = 8.8 Hz, 1 H), 7.33 (dd, *J* = 2.3, 8.5 Hz, 1 H), 7.30 (d, *J* = 8.2 Hz, 1 H), 7.17 (d, *J* = 2.0 Hz, 1 H), 7.07 - 6.99 (m, 6 H), 6.95 (t, *J* = 7.3 Hz, 1 H), 6.76 (br s, 1 H), 6.67 (dd, *J* = 2.0, 8.5 Hz, 3 H), 5.49 (dd, *J* = 7.3, 8.8 Hz, 1 H), 5.17 (dt, *J* = 3.5, 9.5 Hz, 1 H), 4.98 - 4.88 (m, 2 H), 4.69 (sxt, *J* = 6.3 Hz, 1 H), 4.42 (s, 2 H), 4.15 (t, *J* = 4.4 Hz, 2 H), 3.38 (t, *J* = 4.1 Hz, 2 H), 3.11 (s, 3 H), 3.07 - 2.96 (m, 2 H), 2.85 - 2.75 (m, 1 H), 2.70 - 2.53 (m, 4 H), 2.16 (dd, *J* = 11.5, 14.2 Hz, 1 H), 1.85 (spt, *J* = 6.7 Hz, 2 H), 1.76 (d, *J* = 14.3 Hz, 1 H), 1.49 (s, 3 H), 1.55 - 1.45 (m, 1 H), 1.44 - 1.31 (m, 1 H), 1.14 (d, *J* = 6.1 Hz, 3 H), 0.93 (d, *J* = 7.0 Hz, 3 H); $^{13}$C-NMR, HSQC/HMBC (100 MHz, DMSO-$d_6$): $\delta$ = 175.3, 170.6, 170.4, 169.6, 167.9, 159.1, 156.5, 147.2, 143.2, 139.0, 136.4, 133.1, 127.4, 123.9, 123.7, 121.1, 120.4, 119.0, 118.3, 115.5, 115.2, 113.5, 111.5, 109.8, 108.1, 71.2, 67.2, 64.4, 55.4, 49.2, 48.4, 43.1, 41.6, 40.0, 38.2, 35.2, 30.7, 25.3, 23.8, 19.8, 19.3, 17.2; UV-VIS (PBS/MeCN, 2:1): $\lambda_{max}$ ($\varepsilon$) = 414 nm (24.9 × $10^3$ l·mol$^{-1}$·cm$^{-1}$); HRMS (ESI): *m/Z* calcd for $C_{51}H_{59}N_8O_9$ $[M+H]^+$: 927.440, found: 927.440.

*cyclo*-[(2*S*,4*E*,8*S*)-Hdn-L-Dap(2-(4′-((4′′-benzyl-3′′,4′′-dihydro-2′′*H*-benzo[*b*][1′′,4′′]-oxazin-7′′-yl)diazenyl)phenoxy)acetyl)-D-*N*MeTrp-L-*β*Tyr]; neo Optojasp/nOJ (**6**)

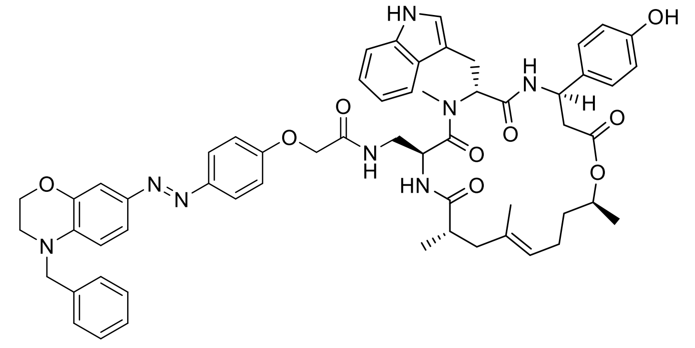

Standard Procedure **SP4** with cyclodepsipeptide **S54** (10.0 mg, 11.3 µmol, 1.0 equiv.), 2-(4′-((4′′-benzyl-3′′,4′′-dihydro-2′′*H*-benzo[b][1′′,4′′]oxazin-7′′-yl)diazenyl)phenoxy) acetic acid (4.5 mg, 11.3 µmol, 1.0 equiv.), HATU (10.3 mg, 27.0 µmol, 2.4 equiv.) and DIPEA (15.3 µl, 90.1 µmol, 8.0 equiv.) gave conjugate **6** (9.5 mg, 83%) as an orange solid after purification by silica-gel chromatography ($CH_2Cl_2$/acetone, 2:1 → 1:2). $R_f$ = 0.39 ($CH_2Cl_2$/MeOH, 14:1); $[\alpha]_D^{24}$ = 23.7 (c = 0.10 in MeCN); $^1$H-NMR (400 MHz, DMSO-$d_6$): $\delta$ = 10.80 (d, $J$ = 1.8 Hz, 1 H), 9.30 (br s, 1 H), 8.58 (d, $J$ = 8.8 Hz, 1 H), 7.95 (t, $J$ = 6.0 Hz, 1 H), 7.72 (d, $J$ = 9.1 Hz, 2 H), 7.68 (d, $J$ = 7.6 Hz, 1 H), 7.61 (d, $J$ = 8.8 Hz, 1 H), 7.40 - 7.27 (m, 7 H), 7.23 (d, $J$ = 2.0 Hz, 1 H), 7.06 (d, $J$ = 8.8 Hz, 2 H), 7.04 - 7.00 (m, 4 H), 6.95 (ddd, $J$ = 1.0, 6.9, 7.6 Hz, 1 H), 6.80 (d, $J$ = 8.8 Hz, 1 H), 6.67 (d, $J$ = 8.5 Hz, 2 H), 5.50 (dd, $J$ = 7.0, 9.1 Hz, 1 H), 5.18 (dt, $J$ = 3.4, 9.6 Hz, 1 H), 4.99 - 4.90 (m, 2 H), 4.66 (s, 2 H), 4.69 (sxt, $J$ = 6.4 Hz, 1 H), 4.43 (br. s, 2 H), 4.28 (t, $J$ = 4.4 Hz, 2 H), 3.55 (t, $J$ = 4.4 Hz, 2 H), 3.12 (s, 3 H), 3.08 - 2.97 (m, 2 H), 2.83 - 2.73 (m, 1 H), 2.72 - 2.54 (m, 4 H), 2.16 (dd, $J$ = 11.7, 14.0 Hz, 1 H), 1.85 (spt, $J$ = 7.0 Hz, 2 H), 1.76 (d, $J$ = 14.3 Hz, 1 H), 1.49 (s, 3 H), 1.55 - 1.44 (m, 1 H), 1.44 - 1.33 (m, 1 H), 1.15 (d, $J$ = 6.1 Hz, 3 H), 0.93 (d, $J$ = 6.7 Hz, 3 H); $^{13}$C-NMR (126 MHz, DMSO-$d_6$): $\delta$ = 175.4, 170.7, 170.5, 170.0, 168.1, 159.4, 156.7, 147.3, 143.9, 143.6, 138.9, 138.0, 136.6, 133.5, 133.2, 129.1, 127.5, 127.4, 127.4, 124.0, 123.8, 123.7, 121.3, 121.3, 119.2, 118.5, 115.6, 115.5, 115.1, 111.6, 111.5, 110.0, 107.3, 71.2, 67.4, 64.3, 55.3, 54.0, 49.4, 48.6, 47.6, 43.2, 41.9, 40.7, 38.3, 35.4, 31.0, 25.7, 24.1, 20.0, 19.8, 17.2; IR: $\tilde{\nu}_{max}$ = 3348, 2932, 1728, 1667, 1601, 1516, 1454, 1319, 1242, 1157, 1053, 837, 745, 652; UV-VIS (PBS/MeCN, 2:1): $\lambda_{max}$ ($\varepsilon$) = 448 nm (22.1 × $10^3$ l·mol$^{-1}$·cm$^{-1}$); HRMS (ESI): $m/Z$ calcd for $C_{58}H_{65}N_8O_9$ $[M+H]^+$: 1017.487, found: 1017.487.

*cyclo*-[(2*S*,4*E*,8*S*)-Hdn-L-Dap(2-(4´-((4´´-(4´´´-methoxybenzyl)-3´´,4´´-dihydro-2´´*H*-benzo[*b*][1´´,4´´]oxazin-7´´-yl)diazenyl)phenoxy)acetyl)-D-*N*MeTrp-L-*β*Tyr]; (**7**)

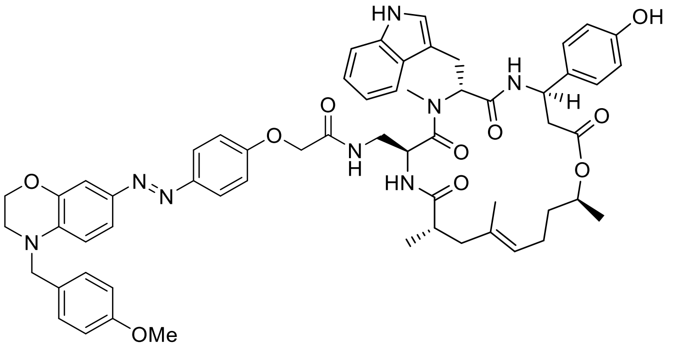

Standard Procedure **SP4** with cyclodepsipeptide **S54** (2.5 mg, 2.8 µmol, 1.0 equiv.), azobenzene **S55** (1.5 mg, 3.4 µmol, 1.2 equiv.), HATU (2.6 mg, 6.8 µmol, 2.4 equiv.) and DIPEA (3.8 µl, 22.5 µmol, 8.0 equiv.) gave conjugate **7** (2.2 mg, 75%) as a red solid after purification by preparative HPLC ($H_2O$/MeCN, 60:40 → 0:100). $R_f$ = 0.39 ($CH_2Cl_2$/MeOH, 14:1); $[\alpha]_D^{24}$ = 60.8 (c = 0.10 in MeCN); $^1$H-NMR (400 MHz, DMSO-$d_6$): $\delta$ = 10.78 (d, *J* = 1.8 Hz, 1 H), 9.28 (s, 1 H), 8.55 (d, *J* = 8.8 Hz, 1 H), 7.92 (t, *J* = 6.0 Hz, 1 H), 7.72 (d, *J* = 9.1 Hz, 2 H), 7.68 (d, *J* = 7.9 Hz, 1 H), 7.58 (d, *J* = 8.8 Hz, 1 H), 7.37 (dd, *J* = 2.2, 8.6 Hz, 1 H), 7.30 (d, *J* = 8.2 Hz, 1 H), 7.25 (d, *J* = 8.8 Hz, 2 H), 7.22 (d, *J* = 2.3 Hz, 1 H), 7.09 - 7.00 (m, 6 H), 6.96 (d, *J* = 7.6 Hz, 1 H), 6.92 (d, *J* = 8.8 Hz, 2 H), 6.86 (d, *J* = 8.8 Hz, 1 H), 6.67 (d, *J* = 8.5 Hz, 2 H), 5.50 (dd, *J* = 7.1, 9.0 Hz, 1 H), 5.18 (ddd, *J* = 3.5, 8.8, 10.8 Hz, 1 H), 4.99 - 4.89 (m, 2 H), 4.69 (sxt, *J* = 6.4 Hz, 1 H), 4.58 (s, 2 H), 4.43 (s, 2 H), 4.25 (t, *J* = 4.2 Hz, 2 H), 3.74 (s, 3 H), 3.51 (t, *J* = 4.4 Hz, 2 H), 3.12 (s, 3 H), 3.08 - 2.98 (m, 2 H), 2.85 - 2.75 (m, 1 H), 2.71 - 2.57 (m, 3 H), 2.55 - 2.53 (m, 1 H), 2.17 (dd, *J* = 11.7, 13.7 Hz, 1 H), 1.86 (spt, *J* = 6.7 Hz, 2 H), 1.76 (d, *J* = 14.3 Hz, 1 H), 1.50 (s, 3 H), 1.55 - 1.45 (m, 1 H), 1.44 - 1.33 (m, 1 H), 1.15 (d, *J* = 6.4 Hz, 3 H), 0.94 (d, *J* = 6.7 Hz, 3 H); $^{13}$C-NMR, HSQC/HMBC (100 MHz, DMSO-$d_6$): $\delta$ = 175.2, 170.5, 169.8, 167.9, 159.2, 158.8, 156.5, 147.3, 143.5, 138.8, 138.3, 136.2, 133.0, 129.5, 128.9, 128.6, 127.3, 126.8, 123.8, 123.8, 123.5, 121.2, 120.9, 119.0, 118.2, 115.5, 115.3, 114.2, 111.4, 111.3, 109.8, 107.2, 70.9, 67.3, 64.3, 55.6, 55.0, 53.2, 49.2, 48.3, 47.1, 43.0, 41.6, 38.2, 35.2, 30.8, 29.3, 25.4, 23.8, 19.6, 19.3, 16.7; UV-VIS (PBS/MeCN, 2:1): $\lambda_{max}$ ($\varepsilon$) = 448 nm (20.6 × $10^3$ l·mol$^{-1}$·cm$^{-1}$); HRMS (ESI): *m/Z* calcd for $C_{59}H_{67}N_8O_{10}$ $[M+H]^+$: 1047.497, found: 1047.498.

*cyclo*-[(2*S*,4*E*,8*S*)-Hdn-L-Dap(2-(4´-((4´´-(4´´´-chlorbenzyl)-3´´,4´´-dihydro-2´´*H*-benzo-[*b*][1´´,4´´]oxazin-7´´-yl)diazenyl)phenoxy)acetyl)-D-*N*MeTrp-L-*β*Tyr]; (**8**)

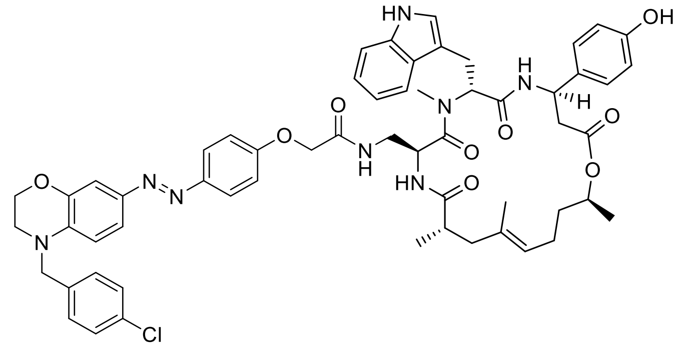

Standard Procedure **SP4** with cyclodepsipeptide **S54** (2.5 mg, 2.8 µmol, 1.0 equiv.), azobenzene **S56** (1.5 mg, 3.4 µmol, 1.2 equiv.), HATU (2.6 mg, 6.8 µmol, 2.4 equiv.) and DIPEA (3.8 µl, 22.5 µmol, 8.0 equiv.) gave conjugate **8** (1.7 mg, 57%) as a red solid after purification by preparative HPLC ($H_2O$/MeCN, 60:40 → 0:100). $R_f$ = 0.39 ($CH_2Cl_2$/MeOH, 14:1); $[\alpha]_D^{24}$ = 53.3 (c = 0.10 in MeCN); $^1$H-NMR (400 MHz, DMSO-$d_6$): $\delta$ = 10.78 (d, *J* = 1.5 Hz, 1 H), 9.28 (s, 1 H), 8.55 (d, *J* = 8.8 Hz, 1 H), 7.92 (t, *J* = 5.8 Hz, 1 H), 7.73 (d, *J* = 9.1 Hz, 2 H), 7.68 (d, *J* = 7.9 Hz, 1 H), 7.58 (d, *J* = 8.8 Hz, 1 H), 7.42 (d, *J* = 8.5 Hz, 2 H), 7.38 - 7.33 (m, 3 H), 7.30 (d, *J* = 8.2 Hz, 1 H), 7.23 (d, *J* = 2.3 Hz, 1 H), 7.08 - 7.00 (m, 6 H), 6.95 (t, *J* = 7.6 Hz, 1 H), 6.78 (d, *J* = 8.8 Hz, 1 H), 6.67 (d, *J* = 8.5 Hz, 2 H), 5.50 (dd, *J* = 7.3, 8.5 Hz, 1 H), 5.18 (ddd, *J* = 3.6, 9.4, 10.0 Hz, 1 H), 4.99 - 4.89 (m, 2 H), 4.69 (sxt, *J* = 6.1 Hz, 1 H), 4.65 (s, 2 H), 4.43 (d, *J* = 0.9 Hz, 2 H), 4.28 (t, *J* = 4.4 Hz, 2 H), 3.55 (t, *J* = 4.4 Hz, 2 H), 3.12 (s, 3 H), 3.09 - 2.97 (m, 2 H), 2.85 - 2.75 (m, 1 H), 2.71 - 2.57 (m, 3 H), 2.55 - 2.54 (m, 1 H), 2.17 (dd, *J* = 12.0, 14.0 Hz, 1 H), 1.86 (spt, *J* = 6.7 Hz, 2 H), 1.76 (d, *J* = 13.7 Hz, 1 H), 1.50 (s, 3 H), 1.55 - 1.45 (m, 1 H), 1.44 - 1.34 (m, 1 H), 1.15 (d, *J* = 6.4 Hz, 3 H), 0.94 (d, *J* = 6.7 Hz, 3 H); $^{13}$C-NMR, HSQC/HMBC (100 MHz, DMSO-$d_6$): $\delta$ = 175.2, 170.6, 170.3, 169.6, 167.7, 159.4, 156.4, 147.0, 143.6, 138.4, 137.1, 136.3, 133.3, 131.8, 129.2, 129.0, 127.3, 127.0, 123.9, 123.9, 123.4, 121.1, 120.8, 118.9, 118.4, 115.5, 115.3, 111.5, 111.3, 109.7, 107.1, 71.0, 67.2, 64.3, 55.3, 53.1, 49.0, 48.5, 47.3, 43.0, 41.7, 40.2, 35.2, 30.8, 29.3, 25.4, 23.8, 19.7, 19.5, 17.0; UV-VIS (PBS/MeCN, 2:1): $\lambda_{max}$ ($\varepsilon$) = 443 nm (22.1 × $10^3$ l·mol$^{-1}$·cm$^{-1}$); HRMS (ESI): *m/Z* calcd for $C_{58}H_{64}ClN_8O_9$ $[M+H]^+$: 1051.448, found: 1051.447.

*cyclo*-[(2*S*,4*E*,8*S*)-Hdn-L-Dap(2-(4´-((4´´-(3´´´,4´´´-dichlorbenzyl)-3´´,4´´-dihydro-2´´*H*-benzo[*b*][1´´,4´´]oxazin-7´´-yl)diazenyl)phenoxy)acetyl)-D-*N*MeTrp-L-*β*Tyr]; (**9**)

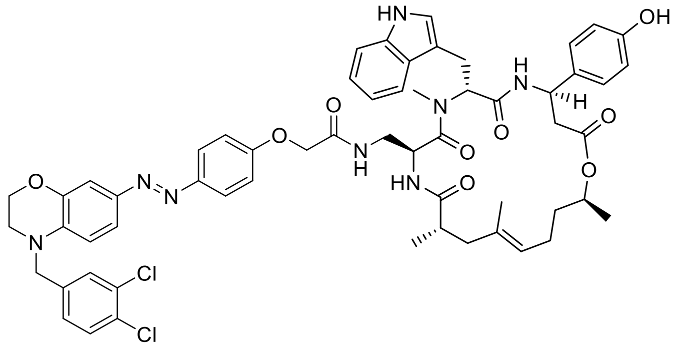

Standard Procedure **SP4** with cyclodepsipeptide **S54** (2.5 mg, 2.8 µmol, 1.0 equiv.), azobenzene **S57** (1.6 mg, 3.4 µmol, 1.2 equiv.), HATU (2.6 mg, 6.8 µmol, 2.4 equiv.) and DIPEA (3.8 µl, 22.5 µmol, 8.0 equiv.) gave conjugate **9** (1.8 mg, 59%) as an orange solid after purification by preparative HPLC ($H_2O$/MeCN, 60:40 → 0:100). $R_f$ = 0.39 ($CH_2Cl_2$/MeOH, 14:1); $[\alpha]_D^{24}$ = 61.6 (c = 0.05 in MeCN); $^1$H-NMR (400 MHz, DMSO-$d_6$): $\delta$ = 10.78 (d, $J$ = 1.8 Hz, 1 H), 9.27 (s, 1 H), 8.54 (d, $J$ = 8.8 Hz, 1 H), 7.91 (t, $J$ = 6.0 Hz, 1 H), 7.72 (d, $J$ = 9.1 Hz, 2 H), 7.67 (d, $J$ = 7.6 Hz, 1 H), 7.62 (d, $J$ = 8.2 Hz, 1 H), 7.60 - 7.55 (m, 2 H), 7.36 (dd, $J$ = 2.2, 8.6 Hz, 1 H), 7.30 (dd, $J$ = 1.9, 8.3 Hz, 1 H), 7.30 (d, $J$ = 8.2 Hz, 1 H), 7.23 (d, $J$ = 2.3 Hz, 1 H), 7.08 - 6.99 (m, 6 H), 6.95 (t, $J$ = 7.3 Hz, 1 H), 6.76 (d, $J$ = 8.8 Hz, 1 H), 6.67 (d, $J$ = 8.5 Hz, 2 H), 5.49 (dd, $J$ = 7.3, 8.8 Hz, 1 H), 5.17 (dt, $J$ = 3.5, 9.6 Hz, 1 H), 4.99 - 4.88 (m, 2 H), 4.69 (sxt, $J$ = 6.4 Hz, 1 H), 4.65 (s, 2 H), 4.43 (s, 2 H), 4.28 (t, $J$ = 4.1 Hz, 2 H), 3.56 (t, $J$ = 4.2 Hz, 2 H), 3.11 (s, 3 H), 3.07 - 2.96 (m, 2 H), 2.80 (td, $J$ = 4.9, 13.3 Hz, 1 H), 2.70 - 2.52 (m, 4 H), 2.16 (dd, $J$ = 11.8, 14.5 Hz, 1 H), 1.85 (spt, $J$ = 7.0 Hz, 2 H), 1.76 (d, $J$ = 14.3 Hz, 1 H), 1.49 (s, 3 H), 1.57 - 1.44 (m, 1 H), 1.43 - 1.33 (m, 1 H), 1.14 (d, $J$ = 6.4 Hz, 3 H), 0.93 (d, $J$ = 6.7 Hz, 3 H); $^{13}$C-NMR, HSQC/HMBC (100 MHz, DMSO-$d_6$): $\delta$ = 175.2, 170.3, 169.5, 167.9, 159.4, 156.5, 147.2,,143.8, 139.4, 138.1, 136.4, 133.3, 133.0, 131.8, 131.0, 130.1, 129.4, 129.2, 127.6, 127.2, 124.0, 123.7, 123.2, 121.1, 120.8, 118.9, 118.2, 115.5, 115.2, 111.5, 111.2, 109.9, 107.3, 71.1, 67.3, 64.3, 55.1, 53.0, 48.9, 48.5, 47.3, 43.1, 41.6, 38.1, 35.4, 30.8, 29.1, 25.5, 23.9, 19.8, 19.4, 17.0; UV-VIS (PBS/MeCN, 2:1): $\lambda_{max}$ ($\varepsilon$) = 438 nm (22.2 × $10^3$ l·mol$^{-1}$·cm$^{-1}$); HRMS (ESI): *m/Z* calcd for $C_{58}H_{63}Cl_2N_8O_9$ $[M+H]^+$: 1085.409, found: 1085.410.

*cyclo*-[(2*S*,4*E*,8*S*)-Hdn-L-Dap(2-(4´-((4´´-(4´´´-methylbenzyl)-3´´,4´´-dihydro-2´´*H*-benzo[*b*][1´´,4´´]oxazin-7´´-yl)diazenyl)phenoxy)acetyl)-D-*N*MeTrp-L-*β*Tyr]; (**10**)

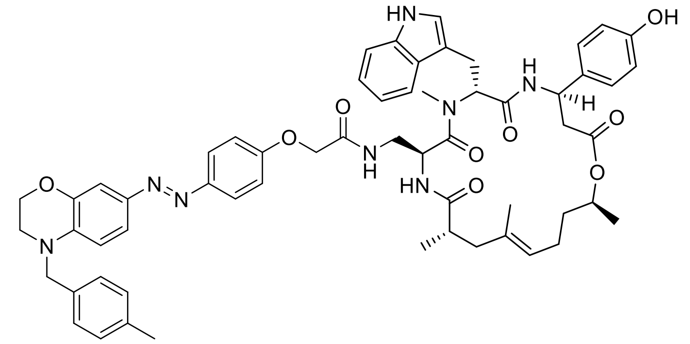

Standard Procedure **SP4** with cyclodepsipeptide **S54** (2.5 mg, 2.8 µmol, 1.0 equiv.), azobenzene **S58** (1.4 mg, 3.4 µmol, 1.2 equiv.), HATU (2.6 mg, 6.8 µmol, 2.4 equiv.) and DIPEA (3.8 µl, 22.5 µmol, 8.0 equiv.) gave conjugate **10** (1.7 mg, 59%) as a red solid after purification by preparative HPLC ($H_2O$/MeCN, 60:40 → 0:100). $R_f$ = 0.39 ($CH_2Cl_2$/MeOH, 14:1); $[\alpha]_D^{24}$ = 75.8 (c = 0.10 in MeCN); $^1$H-NMR (400 MHz, DMSO-$d_6$): $\delta$ = 10.78 (d, $J$ = 1.5 Hz, 1 H), 9.27 (s, 1 H), 8.54 (d, $J$ = 8.8 Hz, 1 H), 7.91 (t, $J$ = 6.0 Hz, 1 H), 7.71 (d, $J$ = 9.1 Hz, 2 H), 7.67 (d, $J$ = 7.6 Hz, 1 H), 7.57 (d, $J$ = 8.8 Hz, 1 H), 7.35 (dd, $J$ = 2.2, 8.6 Hz, 1 H), 7.30 (d, $J$ = 7.9 Hz, 1 H), 7.23 - 7.14 (m, 5 H), 7.08 - 6.99 (m, 6 H), 6.95 (t, $J$ = 7.6 Hz, 1 H), 6.81 (d, $J$ = 9.1 Hz, 1 H), 6.67 (d, $J$ = 8.5 Hz, 2 H), 5.49 (dd, $J$ = 7.0, 8.5 Hz, 1 H), 5.17 (dt, $J$ = 3.8, 9.6 Hz, 1 H), 4.98 - 4.89 (m, 2 H), 4.69 (sxt, $J$ = 6.4 Hz, 1 H), 4.60 (s, 2 H), 4.43 (s, 2 H), 4.25 (t, $J$ = 4.2 Hz, 2 H), 3.52 (t, $J$ = 4.4 Hz, 2 H), 3.11 (s, 3 H), 3.07 - 2.97 (m, 2 H), 2.84 - 2.75 (m, 1 H), 2.71 - 2.57 (m, 3 H), 2.56 - 2.53 (m, 1 H), 2.28 (s, 3 H), 2.16 (dd, $J$ = 11.7, 13.7 Hz, 1 H), 1.85 (spt, $J$ = 7.0 Hz, 2 H), 1.76 (d, $J$ = 14.3 Hz, 1 H), 1.49 (s, 3 H), 1.55 - 1.44 (m, 1 H), 1.43 - 1.32 (m, 1 H), 1.14 (d, $J$ = 6.4 Hz, 3 H), 0.93 (d, $J$ = 6.7 Hz, 3 H); $^{13}$C-NMR, HSQC/HMBC (100 MHz, DMSO-$d_6$): $\delta$ = 175.2, 170.4, 169.8, 167.8, 159.3, 156.4, 147.2, 143.6, 138.8, 136.5, 134.7, 132.9, 129.6, 129.3, 127.6, 127.3, 123.8, 123.7, 123.5, 121.2, 121.0, 119.0, 118.4, 115.5, 115.4, 111.4, 111.3, 109.8, 107.1, 71.1, 67.3, 64.2, 55.1, 53.4, 49.2, 48.2, 47.1, 43.0, 41.9, 38.0, 35.2, 30.9, 29.1, 25.5, 23.8, 20.8, 19.9, 19.6, 16.9; UV-VIS (PBS/MeCN, 2:1): $\lambda_{max}$ ($\varepsilon$) = 448 nm (21.6 × $10^3$ l·mol$^{-1}$·cm$^{-1}$); HRMS (ESI): $m/Z$ calcd for $C_{59}H_{67}N_8O_9$ $[M+H]^+$: 1031.503, found: 1031.503.

*cyclo*-[(2*S*,4*E*,8*S*)-Hdn-L-Dap(2-(4′-((4′′-(4′′′-(trifluormethyl)benzyl)-3′′,4′′-dihydro-2′′*H*-benzo[*b*][1′′,4′′]oxazin-7′′-yl)diazenyl)phenoxy)acetyl)-D-*N*MeTrp-L-*β*Tyr]; (**11**)

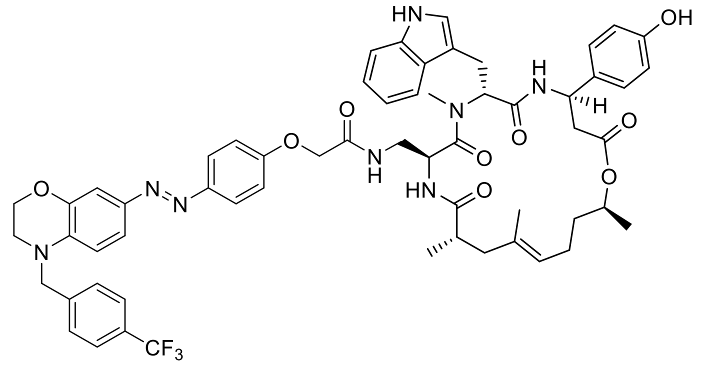

Standard Procedure **SP4** with cyclodepsipeptide **S54** (2.5 mg, 2.8 µmol, 1.0 equiv.), azobenzene **S59** (1.6 mg, 3.4 µmol, 1.2 equiv.), HATU (2.6 mg, 6.8 µmol, 2.4 equiv.) and DIPEA (3.8 µl, 22.5 µmol, 8.0 equiv.) gave conjugate **11** (1.4 mg, 46%) as a red solid after purification by preparative HPLC ($H_2O$/MeCN, 60:40 → 0:100). $R_f$ = 0.39 ($CH_2Cl_2$/MeOH, 14:1); $[\alpha]_D^{24}$ = 70.2 (c = 0.10 in MeCN); $^1$H-NMR (400 MHz, DMSO-$d_6$): $\delta$ = 10.78 (d, *J* = 1.8 Hz, 1 H), 9.27 (s, 1 H), 8.54 (d, *J* = 8.8 Hz, 1 H), 7.91 (t, *J* = 6.0 Hz, 1 H), 7.76 - 7.69 (m, 4 H), 7.67 (d, *J* = 7.6 Hz, 1 H), 7.57 (d, *J* = 8.8 Hz, 1 H), 7.53 (d, *J* = 8.2 Hz, 2 H), 7.35 (dd, *J* = 2.2, 8.6 Hz, 1 H), 7.29 (d, *J* = 7.9 Hz, 1 H), 7.24 (d, *J* = 2.3 Hz, 1 H), 7.08 - 6.99 (m, 6 H), 6.95 (t, *J* = 7.6 Hz, 1 H), 6.75 (d, *J* = 8.8 Hz, 1 H), 6.66 (d, *J* = 8.8 Hz, 2 H), 5.49 (dd, *J* = 7.0, 8.5 Hz, 1 H), 5.17 (dt, *J* = 3.5, 9.6 Hz, 1 H), 4.99 - 4.87 (m, 2 H), 4.76 (s, 2 H), 4.68 (sxt, *J* = 6.3 Hz, 1 H), 4.43 (s, 2 H), 4.30 (t, *J* = 4.2 Hz, 2 H), 3.58 (t, *J* = 4.4 Hz, 2 H), 3.11 (s, 3 H), 3.06 - 2.97 (m, 2 H), 2.85 - 2.74 (m, 1 H), 2.71 - 2.53 (m, 4 H), 2.16 (dd, *J* = 12.0, 14.0 Hz, 1 H), 1.85 (spt, *J* = 7.0 Hz, 2 H), 1.76 (d, *J* = 14.6 Hz, 1 H), 1.49 (s, 3 H), 1.55 - 1.44 (m, 1 H), 1.43 - 1.32 (m, 1 H), 1.14 (d, *J* = 6.4 Hz, 3 H), 0.93 (d, *J* = 6.7 Hz, 3 H); $^{13}$C-NMR, HSQC/HMBC (100 MHz, DMSO-$d_6$): $\delta$ = 175.3, 170.6, 170.3, 169.7, 167.8, 159.5, 156.5, 147.1, 143.7, 143.0, 138.3, 136.3, 135.2, 133.0, 129.1, 128.0, 127.4, 126.7, 125.8, 124.0, 123.8, 123.2, 121.2, 120.9, 119.1, 118.4, 115.4, 115.2, 111.4, 111.2, 109.9, 107.3, 71.0, 67.3, 64.3, 55.2, 53.5, 49.2, 48.4, 47.5, 43.0, 41.7, 38.1, 35.3, 30.8, 29.1, 25.5, 23.8, 19.8, 19.4, 16.8; UV-VIS (PBS/MeCN, 2:1): $\lambda_{max}$ ($\varepsilon$) = 437 nm (21.3 × $10^3$ l·mol$^{-1}$·cm$^{-1}$); HRMS (ESI): *m/Z* calcd for $C_{59}H_{64}F_3N_8O_9$ $[M+H]^+$: 1085.474, found: 1085.474.

*cyclo*-[(2*S*,4*E*,8*S*)-Hdn-L-Dap(2-(4´-((4´´-(4´´´-(*tert*-butyl)benzyl)-3´´,4´´-dihydro-2´´*H*-benzo[*b*][1´´,4´´]oxazin-7´´-yl)diazenyl)phenoxy)acetyl)-D-*N*MeTrp-L-*β*Tyr]; (**12**)

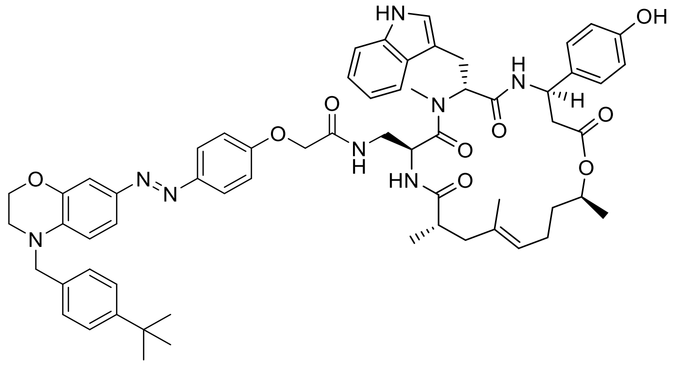

Standard Procedure **SP4** with cyclodepsipeptide **S54** (2.5 mg, 2.8 µmol, 1.0 equiv.), azobenzene **S60** (1.6 mg, 3.4 µmol, 1.2 equiv.), HATU (2.6 mg, 6.8 µmol, 2.4 equiv.) and DIPEA (3.8 µl, 22.5 µmol, 8.0 equiv.) gave conjugate **12** (1.8 mg, 60%) as a red solid after purification by preparative HPLC ($H_2O$/MeCN, 60:40 → 0:100). $R_f$ = 0.39 ($CH_2Cl_2$/MeOH, 14:1); $[\alpha]_D^{24}$ = 43.9 (c = 0.05 in MeCN); $^1$H-NMR (400 MHz, DMSO-$d_6$): $\delta$ = 10.78 (d, $J$ = 1.8 Hz, 1 H), 9.27 (s, 1 H), 8.54 (d, $J$ = 8.8 Hz, 1 H), 7.91 (t, $J$ = 5.8 Hz, 1 H), 7.72 (d, $J$ = 8.8 Hz, 2 H), 7.67 (d, $J$ = 7.9 Hz, 1 H), 7.57 (d, $J$ = 8.8 Hz, 1 H), 7.37 (d, $J$ = 8.2 Hz, 2 H), 7.36 (dd, $J$ = 2.6, 8.5 Hz, 1 H), 7.30 (d, $J$ = 7.9 Hz, 1 H), 7.26 - 7.20 (m, 3 H), 7.08 - 6.99 (m, 6 H), 6.95 (t, $J$ = 7.4 Hz, 1 H), 6.82 (d, $J$ = 8.8 Hz, 1 H), 6.67 (d, $J$ = 8.5 Hz, 2 H), 5.49 (dd, $J$ = 7.3, 8.5 Hz, 1 H), 5.17 (dt, $J$ = 3.5, 9.7 Hz, 1 H), 4.99 - 4.88 (m, 2 H), 4.69 (sxt, $J$ = 6.4 Hz, 1 H), 4.61 (s, 2 H), 4.43 (s, 2 H), 4.26 (t, $J$ = 4.1 Hz, 2 H), 3.53 (t, $J$ = 4.2 Hz, 2 H), 3.11 (s, 3 H), 3.07 - 2.98 (m, 2 H), 2.84 - 2.75 (m, 1 H), 2.70 - 2.54 (m, 4 H), 2.16 (dd, $J$ = 11.4, 13.7 Hz, 1 H), 1.85 (spt, $J$ = 7.0 Hz, 2 H), 1.76 (d, $J$ = 14.3 Hz, 1 H), 1.49 (s, 3 H), 1.55 - 1.45 (m, 1 H), 1.43 - 1.32 (m, 1 H), 1.26 (s, 9 H), 1.14 (d, $J$ = 6.1 Hz, 3 H), 0.93 (d, $J$ = 6.7 Hz, 3 H); $^{13}$C-NMR, HSQC/HMBC (100 MHz, DMSO-$d_6$): $\delta$ = 175.2, 170.8, 170.3, 169.5, 167.8, 159.2, 156.6, 149.6, 147.1, 143.5, 139.0, 138.7, 136.3, 134.7, 133.1, 128.7, 127.4, 126.9, 125.7, 124.0, 123.8, 123.3, 121.2, 121.0, 119.0, 118.3, 115.4, 115.2, 111.3, 111.2, 109.9, 107.1, 71.1, 67.1, 63.9, 55.1, 53.4, 49.3, 48.3, 47.2, 43.3, 41.5, 38.4, 35.3, 34.5, 31.2, 30.8, 29.1, 25.4, 23.7, 19.9, 19.4, 17.1; UV-VIS (PBS/MeCN, 2:1): $\lambda_{max}$ ($\varepsilon$) = 449 nm (22.1 × $10^3$ l·mol$^{-1}$·cm$^{-1}$); HRMS (ESI): *m/Z* calcd for $C_{62}H_{73}N_8O_9$ $[M+H]^+$: 1073.550, found: 1073.551.

*cyclo*-[(2*S*,4*E*,8*S*)-Hdn-L-Dap(2-(4′-((4′′-phenyl-3′′,4′′-dihydro-2′′*H*-benzo[*b*][1′′,4′′]-oxazin-7′′-yl)diazenyl)phenoxy)acetyl)-D-*N*MeTrp-L-*β*Tyr]; (**13**)

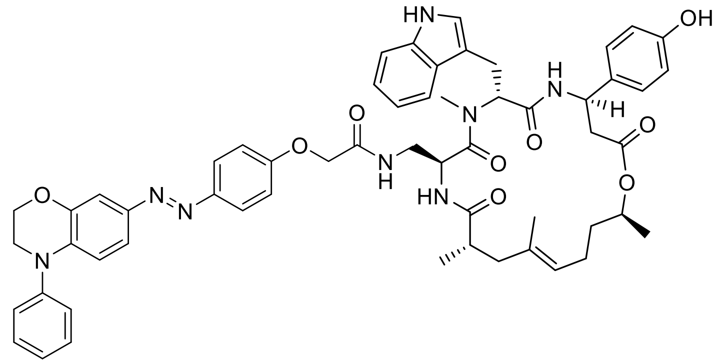

Standard Procedure **SP4** with cyclodepsipeptide **S54** (10.0 mg, 11.3 µmol, 1.0 equiv.), azobenzene **S61** (4.4 mg, 11.3 µmol, 1.0 equiv.), HATU (10.3 mg, 27.0 µmol, 2.4 equiv.) and DIPEA (15.3 µl, 90.1 µmol, 8.0 equiv.) gave conjugate **13** (3.7 mg, 33%) as an orange solid after purification by silica-gel chromatography ($CH_2Cl_2$/acetone, 2:1 → 1:2). $R_f$ = 0.39 ($CH_2Cl_2$/MeOH, 14:1); $[\alpha]_D^{24}$ = 60.3 (c = 0.05 in MeCN); $^1$H-NMR (500 MHz, DMSO-*d*$_6$): $\delta$ = 10.79 (s, 1 H), 9.29 (s, 1 H), 8.57 (d, *J* = 8.5 Hz, 1 H), 7.95 (t, *J* = 5.8 Hz, 1 H), 7.76 (d, *J* = 8.9 Hz, 2 H), 7.67 (d, *J* = 7.9 Hz, 1 H), 7.61 (d, *J* = 8.5 Hz, 1 H), 7.50 - 7.43 (m, 2 H), 7.37 (d, *J* = 7.9 Hz, 2 H), 7.35 - 7.27 (m, 3 H), 7.23 (t, *J* = 7.3 Hz, 1 H), 7.11 - 6.99 (m, 6 H), 6.95 (t, *J* = 7.3 Hz, 1 H), 6.84 (d, *J* = 7.9 Hz, 1 H), 6.66 (d, *J* = 8.5 Hz, 2 H), 5.50 (dd, *J* = 6.7, 9.2 Hz, 1 H), 5.17 (ddd, *J* = 3.0, 8.4, 11.5 Hz, 1 H), 5.01 - 4.86 (m, 2 H), 4.68 (sxt, *J* = 6.3 Hz, 1 H), 4.49 - 4.39 (m, 2 H), 4.35 (t, *J* = 4.1 Hz, 2 H), 3.79 (t, *J* = 4.3 Hz, 2 H), 3.11 (s, 3 H), 3.16 - 2.95 (m, 2 H), 2.85 - 2.72 (m, 1 H), 2.72 - 2.53 (m, 4 H), 2.16 (dd, *J* = 11.9, 14.3 Hz, 1 H), 1.85 (spt, *J* = 7.6 Hz, 2 H), 1.76 (d, *J* = 14.0 Hz, 1 H), 1.49 (s, 3 H), 1.56 - 1.43 (m, 1 H), 1.43 - 1.33 (m, 1 H), 1.14 (d, *J* = 6.4 Hz, 3 H), 0.93 (d, *J* = 6.7 Hz, 3 H); $^{13}$C-NMR (126 MHz, DMSO-*d*$_6$): $\delta$ = 175.4, 170.7, 170.5, 170.0, 168.0, 159.8, 156.7, 147.2, 145.5, 145.3, 144.5, 136.6, 136.5, 133.5, 133.3, 130.2, 127.6, 127.4, 125.4, 124.9, 124.2, 123.8, 123.7, 121.3, 119.2, 119.1, 118.5, 115.7, 115.5, 114.4, 111.6, 110.0, 109.1, 71.2, 67.4, 64.7, 55.3, 49.3, 48.5, 48.4, 43.2, 41.9, 38.3, 35.4, 31.0, 25.8, 24.1, 20.0, 19.8, 17.2; UV-VIS (PBS/MeCN, 2:1): $\lambda_{max}$ ($\varepsilon$) = 438 nm ($28.6 \times 10^3$ l·mol$^{-1}$·cm$^{-1}$); HRMS (ESI): *m/Z* calcd for $C_{57}H_{63}N_8O_9$ $[M+H]^+$: 1003.471, found: 1003.472.

*cyclo*-[(2*S*,4*E*,8*S*)-Hdn-L-Dap(2-(4′-((4′′-phenethyl-3′′,4′′-dihydro-2′′*H*-benzo[*b*]-[1′′,4′′]oxazin-7′′-yl)diazenyl)phenoxy)acetyl)-D-*N*MeTrp-L-*β*Tyr], (**14**)

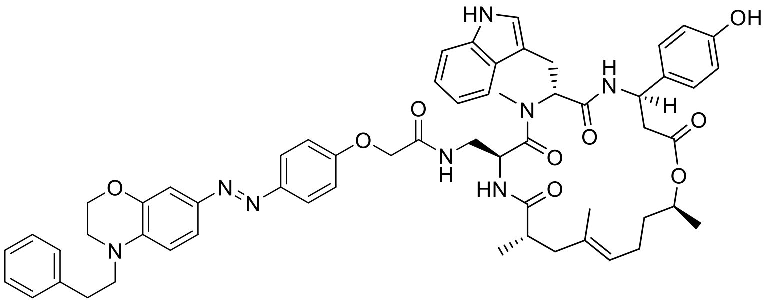

Standard Procedure **SP4** with cyclodepsipeptide **S54** (2.5 mg, 2.8 µmol, 1.0 equiv.), azobenzene **S62** (1.4 mg, 3.4 µmol, 1.2 equiv.), HATU (2.6 mg, 6.8 µmol, 2.4 equiv.) and DIPEA (3.8 µl, 22.5 µmol, 8.0 equiv.) gave conjugate **14** (1.9 mg, 64%) as a red solid after purification by preparative HPLC ($H_2O$/MeCN, 60:40 → 0:100). $R_f$ = 0.39 ($CH_2Cl_2$/MeOH, 14:1); $[\alpha]_D^{24}$ = 47.8 (c = 0.10 in MeCN); $^1$H-NMR (600 MHz, DMSO-$d_6$): $\delta$ = 10.80 (d, *J* = 1.8 Hz, 1 H), 9.29 (s, 1 H), 8.57 (d, *J* = 8.6 Hz, 1 H), 7.94 (t, *J* = 6.0 Hz, 1 H), 7.73 (d, *J* = 9.0 Hz, 2 H), 7.68 (d, *J* = 7.9 Hz, 1 H), 7.61 (d, *J* = 8.6 Hz, 1 H), 7.44 (dd, *J* = 2.4, 8.6 Hz, 1 H), 7.35 - 7.29 (m, 5 H), 7.23 (sxt, *J* = 4.2 Hz, 1 H), 7.20 (d, *J* = 2.4 Hz, 1 H), 7.08 - 7.01 (m, 6 H), 6.96 (ddd, *J* = 0.7, 7.0, 7.9 Hz, 1 H), 6.91 (d, *J* = 8.8 Hz, 1 H), 6.67 (d, *J* = 8.6 Hz, 2 H), 5.50 (dd, *J* = 6.5, 9.6 Hz, 1 H), 5.18 (ddd, *J* = 3.1, 8.8, 11.6 Hz, 1 H), 4.98 - 4.91 (m, 2 H), 4.69 (sxt, *J* = 6.3 Hz, 1 H), 4.44 (d, *J* = 3.9 Hz, 2 H), 4.13 (t, *J* = 4.4 Hz, 2 H), 3.64 (t, *J* = 7.6 Hz, 2 H), 3.38 (t, *J* = 4.2 Hz, 2 H), 3.12 (s, 3 H), 3.08 - 2.96 (m, 2 H), 2.89 (t, *J* = 7.5 Hz, 2 H), 2.82 - 2.76 (m, 1 H), 2.70 - 2.57 (m, 3 H), 2.55 - 2.53 (m, 1 H), 2.17 (dd, *J* = 11.6, 14.3 Hz, 1 H), 1.91 - 1.80 (m, 2 H), 1.77 (d, *J* = 14.1 Hz, 1 H), 1.50 (s, 3 H), 1.54 - 1.47 (m, 1 H), 1.42 - 1.35 (m, 1 H), 1.15 (d, *J* = 6.4 Hz, 3 H), 0.94 (d, *J* = 6.8 Hz, 3 H); $^{13}$C-NMR, HSQC/HMBC (150 MHz, DMSO-$d_6$): $\delta$ = 175.3, 170.6, 170.3, 169.8, 167.9, 159.2, 156.5, 147.2, 143.9, 142.9, 139.5, 138.3, 136.3, 133.3, 133.1, 129.8, 129.1, 127.4, 127.2, 126.6, 124.0, 123.7, 121.4, 121.0, 119.0, 118.3, 115.5, 115.2, 111.4, 110.7, 109.8, 107.2, 71.2, 67.2, 63.9, 54.9, 52.0, 48.9, 48.1, 46.7, 43.0, 41.9, 35.2, 32.1, 30.9, 29.2, 25.6, 23.7, 20.0, 19.5, 17.0; UV-VIS (PBS/MeCN, 2:1): $\lambda_{max}$ ($\varepsilon$) = 454 nm (18.5 × $10^3$ l·mol$^{-1}$·cm$^{-1}$); HRMS (ESI): *m/Z* calcd for $C_{59}H_{67}N_8O_9$ $[M+H]^+$: 1031.503, found: 1031.503.

*cyclo*-[(2*S*,4*E*,8*S*)-Hdn-L-Dab(2-(4´-((4´´-benzyl-3´´,4´´-dihydro-2´´*H*-benzo[*b*][1´´,4´´]-oxazin-7´´-yl)diazenyl)phenoxy)acetyl)-D-*N*MeTrp-L-*β*Tyr]; (**15**)

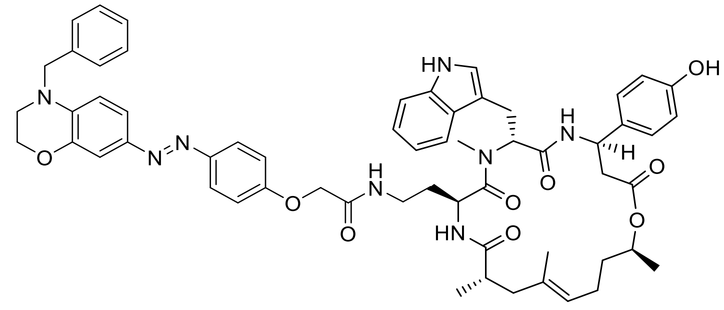

Standard Procedure **SP4** with cyclodepsipeptide **S53** (10.0 mg, 11.3 µmol, 1.0 equiv.), 2-(4´-((4´´-benzyl-3´´,4´´-dihydro-2´´*H*-benzo[b][1´´,4´´]oxazin-7´´-yl)diazenyl)phenoxy) acetic acid (4.5 mg, 11.3 µmol, 1.0 equiv.), HATU (10.3 mg, 27.0 µmol, 2.4 equiv.) and DIPEA (15.3 µl, 90.1 µmol, 8.0 equiv.) gave conjugate **15** (5.8 mg, 50%) as an orange solid after purification by preparative HPLC ($H_2O$/MeCN, 60:40 → 0:100). $R_f$ = 0.39 ($CH_2Cl_2$/MeOH, 14:1); $^1$H-NMR (400 MHz, DMSO-$d_6$, isomeric mixture 3:2) main isomer: $\delta$ = 10.74 (d, $J$ = 1.8 Hz, 1 H), 9.28 (s, 1 H), 8.51 (d, $J$ = 8.8 Hz, 1 H), 7.87 (t, $J$ = 5.8 Hz, 1 H), 7.84 - 7.72 (m, 3 H), 7.64 (d, $J$ = 7.9 Hz, 1 H), 7.40 - 7.27 (m, 7 H), 7.25 (d, $J$ = 2.3 Hz, 1 H), 7.14 - 6.91 (m, 7 H), 6.81 (d, $J$ = 8.8 Hz, 1 H), 6.67 (d, $J$ = 8.5 Hz, 2 H), 5.48 (dd, $J$ = 7.0, 8.8 Hz, 1 H), 5.16 (t, $J$ = 9.6 Hz, 1 H), 4.96 (q, $J$ = 7.3 Hz, 1 H), 4.76 - 4.67 (m, 2 H), 4.66 (s, 2 H), 4.53 (s, 2 H), 4.28 (t, $J$ = 4.2 Hz, 2 H), 3.55 (t, $J$ = 4.2 Hz, 2 H), 3.00 (s, 3 H), 3.08 - 2.88 (m, 3 H), 2.85 - 2.72 (m, 1 H), 2.71 - 2.54 (m, 4 H), 2.24 - 2.12 (m, 1 H), 1.92 - 1.82 (m, 2 H), 1.79 (d, $J$ = 15.2 Hz, 1 H), 1.50 (s, 3 H), 1.54 - 1.45 (m, 1 H), 1.43 - 1.21 (m, 2 H), 1.13 (d, $J$ = 6.4 Hz, 3 H), 0.97 (d, $J$ = 6.7 Hz, 3 H); $^{13}$C-NMR (101 MHz, DMSO-$d_6$, isomeric mixture 3:2) main isomer: $\delta$ = 174.9, 170.2, 170.2, 169.5, 167.0, 159.1, 156.2, 146.9, 143.4, 143.2, 138.4, 137.5, 136.1, 133.1, 132.7, 128.6, 127.0, 127.0, 126.9, 123.5, 123.4, 123.2, 120.8, 120.7, 118.5, 118.1, 115.2, 115.0, 115.0, 111.2, 111.0, 109.6, 106.9, 70.6, 67.1, 63.8, 54.9, 53.5, 48.8, 47.1, 46.1, 42.8, 41.3, 40.9, 37.8, 34.9, 31.3, 30.4, 25.2, 23.5, 19.5, 19.4, 16.7; UV-VIS (PBS/MeCN, 2:1): $\lambda_{max}$ ($\varepsilon$) = 448 nm ($21.3 \times 10^3$ l·mol$^{-1}$·cm$^{-1}$); HRMS (ESI): *m/Z* calcd for $C_{59}H_{67}N_8O_9$ $[M+H]^+$: 1031.502, found: 1031.503.

*cyclo*-[(2*S*,4*E*,8*S*)-Adn-L-Dap(2-(4´-((4´´-benzyl-3´´,4´´-dihydro-2´´*H*-benzo[*b*]-[1´´,4´´]-oxazin-7´´-yl)diazenyl)phenoxy)acetyl)-D-*N*MeTrp-L-*β*Tyr]; (**16**)

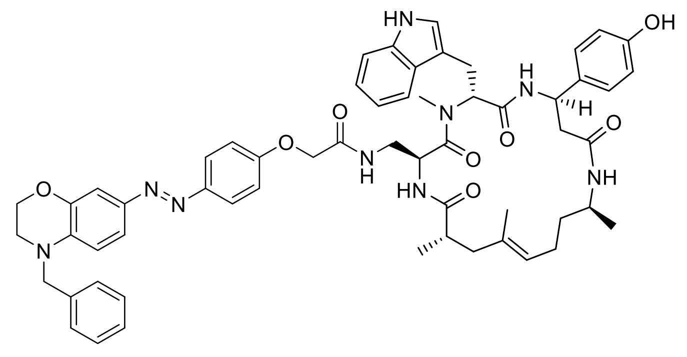

Standard Procedure **SP4** with cyclopeptide **S63** (8.0 mg, 9.0 µmol, 1.0 equiv.), 2-(4´-((4´´-benzyl-3´´,4´´-dihydro-2´´*H*-benzo[b][1´´,4´´]oxazin-7´´-yl)diazenyl)phenoxy) acetic acid (3.6 mg, 9.0 µmol, 1.0 equiv.), HATU (8.2 mg, 21.6 µmol, 2.4 equiv.) and DIPEA (12.3 µl, 72.1 µmol, 8.0 equiv.) gave conjugate **16** (7.7 mg, 84%) as an orange solid after purification by silica-gel chromatography ($CH_2Cl_2$/acetone, 4:1 → 1:1 + 1% MeOH). $R_f$ = 0.20 ($CH_2Cl_2$/MeOH, 19:1); $[\alpha]_D^{24}$ = 31.6 (c = 0.10 in MeCN); $^1$H-NMR (500 MHz, DMSO-$d_6$): $\delta$ = 10.81 (d, *J* = 1.8 Hz, 1 H), 9.25 (s, 1 H), 8.49 (d, *J* = 8.2 Hz, 1 H), 7.91 (t, *J* = 6.0 Hz, 1 H), 7.72 (d, *J* = 9.2 Hz, 2 H), 7.62 (d, *J* = 7.9 Hz, 1 H), 7.53 (d, *J* = 8.5 Hz, 1 H), 7.43 (d, *J* = 7.9 Hz, 1 H), 7.39 - 7.34 (m, 3 H), 7.33 - 7.30 (m, 3 H), 7.30 - 7.26 (m, 1 H), 7.23 (d, *J* = 2.1 Hz, 1 H), 7.05 (ddd, *J* = 1.5, 6.7, 7.9 Hz, 1 H), 7.03 (d, *J* = 9.2 Hz, 2 H), 7.05 - 6.98 (m, 3 H), 6.96 (ddd, *J* = 0.9, 7.0, 7.9 Hz, 1 H), 6.80 (d, *J* = 8.9 Hz, 1 H), 6.67 (d, *J* = 8.5 Hz, 2 H), 5.50 (dd, *J* = 7.0, 9.2 Hz, 1 H), 5.08 (ddd, *J* = 3.0, 7.6, 10.4 Hz, 1 H), 4.98 (dt, *J* = 3.8, 8.9 Hz, 1 H), 4.94 (t, *J* = 6.6 Hz, 1 H), 4.66 (s, 2 H), 4.42 (s, 2 H), 4.28 (t, *J* = 4.3 Hz, 2 H), 3.65 (qud, *J* = 6.9, 14.0 Hz, 1 H), 3.55 (t, *J* = 4.4 Hz, 2 H), 3.11 (s, 3 H), 3.09 - 2.99 (m, 2 H), 2.83 (td, *J* = 4.7, 13.3 Hz, 1 H), 2.66 - 2.53 (m, 2 H), 2.46 (dd, *J* = 10.2, 14.2 Hz, 1 H), 2.30 (dd, *J* = 3.1, 14.0 Hz, 1 H), 2.17 (dd, *J* = 11.7, 14.2 Hz, 1 H), 1.81 - 1.71 (m, 3 H), 1.48 (s, 3 H), 1.36 - 1.26 (m, 1 H), 1.25 - 1.12 (m, 1 H), 1.01 (d, *J* = 6.7 Hz, 3 H), 0.94 (d, *J* = 7.0 Hz, 3 H); $^{13}$C-NMR (126 MHz, DMSO-$d_6$): $\delta$ = 175.4, 170.6, 169.6, 169.1, 168.0, 159.4, 156.5, 147.3, 143.9, 143.6, 138.9, 138.0, 136.6, 134.0, 133.1, 129.1, 127.5, 127.4, 127.4, 124.4, 124.0, 123.6, 121.4, 121.3, 119.0, 118.6, 115.7, 115.4, 111.7, 111.5, 110.1, 107.3, 67.4, 64.3, 55.6, 54.0, 49.7, 48.7, 47.6, 44.3, 43.3, 43.0, 38.4, 36.4, 31.0, 25.5, 24.5, 20.6, 20.0, 17.1; IR: $\tilde{\nu}_{max}$ = 3726, 3391, 2935, 1647, 1498, 1438, 1389, 1253, 1103, 1061, 748, 660; UV-VIS (PBS/MeCN, 2:1): $\lambda_{max}$ ($\varepsilon$) = 447 nm (24.4 × $10^3$ l·mol$^{-1}$·cm$^{-1}$); HRMS (ESI): *m/Z* calcd for $C_{58}H_{66}N_9O_8$ $[M+H]^+$: 1016.503, found: 1016.503.

*cyclo*-[(2*S*,4*E*)-Hdo-L-Dap(2-(4´-((3´´,4´´,5´´-trimethoxyphenyl)diazenyl)phenoxy)-acetyl)-D-*N*MeTrp-L-*β*Tyr]; (**S1**)

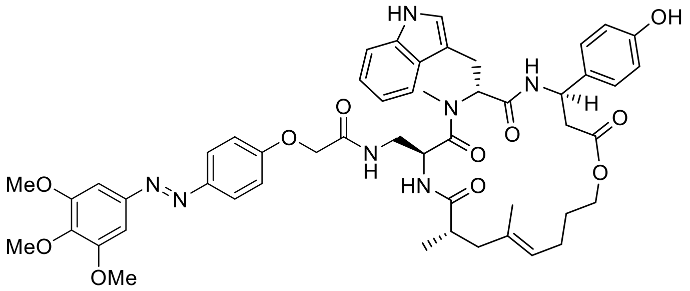

Standard Procedure **SP4** with cyclodepsipeptide **S64** (10.0 mg, 11.4 µmol, 1.0 equiv.), 2-(4´-((3´´,4´´,5´´-trimethoxyphenyl)diazenyl)phenoxy)acetic acid (4.0 mg, 11.4 µmol, 1.0 equiv.), HATU (10.4 mg, 27.5 µmol, 2.4 equiv.) and DIPEA (15.6 µl, 91.5 µmol, 8.0 equiv.) gave conjugate **S1** (5.4 mg, 50%) as a yellow solid after purification by silica-gel chromatography ($CH_2Cl_2$/acetone, 4:1 → 1:1 + 1% MeOH). $R_f$ = 0.38 ($CH_2Cl_2$/MeOH, 14:1); $[\alpha]_D^{24}$ = 24.5 (c = 0.09 in MeCN); $^1$H-NMR (500 MHz, DMSO-$d_6$): $\delta$ = 10.83 (s, 1 H), 9.49 (br s, 1 H), 8.61 (d, $J$ = 8.5 Hz, 1 H), 8.03 (t, $J$ = 5.5 Hz, 1 H), 7.86 (d, $J$ = 9.2 Hz, 2 H), 7.68 (d, $J$ = 8.2 Hz, 1 H), 7.67 (d, $J$ = 9.2 Hz, 1 H), 7.30 (d, $J$ = 8.2 Hz, 1 H), 7.22 (s, 2 H), 7.09 (d, $J$ = 8.8 Hz, 2 H), 7.07 (d, $J$ = 9.2 Hz, 2 H), 7.05 - 7.02 (m, 2 H), 6.95 (t, $J$ = 7.5 Hz, 1 H), 6.69 (d, $J$ = 8.5 Hz, 2 H), 5.51 (dd, $J$ = 7.0, 9.5 Hz, 1 H), 5.22 (ddd, $J$ = 3.4, 8.2, 11.9 Hz, 1 H), 4.99 - 4.91 (m, 2 H), 4.46 (d, $J$ = 4.6 Hz, 2 H), 4.11 - 4.02 (m, 1 H), 3.89 (s, 6 H), 3.88 - 3.81 (m, 1 H), 3.76 (s, 3 H), 3.09 (s, 3 H), 3.08 - 3.02 (m, 2 H), 2.76 - 2.61 (m, 3 H), 2.15 (dd, $J$ = 11.9, 14.3 Hz, 1 H), 1.86 (q, $J$ = 7.5 Hz, 2 H), 1.78 (d, $J$ = 14.0 Hz, 1 H), 1.61 (s, 2 H), 1.52 (s, 3 H), 1.50 - 1.39 (m, 2 H), 0.94 (d, $J$ = 6.7 Hz, 3 H); $^{13}$C-NMR (126 MHz, DMSO-$d_6$): $\delta$ = 175.4, 170.8, 170.6, 170.1, 167.9, 160.6, 156.8, 153.8, 148.3, 146.8, 140.3, 136.6, 133.6, 133.2, 127.5, 127.4, 124.8, 123.7, 123.6, 121.3, 119.2, 118.5, 115.8, 115.5, 111.6, 110.1, 100.4, 67.3, 64.3, 60.7, 56.5, 55.3, 49.2, 48.4, 43.3, 42.1, 40.7, 38.3, 31.2, 28.6, 25.9, 24.4, 19.7, 17.1; IR: $\tilde{\nu}_{max}$ = 3314, 2936, 1732, 1667, 1651, 1597, 1501, 1458, 1331, 1231, 1126, 1003, 845, 745, 652; UV-VIS (PBS/MeCN, 2:1): $\lambda_{max}$ ($\varepsilon$) = 360 nm (20.5 × $10^3$ l·mol$^{-1}$·cm$^{-1}$); HRMS (ESI): *m/Z* calcd for $C_{51}H_{60}N_7O_{11}$ [M+H]$^+$: 946.4345, found: 946.4344.

*cyclo*-[(2*S*,4*E*)-Hdo-L-Dap(4-((5´-methoxy-2´-((3´´,4´´,5´´-trimethoxyphenyl)diazenyl)-phenyl)amino)-4-oxobutyryl)-D-*N*MeTrp-L-$\beta$Tyr]; (**S2**)

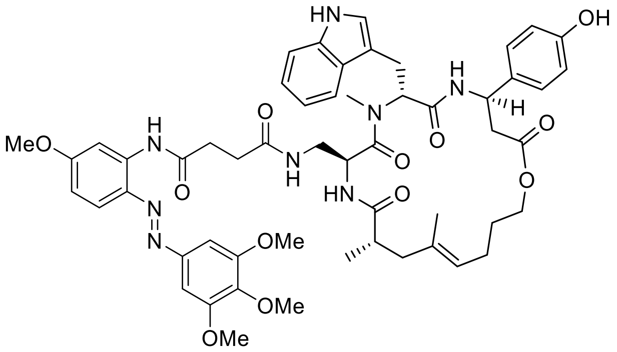

Standard Procedure **SP4** with cyclodepsipeptide **S64** (10.0 mg, 11.4 µmol, 1.0 equiv.), *N*-(5´-Methoxy-2´-((3´´,4´´,5´´-trimethoxyphenyl)diazenyl)phenyl)succinamic acid (5.7 mg, 11.4 µmol, 1.0 equiv.), HATU (10.4 mg, 27.5 µmol, 2.4 equiv.) and DIPEA (15.6 µl, 91.5 µmol, 8.0 equiv.) gave conjugate **S2** (8.0 mg, 69%) as an orange solid after purification by silica-gel chromatography ($CH_2Cl_2$/acetone, 4:1 → 1:1 + 1% MeOH). $R_f$ = 0.29 ($CH_2Cl_2$/MeOH, 14:1); $[\alpha]_D^{24}$ = 25.6 (c = 0.10 in MeCN); $^1$H-NMR (500 MHz, DMSO-$d_6$): $\delta$ = 10.79 (br s, 1 H), 10.27 (s, 1 H), 9.48 (br s, 1 H), 8.57 (d, *J* = 8.5 Hz, 1 H), 7.99 (d, *J* = 2.7 Hz, 1 H), 7.75 (d, *J* = 8.8 Hz, 1 H), 7.67 (d, *J* = 7.9 Hz, 1 H), 7.72 - 7.63 (m, 1 H), 7.59 (d, *J* = 8.5 Hz, 1 H), 7.36 (s, 2 H), 7.29 (d, *J* = 7.9 Hz, 1 H), 7.08 (d, *J* = 8.2 Hz, 2 H), 7.02 (t, *J* = 7.6 Hz, 1 H), 6.99 (d, *J* = 1.2 Hz, 1 H), 6.94 (t, *J* = 7.6 Hz, 1 H), 6.79 (dd, *J* = 2.6, 9.0 Hz, 1 H), 6.69 (d, *J* = 8.5 Hz, 2 H), 5.49 (dd, *J* = 6.0, 10.5 Hz, 1 H), 5.21 (ddd, *J* = 3.7, 8.2, 11.9 Hz, 1 H), 4.93 (t, *J* = 7.2 Hz, 1 H), 4.88 (dt, *J* = 4.0, 9.0 Hz, 1 H), 4.09 - 4.01 (m, 1 H), 3.89 (s, 6 H), 3.84 (s, 3 H), 3.87 - 3.80 (m, 1 H), 3.76 (s, 3 H), 3.09 - 3.02 (m, 1 H), 3.03 (s, 3 H), 2.99 (dd, *J* = 10.7, 15.0 Hz, 1 H), 2.73 - 2.61 (m, 5 H), 2.35 - 2.26 (m, 2 H), 2.10 (dd, *J* = 11.9, 14.0 Hz, 1 H), 1.83 (q, *J* = 7.4 Hz, 2 H), 1.73 (d, *J* = 14.0 Hz, 1 H), 1.67 (s, 2 H), 1.49 (s, 3 H), 1.47 - 1.38 (m, 2 H), 0.92 (d, *J* = 6.7 Hz, 3 H); $^{13}$C-NMR (126 MHz, DMSO-$d_6$): $\delta$ = 175.2, 172.0, 171.7, 170.8, 170.6, 170.1, 163.0, 156.8, 153.7, 148.6, 140.3, 139.2, 136.5, 134.9, 133.6, 133.2, 127.5, 127.4, 123.7, 123.6, 121.3, 119.2, 118.8, 118.5, 115.5, 111.6, 110.5, 110.1, 106.2, 101.1, 64.3, 60.7, 56.5, 56.0, 55.2, 49.1, 48.5, 43.3, 42.1, 40.7, 38.2, 32.4, 31.0, 30.6, 28.6, 26.0, 24.3, 19.7, 17.0; IR: $\tilde{\nu}_{max}$ = 3364, 2936, 1734, 1662, 1608, 1520, 1458, 1427, 1288, 1234, 1126, 1003, 840, 745, 652; UV-VIS (PBS/MeCN, 2:1): $\lambda_{max}$ ($\varepsilon$) = 385 nm (20.7 × $10^3$ l·mol$^{-1}$·cm$^{-1}$); HRMS (ESI): *m/Z* calcd for $C_{54}H_{65}N_8O_{12}$ $[M+H]^+$: 1017.472, found: 1017.472.

*cyclo*-[(2*S*,4*E*)-Hdo-L-Lys(4-((5´-methoxy-2´-((3´´,4´´,5´´-trimethoxyphenyl)diazenyl)-phenyl)amino)-4-oxobutyryl)-D-*N*MeTrp-L-*β*Tyr]; (**S3**)

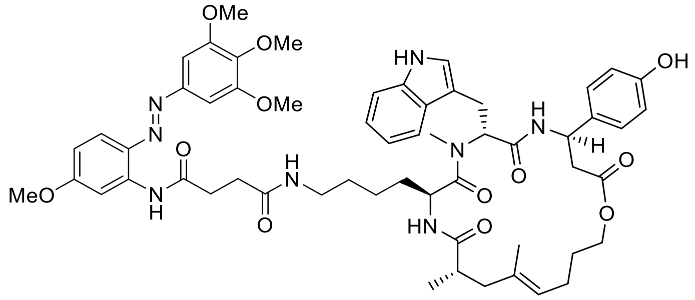

Standard Procedure **SP4** with cyclodepsipeptide **S65** (10.0 mg, 10.9 µmol, 1.0 equiv.), *N*-(5´-Methoxy-2´-((3´´,4´´,5´´-trimethoxyphenyl)diazenyl)phenyl)succinamic acid (4.6 mg, 10.9 µmol, 1.0 equiv.), HATU (10.0 mg, 26.2 µmol, 2.4 equiv.) and DIPEA (14.9 µl, 87.3 µmol, 8.0 equiv.) gave conjugate **S3** (9.4 mg, 81%) as an orange solid after purification by silica-gel chromatography ($CH_2Cl_2$/acetone, 4:1 → 1:1 + 1% MeOH). $R_f$ = 0.30 ($CH_2Cl_2$/MeOH, 14:1); $[\alpha]_D^{24}$ = 21.2 (c = 0.11 in MeCN); $^1$H-NMR (500 MHz, DMSO-$d_6$): $\delta$ = 10.77 (d, $J$ = 1.5 Hz, 1 H), 10.28 (s, 1 H), 9.32 (br s, 1 H), 8.64 (d, $J$ = 8.9 Hz, 1 H), 8.04 (d, $J$ = 2.7 Hz, 1 H), 7.83 (t, $J$ = 5.6 Hz, 1 H), 7.74 (d, $J$ = 8.8 Hz, 1 H), 7.67 (d, $J$ = 8.2 Hz, 1 H), 7.64 (d, $J$ = 8.5 Hz, 1 H), 7.39 (s, 2 H), 7.27 (d, $J$ = 8.2 Hz, 1 H), 7.15 (d, $J$ = 8.5 Hz, 2 H), 7.05 (d, $J$ = 2.1 Hz, 1 H), 6.99 (ddd, $J$ = 0.9, 7.1, 8.0 Hz, 1 H), 6.93 (ddd, $J$ = 0.9, 7.0, 8.0 Hz, 1 H), 6.78 (dd, $J$ = 2.7, 9.2 Hz, 1 H), 6.71 (d, $J$ = 8.9 Hz, 2 H), 5.51 (dd, $J$ = 6.4, 10.1 Hz, 1 H), 5.23 (ddd, $J$ = 3.6, 8.7, 11.0 Hz, 1 H), 4.92 (t, $J$ = 6.7 Hz, 1 H), 4.55 (dt, $J$ = 3.8, 8.5 Hz, 1 H), 4.12 - 4.04 (m, 1 H), 3.90 (s, 6 H), 3.83 (s, 3 H), 3.87 - 3.80 (m, 1 H), 3.75 (s, 3 H), 2.98 (s, 3 H), 3.05 - 2.94 (m, 2 H), 2.88 - 2.76 (m, 2 H), 2.74 (t, $J$ = 7.0 Hz, 2 H), 2.71 - 2.62 (m, 2 H), 2.54 - 2.52 (m, 1 H), 2.46 (t, $J$ = 7.0 Hz, 2 H), 2.13 (dd, $J$ = 12.1, 14.5 Hz, 1 H), 1.83 (spt, $J$ = 7.6 Hz, 2 H), 1.74 (d, $J$ = 14.3 Hz, 1 H), 1.50 (s, 3 H), 1.49 - 1.39 (m, 2 H), 1.05 (quq, $J$ = 6.4, 13.4 Hz, 2 H), 0.91 (d, $J$ = 6.7 Hz, 3 H), 0.82 - 0.67 (m, 3 H), 0.64 - 0.55 (m, 1 H); $^{13}$C-NMR (126 MHz, DMSO-$d_6$): $\delta$ = 174.8, 172.8, 171.8, 171.6, 170.7, 170.6, 163.0, 156.7, 153.7, 148.6, 140.3, 139.3, 136.6, 134.8, 133.5, 133.4, 127.5, 127.4, 123.7, 123.5, 121.3, 119.1, 118.6, 118.5, 115.6, 111.6, 110.4, 110.1, 106.0, 101.1, 64.5, 60.7, 56.5, 56.0, 55.1, 49.1, 47.9, 43.2, 42.3, 38.8, 38.0, 32.9, 31.2, 30.9, 29.0, 28.7, 28.5, 26.1, 24.4, 22.4, 19.9, 17.2; IR: $\tilde{\nu}_{max}$ = 3311, 2940, 1732, 1631, 1601, 1520, 1458, 1334, 1288, 1234, 1126, 1003, 841, 745, 652; UV-VIS (PBS/MeCN, 2:1): $\lambda_{max}$ ($\varepsilon$) = 387 nm (20.3 × $10^3$ l·mol$^{-1}$·cm$^{-1}$); HRMS (ESI): *m/Z* calcd for $C_{57}H_{71}N_8O_{12}$ $[M+H]^+$: 1059.519, found: 1059.518.

*cyclo*-[(2*S*,4*E*,6*R*,8*S*)-Htn-L-Dap(4-((5´-methoxy-2´-((3´´,4´´,5´´-trimethoxyphenyl)-diazenyl)phenyl)amino)-4-oxobutyryl)-D-*N*MeTrp-L-*β*Tyr]; (**S6**)

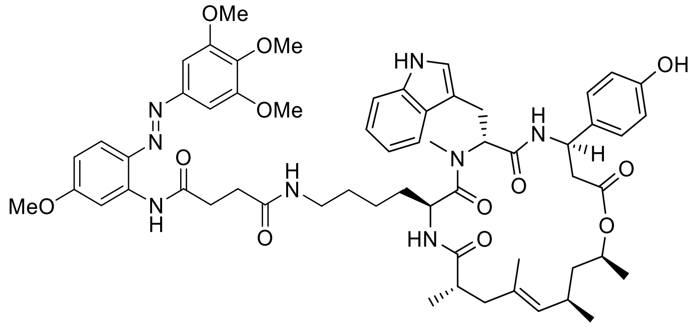

Standard Procedure **SP4** with cyclodepsipeptide **S66** (11.0 mg, 11.7 µmol, 1.0 equiv.), *N*-(5´-Methoxy-2´-((3´´,4´´,5´´-trimethoxyphenyl)diazenyl)phenyl)succinamic acid (4.9 mg, 11.7 µmol, 1.0 equiv.), HATU (10.6 mg, 28.0 µmol, 2.4 equiv.) and DIPEA (15.9 µl, 93.2 µmol, 8.0 equiv.) gave conjugate **S6** (5.0 mg, 39%) as an orange solid after purification by silica-gel chromatography ($CH_2Cl_2$/acetone, 4:1 → 1:1 + 1% MeOH). $R_f$ = 0.38 ($CH_2Cl_2$/MeOH, 14:1); $[\alpha]_D^{24}$ = 29.8 (c = 0.11 in MeCN); $^1$H-NMR (400 MHz, DMSO-$d_6$): $\delta$ = 10.81 (d, *J* = 1.5 Hz, 1 H), 10.29 (s, 1 H), 9.42 (br s, 1 H), 8.25 (d, *J* = 8.8 Hz, 1 H), 8.04 (d, *J* = 2.6 Hz, 1 H), 7.87 (t, *J* = 5.4 Hz, 1 H), 7.75 (d, *J* = 9.1 Hz, 1 H), 7.63 (d, *J* = 8.2 Hz, 1 H), 7.60 (d, *J* = 7.9 Hz, 1 H), 7.39 (s, 2 H), 7.28 (d, *J* = 7.9 Hz, 1 H), 7.07 (d, *J* = 8.5 Hz, 2 H), 7.04 (d, *J* = 2.0 Hz, 1 H), 7.00 (ddd, *J* = 0.9, 6.7, 8.2 Hz, 1 H), 6.93 (ddd, *J* = 0.9, 6.8, 8.2 Hz, 1 H), 6.78 (dd, *J* = 2.8, 9.2 Hz, 1 H), 6.68 (d, *J* = 8.5 Hz, 2 H), 5.44 (t, *J* = 8.0 Hz, 1 H), 5.16 (dt, *J* = 4.7, 8.8 Hz, 1 H), 4.79 (d, *J* = 9.1 Hz, 1 H), 4.62 (sxt, *J* = 6.7 Hz, 1 H), 4.60 - 4.51 (m, 1 H), 3.91 (s, 6 H), 3.83 (s, 3 H), 3.75 (s, 3 H), 2.97 (s, 3 H), 3.08 - 2.93 (m, 2 H), 2.74 (t, *J* = 6.9 Hz, 2 H), 2.92 - 2.70 (m, 3 H), 2.62 - 2.56 (m, 1 H), 2.47 (t, *J* = 7.0 Hz, 2 H), 2.36 - 2.23 (m, 2 H), 2.10 (dd, *J* = 10.4, 15.3 Hz, 1 H), 1.76 (dd, *J* = 2.3, 15.4 Hz, 1 H), 1.50 (s, 3 H), 1.58 - 1.44 (m, 1 H), 1.33 - 1.21 (m, 2 H), 1.11 (d, *J* = 6.1 Hz, 2 H), 1.04 (d, *J* = 6.4 Hz, 3 H), 0.94 (d, *J* = 6.7 Hz, 3 H), 0.82 (d, *J* = 6.4 Hz, 3 H), 0.98 - 0.79 (m, 2 H), 0.77 - 0.70 (m, 1 H); $^{13}$C-NMR (101 MHz, DMSO-$d_6$): $\delta$ = 174.9, 172.4, 171.8, 171.6, 170.5, 169.7, 163.0, 156.8, 153.7, 148.6, 140.3, 139.3, 136.6, 134.8, 132.5, 132.1, 129.7, 127.7, 127.4, 123.7, 121.3, 118.9, 118.6, 118.6, 115.4, 111.7, 110.4, 110.0, 106.0, 101.1, 70.2, 60.7, 56.5, 56.1, 55.5, 49.1, 48.5, 47.3, 42.8, 42.2, 41.7, 38.9, 38.6, 32.9, 31.6, 30.9, 30.6, 29.3, 29.2, 22.3, 22.1, 20.1, 19.9, 18.2; IR: $\tilde{\nu}_{max}$ = 3310, 2932, 1732, 1667, 1597, 1520, 1458, 1288, 1231, 1126, 1003, 837, 745, 652; UV-VIS (PBS/MeCN, 2:1): $\lambda_{max}$ ($\varepsilon$) = 388 nm (21.0 × $10^3$ l·mol$^{-1}$·cm$^{-1}$); HRMS (ESI): *m/Z* calcd for $C_{59}H_{75}N_8O_{12}$ $[M+H]^+$: 1087.550, found: 1087.550.

*cyclo*-[(2*S*,4*Z*,6*R*,8*S*)-Htn-L-Dap(4-((5′-methoxy-2′-((3′′,4′′,5′′-trimethoxyphenyl)-diazenyl)phenyl)amino)-4-oxobutyryl)-D-*N*MeTrp-L-*β*Tyr]; (**S7**)

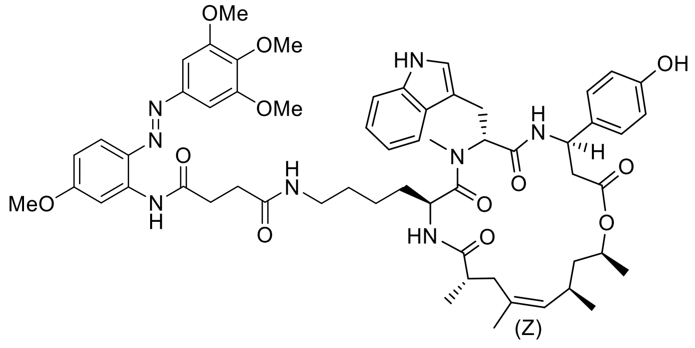

Standard Procedure **SP4** with cyclodepsipeptide **S67** (10.0 mg, 10.6 µmol, 1.0 equiv.), *N*-(5′-Methoxy-2′-((3′′,4′′,5′′-trimethoxyphenyl)diazenyl)phenyl)succinamic acid (4.4 mg, 10.6 µmol, 1.0 equiv.), HATU (9.7 mg, 25.4 µmol, 2.4 equiv.) and DIPEA (14.4 µl, 84.7 µmol, 8.0 equiv.) gave conjugate **S7** (3.7 mg, 32%) as an orange solid after purification by silica-gel chromatography ($CH_2Cl_2$/acetone, 4:1 → 1:1 + 1% MeOH). $R_f$ = 0.38 ($CH_2Cl_2$/MeOH, 14:1); $[\alpha]_D^{24}$ = 9.9 (c = 0.11 in MeCN); $^1$H-NMR (500 MHz, DMSO-$d_6$): $\delta$ = 10.78 (s, 1 H), 10.28 (s, 1 H), 9.36 (br s, 1 H), 8.35 (d, *J* = 8.5 Hz, 1 H), 8.04 (d, *J* = 2.7 Hz, 1 H), 7.88 (qu, *J* = 5.5 Hz, 1 H), 7.81 (d, *J* = 7.9 Hz, 1 H), 7.75 (d, *J* = 9.2 Hz, 1 H), 7.58 (d, *J* = 7.9 Hz, 1 H), 7.39 (s, 2 H), 7.27 (d, *J* = 8.2 Hz, 1 H), 7.14 - 7.06 (m, 3 H), 7.00 (ddd, *J* = 0.9, 7.0, 7.9 Hz, 1 H), 6.92 (ddd, *J* = 0.9, 7.0, 7.9 Hz, 1 H), 6.79 (dd, *J* = 2.7, 9.2 Hz, 1 H), 6.68 (d, *J* = 8.5 Hz, 2 H), 5.42 (dd, *J* = 6.3, 9.6 Hz, 1 H), 5.16 (dt, *J* = 5.8, 9.5 Hz, 1 H), 5.02 (d, *J* = 9.2 Hz, 1 H), 4.41 - 4.33 (m, 2 H), 3.91 (s, 6 H), 3.84 (s, 3 H), 3.75 (s, 3 H), 2.99 (s, 3 H), 3.06 - 2.94 (m, 1 H), 2.92 - 2.80 (m, 3 H), 2.75 (t, *J* = 7.0 Hz, 2 H), 2.78 - 2.70 (m, 1 H), 2.62 - 2.54 (m, 3 H), 2.48 (t, *J* = 7.0 Hz, 2 H), 2.29 (dd, *J* = 8.2, 13.4 Hz, 1 H), 1.85 - 1.72 (m, 2 H), 1.63 (s, 3 H), 1.68 - 1.61 (m, 1 H), 1.52 - 1.45 (m, 1 H), 1.33 - 1.19 (m, 2 H), 1.16 - 1.03 (m, 2 H), 0.99 (d, *J* = 6.4 Hz, 3 H), 0.96 - 0.91 (m, 1 H), 0.88 (d, *J* = 6.7 Hz, 3 H), 0.90 - 0.83 (m, 1 H), 0.82 (d, *J* = 6.7 Hz, 3 H), 0.79 - 0.73 (m, 1 H); $^{13}$C-NMR (126 MHz, DMSO-$d_6$): $\delta$ = 175.3, 172.5, 171.8, 171.6, 170.4, 169.8, 163.0, 156.8, 153.7, 148.6, 140.3, 139.3, 136.6, 134.8, 133.4, 132.8, 131.2, 127.8, 127.5, 123.7, 121.2, 118.9, 118.6, 118.5, 115.4, 111.6, 110.4, 110.1, 106.0, 101.1, 71.7, 60.7, 56.5, 56.1, 55.3, 49.0, 48.9, 44.4, 38.7, 37.3, 35.2, 32.9, 30.9, 30.6, 29.4, 29.1, 26.8, 26.7, 26.3, 26.2, 23.0, 22.4, 20.0, 18.2; IR: $\tilde{\nu}_{max}$ = 3317, 2932, 1728, 1651, 1597, 1519, 1458, 1331, 1288, 1234, 1126, 1006, 841, 745, 648; UV-VIS (PBS/MeCN, 2:1): $\lambda_{max}$ ($\varepsilon$) = 384 nm (21.9 × $10^3$ l·mol$^{-1}$·cm$^{-1}$); HRMS (ESI): *m/Z* calcd for $C_{59}H_{75}N_8O_{12}$ [M+H]$^+$: 1087.550, found: 1087.550.

*cyclo*-[(2*S*,4*E*,8*S*)-Adn-L-Dap(2-(4´-((3´´,4´´,5´´-trimethoxyphenyl)diazenyl)phenoxy)-acetyl)-D-*N*MeTrp-L-*β*Tyr]; (**S8**)

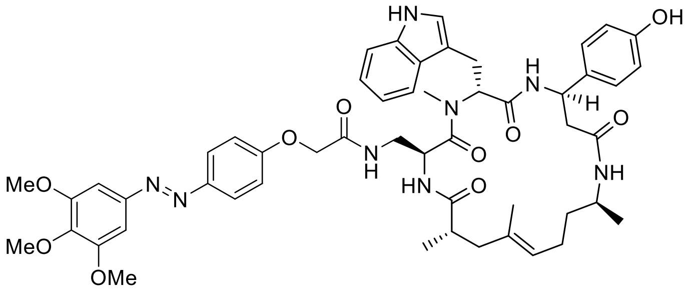

Standard Procedure **SP4** with cyclopeptide **S63** (8.0 mg, 9.0 µmol, 1.0 equiv2-(4´-((3´´,4´´,5´´-trimethoxyphenyl)diazenyl)phenoxy)acetic acid (3.1 mg, 9.0 µmol, 1.0 equiv.), HATU (8.2 mg, 21.6 µmol, 2.4 equiv.) and DIPEA (12.3 µl, 72.1 µmol, 8.0 equiv.) gave conjugate **S8** (7.5 mg, 87%) as a yellow solid after purification by silica-gel chromatography ($CH_2Cl_2$/acetone, 4:1 → 1:1 + 1% MeOH). $R_f$ = 0.20 ($CH_2Cl_2$/MeOH, 19:1); $[\alpha]_D^{24}$ = 30.4 (c = 0.10 in MeCN); $^1$H-NMR (500 MHz, DMSO-$d_6$): $\delta$ = 10.81 (d, $J$ = 1.8 Hz, 1 H), 9.25 (s, 1 H), 8.50 (d, $J$ = 8.2 Hz, 1 H), 7.96 (t, $J$ = 6.0 Hz, 1 H), 7.86 (d, $J$ = 9.2 Hz, 2 H), 7.63 (d, $J$ = 7.9 Hz, 1 H), 7.57 (d, $J$ = 8.2 Hz, 1 H), 7.44 (d, $J$ = 7.9 Hz, 1 H), 7.31 (d, $J$ = 8.2 Hz, 1 H), 7.22 (s, 2 H), 7.09 (d, $J$ = 9.2 Hz, 2 H), 7.05 (ddd, $J$ = 1.2, 7.3, 8.2 Hz, 1 H), 7.02 - 6.98 (m, 3 H), 6.96 (ddd, $J$ = 1.1, 7.0, 8.0 Hz, 1 H), 6.67 (d, $J$ = 8.5 Hz, 2 H), 5.50 (dd, $J$ = 6.7, 9.2 Hz, 1 H), 5.08 (ddd, $J$ = 3.3, 7.3, 10.7 Hz, 1 H), 4.99 (dt, $J$ = 3.8, 8.9 Hz, 1 H), 4.94 (t, $J$ = 6.7 Hz, 1 H), 4.47 (d, $J$ = 1.5 Hz, 2 H), 3.89 (s, 6 H), 3.76 (s, 3 H), 3.65 (spt, $J$ = 7.0 Hz, 1 H), 3.11 (s, 3 H), 3.10 - 3.01 (m, 2 H), 2.84 (td, $J$ = 4.8, 13.0 Hz, 1 H), 2.62 - 2.54 (m, 2 H), 2.46 (dd, $J$ = 10.1, 14.0 Hz, 1 H), 2.30 (dd, $J$ = 3.2, 13.9 Hz, 1 H), 2.17 (dd, $J$ = 11.9, 14.3 Hz, 1 H), 1.81 - 1.72 (m, 3 H), 1.49 (s, 3 H), 1.37 - 1.26 (m, 1 H), 1.23 - 1.15 (m, 1 H), 1.01 (d, $J$ = 6.7 Hz, 3 H), 0.95 (d, $J$ = 6.7 Hz, 3 H); $^{13}$C-NMR (126 MHz, DMSO-$d_6$): $\delta$ = 175.4, 170.6, 169.6, 169.1, 167.9, 160.6, 156.5, 153.8, 148.3, 146.9, 140.3, 136.6, 134.0, 133.1, 127.4, 127.4, 124.8, 124.4, 123.6, 121.4, 119.0, 118.6, 115.8, 115.4, 111.7, 110.1, 100.4, 67.4, 66.2, 60.7, 56.5, 55.6, 49.7, 48.7, 44.3, 43.3, 43.0, 38.4, 36.4, 31.0, 25.5, 24.5, 20.6, 20.0, 17.1; IR: $\tilde{\nu}_{max}$ = 3734, 3364, 3283, 2932, 1732, 1651, 1543, 1330, 1226, 1126, 1003, 845, 745, 652; UV-VIS (PBS/MeCN, 2:1): $\lambda_{max}$ ($\varepsilon$) = 360 nm (18.3 × $10^3$ l·mol$^{-1}$·cm$^{-1}$); HRMS (ESI): $m/Z$ calcd for $C_{52}H_{63}N_8O_{10}$ $[M+H]^+$: 959.466, found: 959.466

*cyclo*-[(2*S*,4*E*,8*S*)-*N*MeAdn-L-Dap(2-(4´-((3´´,4´´,5´´-trimethoxyphenyl)diazenyl)-phenoxy)acetyl)-D-*N*MeTrp-L-*β*Tyr]; (**S9**)

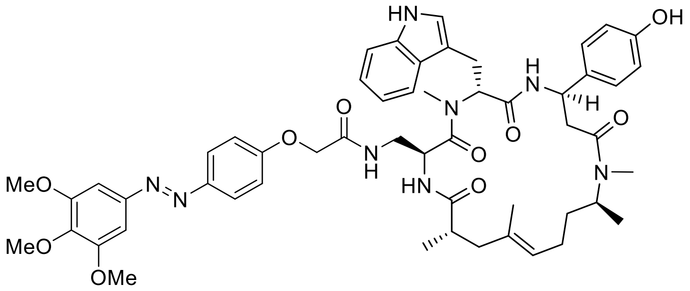

Standard Procedure **SP4** with cyclopeptide **S68** (8.0 mg, 8.9 µmol, 1.0 equiv.), 2-(4´-((3´´,4´´,5´´-trimethoxyphenyl)diazenyl)phenoxy)acetic acid (3.1 mg, 8.9 µmol, 1.0 equiv.), HATU (8.1 mg, 21.3 µmol, 2.4 equiv.) and DIPEA (12.1 µl, 71.0 µmol, 8.0 equiv.) gave conjugate **S9** (6.3 mg, 73%) as a yellow solid after purification by silica-gel chromatography ($CH_2Cl_2$/acetone, 4:1 → 1:1 + 1% MeOH). $R_f$ = 0.21 ($CH_2Cl_2$/MeOH, 19:1); $[\alpha]_D^{24}$ = 10.4 (c = 0.10 in MeCN); $^1$H-NMR (500 MHz, DMSO-$d_6$, isomeric mixture 2:1) main isomer: $\delta$ = 10.78 (s, 1 H), 9.50 (br s, 1 H), 9.25 (s, 1 H), 8.33 (d, *J* = 8.5 Hz, 1 H), 8.00 (t, *J* = 6.0 Hz, 1 H), 7.86 (d, *J* = 8.9 Hz, 2 H), 7.66 (d, *J* = 7.9 Hz, 1 H), 7.57 (d, *J* = 8.2 Hz, 1 H), 7.32 (d, *J* = 7.9 Hz, 1 H), 7.22 (s, 2 H), 7.15 - 7.07 (m, 2 H), 7.05 (ddd, *J* = 0.9, 6.7, 7.9 Hz, 1 H), 6.96 (d, *J* = 7.9 Hz, 2 H), 6.99 - 6.93 (m, 1 H), 6.65 (d, *J* = 8.5 Hz, 2 H), 5.41 (dd, *J* = 7.0, 8.9 Hz, 1 H), 5.24 (ddd, *J* = 2.4, 8.4, 11.0 Hz, 1 H), 5.02 - 4.90 (m, 2 H), 4.48 (s, 2 H), 3.89 (s, 6 H), 3.76 (s, 3 H), 3.57 - 3.53 (m, 1 H), 3.48 (t, *J* = 5.8 Hz, 1 H), 3.28 - 3.23 (m, 1 H), 3.16 - 3.08 (m, 1 H), 3.05 (s, 3 H), 3.02 - 2.86 (m, 2 H), 2.75 (s, 3 H), 2.72 - 2.58 (m, 2 H), 2.10 (dd, *J* = 11.6, 13.4 Hz, 1 H), 1.88 - 1.65 (m, 3 H), 1.49 (s, 3 H), 1.33 - 1.17 (m, 2 H), 0.95 (d, *J* = 6.7 Hz, 6 H); $^{13}$C-NMR (126 MHz, DMSO-$d_6$): $\delta$ = 175.5, 170.0, 169.1, 167.9, 165.9, 160.6, 156.4, 153.8, 148.3, 146.8, 140.3, 136.6, 133.9, 133.1, 127.5, 127.4, 124.9, 124.8, 124.8, 123.7, 121.3, 119.1, 118.6, 115.8, 115.3, 111.7, 110.2, 100.4, 67.4, 60.7, 56.5, 56.3, 53.9, 49.6, 48.4, 47.4, 41.0, 38.6, 38.1, 32.2, 28.7, 26.4, 23.8, 19.4, 19.3, 18.1, 16.8; IR: $\tilde{\nu}_{max}$ = 3734, 3136, 2943, 1732, 1624, 1508, 1454, 1419, 1377, 1038, 918, 748, 652 cm$^{-1}$; UV-VIS (PBS/MeCN, 2:1): $\lambda_{max}$ ($\varepsilon$) = 360 nm (18.7 × $10^3$ l·mol$^{-1}$·cm$^{-1}$); HRMS (ESI): *m/Z* calcd for $C_{53}H_{65}N_8O_{10}$ $[M+H]^+$: 973.4818, found: 973.4820.

*cyclo*-[(2*S*,4*E*,8*S*)-Adn-L-Dap(4-((5´-methoxy-2´-((3´´,4´´,5´´-trimethoxyphenyl)-diazenyl)phenyl)amino)-4-oxobutyryl)-D-*N*MeTrp-L-*β*Tyr]; (**S10**)

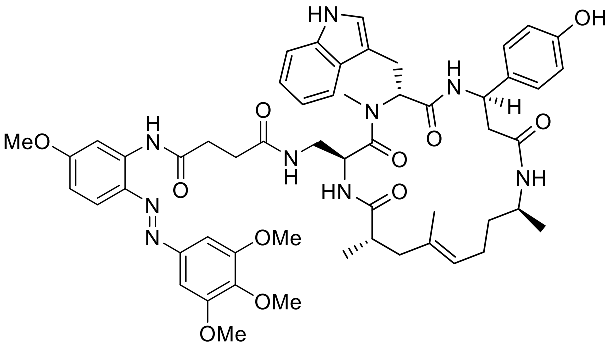

Standard Procedure **SP4** with cyclopeptide **S63** (8.0 mg, 9.0 µmol, 1.0 equiv.), *N*-(5´-Methoxy-2´-((3´´,4´´,5´´-trimethoxyphenyl)diazenyl)phenyl)succinamic acid (3.8 mg, 9.0 µmol, 1.0 equiv.), HATU (8.2 mg, 21.6 µmol, 2.4 equiv.) and DIPEA (12.3 µl, 72.1 µmol, 8.0 equiv.) gave conjugate **S10** (5.2 mg, 56%) as an orange solid after purification by silica-gel chromatography ($CH_2Cl_2$/acetone, 4:1 → 1:1 + 1% MeOH). $R_f$ = 0.10 ($CH_2Cl_2$/MeOH, 19:1); $[\alpha]_D^{24}$ = 15.6 (c = 0.10 in MeCN); $^1$H-NMR (500 MHz, DMSO-$d_6$): $\delta$ = 10.78 (d, $J$ = 2.1 Hz, 1 H), 10.25 (s, 1 H), 9.26 (s, 1 H), 8.49 (d, $J$ = 8.5 Hz, 1 H), 7.99 (d, $J$ = 2.7 Hz, 1 H), 7.75 (d, $J$ = 9.2 Hz, 1 H), 7.66 - 7.59 (m, 2 H), 7.51 (d, $J$ = 8.5 Hz, 1 H), 7.45 (d, $J$ = 7.9 Hz, 1 H), 7.36 (s, 2 H), 7.30 (d, $J$ = 7.9 Hz, 1 H), 7.02 (ddd, $J$ = 0.9, 7.0, 8.2 Hz, 1 H), 7.00 (d, $J$ = 8.5 Hz, 2 H), 6.96 (d, $J$ = 2.1 Hz, 1 H), 6.94 (ddd, $J$ = 0.9, 7.0, 8.2 Hz, 1 H), 6.79 (dd, $J$ = 2.9, 9.0 Hz, 1 H), 6.67 (d, $J$ = 8.5 Hz, 2 H), 5.49 (dd, $J$ = 6.1, 9.8 Hz, 1 H), 5.06 (ddd, $J$ = 2.9, 8.0, 10.5 Hz, 1 H), 4.95 - 4.86 (m, 2 H), 3.89 (s, 6 H), 3.84 (s, 3 H), 3.76 (s, 3 H), 3.66 - 3.58 (m, 2 H), 3.11 - 3.06 (m, 1 H), 3.05 (s, 3 H), 3.04 - 2.95 (m, 2 H), 2.78 - 2.65 (m, 3 H), 2.56 - 2.53 (m, 1 H), 2.35 - 2.29 (m, 2 H), 2.29 - 2.24 (m, 1 H), 2.13 (dd, $J$ = 11.9, 14.0 Hz, 1 H), 1.79 - 1.73 (m, 2 H), 1.71 (d, $J$ = 15.0 Hz, 1 H), 1.46 (s, 3 H), 1.32 - 1.18 (m, 2 H), 1.01 (d, $J$ = 6.4 Hz, 3 H), 0.93 (d, $J$ = 7.0 Hz, 3 H); $^{13}$C-NMR (126 MHz, DMSO-$d_6$): $\delta$ = 174.7, 171.5, 171.2, 170.2, 169.3, 168.6, 162.5, 156.0, 153.3, 148.1, 139.8, 138.7, 136.1, 134.4, 133.6, 132.6, 126.9, 126.9, 123.9, 123.1, 120.9, 118.6, 118.3, 118.1, 114.9, 111.2, 110.0, 109.6, 105.7, 100.6, 60.2, 56.0, 55.6, 54.9, 52.2, 49.3, 48.3, 45.5, 43.9, 42.8, 37.9, 34.2, 31.9, 30.4, 30.1, 25.1, 24.1, 20.0, 19.5, 16.6; IR: $\tilde{\nu}_{max}$ = 3734, 3568, 3001, 2943, 1732, 1601, 1419, 1377, 1037, 918, 748, 652; UV-VIS (PBS/MeCN, 2:1): $\lambda_{max}$ ($\varepsilon$) = 385 nm (19.2 × 10$^3$ l·mol$^{-1}$·cm$^{-1}$); HRMS (ESI): $m/Z$ calcd $C_{55}H_{68}N_9O_{11}$ $[M+H]^+$: 1030.503, found: 1030.502.

*cyclo*-[(2*S*,4*E*,8*S*)-*N*MeAdn-L-Dap(4-((5´-methoxy-2´-((3´´,4´´,5´´-trimethoxyphenyl)-diazenyl)phenyl)amino)-4-oxobutyryl)-D-*N*MeTrp-L-*β*Tyr]; (**S11**)

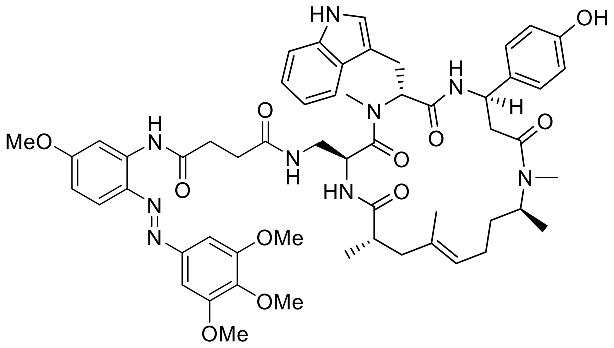

Standard Procedure **SP4** with cyclopeptide **S68** (8.0 mg, 8.9 µmol, 1.0 equiv.), *N*-(5´-Methoxy-2´-((3´´,4´´,5´´-trimethoxyphenyl)diazenyl)phenyl)succinamic acid (3.7 mg, 8.9 µmol, 1.0 equiv.), HATU (8.1 mg, 21.3 µmol, 2.4 equiv.) and DIPEA (12.1 µl, 71.0 µmol, 8.0 equiv.) gave conjugate **S11** (4.6 mg, 50%) as an orange solid after purification by silica-gel chromatography ($CH_2Cl_2$/acetone, 4:1 → 1:1 + 1% MeOH). $R_f$ = 0.18 ($CH_2Cl_2$/MeOH, 19:1); $[\alpha]_D^{24}$ = 26.2 (c = 0.10 in MeCN); $^1$H-NMR (500 MHz, DMSO-$d_6$, isomeric mixture 3:1) main isomer: δ = 10.75 (d, *J* = 1.5 Hz, 1 H), 10.26 (s, 1 H), 9.24 (s, 1 H), 8.31 (d, *J* = 8.5 Hz, 1 H), 7.99 (d, *J* = 2.7 Hz, 1 H), 7.75 (d, *J* = 9.2 Hz, 2 H), 7.64 (d, *J* = 7.3 Hz, 1 H), 7.49 (d, *J* = 8.2 Hz, 1 H), 7.36 (s, 2 H), 7.30 (d, *J* = 7.9 Hz, 1 H), 7.05 - 7.00 (m, 2 H), 6.98 - 6.92 (m, 3 H), 6.79 (dd, *J* = 2.7, 9.2 Hz, 1 H), 6.65 (d, *J* = 8.5 Hz, 2 H), 5.40 (dd, *J* = 7.0, 8.9 Hz, 1 H), 5.23 (ddd, *J* = 2.2, 8.2, 10.4 Hz, 1 H), 4.94 (t, *J* = 7.2 Hz, 1 H), 4.88 (dt, *J* = 4.7, 8.3 Hz, 1 H), 4.52 (sxt, *J* = 7.2 Hz, 1 H), 3.89 (s, 6 H), 3.84 (s, 3 H), 3.76 (s, 3 H), 3.09 (dd, *J* = 6.4, 14.6 Hz, 1 H), 3.00 (s, 3 H), 2.97 - 2.89 (m, 3 H), 2.87 - 2.80 (m, 1 H), 2.75 (s, 3 H), 2.72 - 2.65 (m, 3 H), 2.61 - 2.56 (m, 1 H), 2.35 - 2.29 (m, 2 H), 2.07 (dd, *J* = 11.6, 14.0 Hz, 1 H), 1.80 - 1.66 (m, 3 H), 1.45 (s, 3 H), 1.31 - 1.17 (m, 2 H), 0.95 (d, *J* = 6.7 Hz, 3 H), 0.93 (d, *J* = 6.4 Hz, 3 H); $^{13}$C-NMR (126 MHz, DMSO-$d_6$): *δ* = 175.2, 172.0, 171.7, 170.5, 170.0, 169.2, 163.0, 156.4, 153.7, 148.6, 140.3, 139.2, 136.6, 134.9, 133.9, 133.1, 128.0, 127.4, 124.8, 123.7, 121.3, 119.1, 118.8, 118.5, 115.3, 111.6, 110.5, 110.2, 106.2, 101.0, 60.7, 56.5, 56.1, 49.6, 48.5, 47.4, 43.9, 41.2, 40.8, 38.6, 33.3, 32.4, 32.4, 30.9, 30.6, 28.7, 25.4, 24.7, 19.4, 18.0, 16.7.2, 40.8, 38.6, 33.3, 32.4, 32.4, 30.9, 30.6, 30.6, 28.7, 25.4, 24.7, 19.4, 18.0, 16.7; IR: $\tilde{\nu}$ = 3630, 3383, 2928, 1732, 1608, 1520, 1458, 1234, 1126, 1029, 1003, 845, 745, 652; UV-VIS (PBS/MeCN, 2:1): $\lambda_{max}$ ($\varepsilon$) = 385 nm (18.0 × $10^3$ l·mol$^{-1}$·cm$^{-1}$); HRMS (ESI): *m/Z* calcd for $C_{56}H_{70}N_9O_{11}$ $[M+H]^+$: 1044.519, found: 1044.518.

*cyclo*-[(2*S*,4*E*,8*S*)-*N*MeAdn-L-Dap(2-(4′-((4′′-benzyl-3′′,4′′-dihydro-2′′*H*-benzo[*b*]-[1′′,4′′]-oxazin-7′′-yl)diazenyl)phenoxy)acetyl)-D-*N*MeTrp-L-*β*Tyr]; (**S12**)

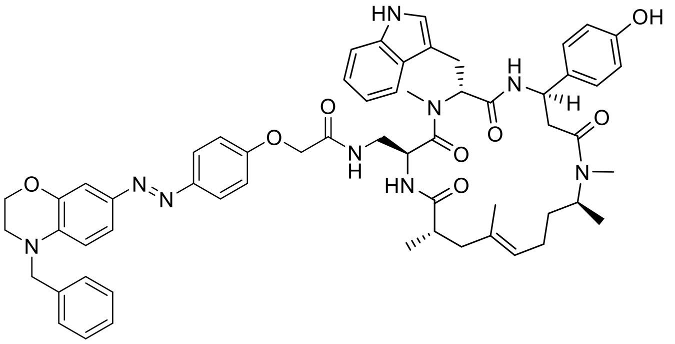

Standard Procedure **SP4** with cyclopeptide **S68** (8.0 mg, 8.9 µmol, 1.0 equiv.), 2-(4′-((4′′-benzyl-3′′,4′′-dihydro-2′′*H*-benzo[b][1′′,4′′]oxazin-7′′-yl)diazenyl)phenoxy) acetic acid (3.6 mg, 8.9 µmol, 1.0 equiv.), HATU (8.1 mg, 21.3 µmol, 2.4 equiv.) and DIPEA (12.1 µl, 71.0 µmol, 8.0 equiv.) gave conjugate **S12** (4.1 mg, 45%) as an orange solid after purification by silica-gel chromatography ($CH_2Cl_2$/acetone, 4:1 → 1:1 + 1% MeOH). $R_f$ = 0.22 ($CH_2Cl_2$/MeOH, 19:1); $[\alpha]_D^{24}$ = 43.7 (c = 0.10 in MeCN); $^1$H-NMR (500 MHz, DMSO-$d_6$, isomeric mixture 2:1) main isomer: $\delta$ = 10.78 (d, $J$ = 2.1 Hz, 1 H), 9.24 (s, 1 H), 8.33 (d, $J$ = 8.5 Hz, 1 H), 7.94 (t, $J$ = 6.0 Hz, 1 H), 7.72 (d, $J$ = 8.9 Hz, 2 H), 7.66 (d, $J$ = 7.9 Hz, 1 H), 7.53 (d, $J$ = 8.2 Hz, 1 H), 7.39 - 7.34 (m, 3 H), 7.34 - 7.26 (m, 4 H), 7.23 (d, $J$ = 2.1 Hz, 1 H), 7.08 - 7.02 (m, 4 H), 6.99 - 6.94 (m, 3 H), 6.80 (d, $J$ = 8.9 Hz, 1 H), 6.65 (d, $J$ = 8.5 Hz, 2 H), 5.41 (dd, $J$ = 7.0, 8.9 Hz, 1 H), 5.23 (ddd, $J$ = 1.8, 8.2, 10.7 Hz, 1 H), 5.00 - 4.91 (m, 2 H), 4.66 (s, 2 H), 4.55 (qu, $J$ = 6.7 Hz, 1 H), 4.43 (s, 2 H), 4.28 (t, $J$ = 4.3 Hz, 2 H), 3.56 (t, $J$ = 4.3 Hz, 2 H), 3.15 - 3.08 (m, 1 H), 3.05 (s, 3 H), 3.02 - 2.89 (m, 2 H), 2.75 (s, 3 H), 2.73 - 2.66 (m, 1 H), 2.53 - 2.52 (m, 2 H), 2.38 - 2.33 (m, 1 H), 2.10 (dd, $J$ = 11.9, 13.7 Hz, 1 H), 1.84 - 1.58 (m, 3 H), 1.48 (s, 3 H), 1.32 - 1.16 (m, 2 H), 0.95 (d, $J$ = 3.7 Hz, 3 H), 0.94 (d, $J$ = 3.7 Hz, 3 H); $^{13}$C-NMR (126 MHz, DMSO-$d_6$): $\delta$ = 175.4, 170.4, 170.0, 169.1, 168.0, 159.5, 156.4, 147.3, 143.9, 143.6, 138.9, 138.0, 136.6, 133.9, 133.1, 129.1, 127.5, 127.5, 127.4, 124.9, 124.0, 124.0, 123.7, 121.3, 121.3, 119.1, 118.6, 115.7, 115.3, 111.7, 111.5, 110.2, 107.3, 67.4, 64.3, 56.3, 54.0, 49.6, 48.5, 47.6, 47.4, 43.9, 40.9, 38.6, 33.3, 31.0, 29.5, 28.7, 25.4, 24.8, 19.4, 18.0, 16.8; IR: $\tilde{\nu}_{max}$ = 3734, 3630, 2943, 1624, 1517, 1419, 1377, 1037, 918, 748, 652; UV-VIS (PBS/MeCN, 2:1): $\lambda_{max}$ ($\varepsilon$) = 446 nm (20.7 × 10$^3$ l·mol$^{-1}$·cm$^{-1}$); HRMS (ESI): *m/Z* calcd for $C_{59}H_{68}N_9O_8$ [M+H]$^+$: 1030.519, found: 1030.518.

*cyclo*-[(2*S*,4*E*,8*S*)-Hdn-L-Dap(4-((2′-((4′′-(dimethylamino)phenyl)diazenyl)-5′-methoxyphenyl)amino)-4-oxobutyryl)-D-*N*MeTrp-L-*β*Tyr]; (**S17**)

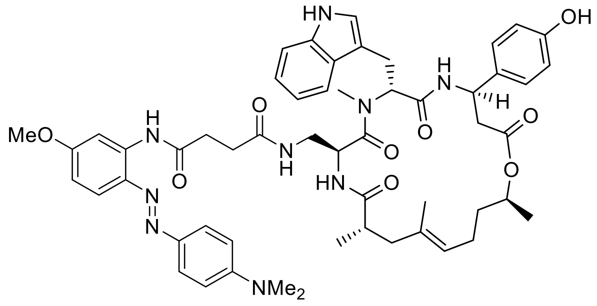

Standard Procedure **SP4** with cyclodepsipeptide **S54** (11.0 mg, 12.4 µmol, 1.0 equiv.), *N*-(2′-((4′′-(Dimethylamino)phenyl)diazenyl)-5′-methoxyphenyl)succinamic acid (4.6 mg, 12.4 µmol, 1.0 equiv.), HATU (9.4 mg, 24.8 µmol, 2.4 equiv.) and DIPEA (16.9 µl, 99.1 µmol, 8.0 equiv.) gave conjugate **S17** (8.5 mg, 70%) as a yellow solid after purification by silica-gel chromatography ($CH_2Cl_2$/petroleum ether/aceton, 3:1:1 → 1:1:3). $R_f$ = 0.29 ($CH_2Cl_2$/MeOH, 14:1); $[\alpha]_D^{24}$ = 73.8 (c = 0.10 in MeCN); $^1$H-NMR (500 MHz, DMSO-$d_6$): $\delta$ = 10.78 (d, $J$ = 1.8 Hz, 1 H), 10.23 (s, 1 H), 9.32 (br s, 1 H), 8.55 (d, $J$ = 8.9 Hz, 1 H), 8.02 (d, $J$ = 2.7 Hz, 1 H), 7.88 (d, $J$ = 9.2 Hz, 2 H), 7.71 (t, $J$ = 6.0 Hz, 1 H), 7.68 (d, $J$ = 9.2 Hz, 1 H), 7.67 (d, $J$ = 7.6 Hz, 1 H), 7.62 (d, $J$ = 8.8 Hz, 1 H), 7.30 (d, $J$ = 8.2 Hz, 1 H), 7.08 - 6.98 (m, 4 H), 6.95 (ddd, $J$ = 0.9, 7.0, 7.9 Hz, 1 H), 6.83 (d, $J$ = 9.2 Hz, 2 H), 6.74 (dd, $J$ = 2.9, 9.0 Hz, 1 H), 6.67 (d, $J$ = 8.5 Hz, 2 H), 5.49 (t, $J$ = 8.1 Hz, 1 H), 5.17 (ddd, $J$ = 3.1, 8.9, 11.3 Hz, 1 H), 4.94 (t, $J$ = 6.7 Hz, 1 H), 4.91 (dt, $J$ = 4.3, 8.9 Hz, 1 H), 4.68 (sxt, $J$ = 6.3 Hz, 1 H), 3.81 (s, 3 H), 3.10 (s, 3 H), 3.05 (s, 6 H), 3.00 (d, $J$ = 8.2 Hz, 2 H), 2.77 - 2.71 (m, 1 H), 2.71 - 2.57 (m, 4 H), 2.53 - 2.52 (m, 2 H), 2.32 (q, $J$ = 7.6 Hz, 2 H), 2.16 (dd, $J$ = 11.4, 14.5 Hz, 1 H), 1.85 (dquin, $J$ = 7.4, 14.7 Hz, 2 H), 1.74 (d, $J$ = 14.0 Hz, 1 H), 1.48 (s, 3 H), 1.53 - 1.46 (m, 1 H), 1.42 - 1.33 (m, 1 H), 1.14 (d, $J$ = 6.1 Hz, 3 H), 0.96 (d, $J$ = 6.7 Hz, 3 H); $^{13}$C-NMR (126 MHz, DMSO-$d_6$): $\delta$ = 175.3, 172.1, 171.4, 170.7, 170.6, 170.0, 161.4, 156.7, 152.6, 143.3, 137.6, 136.6, 135.0, 133.6, 133.3, 127.5, 127.4, 125.3, 123.8, 123.7, 121.3, 119.2, 119.1, 118.5, 115.5, 112.0, 111.6, 110.0, 110.0, 105.7, 71.2, 55.9, 55.2, 49.3, 48.6, 43.2, 41.9, 40.7, 38.3, 35.3, 32.6, 31.0, 30.5, 25.8, 24.1, 20.0, 19.9, 17.2; IR: $\tilde{\nu}_{max}$ = 3325, 2931, 1732, 1670, 1597, 1520, 1458, 1366, 1285, 1231, 1150, 1011, 826, 745, 652; UV-VIS (PBS/MeCN, 2:1): $\lambda_{max}$ ($\varepsilon$) = 466 nm (25.1 × $10^3$ l·mol$^{-1}$·cm$^{-1}$); HRMS (ESI): *m/Z* calcd for $C_{54}H_{66}N_9O_9$ $[M+H]^+$: 984.4978, found: 984.4978.

*cyclo*-[(2*S*,4*E*,8*S*)-Hdn-L-Dab(4-((2´-((4´´-(dimethylamino)phenyl)diazenyl)-5´-methoxyphenyl)amino)-4-oxobutyryl)-D-*N*MeTrp-L-*β*Tyr]; (**S18**)

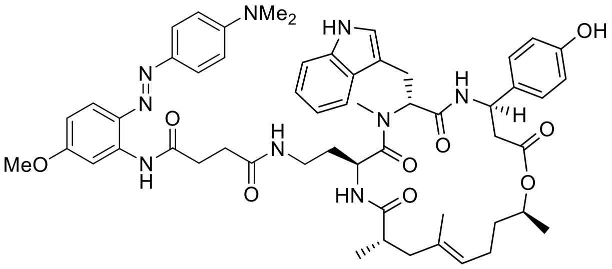

Standard Procedure **SP4** with cyclodepsipeptide **S53** (10.0 mg, 11.1 µmol, 1.0 equiv.), *N*-(2´-((4´´-(Dimethylamino)phenyl)diazenyl)-5´-methoxyphenyl)succinamic acid (4.1 mg, 11.1 µmol, 1.0 equiv.), HATU (10.1 mg, 26.6 µmol, 2.4 equiv.) and DIPEA (15.1 µl, 88.7 µmol, 8.0 equiv.) gave conjugate **S18** (6.8 mg, 61%) as a yellow solid after purification by silica-gel chromatography ($CH_2Cl_2$/petroleum ether/aceton, 1:1:1). $R_f$ = 0.31 ($CH_2Cl_2$/MeOH, 14:1); $[\alpha]_D^{24}$ = 15.6 (c = 0.10 in MeCN); $^1$H-NMR (500 MHz, DMSO-$d_6$): $\delta$ = 10.69 (d, $J$ = 1.5 Hz, 1 H), 10.28 (s, 1 H), 9.31 (s, 1 H), 8.56 (d, $J$ = 8.5 Hz, 1 H), 8.05 (d, $J$ = 2.4 Hz, 1 H), 7.90 (d, $J$ = 9.2 Hz, 2 H), 7.79 (d, $J$ = 8.5 Hz, 1 H), 7.70 - 7.61 (m, 3 H), 7.30 (d, $J$ = 8.2 Hz, 1 H), 7.07 (d, $J$ = 8.5 Hz, 2 H), 7.05 - 7.01 (m, 1 H), 6.98 - 6.93 (m, 2 H), 6.81 (d, $J$ = 9.2 Hz, 2 H), 6.73 (dd, $J$ = 2.7, 9.2 Hz, 1 H), 6.68 (d, $J$ = 8.5 Hz, 2 H), 5.45 (dd, $J$ = 7.0, 8.9 Hz, 1 H), 5.17 (ddd, $J$ = 3.1, 8.5, 11.6 Hz, 1 H), 4.94 (t, $J$ = 6.7 Hz, 1 H), 4.74 - 4.65 (m, 2 H), 3.79 (s, 3 H), 3.04 (s, 3 H), 3.04 (s, 6 H), 3.01 - 2.94 (m, 2 H), 2.90 - 2.83 (m, 1 H), 2.73 (t, $J$ = 7.0 Hz, 2 H), 2.76 - 2.65 (m, 2 H), 2.58 (dd, $J$ = 3.4, 15.0 Hz, 1 H), 2.53 - 2.52 (m, 2 H), 2.44 (t, $J$ = 7.0 Hz, 2 H), 2.18 (dd, $J$ = 11.4, 14.2 Hz, 1 H), 1.84 (dquin, $J$ = 7.8, 15.7 Hz, 2 H), 1.76 (d, $J$ = 14.0 Hz, 1 H), 1.49 (s, 3 H), 1.54 - 1.45 (m, 1 H), 1.43 - 1.34 (m, 1 H), 1.14 (d, $J$ = 6.1 Hz, 3 H), 1.12 - 1.05 (m, 1 H), 0.95 (d, $J$ = 7.0 Hz, 3 H); $^{13}$C-NMR (126 MHz, DMSO-$d_6$): $\delta$ = 175.2, 172.2, 171.6, 171.5, 170.7, 170.1, 161.5, 156.7, 152.6, 143.3, 137.7, 136.6, 135.0, 133.6, 133.3, 127.5, 127.4, 125.3, 123.8, 123.6, 121.3, 119.0, 119.0, 118.6, 115.5, 112.0, 111.7, 110.0, 110.0, 105.7, 71.2, 55.9, 55.3, 49.4, 46.7, 43.2, 41.9, 40.6, 38.2, 35.7, 35.3, 32.8, 31.7, 30.9, 30.7, 25.8, 24.0, 20.0, 17.2; IR: $\tilde{\nu}_{max}$ = 3325, 2932, 1728, 1651, 1597, 1520, 1454, 1366, 1285, 1150, 1034, 825, 745, 652; UV-VIS (PBS/MeCN, 2:1): $\lambda_{max}$ ($\varepsilon$) = 465 nm (20.8 × $10^3$ l·mol$^{-1}$·cm$^{-1}$); HRMS (ESI): *m/Z* calcd for $C_{55}H_{68}N_9O_9$ $[M+H]^+$: 998.5135, found: 998.5134.

*cyclo*-[(2*S*,4*E*,8*S*)-Hdn-L-Orn(4-((2′-((4′′-(dimethylamino)phenyl)diazenyl)-5′-methoxyphenyl)amino)-4-oxobutyryl)-D-*N*MeTrp-L-*β*Tyr]; (**S19**)

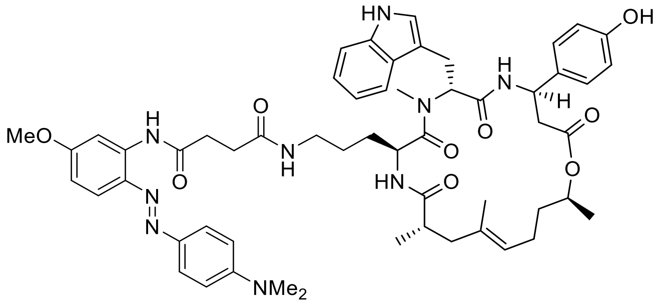

Standard Procedure **SP4** with cyclodepsipeptide **S52** (10.0 mg, 10.9 µmol, 1.0 equiv.), *N*-(2′-((4′′-(Dimethylamino)phenyl)diazenyl)-5′-methoxyphenyl)succinamic acid (4.0 mg, 10.9 µmol, 1.0 equiv.), HATU (10.0 mg, 26.2 µmol, 2.4 equiv.) and DIPEA (14.9 µl, 87.3 µmol, 8.0 equiv.) gave conjugate **S19** (8.6 mg, 78%) as an orange solid after purification by silica-gel chromatography ($CH_2Cl_2$/petroleum ether/aceton, 1:1:1). $R_f$ = 0.35 ($CH_2Cl_2$/MeOH, 14:1); $[\alpha]_D^{24}$ = 61.1 (c = 0.10 in MeCN); $^1$H-NMR (500 MHz, DMSO-$d_6$): $\delta$ = 10.76 (s, 1 H), 10.27 (s, 1 H), 9.33 (br s, 1 H), 8.61 (d, *J* = 8.9 Hz, 1 H), 8.05 (d, *J* = 2.7 Hz, 1 H), 7.90 (d, *J* = 9.2 Hz, 2 H), 7.72 - 7.67 (m, 3 H), 7.66 (d, *J* = 7.9 Hz, 1 H), 7.29 (d, *J* = 8.2 Hz, 1 H), 7.11 (d, *J* = 8.9 Hz, 2 H), 7.04 - 7.00 (m, 2 H), 6.97 - 6.92 (m, 1 H), 6.82 (d, *J* = 9.2 Hz, 2 H), 6.74 (dd, *J* = 2.9, 9.0 Hz, 1 H), 6.69 (d, *J* = 8.2 Hz, 2 H), 5.50 (dd, *J* = 5.5, 10.7 Hz, 1 H), 5.18 (ddd, *J* = 2.7, 8.9, 11.3 Hz, 1 H), 4.93 (t, *J* = 6.9 Hz, 1 H), 4.68 (sxt, *J* = 6.3 Hz, 1 H), 4.58 (dt, *J* = 5.0, 8.5 Hz, 1 H), 3.80 (s, 3 H), 3.04 (s, 6 H), 3.02 (s, 3 H), 3.01 - 2.97 (m, 1 H), 2.97 - 2.91 (m, 1 H), 2.83 - 2.74 (m, 2 H), 2.72 (t, *J* = 6.9 Hz, 2 H), 2.66 (dd, *J* = 3.2, 14.5 Hz, 1 H), 2.59 (dd, *J* = 2.7, 14.6 Hz, 1 H), 2.53 - 2.52 (m, 2 H), 2.42 (t, *J* = 6.9 Hz, 2 H), 2.17 (dd, *J* = 11.9, 14.3 Hz, 1 H), 1.91 - 1.77 (m, 2 H), 1.73 (d, *J* = 14.3 Hz, 1 H), 1.48 (s, 3 H), 1.54 - 1.45 (m, 1 H), 1.42 - 1.34 (m, 1 H), 1.15 (d, *J* = 6.1 Hz, 3 H), 1.00 - 0.96 (m, 1 H), 0.92 (d, *J* = 7.0 Hz, 3 H), 0.88 - 0.78 (m, 2 H); $^{13}$C-NMR (126 MHz, DMSO-$d_6$): $\delta$ = 174.9, 172.6, 171.6, 171.5, 170.8, 170.4, 161.5, 156.7, 152.6, 143.3, 137.7, 136.6, 135.0, 133.6, 133.4, 127.5, 127.4, 125.3, 123.8, 123.6, 121.3, 119.1, 119.0, 118.5, 115.5, 112.0, 111.6, 110.0, 109.9, 105.7, 71.3, 55.9, 55.1, 49.4, 48.0, 43.1, 42.1, 40.7, 38.7, 38.2, 35.3, 32.8, 31.0, 30.7, 29.1, 25.9, 25.3, 24.1, 20.1, 20.0, 17.3; IR: $\tilde{\nu}_{max}$ = 3310, 2932, 1728, 1647, 1597, 1516, 1454, 1366, 1285, 1150, 1030, 825, 744, 652; UV-VIS (PBS/MeCN, 2:1): $\lambda_{max}$ ($\varepsilon$) = 465 nm (25.6 × $10^3$ l·mol$^{-1}$·cm$^{-1}$); HRMS (ESI): *m/Z* calcd for $C_{56}H_{70}N_9O_9$ $[M+H]^+$: 1012.529, found: 1012.530.

*cyclo*-[(2*S*,4*E*,8*S*)-Hdn-L-Lys(4-((2´-((4´´-(dimethylamino)phenyl)diazenyl)-5´-methoxyphenyl)amino)-4-oxobutyryl)-D-*N*MeTrp-L-*β*Tyr]; (**S20**)

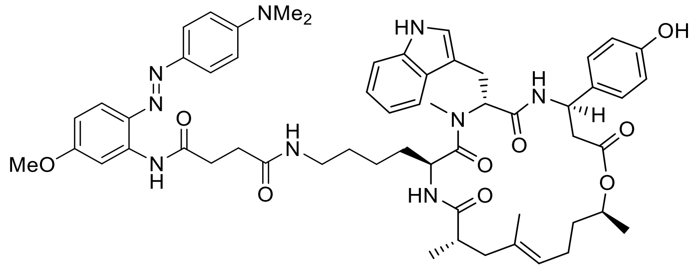

Standard Procedure **SP4** with cyclodepsipeptide **S51** (10.0 mg, 10.8 µmol, 1.0 equiv.), *N*-(2´-((4´´-(Dimethylamino)phenyl)diazenyl)-5´-methoxyphenyl)succinamic acid (4.0 mg, 10.8 µmol, 1.0 equiv.), HATU (9.8 mg, 25.8 µmol, 2.4 equiv.) and DIPEA (14.3 µl, 86.0 µmol, 8.0 equiv.) gave conjugate **S20** (5.5 mg, 50%) as an orange solid after purification by silica-gel chromatography ($CH_2Cl_2$/aceton, 4:1 → 1:1). $R_f$ = 0.33 ($CH_2Cl_2$/MeOH, 14:1); $[\alpha]_D^{24}$ = 69.3 (c = 0.10 in MeCN); $^1$H-NMR (500 MHz, DMSO-$d_6$): $\delta$ = 10.81 (s, 1 H), 10.28 (s, 1 H), 9.38 (br s, 1 H), 8.65 (d, $J$ = 8.9 Hz, 1 H), 8.06 (d, $J$ = 2.7 Hz, 1 H), 7.91 (d, $J$ = 9.2 Hz, 2 H), 7.87 (t, $J$ = 5.6 Hz, 1 H), 7.71 - 7.64 (m, 3 H), 7.29 (d, $J$ = 7.9 Hz, 1 H), 7.13 (d, $J$ = 8.5 Hz, 2 H), 7.04 (d, $J$ = 1.8 Hz, 1 H), 7.01 (t, $J$ = 7.2 Hz, 1 H), 6.94 (t, $J$ = 7.0 Hz, 1 H), 6.81 (d, $J$ = 9.2 Hz, 2 H), 6.73 (dd, $J$ = 2.7, 9.2 Hz, 1 H), 6.70 (d, $J$ = 8.5 Hz, 2 H), 5.51 (dd, $J$ = 5.0, 11.4 Hz, 1 H), 5.18 (ddd, $J$ = 2.7, 9.2, 11.3 Hz, 1 H), 4.92 (t, $J$ = 6.6 Hz, 1 H), 4.67 (sxt, $J$ = 6.3 Hz, 1 H), 4.58 - 4.52 (m, 1 H), 3.80 (s, 3 H), 3.04 (s, 6 H), 3.02 (s, 3 H), 3.03 - 2.99 (m, 1 H), 2.95 - 2.78 (m, 3 H), 2.74 (t, $J$ = 7.0 Hz, 2 H), 2.68 (dd, $J$ = 11.4, 14.8 Hz, 1 H), 2.59 (dd, $J$ = 2.7, 14.6 Hz, 1 H), 2.53 - 2.52 (m, 2 H), 2.46 (t, $J$ = 6.7 Hz, 2 H), 2.17 (dd, $J$ = 11.9, 14.6 Hz, 1 H), 1.90 - 1.76 (m, 2 H), 1.72 (d, $J$ = 14.3 Hz, 1 H), 1.48 (s, 3 H), 1.53 - 1.47 (m, 1 H), 1.41 - 1.33 (m, 1 H), 1.16 (d, $J$ = 6.4 Hz, 3 H), 1.13 - 1.07 (m, 1 H), 0.92 (d, $J$ = 6.7 Hz, 3 H), 0.85 - 0.75 (m, 3 H), 0.70 - 0.63 (m, 1 H); $^{13}$C-NMR (126 MHz, DMSO-$d_6$): $\delta$ = 174.9, 172.7, 171.6, 171.5, 170.8, 170.5, 161.4, 156.8, 152.5, 143.3, 137.7, 136.6, 135.0, 133.6, 133.5, 127.5, 127.4, 125.3, 123.8, 123.4, 121.3, 119.1, 118.9, 118.5, 115.5, 112.0, 111.6, 110.0, 109.9, 105.7, 71.4, 55.9, 55.0, 49.5, 48.1, 42.9, 42.1, 40.7, 38.8, 38.1, 35.3, 32.9, 31.0, 30.8, 29.1, 26.0, 26.0, 24.2, 22.5, 20.2, 19.9, 17.4; IR: $\tilde{\nu}_{max}$ = 3322, 2932, 1728, 1649, 1597, 1519, 1454, 1366, 1285, 1150, 1031, 825, 744, 652; UV-VIS (PBS/MeCN, 2:1): $\lambda_{max}$ ($\varepsilon$) = 465 nm (24.6 × $10^3$ l·mol$^{-1}$·cm$^{-1}$); HRMS (ESI): *m/Z* calcd for $C_{57}H_{72}N_9O_9$ $[M+H]^+$: 1026.545, found: 1026.545.

*cyclo*-[(2*S*,4*E*,8*S*)-Hdn-L-Dap(4-((5-methoxy-2-((4-morpholinophenyl)diazenyl)-phenyl)amino)-4-oxobutyryl)-D-*N*MeTrp-L-*β*Tyr]; (**S21**)

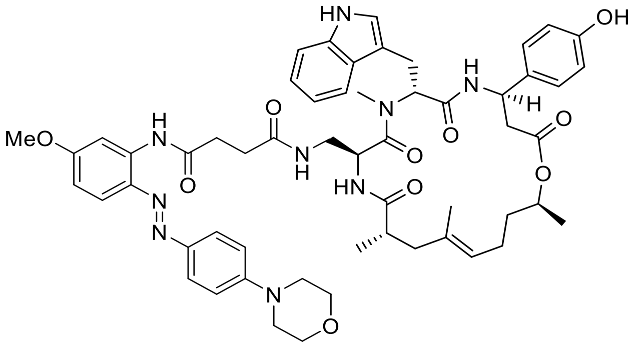

Standard Procedure **SP4** with cyclodepsipeptide **S54** (6.6 mg, 7.4 µmol, 1.0 equiv.), *N*-(5′-Methoxy-2′-((4′′-morpholinophenyl)diazenyl)phenyl)succinamic acid (3.1 mg, 7.4 µmol, 1.0 equiv.), HATU (6.8 mg, 17.8 µmol, 2.4 equiv.) and DIPEA (10.1 µl, 59.4 µmol, 8.0 equiv.) gave conjugate **S21** (6.5 mg, 85%) as an orange solid after purification by silica-gel chromatography ($CH_2Cl_2$/acetone, 4:1 → 1:1). $R_f$ = 0.34 ($CH_2Cl_2$/MeOH, 14:1); $[\alpha]_D^{24}$ = 20.6 (c = 0.10 in MeCN); $^1$H-NMR (400 MHz, DMSO-$d_6$): $\delta$ = 10.78 (br s, 1 H), 10.25 (s, 1 H), 9.33 (br s, 1 H), 8.55 (d, *J* = 8.8 Hz, 1 H), 8.02 (d, *J* = 2.6 Hz, 1 H), 7.91 (d, *J* = 9.1 Hz, 2 H), 7.75 - 7.60 (m, 4 H), 7.30 (d, *J* = 7.9 Hz, 1 H), 7.11 - 7.01 (m, 5 H), 6.99 (d, *J* = 1.5 Hz, 1 H), 6.94 (t, *J* = 7.6 Hz, 1 H), 6.76 (dd, *J* = 2.6, 9.1 Hz, 1 H), 6.67 (d, *J* = 8.2 Hz, 2 H), 5.49 (t, *J* = 7.9 Hz, 1 H), 5.17 (ddd, *J* = 3.2, 8.0, 11.0 Hz, 1 H), 4.94 (t, *J* = 7.6 Hz, 1 H), 4.91 (dt, *J* = 4.1, 9.4 Hz, 1 H), 4.68 (sxt, *J* = 6.1 Hz, 1 H), 3.82 (s, 3 H), 3.76 (t, *J* = 4.7 Hz, 4 H), 3.31 (t, *J* = 4.7 Hz, 4 H), 3.09 (s, 3 H), 3.00 (d, *J* = 7.6 Hz, 2 H), 2.79 - 2.70 (m, 2 H), 2.70 - 2.59 (m, 5 H), 2.35 - 2.28 (m, 2 H), 2.16 (dd, *J* = 11.7, 14.3 Hz, 1 H), 1.85 (spt, *J* = 6.6 Hz, 2 H), 1.74 (d, *J* = 14.6 Hz, 1 H), 1.48 (s, 3 H), 1.55 - 1.45 (m, 1 H), 1.44 - 1.32 (m, 1 H), 1.14 (d, *J* = 6.4 Hz, 3 H), 0.95 (d, *J* = 6.7 Hz, 3 H); $^{13}$C-NMR (101 MHz, DMSO-$d_6$): $\delta$ = 175.3, 172.1, 171.5, 170.7, 170.7, 170.0, 161.9, 156.7, 153.2, 145.1, 138.0, 136.5, 135.0, 133.6, 133.2, 127.5, 127.4, 124.9, 123.8, 123.7, 121.3, 119.2, 119.2, 118.5, 115.5, 114.5, 111.6, 110.1, 110.0, 105.8, 71.2, 66.4, 55.9, 55.2, 49.3, 48.5, 47.7, 43.2, 41.8, 41.1, 38.3, 35.3, 32.6, 31.0, 30.5, 25.8, 24.1, 20.0, 19.8, 17.2; IR: $\tilde{\nu}_{max}$ = 3291, 2932, 1732, 1670, 1597, 1516, 1454, 1377, 1234, 1153, 1114, 1030, 926, 745, 652; UV-VIS (PBS/MeCN, 2:1): $\lambda_{max}$ ($\varepsilon$) = 411 nm (22.1 × $10^3$ l·mol$^{-1}$·cm$^{-1}$); HRMS (ESI): *m/Z* calcd for $C_{56}H_{68}N_9O_{10}$ [M+H]$^+$: 1026.508, found: 1026.508.

*cyclo*-[(2*S*,4*E*,8*S*)-Hdn-L-Dap(4-((5′-methoxy-2′-((4′′-(4′′′-methylpiperazin-1′′′-yl)-phenyl)diazenyl)phenyl)amino)-4-oxobutyryl)-D-*N*MeTrp-L-*β*Tyr]; (**S22**)

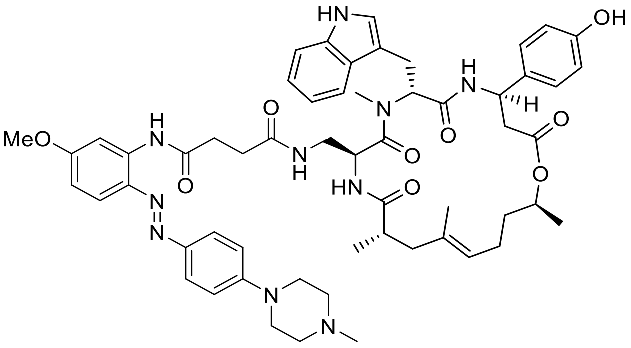

Standard Procedure **SP4** with cyclodepsipeptide **S54** (10.0 mg, 11.3 µmol, 1.0 equiv.), *N*-(5′-Methoxy-2′-((4′′-(4′′′-methylpiperazin-1′′′-yl)phenyl)diazenyl)phenyl)succinamic acid (4.8 mg, 11.3 µmol, 1.0 equiv.), HATU (10.3 mg, 27.0 µmol, 2.4 equiv.) and DIPEA (15.3 µl, 90.1 µmol, 8.0 equiv.) gave conjugate **S22** (10.2 mg, 87%) as an orange solid after purification by silica-gel chromatography ($CH_2Cl_2$/MeOH, 96:4 → 90:10). $R_f$ = 0.24 ($CH_2Cl_2$/MeOH, 14:1); $[\alpha]_D^{24}$ = 37.5 (c = 0.10 in MeCN); $^1$H-NMR (400 MHz, DMSO-$d_6$): $\delta$ = 10.79 (d, $J$ = 1.2 Hz, 1 H), 10.25 (s, 1 H), 8.55 (d, $J$ = 8.5 Hz, 1 H), 8.01 (d, $J$ = 2.6 Hz, 1 H), 7.87 (d, $J$ = 9.1 Hz, 2 H), 7.76 (t, $J$ = 5.7 Hz, 1 H), 7.71 - 7.63 (m, 3 H), 7.29 (d, $J$ = 8.2 Hz, 1 H), 7.09 - 7.00 (m, 5 H), 6.98 (d, $J$ = 1.8 Hz, 1 H), 6.96 - 6.91 (m, 1 H), 6.75 (dd, $J$ = 2.6, 9.1 Hz, 1 H), 6.66 (d, $J$ = 8.5 Hz, 2 H), 5.48 (t, $J$ = 8.0 Hz, 1 H), 5.16 (ddd, $J$ = 3.5, 8.0, 11.0 Hz, 1 H), 4.94 (t, $J$ = 6.7 Hz, 1 H), 4.90 (dt, $J$ = 4.4, 9.1 Hz, 1 H), 4.67 (sxt, $J$ = 6.1 Hz, 1 H), 3.81 (s, 3 H), 3.34 (t, $J$ = 4.4 Hz, 4 H), 3.09 (s, 3 H), 3.00 (d, $J$ = 7.6 Hz, 2 H), 2.79 - 2.70 (m, 2 H), 2.70 - 2.54 (m, 5 H), 2.44 (t, $J$ = 4.7 Hz, 4 H), 2.35 - 2.28 (m, 2 H), 2.22 (s, 3 H), 2.16 (dd, $J$ = 11.8, 14.2 Hz, 1 H), 1.84 (spt, $J$ = 6.3 Hz, 2 H), 1.73 (d, $J$ = 12.0 Hz, 1 H), 1.47 (s, 3 H), 1.55 - 1.43 (m, 1 H), 1.42 - 1.32 (m, 1 H), 1.14 (d, $J$ = 6.4 Hz, 3 H), 0.95 (d, $J$ = 6.7 Hz, 3 H); $^{13}$C-NMR (101 MHz, DMSO-$d_6$): $\delta$ = 175.3, 172.1, 171.5, 170.7, 170.7, 170.0, 161.8, 156.8, 153.1, 144.7, 137.9, 136.6, 135.0, 133.6, 133.2, 127.5, 127.4, 125.0, 123.8, 123.7, 121.3, 119.3, 119.2, 118.5, 115.5, 114.6, 111.6, 110.1, 110.0, 105.8, 71.2, 55.9, 55.2, 54.8, 49.3, 48.6, 47.4, 46.2, 43.2, 41.9, 40.8, 38.3, 35.3, 32.6, 31.0, 30.5, 25.8, 24.1, 20.0, 19.9, 17.2; IR: $\tilde{\nu}_{max}$ = 3317, 2931, 1732, 1670, 1597, 1516, 1454, 1377, 1288, 1238, 1145, 1007, 829, 745, 652; UV-VIS (PBS/MeCN, 2:1): $\lambda_{max}$ ($\varepsilon$) = 412 nm (20.7 × $10^3$ l·mol$^{-1}$·cm$^{-1}$); HRMS (ESI): *m/Z* calcd for $C_{57}H_{71}N_{10}O_9$ $[M+H]^+$: 1039.540, found: 1039.540.

*cyclo*-[(2*S*,4*E*,8*S*)-Hdn-L-Dap(2-(4´-((4´´-(dimethylamino)phenyl)diazenyl)phenoxy)-acetyl)-D-*N*MeTrp-L-*β*Tyr]; (**S23**)

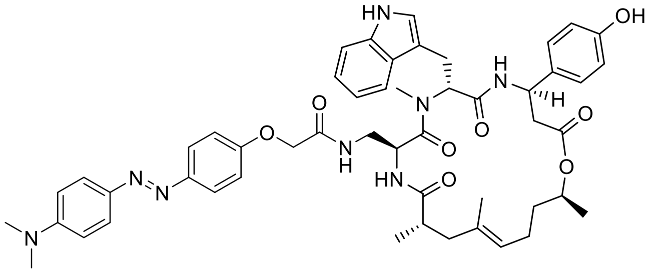

Standard Procedure **SP4** with cyclodepsipeptide **S54** (10.0 mg, 11.3 µmol, 1.0 equiv.), 2-(4´-((4´´-(dimethylamino)phenyl)diazenyl)phenoxy)acetic acid (4.0 mg, 11.3 µmol, 1.0 equiv.), HATU (10.3 mg, 27.0 µmol, 2.4 equiv.) and DIPEA (15.3 µl, 90.1 µmol, 8.0 equiv.) gave conjugate **S23** (6.8 mg, 66%) as an orange solid after purification by silica-gel chromatography ($CH_2Cl_2$/MeOH, 98:2 → 90:10). $R_f$ = 0.36 ($CH_2Cl_2$/MeOH, 14:1); $[\alpha]_D^{24}$ = 44.3 (c = 0.05 in MeCN); $^1$H-NMR (500 MHz, DMSO-$d_6$): $\delta$ = 10.80 (d, $J$ = 1.8 Hz, 1 H), 9.31 (s, 1 H), 8.58 (d, $J$ = 8.9 Hz, 1 H), 7.95 (t, $J$ = 5.8 Hz, 1 H), 7.76 (d, $J$ = 3.1 Hz, 2 H), 7.74 (d, $J$ = 2.7 Hz, 2 H), 7.68 (d, $J$ = 7.9 Hz, 1 H), 7.62 (d, $J$ = 8.9 Hz, 1 H), 7.30 (d, $J$ = 7.9 Hz, 1 H), 7.08 - 7.00 (m, 6 H), 6.96 (ddd, $J$ = 0.6, 7.0, 7.9 Hz, 1 H), 6.83 (d, $J$ = 9.2 Hz, 2 H), 6.67 (d, $J$ = 8.9 Hz, 2 H), 5.50 (dd, $J$ = 6.7, 9.5 Hz, 1 H), 5.24 - 5.11 (m, 1 H), 4.98 - 4.90 (m, 2 H), 4.69 (sxt, $J$ = 6.3 Hz, 1 H), 4.44 (d, $J$ = 2.4 Hz, 2 H), 3.12 (s, 3 H), 3.05 (s, 6 H), 3.08 - 2.96 (m, 3 H), 2.84 - 2.74 (m, 1 H), 2.70 - 2.56 (m, 3 H), 2.17 (dd, $J$ = 11.6, 14.3 Hz, 1 H), 1.85 (qud, $J$ = 7.5, 15.2 Hz, 2 H), 1.76 (d, $J$ = 14.0 Hz, 1 H), 1.50 (s, 3 H), 1.55 - 1.45 (m, 1 H), 1.44 - 1.34 (m, 1 H), 1.15 (d, $J$ = 6.4 Hz, 3 H), 0.94 (d, $J$ = 6.7 Hz, 3 H); $^{13}$C-NMR (126 MHz, DMSO-$d_6$): $\delta$ = 175.4, 170.7, 170.5, 170.0, 168.1, 159.3, 156.7, 152.6, 147.5, 143.0, 136.6, 133.5, 133.3, 127.6, 127.4, 124.8, 123.9, 123.8, 123.7, 121.3, 119.2, 118.5, 115.6, 115.5, 112.0, 111.6, 110.0, 71.2, 67.4, 55.3, 49.3, 48.6, 43.2, 41.9, 40.7, 38.3, 35.4, 31.0, 25.7, 24.1, 20.0, 19.8, 17.2; IR: $\tilde{\nu}_{max}$ = 3325, 2932, 1732, 1670, 1601, 1516, 1366, 1234, 1153, 1060, 837, 745, 656; UV-VIS (PBS/MeCN, 2:1): $\lambda_{max}$ ($\varepsilon$) = 439 nm (27.0 × 10$^3$ l·mol$^{-1}$·cm$^{-1}$); HRMS (ESI): *m/Z* calcd for $C_{51}H_{62}N_8O_8$ $[M+H]^+$: 913.4607, found: 913.4609.

*cyclo*-[(2*S*,4*E*,8*S*)-Hdn-L-Dap(2-(4´-((4´´-(4´´´-morpholinophenyl)phenyl)diazenyl)-phenoxy)acetyl)-D-*N*MeTrp-L-*β*Tyr]; (**S24**)

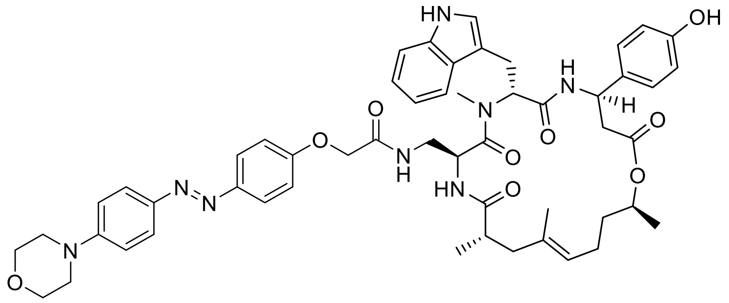

Standard Procedure **SP4** with cyclodepsipeptide **S54** (10.0 mg, 11.3 µmol, 1.0 equiv.), 2-(4´-((4´´-morpholinophenyl)diazenyl)phenoxy)acetic acid (3.8 mg, 11.3 µmol, 1.0 equiv.), HATU (10.3 mg, 27.0 µmol, 2.4 equiv.) and DIPEA (15.3 µl, 90.1 µmol, 8.0 equiv.) gave conjugate **S24** (6.4 mg, 59%) as an orange solid after purification by silica-gel chromatography ($CH_2Cl_2$/MeOH, 96:4 → 90:10). $R_f$ = 0.36 ($CH_2Cl_2$/MeOH, 14:1); $[\alpha]_D^{24}$ = 34.8 (c = 0.05 in MeCN); $^1$H-NMR (500 MHz, DMSO-$d_6$): $\delta$ = 10.80 (d, *J* = 1.8 Hz, 1 H), 9.31 (s, 1 H), 8.57 (d, *J* = 8.5 Hz, 1 H), 7.96 (t, *J* = 6.0 Hz, 1 H), 7.78 (d, *J* = 2.4 Hz, 2 H), 7.77 (d, *J* = 2.7 Hz, 2 H), 7.68 (d, *J* = 7.9 Hz, 1 H), 7.62 (d, *J* = 8.9 Hz, 1 H), 7.30 (d, *J* = 7.9 Hz, 1 H), 7.10 - 7.02 (m, 8 H), 6.96 (ddd, *J* = 0.9, 6.7, 7.9 Hz, 1 H), 6.67 (d, *J* = 8.5 Hz, 2 H), 5.50 (dd, *J* = 6.6, 9.3 Hz, 1 H), 5.21 - 5.15 (m, 1 H), 5.01 - 4.88 (m, 2 H), 4.69 (sxt, *J* = 6.4 Hz, 1 H), 4.45 (d, *J* = 2.7 Hz, 2 H), 3.76 (t, *J* = 4.9 Hz, 4 H), 3.30 (t, *J* = 4.6 Hz, 4 H), 3.12 (s, 3 H), 3.12 - 2.97 (m, 3 H), 2.83 - 2.76 (m, 1 H), 2.70 - 2.56 (m, 3 H), 2.17 (dd, *J* = 11.6, 14.3 Hz, 1 H), 1.86 (tt, *J* = 7.6, 15.4 Hz, 2 H), 1.76 (d, *J* = 14.3 Hz, 1 H), 1.50 (s, 3 H), 1.56 - 1.46 (m, 1 H), 1.44 - 1.32 (m, 1 H), 1.15 (d, *J* = 6.4 Hz, 3 H), 0.94 (d, *J* = 6.7 Hz, 3 H); $^{13}$C-NMR (126 MHz, DMSO-$d_6$): $\delta$ = 175.4, 170.7, 170.5, 170.0, 168.0, 159.7, 156.7, 153.3, 147.3, 144.8, 136.6, 133.5, 133.2, 127.6, 127.4, 124.4, 124.2, 123.8, 123.7, 121.3, 119.2, 118.5, 115.7, 115.5, 114.6, 111.6, 110.0, 71.2, 67.4, 66.4, 55.3, 49.3, 48.6, 47.7, 43.2, 41.9, 40.7, 38.3, 35.4, 31.0, 25.7, 24.1, 20.0, 19.8, 17.2; IR: $\tilde{\nu}_{max}$ = 3317, 2970, 1732, 1670, 1597, 1377, 1231, 1157, 1115, 1053, 926, 837, 744, 652; UV-VIS (PBS/MeCN, 2:1): $\lambda_{max}$ ($\varepsilon$) = 391 nm (25.5 × $10^3$ l·mol$^{-1}$·cm$^{-1}$); HRMS (ESI): *m/Z* clacd for $C_{53}H_{63}N_8O_9$ $[M+H]^+$: 955.4713, found: 955.4720.

*cyclo*-[(2*S*,4*E*,8*S*)-Hdn-L-Dap(2-(4′-((4′′-(4′′′-methylpiperazin-1′′′-yl)phenyl)-diazenyl)phenoxy)acetyl)-D-*N*MeTrp-L-*β*Tyr]; (**S25**)

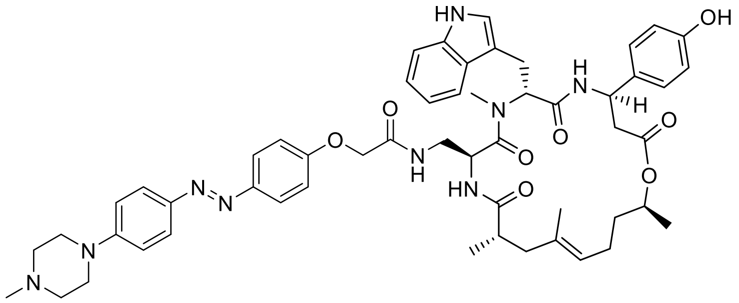

Standard Procedure **SP4** with cyclodepsipeptide **S54** (10.0 mg, 11.3 µmol, 1.0 equiv.), 2-(4′-((4′′-(4′′′-methylpiperazin-1′′′-yl)phenyl)diazenyl)phenoxy)acetic acid (3.0 mg, 11.3 µmol, 1.0 equiv.), HATU (10.3 mg, 27.0 µmol, 2.4 equiv.) and DIPEA (15.3 µl, 90.1 µmol, 8.0 equiv.) gave conjugate **S25** (7.8 mg, 71%) as an orange solid after purification by silica-gel chromatography ($CH_2Cl_2$/MeOH, 96:4 → 90:10). $R_f$ = 0.23 ($CH_2Cl_2$/MeOH, 14:1); $[\alpha]_D^{24}$ = 36.4 (c = 0.05 in MeCN); $^1$H-NMR (500 MHz, DMSO-$d_6$): $\delta$ = 10.81 (d, $J$ = 1.8 Hz, 1 H), 9.36 (br s, 1 H), 8.58 (d, $J$ = 8.5 Hz, 1 H), 7.98 (t, $J$ = 5.8 Hz, 1 H), 7.77 (d, $J$ = 8.9 Hz, 2 H), 7.75 (d, $J$ = 9.5 Hz, 2 H), 7.68 (d, $J$ = 7.9 Hz, 1 H), 7.63 (d, $J$ = 8.9 Hz, 1 H), 7.30 (d, $J$ = 7.9 Hz, 1 H), 7.10 - 7.00 (m, 8 H), 6.96 (ddd, $J$ = 0.9, 7.0, 7.9 Hz, 1 H), 6.67 (d, $J$ = 8.5 Hz, 2 H), 5.50 (dd, $J$ = 6.7, 9.5 Hz, 1 H), 5.18 (ddd, $J$ = 3.4, 8.9, 11.6 Hz, 1 H), 4.99 - 4.90 (m, 2 H), 4.69 (sxt, $J$ = 6.3 Hz, 1 H), 4.45 (d, $J$ = 2.4 Hz, 2 H), 3.12 (s, 3 H), 3.09 - 2.94 (m, 3 H), 2.80 (td, $J$ = 4.8, 13.2 Hz, 1 H), 2.70 - 2.56 (m, 3 H), 2.48 - 2.43 (m, 4 H), 2.23 (s, 3 H), 2.17 (dd, $J$ = 11.4, 14.5 Hz, 1 H), 1.92 - 1.80 (m, 2 H), 1.77 (d, $J$ = 14.0 Hz, 1 H), 1.50 (s, 3 H), 1.54 - 1.46 (m, 1 H), 1.43 - 1.34 (m, 1 H), 1.15 (d, $J$ = 6.1 Hz, 3 H), 0.94 (d, $J$ = 6.7 Hz, 3 H); $^{13}$C-NMR (126 MHz, DMSO-$d_6$): $\delta$ = 175.4, 170.7, 170.5, 170.0, 168.1, 159.6, 156.7, 153.1, 147.3, 144.5, 136.6, 133.5, 133.2, 127.6, 127.4, 124.5, 124.1, 123.8, 123.7, 121.3, 119.2, 118.5, 115.7, 115.5, 114.7, 111.6, 110.0, 71.2, 67.4, 55.3, 54.8, 49.3, 48.6, 47.4, 46.2, 43.2, 41.9, 40.7, 38.3, 35.4, 31.0, 25.7, 24.1, 20.0, 19.8, 17.2; IR: $\tilde{\nu}_{max}$ = 3325, 2940, 1732, 1670, 1597, 1504, 1454, 1377, 1234, 1157, 1053, 873, 745, 652; UV-VIS (PBS/MeCN, 2:1): $\lambda_{max}$ ($\varepsilon$) = 391 nm (22.9 × $10^3$ l·mol$^{-1}$·cm$^{-1}$); HRMS (ESI): $m/Z$ calcd for $C_{54}H_{66}N_9O_8$ $[M+H]^+$: 968.5029, found: 968.5023.

*cyclo*-[(2*S*,4*E*,8*S*)-Hdn-L-Dap(2-(4´-((4´´-methyl-3´´,4´´-dihydro-2´´*H*-benzo[*b*][1´´,4´´]-oxazin-7´´-yl)diazenyl)phenoxy)acetyl)-D-*N*MeTrp-L-*β*Tyr]; (**S26**)

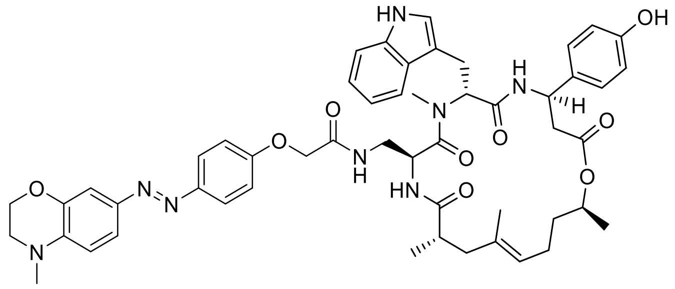

Standard Procedure **SP4** with cyclodepsipeptide **S54** (10.0 mg, 11.3 µmol, 1.0 equiv.), 2-(4´-((4´´-methyl-3´´,4´´-dihydro-2´´*H*-benzo[b][1´´,4´´]oxazin-7´´-yl)diazenyl)phenoxy) acetic acid (3.7 mg, 11.3 µmol, 1.0 equiv.), HATU (10.3 mg, 27.0 µmol, 2.4 equiv.) and DIPEA (15.3 µl, 90.1 µmol, 8.0 equiv.) gave conjugate **S26** (9.0 mg, 85%) as an orange solid after purification by silica-gel chromatography ($CH_2Cl_2$/MeOH, 98:2 → 90:10). $R_f$ = 0.36 ($CH_2Cl_2$/MeOH, 14:1); $[\alpha]_D^{24}$ = 33.6 (c = 0.10 in MeCN); $^1$H-NMR (500 MHz, DMSO-$d_6$): $\delta$ = 10.80 (d, $J$ = 1.2 Hz, 1 H), 9.30 (br s, 1 H), 8.58 (d, $J$ = 8.5 Hz, 1 H), 7.95 (t, $J$ = 5.8 Hz, 1 H), 7.74 (d, $J$ = 8.9 Hz, 2 H), 7.68 (d, $J$ = 7.9 Hz, 1 H), 7.61 (d, $J$ = 8.8 Hz, 1 H), 7.43 (dd, $J$ = 2.1, 8.5 Hz, 1 H), 7.30 (d, $J$ = 7.9 Hz, 1 H), 7.19 (d, $J$ = 2.1 Hz, 1 H), 7.09 - 7.00 (m, 6 H), 6.96 (t, $J$ = 7.6 Hz, 1 H), 6.81 (d, $J$ = 8.9 Hz, 1 H), 6.67 (d, $J$ = 8.5 Hz, 2 H), 5.50 (dd, $J$ = 6.7, 9.5 Hz, 1 H), 5.18 (ddd, $J$ = 3.4, 8.9, 11.6 Hz, 1 H), 4.99 - 4.90 (m, 2 H), 4.69 (sxt, $J$ = 6.3 Hz, 1 H), 4.44 (d, $J$ = 2.7 Hz, 2 H), 4.25 (t, $J$ = 4.4 Hz, 2 H), 3.42 - 3.39 (m, 2 H), 3.12 (s, 3 H), 2.99 (s, 3 H), 3.08 - 2.94 (m, 3 H), 2.84 - 2.74 (m, 1 H), 2.71 - 2.57 (m, 3 H), 2.17 (dd, $J$ = 12.1, 13.9 Hz, 1 H), 1.85 (tt, $J$ = 7.6, 15.0 Hz, 2 H), 1.76 (d, $J$ = 14.0 Hz, 1 H), 1.50 (s, 3 H), 1.57 - 1.44 (m, 1 H), 1.43 - 1.33 (m, 1 H), 1.15 (d, $J$ = 6.4 Hz, 3 H), 0.94 (d, $J$ = 7.0 Hz, 3 H); $^{13}$C-NMR (126 MHz, DMSO-$d_6$): $\delta$ = 175.4, 170.7, 170.5, 170.0, 168.1, 159.4, 156.7, 147.3, 144.1, 143.9, 140.1, 136.6, 133.5, 133.3, 127.6, 127.4, 124.0, 123.8, 123.7, 121.3, 121.3, 119.2, 118.5, 115.7, 115.5, 111.6, 111.4, 110.0, 106.7, 71.2, 67.4, 64.5, 55.3, 49.3, 48.6, 48.5, 43.2, 41.9, 40.7, 38.4, 38.3, 35.4, 31.0, 25.7, 24.1, 20.0, 19.8, 17.2; IR: $\tilde{\nu}_{max}$ = 3325, 2940, 1732, 1670, 1597, 1504, 1454, 1377, 1234, 1157, 1053, 873, 745, 652: UV-VIS (PBS/MeCN, 2:1): $\lambda_{max}$ ($\varepsilon$) = 447 nm (27.3 × 10$^3$ l·mol$^{-1}$·cm$^{-1}$); HRMS (ESI): *m/Z* calcd for $C_{52}H_{67}N_8O_9$ [M+H]$^+$: 941.4556, found: 941.4569.

*cyclo*-[(2*S*,4*E*,8*S*)-Hdn-L-Dap(2-(4´-((3´´,4´´-dimethoxyphenyl)diazenyl)phenoxy)-acetyl)-D-*N*MeTrp-L-*β*Tyr]; (**S27**)

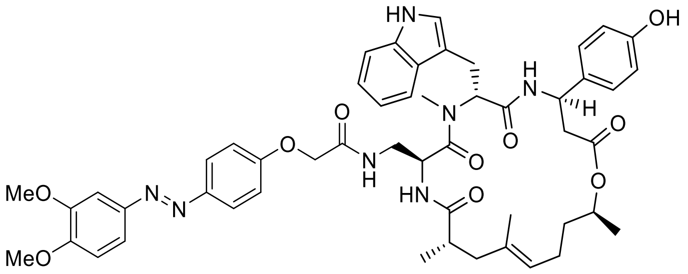

Standard Procedure **SP4** with cyclodepsipeptide **S54** (10.0 mg, 11.3 µmol, 1.0 equiv.), 2-(4´-((3´´,4´´-dimethoxyphenyl)diazenyl)phenoxy)acetic acid (4.3 mg, 11.3 µmol, 1.0 equiv.), HATU (10.3 mg, 27.0 µmol, 2.4 equiv.) and DIPEA (15.3 µl, 90.1 µmol, 8.0 equiv.) gave conjugate **S27** (4.4 mg, 42%) as a yellow solid after purification by silica-gel chromatography ($CH_2Cl_2$/MeOH, 98:2 → 90:10). $R_f$ = 0.36 ($CH_2Cl_2$/MeOH, 14:1); $[\alpha]_D^{24}$ = 40.2 (c = 0.09 in MeCN); $^1$H-NMR (500 MHz, DMSO-$d_6$): $\delta$ = 10.81 (s, 1 H), 9.20 (br s, 1 H), 8.58 (d, $J$ = 8.5 Hz, 1 H), 8.00 (t, $J$ = 5.8 Hz, 1 H), 7.83 (d, $J$ = 8.9 Hz, 2 H), 7.68 (d, $J$ = 7.9 Hz, 1 H), 7.65 (d, $J$ = 8.5 Hz, 1 H), 7.55 (dd, $J$ = 2.3, 8.4 Hz, 1 H), 7.43 (d, $J$ = 2.1 Hz, 1 H), 7.31 (d, $J$ = 7.9 Hz, 1 H), 7.17 (d, $J$ = 8.9 Hz, 1 H), 7.08 - 7.00 (m, 6 H), 6.96 (t, $J$ = 7.5 Hz, 1 H), 6.67 (d, $J$ = 8.5 Hz, 2 H), 5.50 (dd, $J$ = 6.9, 9.3 Hz, 1 H), 5.18 (ddd, $J$ = 3.1, 8.0, 11.6 Hz, 1 H), 4.98 - 4.90 (m, 2 H), 4.69 (sxt, $J$ = 6.4 Hz, 1 H), 4.47 (d, $J$ = 3.1 Hz, 2 H), 3.87 (s, 3 H), 3.85 (s, 3 H), 3.12 (s, 3 H), 3.10 - 2.97 (m, 3 H), 2.84 - 2.77 (m, 1 H), 2.70 - 2.57 (m, 3 H), 2.17 (dd, $J$ = 11.4, 14.2 Hz, 1 H), 1.86 (tt, $J$ = 7.3, 15.3 Hz, 2 H), 1.77 (d, $J$ = 14.0 Hz, 1 H), 1.50 (s, 3 H), 1.57 - 1.46 (m, 1 H), 1.44 - 1.34 (m, 1 H), 1.15 (d, $J$ = 6.4 Hz, 3 H), 0.95 (d, $J$ = 6.7 Hz, 3 H); $^{13}$C-NMR (126 MHz, DMSO-$d_6$): $\delta$ = 175.5, 170.7, 170.5, 170.0, 168.0, 160.2, 156.7, 152.0, 149.9, 147.0, 146.5, 136.6, 133.5, 133.2, 127.6, 127.4, 124.5, 123.8, 123.7, 121.3, 120.3, 119.2, 118.5, 115.7, 115.5, 111.7, 111.6, 110.0, 102.3, 71.2, 67.4, 56.3, 55.9, 55.3, 49.3, 48.6, 43.2, 41.9, 40.7, 38.3, 35.4, 31.0, 25.8, 24.1, 20.0, 19.8, 17.2; IR: $\tilde{\nu}_{max}$ = 3318, 2932, 1728, 1670, 1597, 1504, 1257, 1234, 1114, 1022, 837, 745, 629; UV-VIS (PBS/MeCN, 2:1): $\lambda_{max}$ ($\varepsilon$) = 369 nm (20.4 × $10^3$ l·mol$^{-1}$·cm$^{-1}$); HRMS (ESI): *m/Z* calcd for $C_{51}H_{60}N_7O_{10}$ [M+H]$^+$: 930.4396, found: 930.4401.

*cyclo*-[(2*S*,4*E*,8*S*)-Hdn-L-Dap(2-(4′-((4′′-methoxyphenyl)diazenyl)phenoxy)acetyl)-D-*N*MeTrp-L-*β*Tyr]; (**S28**)

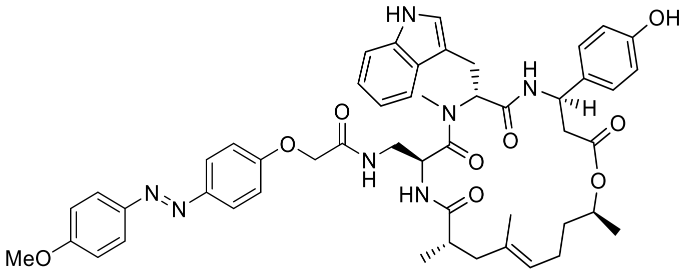

Standard Procedure **SP4** with cyclodepsipeptide **S54** (10.0 mg, 11.3 µmol, 1.0 equiv.), 2-(4′-((4′′-methoxyphenyl)diazenyl)phenoxy)acetic acid (3.9 mg, 11.3 µmol, 1.0 equiv.), HATU (10.3 mg, 27.0 µmol, 2.4 equiv.) and DIPEA (15.3 µl, 90.1 µmol, 8.0 equiv.) gave conjugate **S28** (6.5 mg, 64%) as a yellow solid after purification by silica-gel chromatography ($CH_2Cl_2$/MeOH, 98:2 → 90:10). $R_f$ = 0.36 ($CH_2Cl_2$/MeOH, 14:1); $[\alpha]_D^{24}$ = 46.7 (c = 0.09 in MeCN); $^1$H-NMR (500 MHz, DMSO-$d_6$): $\delta$ = 10.80 (d, $J$ = 1.2 Hz, 1 H), 9.28 (br s, 1 H), 8.58 (d, $J$ = 8.5 Hz, 1 H), 7.99 (t, $J$ = 6.0 Hz, 1 H), 7.85 (d, $J$ = 9.2 Hz, 2 H), 7.82 (d, $J$ = 8.9 Hz, 2 H), 7.68 (d, $J$ = 7.9 Hz, 1 H), 7.63 (d, $J$ = 8.5 Hz, 1 H), 7.30 (d, $J$ = 8.2 Hz, 1 H), 7.12 (d, $J$ = 9.2 Hz, 2 H), 7.09 - 7.00 (m, 6 H), 6.96 (t, $J$ = 7.5 Hz, 1 H), 6.67 (d, $J$ = 8.5 Hz, 2 H), 5.50 (dd, $J$ = 6.9, 9.3 Hz, 1 H), 5.18 (ddd, $J$ = 3.4, 8.5, 11.3 Hz, 1 H), 4.98 - 4.90 (m, 2 H), 4.69 (sxt, $J$ = 6.4 Hz, 1 H), 4.46 (d, $J$ = 2.7 Hz, 2 H), 3.86 (s, 3 H), 3.12 (s, 3 H), 3.10 - 2.96 (m, 3 H), 2.84 - 2.76 (m, 1 H), 2.70 - 2.57 (m, 3 H), 2.17 (dd, $J$ = 11.3, 14.3 Hz, 1 H), 1.93 - 1.80 (m, 2 H), 1.77 (d, $J$ = 14.0 Hz, 1 H), 1.50 (s, 3 H), 1.57 - 1.44 (m, 1 H), 1.43 - 1.33 (m, 1 H), 1.15 (d, $J$ = 6.4 Hz, 3 H), 0.94 (d, $J$ = 6.7 Hz, 3 H); $^{13}$C-NMR (126 MHz, DMSO-$d_6$): $\delta$ = 175.4, 170.7, 170.5, 170.0, 168.0, 162.0, 160.2, 156.7, 147.0, 146.6, 136.6, 133.5, 133.2, 127.6, 127.4, 124.7, 124.5, 123.8, 123.7, 121.3, 119.2, 118.5, 115.7, 115.5, 115.0, 111.6, 110.0, 71.2, 67.4, 56.1, 55.3, 49.3, 48.5, 43.2, 41.9, 40.7, 38.3, 35.4, 31.0, 25.8, 24.1, 20.0, 19.8, 17.2; IR: $\tilde{\nu}_{max}$ = 3317, 2931, 1728, 1667, 1597, 1501, 1454, 1246, 1150, 1026, 841, 745, 652; UV-VIS (PBS/MeCN, 2:1): $\lambda_{max}$ ($\varepsilon$) = 357 nm (23.0 × $10^3$ l·mol$^{-1}$·cm$^{-1}$); HRMS (ESI): *m/Z* calcd for $C_{50}H_{58}N_7O_9$ $[M+H]^+$: 900.4291, found: 900.4296.

*cyclo*-[(2*S*,4*E*,8*S*)-Hdn-L-Dap(2-(4´-((3´´,5´´-dimethoxyphenyl)diazenyl)phenoxy)-acetyl)-D-*N*MeTrp-L-*β*Tyr]; (**S29**)

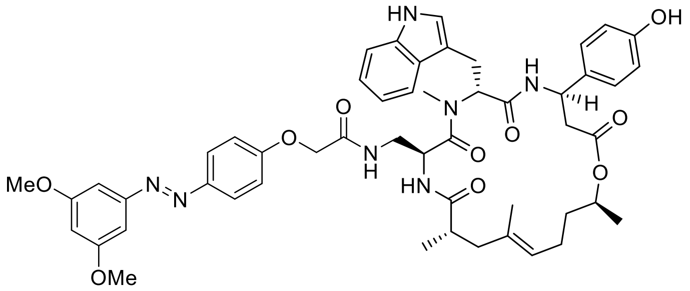

Standard Procedure **SP4** with cyclodepsipeptide **S54** (10.0 mg, 11.3 µmol, 1.0 equiv.), 2-(4´-((3´´,5´´-dimethoxyphenyl)diazenyl)phenoxy)acetic acid (3.6 mg, 11.3 µmol, 1.0 equiv.), HATU (10.3 mg, 27.0 µmol, 2.4 equiv.) and DIPEA (15.3 µl, 90.1 µmol, 8.0 equiv.) gave conjugate **S29** (4.7 mg, 43%) as a yellow solid after purification by silica-gel chromatography ($CH_2Cl_2$/acetone, 3:1 → 1:2). $R_f$ = 0.36 ($CH_2Cl_2$/MeOH, 14:1); $[\alpha]_D^{24}$ = 82.6 (c = 0.09 in MeCN); $^1$H-NMR (400 MHz, DMSO-$d_6$): $\delta$ = 10.81 (br s, 1 H), 9.39 (br s, 1 H), 8.58 (d, $J$ = 8.8 Hz, 1 H), 8.04 (t, $J$ = 5.6 Hz, 1 H), 7.87 (d, $J$ = 9.1 Hz, 2 H), 7.68 (d, $J$ = 7.3 Hz, 1 H), 7.67 (d, $J$ = 8.5 Hz, 1 H), 7.31 (d, $J$ = 8.2 Hz, 1 H), 7.13 - 7.00 (m, 8 H), 6.96 (t, $J$ = 7.3 Hz, 1 H), 6.72 - 6.63 (m, 3 H), 5.50 (dd, $J$ = 7.2, 8.9 Hz, 1 H), 5.18 (dt, $J$ = 3.4, 9.2 Hz, 1 H), 5.00 - 4.87 (m, 2 H), 4.69 (sxt, $J$ = 6.1 Hz, 1 H), 4.54 - 4.42 (m, 2 H), 3.84 (s, 6 H), 3.12 (s, 3 H), 3.10 - 2.96 (m, 2 H), 2.85 - 2.73 (m, 1 H), 2.72 - 2.56 (m, 4 H), 2.17 (dd, $J$ = 11.7, 14.0 Hz, 1 H), 1.86 (spt, $J$ = 7.0 Hz, 2 H), 1.77 (d, $J$ = 13.7 Hz, 1 H), 1.50 (s, 3 H), 1.58 - 1.45 (m, 1 H), 1.44 - 1.32 (m, 1 H), 1.15 (d, $J$ = 6.1 Hz, 3 H), 0.95 (d, $J$ = 6.7 Hz, 3 H); $^{13}$C-NMR (101 MHz, DMSO-$d_6$): $\delta$ = 175.5, 170.7, 170.5, 170.0, 167.9, 161.4, 160.9, 156.7, 154.3, 146.8, 136.6, 133.5, 133.2, 127.5, 127.4, 125.0, 123.8, 123.7, 121.3, 119.2, 118.5, 115.8, 115.5, 111.6, 110.0, 103.6, 100.8, 71.2, 67.4, 56.0, 55.3, 49.3, 48.5, 43.2, 41.9, 40.8, 38.3, 35.4, 31.0, 25.8, 24.1, 20.0, 19.8, 17.2; IR: $\tilde{\nu}_{max}$ = 3287, 2936, 1732, 1670, 1601, 1504, 1458, 1354, 1246, 1153, 1053, 1007, 837, 745, 652; UV-VIS (PBS/MeCN, 2:1): $\lambda_{max}$ ($\varepsilon$) = 347 nm (19.5 × $10^3$ l·mol$^{-1}$·cm$^{-1}$); HRMS (ESI): *m/Z* calcd for $C_{51}H_{60}N_7O_{10}$ $[M+H]^+$: 930.4396, found: 930.4395.

*cyclo*-[(2*S*,4*E*,8*S*)-Hdn-L-Dap(4-((2´-((3´´,4´´-dimethoxyphenyl)diazenyl)-5´-methoxyphenyl)amino)-4-oxobutyryl)-D-*N*MeTrp-L-*β*Tyr]; (**S30**)

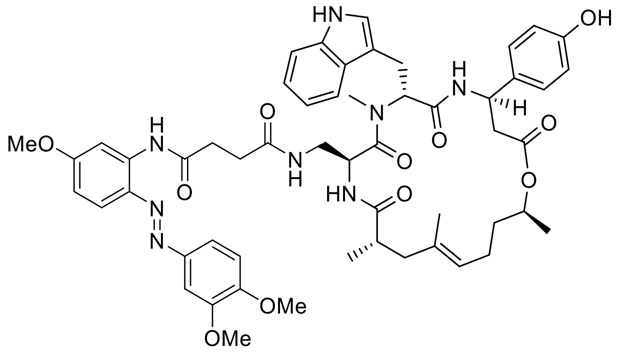

Standard Procedure **SP4** with cyclodepsipeptide **S54** (10.0 mg, 11.3 µmol, 1.0 equiv.), *N*-(2´-((3´´,4´´-Dimethoxyphenyl)diazenyl)-5´-methoxyphenyl)succinamic acid (4.4 mg, 11.3 µmol, 1.0 equiv.), HATU (10.3 mg, 27.0 µmol, 2.4 equiv.) and DIPEA (15.3 µl, 90.1 µmol, 8.0 equiv.) gave conjugate **S30** (9.0 mg, 80%) as a yellow solid after purification by silica-gel chromatography ($CH_2Cl_2$/acetone, 4:1 → 2:1 + 1% MeOH). $R_f$ = 0.35 ($CH_2Cl_2$/MeOH, 14:1); $[\alpha]_D^{24}$ = 18.4 (c = 0.10 in MeCN); $^1$H-NMR (500 MHz, DMSO-$d_6$): $\delta$ = 10.80 (s, 1 H), 10.30 (s, 1 H), 9.49 (br s, 1 H), 8.55 (d, *J* = 8.5 Hz, 1 H), 8.01 (d, *J* = 2.7 Hz, 1 H), 7.80 - 7.73 (m, 1 H), 7.74 (d, *J* = 9.2 Hz, 1 H), 7.69 - 7.60 (m, 4 H), 7.30 (d, *J* = 8.2 Hz, 1 H), 7.15 (d, *J* = 8.2 Hz, 1 H), 7.07 - 7.00 (m, 3 H), 6.98 (s, 1 H), 6.94 (t, *J* = 7.6 Hz, 1 H), 6.78 (dd, *J* = 2.6, 9.0 Hz, 1 H), 6.67 (d, *J* = 8.5 Hz, 2 H), 5.48 (t, *J* = 7.9 Hz, 1 H), 5.16 (ddd, *J* = 3.1, 8.3, 11.3 Hz, 1 H), 4.93 (t, *J* = 6.7 Hz, 1 H), 4.88 (dt, *J* = 4.3, 8.9 Hz, 1 H), 4.68 (sxt, *J* = 6.3 Hz, 1 H), 3.87 (s, 6 H), 3.83 (s, 3 H), 3.07 (s, 3 H), 2.99 (d, *J* = 8.2 Hz, 2 H), 2.77 - 2.55 (m, 5 H), 2.50 - 2.47 (m, 2 H), 2.39 - 2.29 (m, 2 H), 2.15 (dd, *J* = 11.7, 14.2 Hz, 1 H), 1.84 (qud, *J* = 7.1, 14.2 Hz, 2 H), 1.73 (d, *J* = 14.0 Hz, 1 H), 1.47 (s, 3 H), 1.54 - 1.45 (m, 1 H), 1.43 - 1.33 (m, 1 H), 1.14 (d, *J* = 6.4 Hz, 3 H), 0.94 (d, *J* = 6.7 Hz, 3 H); $^{13}$C-NMR (126 MHz, DMSO-$d_6$): $\delta$ = 175.3, 172.1, 171.6, 170.7, 170.6, 170.0, 162.5, 156.8, 152.0, 149.8, 146.9, 138.6, 136.6, 134.9, 133.6, 133.2, 127.5, 127.4, 123.8, 123.7, 121.3, 120.5, 119.2, 119.1, 118.5, 115.5, 111.6, 111.6, 110.3, 110.0, 106.0, 103.3, 71.2, 56.2, 56.0, 56.0, 55.2, 49.4, 48.6, 43.2, 41.9, 40.7, 38.3, 35.3, 32.4, 30.9, 30.5, 25.8, 24.0, 20.0, 19.8, 17.2; IR: $\tilde{\nu}_{max}$ = 3364, 2936, 1732, 1667, 1608, 1519, 1458, 1345, 1261, 1119, 1022, 821, 745, 652: UV-VIS (PBS/MeCN, 2:1): $\lambda_{max}$ ($\varepsilon$) = 392 nm (20.8 × $10^3$ l·mol$^{-1}$·cm$^{-1}$); HRMS (ESI): *m/Z* calcd for $C_{54}H_{65}N_8O_{11}$ [M+H]$^+$: 1001.477, found: 1001.477.

*cyclo*-[(2*S*,4*E*,8*S*)-Hdn-L-Dap(4-((5´-methoxy-2´-((4´´-methoxyphenyl)diazenyl)-phenyl)amino)-4-oxobutyryl)-D-*N*MeTrp-L-*β*Tyr]; (**S31**)

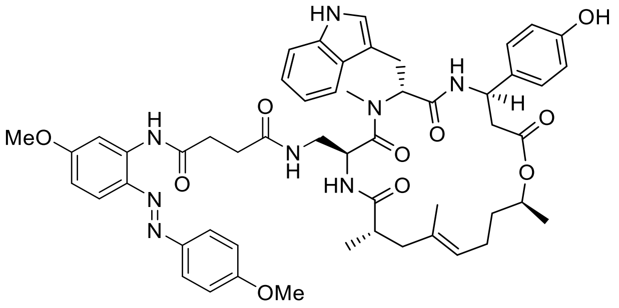

Standard Procedure **SP4** with cyclodepsipeptide **S54** (10.0 mg, 11.3 µmol, 1.0 equiv.), *N*-(5´-Methoxy-2´-((4´´-methoxyphenyl)diazenyl)phenyl)succinamic acid (4.0 mg, 11.3 µmol, 1.0 equiv.), HATU (10.3 mg, 27.0 µmol, 2.4 equiv.) and DIPEA (15.3 µl, 90.1 µmol, 8.0 equiv.) gave conjugate **S31** (7.5 mg, 68%) as a yellow solid after purification by silica-gel chromatography ($CH_2Cl_2$/acetone, 2:1 → 1:1). $R_f$ = 0.36 ($CH_2Cl_2$/MeOH, 14:1); $[\alpha]_D^{24}$ = 66.8 (c = 0.10 in MeCN); $^1$H-NMR (400 MHz, DMSO-$d_6$): $\delta$ = 10.79 (d, $J$ = 1.5 Hz, 1 H), 10.25 (s, 1 H), 9.42 (br s, 1 H), 8.54 (d, $J$ = 8.8 Hz, 1 H), 8.02 (d, $J$ = 2.9 Hz, 1 H), 7.99 (d, $J$ = 8.8 Hz, 2 H), 7.73 (d, $J$ = 9.1 Hz, 2 H), 7.67 (d, $J$ = 7.9 Hz, 1 H), 7.63 (d, $J$ = 8.5 Hz, 1 H), 7.30 (d, $J$ = 8.2 Hz, 1 H), 7.12 (d, $J$ = 9.1 Hz, 2 H), 7.07 - 6.98 (m, 4 H), 6.95 (t, $J$ = 7.3 Hz, 1 H), 6.77 (dd, $J$ = 2.9, 9.1 Hz, 1 H), 6.67 (d, $J$ = 8.5 Hz, 2 H), 5.49 (t, $J$ = 8.0 Hz, 1 H), 5.17 (ddd, $J$ = 3.2, 8.2, 11.0 Hz, 1 H), 4.98 - 4.86 (m, 2 H), 4.68 (sxt, $J$ = 6.3 Hz, 1 H), 3.86 (s, 3 H), 3.83 (s, 3 H), 3.09 (s, 3 H), 3.00 (d, $J$ = 7.9 Hz, 2 H), 2.80 - 2.54 (m, 7 H), 2.40 - 2.27 (m, 2 H), 2.16 (dd, $J$ = 11.5, 14.2 Hz, 1 H), 1.94 - 1.78 (m, 2 H), 1.74 (d, $J$ = 14.0 Hz, 1 H), 1.48 (s, 3 H), 1.55 - 1.44 (m, 1 H), 1.43 - 1.33 (m, 1 H), 1.14 (d, $J$ = 6.4 Hz, 3 H), 0.95 (d, $J$ = 6.7 Hz, 3 H); $^{13}$C-NMR (101 MHz, DMSO-$d_6$): $\delta$ = 175.3, 172.1, 171.6, 170.7, 170.7, 170.0, 162.5, 162.0, 156.7, 146.9, 138.6, 136.6, 134.9, 133.5, 133.2, 127.5, 127.4, 125.2, 123.8, 123.7, 121.3, 119.2, 119.2, 118.5, 115.5, 114.9, 111.6, 110.3, 110.0, 105.9, 71.2, 56.1, 56.0, 55.2, 49.3, 48.6, 43.2, 41.8, 40.7, 38.4, 35.3, 32.5, 31.0, 30.5, 25.7, 24.0, 20.0, 19.8, 17.2; IR: $\tilde{\nu}_{max}$ = 3318, 2932, 1734, 1667, 1597, 1520, 1458 1250, 1145, 1030, 837, 745, 652; UV-VIS (PBS/MeCN, 2:1): $\lambda_{max}$ ($\varepsilon$) = 382 nm (22.2 × $10^3$ l·mol$^{-1}$·cm$^{-1}$); HRMS (ESI): *m/Z* calcd for $C_{53}H_{63}N_8O_{10}$ $[M+H]^+$: 971.4662, found: 971.4666.

*cyclo*-[(2*S*,4*E*,8*S*)-Hdn-L-Dap(4-((4´-methoxy-2´-((3´´,4´´,5´´-trimethoxyphenyl)-diazenyl)phenyl)amino)-4-oxobutyryl)-D-*N*MeTrp-L-*β*Tyr]; (**S32**)

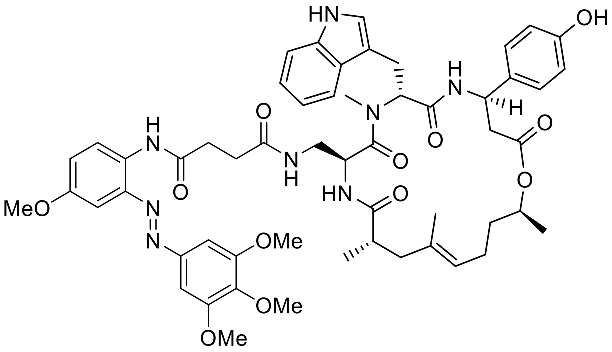

Standard Procedure **SP4** with cyclodepsipeptide **S54** (10.0 mg, 11.3 µmol, 1.0 equiv.), *N*-(4´-Methoxy-2´-((3´´,4´´,5´´-trimethoxyphenyl)diazenyl)phenyl)succinamic acid (4.7 mg, 11.3 µmol, 1.0 equiv.), HATU (10.3 mg, 27.0 µmol, 2.4 equiv.) and DIPEA (15.3 µl, 90.1 µmol, 8.0 equiv.) gave conjugate **S32** (7.9 mg, 68%) as an orange solid after purification by silica-gel chromatography ($CH_2Cl_2$/acetone, 4:1 → 2:1 + 1% MeOH). $R_f$ = 0.27 ($CH_2Cl_2$/MeOH, 14:1); $[\alpha]_D^{24}$ = 22.3 (c = 0.10 in MeCN); $^1$H-NMR (400 MHz, DMSO-$d_6$): $\delta$ = 10.79 (d, $J$ = 1.8 Hz, 1 H), 9.98 (s, 1 H), 9.40 (br s, 1 H), 8.55 (d, $J$ = 8.8 Hz, 1 H), 8.06 (d, $J$ = 9.1 Hz, 1 H), 7.76 - 7.69 (m, 1 H), 7.67 (d, $J$ = 7.9 Hz, 1 H), 7.62 (d, $J$ = 8.5 Hz, 1 H), 7.41 (s, 2 H), 7.29 (d, $J$ = 7.9 Hz, 1 H), 7.22 (d, $J$ = 2.9 Hz, 1 H), 7.12 (dd, $J$ = 2.9, 9.1 Hz, 1 H), 7.07 - 6.97 (m, 4 H), 6.94 (t, $J$ = 7.0 Hz, 1 H), 6.67 (d, $J$ = 8.5 Hz, 2 H), 5.48 (t, $J$ = 8.0 Hz, 1 H), 5.16 (ddd, $J$ = 3.6, 8.0, 11.4 Hz, 1 H), 4.92 (t, $J$ = 6.1 Hz, 1 H), 4.87 (dt, $J$ = 4.4, 9.1 Hz, 1 H), 4.67 (sxt, $J$ = 6.3 Hz, 1 H), 3.90 (s, 6 H), 3.80 (s, 3 H), 3.78 (s, 3 H), 3.07 (s, 3 H), 2.98 (d, $J$ = 7.9 Hz, 2 H), 2.79 - 2.53 (m, 7 H), 2.36 - 2.27 (m, 2 H), 2.14 (dd, $J$ = 11.5, 14.2 Hz, 1 H), 1.91 - 1.78 (m, 2 H), 1.73 (d, $J$ = 13.7 Hz, 1 H), 1.47 (s, 3 H), 1.54 - 1.44 (m, 1 H), 1.43 - 1.32 (m, 1 H), 1.14 (d, $J$ = 6.1 Hz, 3 H), 0.93 (d, $J$ = 6.7 Hz, 3 H); $^{13}$C-NMR (101 MHz, DMSO-$d_6$): $\delta$ = 175.3, 172.1, 171.1, 170.7, 170.6, 170.0, 156.7, 153.7, 148.5, 140.9, 136.6, 133.5, 133.2, 131.7, 127.5, 127.4, 123.8, 123.7, 121.3, 119.6, 119.2, 118.5, 115.5, 111.6, 110.0, 101.5, 99.0, 71.2, 60.7, 56.5, 55.9, 55.2, 49.3, 48.6, 43.2, 41.9, 40.8, 38.3, 35.3, 32.0, 30.9, 30.8, 25.7, 24.0, 20.0, 19.8, 17.2; IR: $\tilde{\nu}_{max}$ = 3294, 2931, 1724, 1654, 1512, 1415, 1307, 1223, 1126, 1002, 841, 745, 652; UV-VIS (PBS/MeCN, 2:1): $\lambda_{max}$ ($\varepsilon$) = 362 nm (18.2 × 10$^3$ l·mol$^{-1}$·cm$^{-1}$); HRMS (ESI): *m/Z* calcd for $C_{55}H_{67}N_8O_{12}$ [M+H]$^+$: 1031.487, found: 1031.488.

*cyclo*-[(2*S*,4*E*,8*S*)-Hdn-L-Dap(2-((4′-((3′′,4′′,5′′-trimethoxyphenyl)diazenyl)naphth-1-yl)-oxy)acetyl)-D-*N*MeTrp-L-*β*Tyr]; (**S33**)

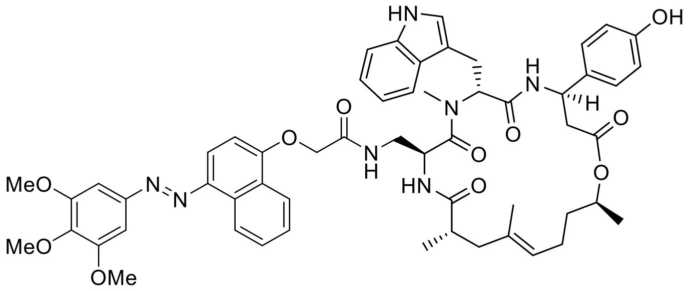

Standard Procedure **SP4** with cyclodepsipeptide **S54** (10.0 mg, 11.3 µmol, 1.0 equiv.), 2-((4′-((3′′,4′′,5′′-Trimethoxyphenyl)diazenyl)naphth-1′-yl)oxy)acetic acid (4.5 mg, 11.3 µmol, 1.0 equiv.), HATU (10.3 mg, 27.0 µmol, 2.4 equiv.) and DIPEA (15.3 µl, 90.1 µmol, 8.0 equiv.) gave conjugate **S33** (8.3 mg, 73%) as a yellow solid after purification by silica-gel chromatography ($CH_2Cl_2$/acetone, 4:1 → 1:2). $R_f$ = 0.35 ($CH_2Cl_2$/MeOH, 14:1); $[\alpha]_D^{24}$ = 33.7 (c = 0.10 in MeCN); $^1$H-NMR (500 MHz, DMSO-$d_6$): $\delta$ = 10.81 (d, *J* = 1.8 Hz, 1 H), 9.30 (s, 1 H), 8.92 (d, *J* = 8.5 Hz, 1 H), 8.57 (d, *J* = 8.5 Hz, 1 H), 8.45 (d, *J* = 8.2 Hz, 1 H), 8.07 (t, *J* = 6.0 Hz, 1 H), 7.83 (d, *J* = 8.5 Hz, 1 H), 7.77 (ddd, *J* = 1.2, 6.9, 8.3 Hz, 1 H), 7.70 (d, *J* = 8.9 Hz, 1 H), 7.68 (d, *J* = 8.2 Hz, 1 H), 7.65 (ddd, *J* = 1.2, 7.0, 8.2 Hz, 1 H), 7.36 (s, 2 H), 7.31 (d, *J* = 8.2 Hz, 1 H), 7.06 (d, *J* = 8.5 Hz, 2 H), 7.05 - 7.02 (m, 3 H), 6.93 (ddd, *J* = 0.9, 7.0, 7.9 Hz, 1 H), 6.67 (d, *J* = 8.5 Hz, 2 H), 5.52 (dd, *J* = 7.0, 9.2 Hz, 1 H), 5.18 (ddd, *J* = 3.3, 8.7, 10.8 Hz, 1 H), 5.00 (dd, *J* = 4.0, 8.9 Hz, 1 H), 4.96 (t, *J* = 6.1 Hz, 1 H), 4.74 - 4.65 (m, 3 H), 3.95 (s, 6 H), 3.78 (s, 3 H), 3.14 (s, 3 H), 3.09 - 2.99 (m, 2 H), 2.90 (td, *J* = 5.0, 13.0 Hz, 1 H), 2.71 - 2.58 (m, 3 H), 2.57 - 2.54 (m, 1 H), 2.18 (dd, *J* = 11.4, 14.2 Hz, 1 H), 1.86 (spt, *J* = 7.6 Hz, 2 H), 1.77 (d, *J* = 14.0 Hz, 1 H), 1.51 (s, 3 H), 1.57 - 1.47 (m, 1 H), 1.44 - 1.35 (m, 1 H), 1.15 (d, *J* = 6.4 Hz, 3 H), 0.93 (d, *J* = 6.7 Hz, 3 H); $^{13}$C-NMR (126 MHz, DMSO-$d_6$): $\delta$ = 175.5, 170.7, 170.5, 169.9, 167.8, 156.7, 156.5, 153.9, 149.0, 141.4, 140.5, 136.6, 133.5, 133.2, 132.2, 128.4, 127.6, 127.5, 126.6, 125.3, 123.9, 123.7, 123.2, 123.0, 121.3, 119.2, 118.5, 115.5, 113.4, 111.6, 110.0, 106.3, 100.8, 71.2, 67.8, 60.8, 56.5, 55.3, 49.3, 48.6, 43.2, 42.7, 41.8, 38.4, 35.4, 31.1, 25.8, 24.0, 20.0, 19.8, 17.2; IR: $\tilde{\nu}_{max}$ = 3352, 2936, 1728, 1670, 1516, 1466, 1331, 1234, 1126, 1002, 845, 768, 745, 656; UV-VIS (PBS/MeCN, 2:1): $\lambda_{max}$ ($\varepsilon$) = 401 nm (19.9 × $10^3$ l·mol$^{-1}$·cm$^{-1}$); HRMS (ESI): *m/Z* calcd for $C_{56}H_{64}N_7O_{11}$ $[M+H]^+$: 1010.466, found: 1010.464.

*cyclo*-[(2*S*,4*E*,8*S*)-Hdn-L-Dap(2-((5′-methoxy-4′-((3′′,4′′,5′′-trimethoxyphenyl)-diazenyl)naphth-1-yl)oxy)acetyl)-D-*N*MeTrp-L-*β*Tyr]; (**S34**)

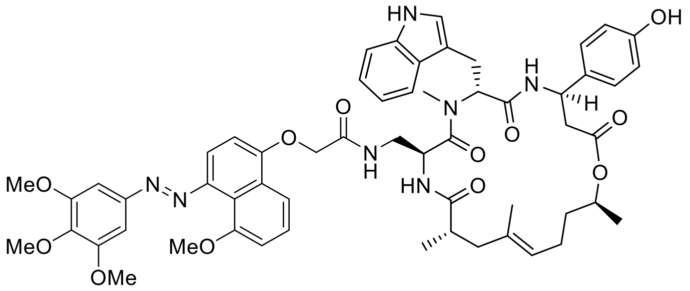

Standard Procedure **SP4** with cyclodepsipeptide **S54** (9.0 mg, 10.1 µmol, 1.0 equiv.), 2-((4′-((3′′,4′′,5′′-Trimethoxyphenyl)diazenyl)-5′-methoxynaphthalen-1′-yl)oxy)acetic acid (4.3 mg, 10.1 µmol, 1.0 equiv.), HATU (9.3 mg, 24.3 µmol, 2.4 equiv.) and DIPEA (13.8 µl, 81.1 µmol, 8.0 equiv.) gave conjugate **S34** (5.3 mg, 50%) as a yellow solid after purification by silica-gel chromatography ($CH_2Cl_2$/acetone, 4:1 → 1:2). $R_f$ = 0.40 ($CH_2Cl_2$/MeOH, 14:1); $[\alpha]_D^{24}$ = 47.7 (c = 0.10 in MeCN); $^1$H-NMR (400 MHz, DMSO-$d_6$): $\delta$ = 11.62 (s, 1 H), 9.28 (s, 1 H), 8.53 (d, $J$ = 8.8 Hz, 1 H), 8.02 (d, $J$ = 8.5 Hz, 1 H), 7.94 (t, $J$ = 6.0 Hz, 1 H), 7.69 (d, $J$ = 7.9 Hz, 1 H), 7.61 (d, $J$ = 8.5 Hz, 1 H), 7.51 (t, $J$ = 8.2 Hz, 1 H), 7.29 (s, 2 H), 7.26 (d, $J$ = 8.5 Hz, 1 H), 7.23 (d, $J$ = 8.2 Hz, 1 H), 7.20 (d, $J$ = 8.2 Hz, 1 H), 7.10 - 6.94 (m, 5 H), 6.65 (d, $J$ = 8.5 Hz, 2 H), 5.62 (dd, $J$ = 7.6, 9.1 Hz, 1 H), 5.17 (ddd, $J$ = 3.0, 9.1, 10.2 Hz, 1 H), 4.99 - 4.87 (m, 2 H), 4.72 - 4.57 (m, 3 H), 3.91 (s, 9 H), 3.77 (s, 3 H), 3.11 (s, 3 H), 3.06 - 2.95 (m, 2 H), 2.79 - 2.57 (m, 4 H), 2.44 - 2.35 (m, 1 H), 2.16 (dd, $J$ = 11.5, 13.9 Hz, 1 H), 1.92 - 1.80 (m, 2 H), 1.74 (d, $J$ = 12.6 Hz, 1 H), 1.48 (s, 3 H), 1.55 - 1.43 (m, 1 H), 1.42 - 1.33 (m, 1 H), 1.14 (d, $J$ = 6.1 Hz, 3 H), 0.91 (d, $J$ = 6.7 Hz, 3 H); $^{13}$C-NMR (101 MHz, DMSO-$d_6$) $\delta$ = 175.0, 170.2, 169.9, 169.0, 167.3, 156.3, 156.2, 154.5, 153.3, 148.5, 144.5, 139.5, 134.4, 133.0, 132.6, 127.1, 126.9, 126.7, 126.5, 124.4, 123.4, 122.1, 121.5, 120.9, 119.0, 118.8, 115.1, 115.0, 113.0, 109.1, 106.0, 105.8, 100.1, 70.6, 67.3, 60.2, 56.4, 55.9, 55.4, 48.7, 48.0, 42.7, 42.7, 41.0, 40.6, 37.9, 34.9, 30.8, 23.4, 19.5, 19.3, 16.7; IR: $\tilde{\nu}_{max}$ = 3350, 2912, 1727, 1670, 1515, 1464, 1234, 1126, 1006, 845, 768, 745, 656; UV-VIS (PBS/MeCN, 2:1): $\lambda_{max}$ ($\varepsilon$) = 390 nm (19.2 × $10^3$ l·mol$^{-1}$·cm$^{-1}$); HRMS (ESI): *m/Z* calcd for $C_{57}H_{66}N_7O_{11}$ $[M+H]^+$: 1040.476, found: 1040.477.

*cyclo*-[(2*S*,4*E*,8*S*)-Hdn-L-Lys(4-((2´-methoxy-5´-((3´´,4´´,5´´-trimethoxyphenyl)-diazenyl)phenyl)amino)-4-oxobutyryl)-D-*N*MeTrp-L-*β*Tyr]; (**S35**)

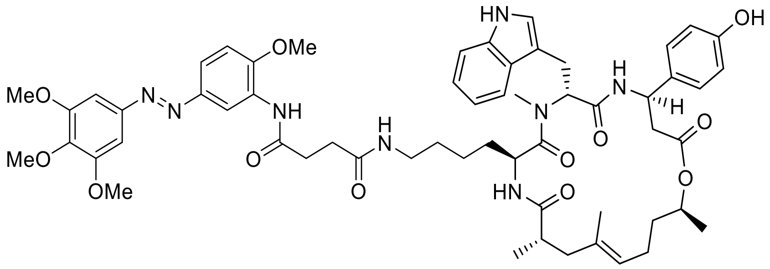

Standard Procedure **SP4** with cyclodepsipeptide **S51** (10.0 mg, 10.8 µmol, 1.0 equiv.), *N*-(2´-Methoxy-5´-((3´´,4´´,5´´-trimethoxyphenyl)diazenyl)phenyl)succinamic acid (4.5 mg, 10.8 µmol, 1.0 equiv.), HATU (9.9 mg, 25.9 µmol, 2.4 equiv.) and DIPEA (14.3 µl, 86.4 µmol, 8.0 equiv.) gave conjugate **S35** (7.0 mg, 60%) as an orange solid after purification by silica-gel chromatography ($CH_2Cl_2$/petroleum ether/acetone, 3:1:1 → 1:1:3). $R_f$ = 0.28 ($CH_2Cl_2$/MeOH, 14:1); $[\alpha]_D^{24}$ = 70.6 (c = 0.06 in MeCN); $^1$H-NMR (500 MHz, DMSO-$d_6$): $\delta$ = 10.81 (d, $J$ = 1.8 Hz, 1 H), 9.34 (s, 1 H), 9.32 (br s, 1 H), 8.65 (d, $J$ = 8.9 Hz, 1 H), 8.63 (d, $J$ = 1.8 Hz, 1 H), 7.81 (t, $J$ = 5.6 Hz, 1 H), 7.71 (dd, $J$ = 2.4, 8.5 Hz, 2 H), 7.66 (d, $J$ = 7.6 Hz, 1 H), 7.29 (d, $J$ = 7.9 Hz, 1 H), 7.24 (d, $J$ = 8.9 Hz, 1 H), 7.22 (s, 2 H), 7.13 (d, $J$ = 8.5 Hz, 2 H), 7.06 (d, $J$ = 2.4 Hz, 1 H), 7.01 (ddd, $J$ = 0.9, 6.7, 7.9 Hz, 1 H), 6.94 (ddd, $J$ = 0.9, 7.0, 7.9 Hz, 1 H), 6.70 (d, $J$ = 8.5 Hz, 2 H), 5.52 (dd, $J$ = 4.9, 11.6 Hz, 1 H), 5.19 (ddd, $J$ = 2.7, 9.2, 11.3 Hz, 1 H), 4.92 (t, $J$ = 6.4 Hz, 1 H), 4.67 (sxt, $J$ = 6.1 Hz, 1 H), 4.59 - 4.53 (m, 1 H), 3.94 (s, 3 H), 3.89 (s, 6 H), 3.75 (s, 3 H), 3.03 (s, 3 H), 3.07 - 2.98 (m, 1 H), 2.92 (dd, $J$ = 5.0, 15.1 Hz, 1 H), 2.89 - 2.78 (m, 2 H), 2.72 - 2.65 (m, 3 H), 2.59 (dd, $J$ = 2.7, 14.6 Hz, 1 H), 2.53 - 2.52 (m, 2 H), 2.41 (t, $J$ = 6.7 Hz, 2 H), 2.17 (dd, $J$ = 11.9, 14.6 Hz, 1 H), 1.91 - 1.77 (m, 2 H), 1.72 (d, $J$ = 14.6 Hz, 1 H), 1.48 (s, 3 H), 1.53 - 1.45 (m, 1 H), 1.42 - 1.33 (m, 1 H), 1.16 (d, $J$ = 6.1 Hz, 3 H), 1.14 - 1.04 (m, 2 H), 0.93 (d, $J$ = 6.7 Hz, 3 H), 0.86 - 0.79 (m, 2 H), 0.72 - 0.64 (m, 1 H); $^{13}$C-NMR (126 MHz, DMSO-$d_6$): $\delta$ = 174.9, 172.8, 171.6, 171.6, 170.8, 170.5, 156.7, 153.8, 152.2, 148.3, 145.9, 140.2, 136.6, 133.6, 133.5, 129.9, 127.6, 127.4, 123.8, 123.4, 121.9, 121.3, 119.1, 118.5, 115.5, 113.7, 111.6, 111.5, 110.0, 100.4, 71.4, 60.7, 56.6, 56.4, 55.4, 55.0, 49.5, 48.1, 43.0, 42.1, 38.7, 38.1, 35.3, 32.1, 31.0, 30.8, 29.1, 26.0, 24.2, 22.5, 20.2, 19.9, 17.4; IR: $\tilde{\nu}_{max}$ = 3313, 2928, 1732, 1651,1597,1519, 1415,1265,1126, 1006, 833, 744, 652; UV-VIS (PBS/MeCN, 2:1): $\lambda_{max}$ ($\varepsilon$) = 371 nm (19.9 × $10^3$ l·mol$^{-1}$·cm$^{-1}$); HRMS (ESI): $m/Z$ calcd for $C_{58}H_{73}N_8O_{12}$ $[M+H]^+$: 1073.534, found: 1073.534.

*cyclo*-[(2*S*,4*E*,8*S*)-Hdn-L-Orn(4-((2′-methoxy-5′-((3′′,4′′,5′′-trimethoxyphenyl)-diazenyl)phenyl)amino)-4-oxobutyryl)-D-*N*MeTrp-L-*β*Tyr]; (**S36**)

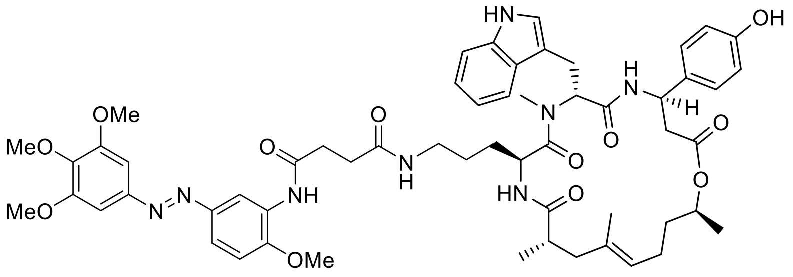

Standard Procedure **SP4** with cyclodepsipeptide **S52** (10.0 mg, 10.9 µmol, 1.0 equiv.), *N*-(2-Methoxy-5-((3′,4′,5′-trimethoxyphenyl)diazenyl)phenyl)succinamic acid (4.6 mg, 10.9 µmol, 1.0 equiv.), HATU (9.9 mg, 25.9 µmol, 2.4 equiv.) and DIPEA (14.3 µl, 86.4 µmol, 8.0 equiv.) gave conjugate **S36** (7.4 mg, 64%) as an orange solid after purification by silica-gel chromatography ($CH_2Cl_2$/petroleum ether/acetone, 3:1:1 → 1:1:3). $R_f$ = 0.33 ($CH_2Cl_2$/MeOH, 14:1); $[\alpha]_D^{24}$ = 76.6 (c = 0.06 in MeCN); $^1$H-NMR (500 MHz, DMSO-$d_6$): $\delta$ = 10.79 (d, $J$ = 1.5 Hz, 1 H), 9.34 (s, 1 H), 9.36 (br. s, 1 H), 8.64 - 8.59 (m, 2 H), 7.72 (d, $J$ = 7.9 Hz, 1 H), 7.70 (dd, $J$ = 1.8, 8.5 Hz, 1 H), 7.68 - 7.64 (m, 2 H), 7.30 (d, $J$ = 7.9 Hz, 1 H), 7.24 (d, $J$ = 8.9 Hz, 1 H), 7.21 (s, 2 H), 7.11 (d, $J$ = 8.5 Hz, 2 H), 7.06 - 7.01 (m, 2 H), 6.95 (ddd, $J$ = 0.9, 7.0, 7.9 Hz, 1 H), 6.69 (d, $J$ = 8.5 Hz, 2 H), 5.51 (dd, $J$ = 5.3, 10.8 Hz, 1 H), 5.18 (ddd, $J$ = 2.7, 8.9, 11.3 Hz, 1 H), 4.93 (t, $J$ = 6.6 Hz, 1 H), 4.68 (sxt, $J$ = 6.3 Hz, 1 H), 4.59 (dt, $J$ = 4.7, 8.6 Hz, 1 H), 3.95 (s, 3 H), 3.89 (s, 6 H), 3.75 (s, 3 H), 3.03 (s, 3 H), 3.06 - 2.98 (m, 1 H), 2.93 (dd, $J$ = 5.2, 14.6 Hz, 1 H), 2.79 (qd, $J$ = 6.6, 13.2 Hz, 1 H), 2.74 - 2.69 (m, 1 H), 2.68 - 2.63 (m, 3 H), 2.59 (dd, $J$ = 2.7, 14.6 Hz, 1 H), 2.53 - 2.52 (m, 2 H), 2.37 (t, $J$ = 6.9 Hz, 2 H), 2.16 (dd, $J$ = 11.7, 14.5 Hz, 1 H), 1.91 - 1.77 (m, 2 H), 1.73 (d, $J$ = 14.0 Hz, 1 H), 1.48 (s, 3 H), 1.54 - 1.45 (m, 1 H), 1.42 - 1.33 (m, 1 H), 1.15 (d, $J$ = 6.1 Hz, 3 H), 1.00 - 0.96 (m, 1 H), 0.93 (d, $J$ = 6.7 Hz, 3 H), 0.88 - 0.78 (m, 2 H); $^{13}$C-NMR (126 MHz, DMSO-$d_6$): $\delta$ = 175.0, 172.6, 171.6, 171.5, 170.8, 170.4, 156.7, 153.8, 152.3, 148.3, 145.9, 140.2, 136.6, 133.6, 133.4, 128.7, 127.5, 127.4, 123.8, 123.6, 121.9, 121.3, 119.1, 118.5, 115.5, 113.8, 111.6, 111.5, 110.0, 100.4, 71.3, 60.7, 56.6, 56.4, 55.1, 49.4, 48.0, 43.1, 42.1, 38.7, 38.1, 35.3, 32.0, 31.0, 30.7, 29.1, 25.9, 25.4, 24.1, 20.1, 20.0, 17.3; IR: $\tilde{\nu}_{max}$ = 3317, 2935, 1651, 1597, 1519, 1415, 1265, 1223, 1126, 1003, 833, 745, 652; UV-VIS (PBS/MeCN, 2:1): $\lambda_{max}$ ($\varepsilon$) = 371 nm (19.2 × 10$^3$ l·mol$^{-1}$·cm$^{-1}$); HRMS (ESI): *m/Z* calcd for $C_{57}H_{71}N_8O_{12}$ [M+H]$^+$: 1059.519, found: 1059.519.

*cyclo*-[(2*S*,4*E*,8*S*)-Hdn-L-Dap(2-(4′-((4′′-(2′′′-hydroxyethoxy)-3′′,5′′-dimethoxyphenyl)diazenyl)phenoxy)acetyl)-D-*N*MeTrp-L-*β*Tyr]; (**S37**)

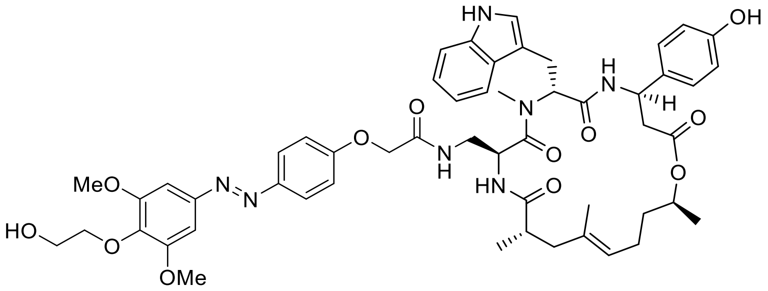

Standard Procedure **SP4** with cyclodepsipeptide **S54** (10.0 mg, 11.3 µmol, 1.0 equiv.), 2-(4′-((4′′-(2′′′-hydroxyethoxy)-3′′,5′′-dimethoxyphenyl)diazenyl)phenoxy)acetic acid (4.2 mg, 11.3 µmol, 1.0 equiv.), HATU (10.3 mg, 27.0 µmol, 2.4 equiv.) and DIPEA (15.3 µl, 90.1 µmol, 8.0 equiv.) gave conjugate **S37** (8.8 mg, 79%) as a yellow solid after purification by silica-gel chromatography ($CH_2Cl_2$/MeOH, 98:2 → 90:10). $R_f$ = 0.32 ($CH_2Cl_2$/MeOH, 14:1); $[\alpha]_D^{24}$ = 34.6 (c = 0.10 in MeCN); $^1$H-NMR (500 MHz, DMSO-$d_6$): $\delta$ = 10.80 (d, $J$ = 1.8 Hz, 1 H), 9.31 (br s, 1 H), 8.58 (d, $J$ = 8.5 Hz, 1 H), 8.00 (t, $J$ = 6.0 Hz, 1 H), 7.86 (d, $J$ = 8.9 Hz, 2 H), 7.68 (d, $J$ = 7.9 Hz, 1 H), 7.65 (d, $J$ = 8.5 Hz, 1 H), 7.31 (d, $J$ = 8.2 Hz, 1 H), 7.22 (s, 2 H), 7.10 - 7.00 (m, 6 H), 6.96 (t, $J$ = 7.5 Hz, 1 H), 6.67 (d, $J$ = 8.5 Hz, 2 H), 5.50 (dd, $J$ = 6.9, 9.3 Hz, 1 H), 5.18 (ddd, $J$ = 3.4, 8.9, 11.3 Hz, 1 H), 5.00 - 4.91 (m, 2 H), 4.69 (sxt, $J$ = 6.3 Hz, 1 H), 4.65 - 4.60 (m, 1 H), 4.52 - 4.42 (m, 2 H), 3.97 (t, $J$ = 5.6 Hz, 2 H), 3.89 (s, 6 H), 3.65 (q, $J$ = 5.7 Hz, 2 H), 3.12 (s, 3 H), 3.07 - 2.97 (m, 2 H), 2.86 - 2.74 (m, 2 H), 2.70 - 2.56 (m, 4 H), 2.18 (dd, $J$ = 11.6, 14.3 Hz, 1 H), 1.93 - 1.80 (m, 2 H), 1.77 (d, $J$ = 14.0 Hz, 1 H), 1.50 (s, 3 H), 1.55 - 1.46 (m, 1 H), 1.15 (d, $J$ = 6.1 Hz, 3 H), 0.95 (d, $J$ = 6.7 Hz, 3 H); $^{13}$C-NMR (126 MHz, DMSO-$d_6$): $\delta$ = 175.5, 170.7, 170.5, 170.0, 168.0, 160.6, 156.7, 153.8, 148.2, 146.9, 139.6, 136.6, 133.5, 133.2, 127.6, 127.4, 124.8, 123.8, 123.7, 121.3, 119.2, 118.5, 115.8, 115.5, 111.6, 110.0, 100.5, 74.8, 71.2, 67.4, 60.7, 56.5, 55.3, 49.3, 48.5, 43.2, 41.9, 40.7, 38.3, 35.4, 31.0, 25.8, 24.1, 20.0, 19.8, 17.2; IR: $\tilde{\nu}_{max}$ = 3318, 2036, 1732, 1670, 1597, 1497, 1458, 1327, 1223, 1126, 841, 745, 656; UV-VIS (PBS/MeCN, 2:1): $\lambda_{max}$ ($\varepsilon$) = 362 nm (24.6 × 10$^3$ l·mol$^{-1}$·cm$^{-1}$); HRMS (ESI): *m/Z* calcd for $C_{53}H_{64}N_7O_{12}$ [M+H]$^+$: 990.461, found: 990.461.

*cyclo*-[(2*S*,4*E*,8*S*)-Hdn-L-Orn(2-(4′-((4′′-(2′′′-hydroxyethoxy)-3′′,5′′-dimethoxyphenyl)diazenyl)phenoxy)acetyl)-D-*N*MeTrp-L-*β*Tyr]; (**S38**)

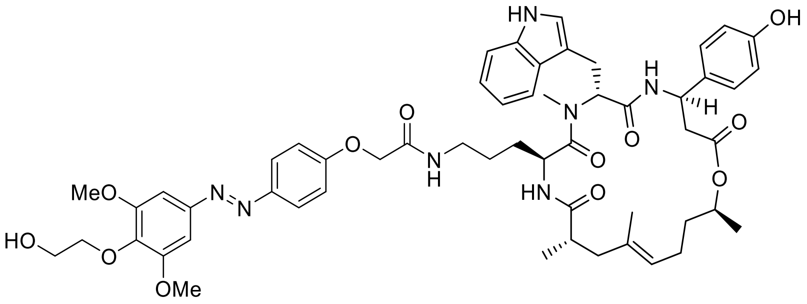

Standard Procedure **SP4** with cyclodepsipeptide **S52** (10.0 mg, 10.9 µmol, 1.0 equiv.), 2-(4′-((4′′-(2′′′-hydroxyethoxy)-3′′,5′′-dimethoxyphenyl)diazenyl)phenoxy)acetic acid (4.1 mg, 10.9 µmol, 1.0 equiv.), HATU (10.0 mg, 26.2 µmol, 2.4 equiv.) and DIPEA (14.9 µl, 87.3 µmol, 8.0 equiv.) gave conjugate **S38** (7.9 mg, 71%) as a yellow solid after purification by silica-gel chromatography ($CH_2Cl_2$/MeOH, 98:2 → 90:10). $R_f$ = 0.31 ($CH_2Cl_2$/MeOH, 14:1); $[\alpha]_D^{24}$ = 40.1 (c = 0.10 in MeCN); $^1$H-NMR (500 MHz, DMSO-$d_6$): $\delta$ = 10.79 (d, $J$ = 1.8 Hz, 1 H), 9.32 (s, 1 H), 8.61 (d, $J$ = 8.8 Hz, 1 H), 7.95 (t, $J$ = 5.6 Hz, 1 H), 7.88 (d, $J$ = 9.2 Hz, 2 H), 7.74 (d, $J$ = 8.5 Hz, 1 H), 7.66 (d, $J$ = 7.9 Hz, 1 H), 7.31 (d, $J$ = 7.9 Hz, 1 H), 7.22 (s, 2 H), 7.12 (d, $J$ = 9.2 Hz, 2 H), 7.10 (d, $J$ = 8.5 Hz, 2 H), 7.06 - 7.02 (m, 2 H), 6.96 (ddd, $J$ = 0.9, 7.0, 7.9 Hz, 1 H), 6.69 (d, $J$ = 8.5 Hz, 2 H), 5.51 (dd, $J$ = 5.5, 11.0 Hz, 1 H), 5.18 (ddd, $J$ = 2.7, 9.2, 11.3 Hz, 1 H), 4.93 (t, $J$ = 6.7 Hz, 1 H), 4.68 (sxt, $J$ = 6.4 Hz, 1 H), 4.64 - 4.57 (m, 2 H), 4.55 (s, 2 H), 3.97 (t, $J$ = 5.6 Hz, 2 H), 3.88 (s, 6 H), 3.65 (q, $J$ = 5.5 Hz, 2 H), 3.04 (s, 3 H), 3.06 - 2.99 (m, 1 H), 2.95 (dd, $J$ = 5.2, 15.0 Hz, 1 H), 2.91 - 2.78 (m, 2 H), 2.68 (dd, $J$ = 11.3, 15.0 Hz, 1 H), 2.59 (dd, $J$ = 3.1, 15.0 Hz, 1 H), 2.54 - 2.52 (m, 2 H), 2.17 (dd, $J$ = 11.7, 14.2 Hz, 1 H), 1.91 - 1.78 (m, 2 H), 1.74 (d, $J$ = 15.0 Hz, 1 H), 1.49 (s, 3 H), 1.54 - 1.45 (m, 1 H), 1.42 - 1.33 (m, 1 H), 1.15 (d, $J$ = 6.4 Hz, 3 H), 1.04 - 1.01 (m, 1 H), 0.93 (d, $J$ = 6.7 Hz, 3 H), 0.91 - 0.79 (m, 2 H); $^{13}$C-NMR (126 MHz, DMSO-$d_6$): $\delta$ = 175.0, 172.5, 170.8, 170.4, 167.4, 160.7, 156.7, 153.8, 148.2, 146.9, 139.6, 136.6, 133.6, 133.4, 127.5, 127.5, 124.8, 123.8, 123.6, 121.3, 119.1, 118.6, 115.8, 115.5, 111.6, 110.0, 100.5, 74.8, 71.3, 67.6, 60.7, 56.5, 55.1, 49.4, 48.0, 43.1, 42.0, 38.5, 38.2, 35.3, 31.0, 29.1, 26.0, 25.3, 24.1, 20.1, 20.0, 17.3; IR: $\tilde{\nu}_{max}$ = 3348, 2943, 1732, 1636, 1501, 1454, 1373, 1219, 1126, 1042, 918, 841, 748, 652; UV-VIS (PBS/MeCN, 2:1): $\lambda_{max}$ ($\varepsilon$) = 362 nm (24.6 × $10^3$ l·mol$^{-1}$·cm$^{-1}$); HRMS (ESI): *m/Z* calcd for $C_{55}H_{68}N_7O_{12}$ $[M+H]^+$: 1018.492, found: 1018.493.

*cyclo*-[(2*S*,4*E*,8*S*)-Hdn-L-Orn(2-(3´-hydroxy-4´-((3´´,4´´,5´´-trimethoxyphenyl)-diazenyl)phenoxy)acetyl)-D-*N*MeTrp-L-*β*Tyr]; (**S39**)

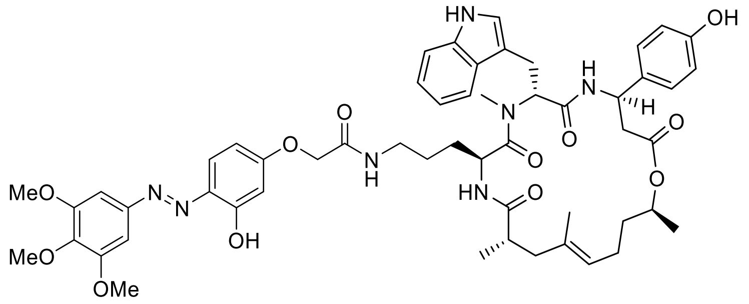

Standard Procedure **SP4** with cyclodepsipeptide **S52** (10.0 mg, 10.9 µmol, 1.0 equiv.), 2-(3´-Hydroxy-4´-((3´´,4´´,5´´-trimethoxyphenyl)diazenyl)phenoxy)acetic acid (4.0 mg, 10.9 µmol, 1.0 equiv.), HATU (10.0 mg, 26.2 µmol, 2.4 equiv.) and DIPEA (14.9 µl, 87.3 µmol, 8.0 equiv.) gave conjugate **S39** (4.5 mg, 41%) as an orange solid after purification by silica-gel chromatography ($CH_2Cl_2$/acetone, 3:1 → 1:2). $R_f$ = 0.40 ($CH_2Cl_2$/MeOH, 14:1); $[\alpha]_D^{24}$ = –9.4 (c = 0.10 in MeCN); $^1$H-NMR (400 MHz, DMSO-*d*$_6$): $\delta$ = 11.74 (br s, 1 H), 10.80 (br s, 1 H), 9.33 (br s, 1 H), 8.63 (d, *J* = 8.5 Hz, 1 H), 7.93 (t, *J* = 5.1 Hz, 1 H), 7.75 (d, *J* = 9.1 Hz, 2 H), 7.66 (d, *J* = 7.9 Hz, 1 H), 7.31 (s, 2 H), 7.36 - 7.26 (m, 1 H), 7.10 (d, *J* = 8.5 Hz, 2 H), 7.06 - 7.01 (m, 2 H), 6.95 (t, *J* = 7.6 Hz, 1 H), 6.69 (d, *J* = 8.5 Hz, 2 H), 6.65 - 6.60 (m, 1 H), 6.56 (br s, 1 H), 5.51 (dd, *J* = 5.6, 10.5 Hz, 1 H), 5.18 (ddd, *J* = 2.8, 8.0, 11.0 Hz, 1 H), 4.93 (t, *J* = 5.8 Hz, 1 H), 4.68 (sxt, *J* = 6.1 Hz, 1 H), 4.65 - 4.56 (m, 1 H), 4.52 (s, 2 H), 3.88 (s, 6 H), 3.75 (s, 3 H), 3.04 (s, 3 H), 3.02 - 2.92 (m, 2 H), 2.92 - 2.77 (m, 2 H), 2.74 - 2.55 (m, 3 H), 2.17 (dd, *J* = 11.7, 14.0 Hz, 1 H), 1.84 (spt, *J* = 6.7 Hz, 2 H), 1.74 (d, *J* = 14.0 Hz, 1 H), 1.48 (s, 3 H), 1.55 - 1.45 (m, 1 H), 1.44 - 1.32 (m, 1 H), 1.15 (d, *J* = 6.4 Hz, 3 H), 1.07 - 0.99 (m, 2 H), 0.93 (d, *J* = 6.4 Hz, 3 H), 0.89 - 0.80 (m, 2 H); $^{13}$C-NMR (101 MHz, DMSO-*d*$_6$): $\delta$ = 175.0, 172.6, 170.8, 170.4, 167.3, 162.3, 156.7, 153.8, 147.4, 140.0, 136.6, 136.6, 133.6, 133.4, 127.5, 127.5, 123.8, 123.6, 121.3, 119.1, 118.5, 115.7, 115.6, 115.5, 111.7, 111.6, 110.0, 103.1, 100.3, 71.3, 67.5, 60.7, 56.5, 55.2, 49.4, 48.0, 43.1, 42.1, 38.5, 38.2, 35.3, 31.0, 29.1, 25.9, 25.3, 24.1, 20.1, 20.0, 17.3; IR: $\tilde{\nu}_{max}$ = 3588, 3290, 3005, 2943, 1720, 1628, 1501, 1442, 1419, 1377, 1126, 1037, 918, 748, 601; UV-VIS (PBS/MeCN, 2:1): $\lambda_{max}$ ($\varepsilon$) = 388 nm (21.4 × 10$^3$ l·mol$^{-1}$·cm$^{-1}$); HRMS (ESI): *m/Z* calcd for $C_{54}H_{66}N_7O_{12}$ $[M+H]^+$: 1004.476, found: 1004.478.

*cyclo*-[(2*S*,4*E*,8*S*)-Hdn-L-Orn(2-(2´-hydroxy-4´-((3´´,4´´,5´´-trimethoxyphenyl)-diazenyl)phenoxy)acetyl)-D-*N*MeTrp-L-*β*Tyr]; (**S40**)

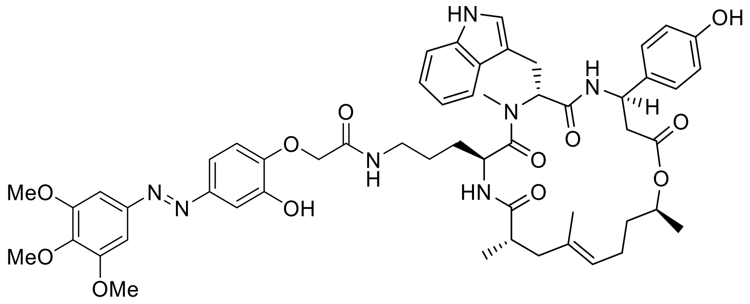

Standard Procedure **SP4** with cyclodepsipeptide **S52** (10.0 mg, 10.9 µmol, 1.0 equiv.), 2-(2´-Hydroxy-4´-((3´´,4´´,5´´-trimethoxyphenyl)diazenyl)phenoxy)acetic acid (4.0 mg, 10.9 µmol, 1.0 equiv.), HATU (10.0 mg, 26.2 µmol, 2.4 equiv.) and DIPEA (14.9 µl, 87.3 µmol, 8.0 equiv.) gave conjugate **S40** (6.8 mg, 62%) as an orange solid after purification by silica-gel chromatography ($CH_2Cl_2$/acetone, 2:1 → 1:2 + 1% MeOH). $R_f$ = 0.36 ($CH_2Cl_2$/MeOH, 14:1); $[\alpha]_D^{24}$ = 52.8 (c = 0.05 in MeCN); $^1$H-NMR (400 MHz, DMSO-$d_6$): $\delta$ = 10.92 (br s, 1 H), 9.34 (br s, 1 H), 9.54 (br s, 1 H), 8.62 (d, $J$ = 8.8 Hz, 1 H), 8.31 (br s, 1 H), 7.78 (d, $J$ = 8.5 Hz, 1 H), 7.65 (d, $J$ = 7.6 Hz, 1 H), 7.40 (d, $J$ = 8.5 Hz, 1 H), 7.33 (s, 1 H), 7.32 (d, $J$ = 5.8 Hz, 1 H), 7.20 (s, 2 H), 7.14 - 7.07 (m, 3 H), 7.07 - 7.01 (m, 2 H), 6.95 (t, $J$ = 7.3 Hz, 1 H), 6.69 (d, $J$ = 8.5 Hz, 2 H), 5.52 (dd, $J$ = 5.4, 10.7 Hz, 1 H), 5.18 (ddd, $J$ = 2.8, 8.2, 11.4 Hz, 1 H), 4.94 (t, $J$ = 6.4 Hz, 1 H), 4.68 (sxt, $J$ = 6.1 Hz, 1 H), 4.62 (dt, $J$ = 5.1, 8.4 Hz, 1 H), 4.53 (s, 2 H), 3.88 (s, 6 H), 3.75 (s, 3 H), 3.04 (s, 3 H), 3.08 - 2.95 (m, 2 H), 2.94 - 2.84 (m, 2 H), 2.74 - 2.60 (m, 2 H), 2.59 (dd, $J$ = 2.8, 14.8 Hz, 1 H), 2.18 (dd, $J$ = 11.7, 14.3 Hz, 1 H), 1.91 - 1.79 (m, 2 H), 1.75 (d, $J$ = 14.0 Hz, 1 H), 1.49 (s, 3 H), 1.56 - 1.45 (m, 1 H), 1.43 - 1.32 (m, 1 H), 1.15 (d, $J$ = 6.4 Hz, 3 H), 1.11 - 1.01 (m, 2 H), 0.94 (d, $J$ = 6.7 Hz, 3 H), 0.97 - 0.81 (m, 2 H); $^{13}$C-NMR (101 MHz, DMSO-$d_6$): $\delta$ = 174.6, 172.1, 170.3, 169.9, 167.1, 156.2, 153.3, 147.8, 139.8, 136.2, 136.1, 133.2, 133.0, 127.1, 127.0, 123.4, 123.1, 120.8, 118.6, 118.1, 115.1, 115.1, 111.2, 109.5, 106.7, 99.9, 70.8, 60.2, 56.0, 54.7, 49.0, 47.6, 42.6, 41.6, 37.9, 37.7, 34.9, 30.6, 28.6, 25.5, 24.9, 23.6, 19.6, 19.5, 16.9; IR: $\tilde{\nu}_{max}$ = 3583, 3310, 2932, 1734, 1659, 1454, 1377, 1273, 1126, 1045, 1003, 918, 745, 656; UV-VIS (PBS/MeCN, 2:1): $\lambda_{max}$ ($\varepsilon$) = 371 nm (19.3 × $10^3$ l·mol$^{-1}$·cm$^{-1}$); HRMS (ESI): *m/Z* calcd for $C_{54}H_{66}N_7O_{12}$ $[M+H]^+$: 1004.476, found: 1004.476.

*cyclo*-[(2*S*,4*E*,8*S*)-Hdn-L-Dap(4-((2′-((3′′,5′′-dimethoxy-4′′-(2′′′-(4′′′′-methyl-piperazin-1′′′′-yl)ethoxy)phenyl)diazenyl)-5-methoxyphenyl)amino)-4-oxobutyryl)-D-*N*MeTrp-L-$\beta$Tyr]; (**S41**)

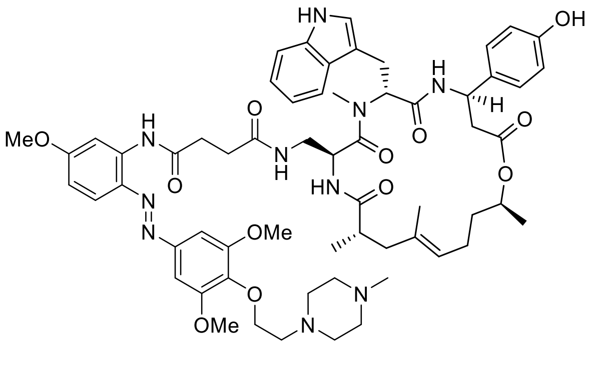

Standard Procedure **SP4** with cyclodepsipeptide **S54** (10.0 mg, 11.3 µmol, 1.0 equiv.), *N*-(2′-((3′′,5′′-Dimethoxy-4′′-(2′′′-(4′′′′-methylpiperazin-1′′′′-yl)ethoxy)phenyl)diazenyl)-5′-methoxyphenyl)succinamic acid (6.0 mg, 11.3 µmol, 1.0 equiv.), HATU (10.3 mg, 27.0 µmol, 2.4 equiv.) and DIPEA (15.3 µl, 90.1 µmol, 8.0 equiv.) gave conjugate **S41** (4.8 mg, 37%) as an orange solid after purification by silica-gel chromatography ($CH_2Cl_2$/MeOH, 95:5 → 85:15). $R_f$ = 0.63 ($CH_2Cl_2$/MeOH, 4:1); $[\alpha]_D^{24}$ = 14.4 (c = 0.11 in MeCN); $^1$H-NMR (500 MHz, DMSO-$d_6$): $\delta$ = 10.78 (d, *J* = 1.7 Hz, 1 H), 10.26 (s, 1 H), 9.39 (br s, 1 H), 8.54 (d, *J* = 8.9 Hz, 1 H), 7.99 (d, *J* = 2.4 Hz, 1 H), 7.75 (d, *J* = 9.2 Hz, 1 H), 7.73 (t, *J* = 5.8 Hz, 1 H), 7.66 (d, *J* = 7.9 Hz, 1 H), 7.61 (d, *J* = 8.9 Hz, 1 H), 7.35 (s, 2 H), 7.29 (d, *J* = 7.9 Hz, 1 H), 7.05 (d, *J* = 8.5 Hz, 2 H), 7.04 - 7.00 (m, 1 H), 6.98 (d, *J* = 2.1 Hz, 1 H), 6.94 (t, *J* = 7.6 Hz, 1 H), 6.79 (dd, *J* = 2.7, 9.2 Hz, 1 H), 6.67 (d, *J* = 8.5 Hz, 2 H), 5.47 (t, *J* = 8.1 Hz, 1 H), 5.16 (ddd, *J* = 3.3, 8.4, 11.0 Hz, 1 H), 4.92 (t, *J* = 6.9 Hz, 1 H), 4.87 (dt, *J* = 4.1, 8.9 Hz, 1 H), 4.67 (sxt, *J* = 6.3 Hz, 1 H), 4.03 (t, *J* = 6.0 Hz, 2 H), 3.88 (s, 6 H), 3.84 (s, 3 H), 3.06 (s, 3 H), 2.98 (d, *J* = 8.2 Hz, 2 H), 2.77 - 2.55 (m, 9 H), 2.44 (br. s, 4 H), 2.35 - 2.28 (m, 1 H), 2.31 (br. s, 4 H), 2.13 (s, 3 H), 2.17 - 2.09 (m, 1 H), 1.84 (qud, *J* = 7.0, 13.4 Hz, 2 H), 1.76 - 1.69 (m, 2 H), 1.47 (s, 3 H), 1.53 - 1.45 (m, 1 H), 1.41 - 1.32 (m, 1 H), 1.14 (d, *J* = 6.1 Hz, 3 H), 0.93 (d, *J* = 6.7 Hz, 3 H); $^{13}$C-NMR (126 MHz, DMSO-$d_6$): $\delta$ = 175.3, 172.0, 171.6, 170.7, 170.6, 170.0, 163.0, 156.7, 153.8, 148.5, 139.4, 139.1, 136.6, 134.9, 133.5, 133.2, 127.5, 127.4, 123.8, 123.7, 121.3, 119.2, 118.8, 118.5, 115.5, 111.6, 110.5, 110.0, 106.2, 101.1, 71.2, 71.0, 57.8, 56.6, 56.0, 55.2, 55.3, 53.4, 49.3, 48.6, 46.3, 43.2, 41.9, 40.7, 38.3, 35.3, 32.4, 30.9, 30.6, 25.8, 24.0, 20.0, 19.8, 17.2; IR: $\tilde{\nu}_{max}$ = 3260, 2936, 1732, 1670, 1597, 1519, 1458, 1284, 1223, 1126, 1026, 1007, 837, 745, 652; UV-VIS (PBS/MeCN, 2:1): $\lambda_{max}$ ($\varepsilon$) = 386 nm (18.6 × $10^3$ l·mol$^{-1}$·cm$^{-1}$); HRMS (ESI): *m/Z* calcd for $C_{61}H_{79}N_{10}O_{12}$ [M+H]$^+$: 1143.587, found: 1143.586.

*cyclo*-[(2*S*,4*E*,8*S*)-Hdn-L-Dap(2-(4´-((3´´,5´´-dimethoxy-4´´-(2´´´-(4´´´´-methylpiperazin-1´´´´-yl)ethoxy)phenyl)diazenyl)phenoxy)acetyl)-D-*N*MeTrp-L-*β*Tyr]; (**S42**)

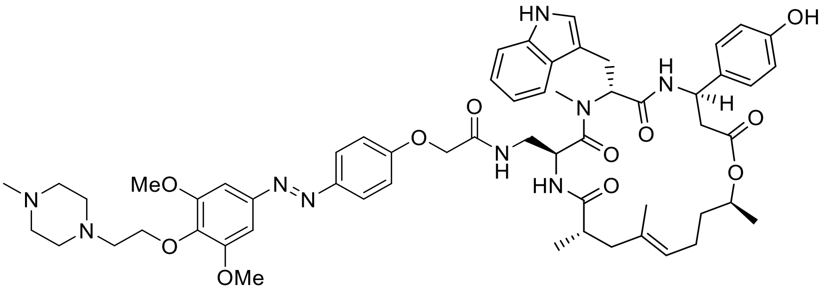

Standard Procedure **SP4** with cyclodepsipeptide **S54** (10.0 mg, 11.3 µmol, 1.0 equiv.), 2-(4´-((3´´,5´´-dimethoxy-4´´-(2´´´-(4´´´´-methylpiperazin-1´´´´-yl)ethoxy)phenyl)diazenyl)-phenoxy)acetic acid (5.2 mg, 11.3 µmol, 1.0 equiv.), HATU (10.3 mg, 27.0 µmol, 2.4 equiv.) and DIPEA (15.3 µl, 90.1 µmol, 8.0 equiv.) gave conjugate **S42** (8.1 mg, 67%) as a yellow solid after purification by silica-gel chromatography ($CH_2Cl_2$/MeOH, 95:5 → 85:15). $R_f$ = 0.63 ($CH_2Cl_2$/MeOH, 4:1); $[\alpha]_D^{24}$ = 27.3 (c = 0.11 in MeCN); $^1$H-NMR (500 MHz, DMSO-$d_6$): $\delta$ = 10.83 (s, 1 H), 8.58 (d, $J$ = 8.5 Hz, 1 H), 8.11 - 8.05 (m, 1 H), 7.86 (d, $J$ = 8.9 Hz, 2 H), 7.70 (d, $J$ = 8.2 Hz, 1 H), 7.68 (d, $J$ = 7.9 Hz, 1 H), 7.31 (d, $J$ = 7.9 Hz, 1 H), 7.21 (s, 2 H), 7.10 - 6.99 (m, 6 H), 6.96 (t, $J$ = 7.5 Hz, 1 H), 6.68 (d, $J$ = 8.5 Hz, 2 H), 5.50 (dd, $J$ = 7.3, 8.9 Hz, 1 H), 5.18 (ddd, $J$ = 3.3, 8.6, 11.6 Hz, 1 H), 4.99 - 4.91 (m, 2 H), 4.69 (sxt, $J$ = 6.3 Hz, 1 H), 4.53 - 4.43 (m, 2 H), 4.03 (t, $J$ = 5.8 Hz, 2 H), 3.88 (s, 6 H), 3.12 (s, 3 H), 3.04 - 2.99 (m, 2 H), 2.85 - 2.77 (m, 1 H), 2.63 (t, $J$ = 6.0 Hz, 2 H), 2.70 - 2.56 (m, 4 H), 2.29 (br s, 4 H), 2.18 (dd, $J$ = 11.7, 14.2 Hz, 1 H), 2.13 (s, 3 H), 1.92 - 1.81 (m, 2 H), 1.93 - 1.72 (m, 5 H), 1.50 (s, 3 H), 1.54 - 1.46 (m, 1 H), 1.44 - 1.34 (m, 1 H), 1.15 (d, $J$ = 6.4 Hz, 3 H), 0.95 (d, $J$ = 6.7 Hz, 3 H); $^{13}$C-NMR (126 MHz, DMSO-$d_6$): $\delta$ = 175.5, 170.7, 170.5, 170.0, 168.0, 160.6, 156.8, 153.9, 148.2, 146.9, 139.5, 136.6, 133.5, 133.1, 127.5, 127.5, 124.8, 123.8, 123.7, 121.3, 119.2, 118.5, 115.8, 115.5, 111.6, 110.0, 100.4, 71.2, 71.0, 67.3, 57.8, 56.5, 55.3, 55.2, 53.4, 49.4, 48.6, 46.3, 43.2, 41.9, 40.7, 38.3, 35.3, 31.0, 25.7, 24.1, 20.0, 19.8, 17.2; IR: $\tilde{\nu}_{max}$ = 3287, 2936, 1728, 1670, 1597, 1497, 1458, 1415, 1327, 1223, 1126, 1006, 837, 745, 652; UV-VIS (PBS/MeCN, 2:1): $\lambda_{max}$ ($\varepsilon$) = 362 nm (24.7 × $10^3$ l·mol$^{-1}$·cm$^{-1}$); HRMS (ESI): $m/Z$ calcd for $C_{58}H_{74}N_9O_{11}$ $[M+H]^+$: 1072.550, found: 1072.551.

*cyclo*-[(2*S*,4*E*,8*S*)-Hdn-L-Dap(2-(4´-((4´´-benzyloxy-3´´,5´´-dimethoxyphenyl)-diazenyl)phenoxy)acetyl)-D-*N*MeTrp-L-*β*Tyr]; (**S43**)

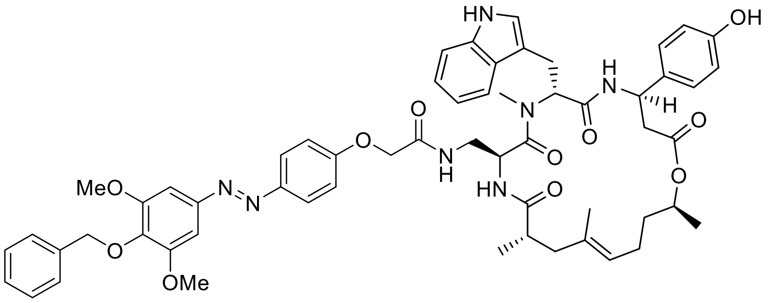

Standard Procedure **SP4** with cyclodepsipeptide **S54** (2.5 mg, 2.8 µmol, 1.0 equiv.), 2-(4´-((4´´-benzyloxy-3´´,5´´-dimethoxyphenyl)diazenyl)phenoxy)acetic acid (1.4 mg, 3.4 µmol, 1.2 equiv.), HATU (2.6 mg, 6.8 µmol, 2.4 equiv.) and DIPEA (3.8 µl, 22.5 µmol, 8.0 equiv.) gave conjugate **S43** (2.1 mg, 72%) as an orange solid after purification by preparative HPLC ($H_2O$/MeCN, 60:40 → 0:100). $R_f$ = 0.39 ($CH_2Cl_2$/MeOH, 14:1); $[\alpha]_D^{24}$ = 36.3 (c = 0.05 in MeCN); $^1$H-NMR (400 MHz, DMSO-$d_6$): $\delta$ = 10.79 (d, $J$ = 1.8 Hz, 1 H), 9.28 (s, 1 H), 8.56 (d, $J$ = 8.8 Hz, 1 H), 7.98 (t, $J$ = 6.0 Hz, 1 H), 7.86 (d, $J$ = 8.8 Hz, 2 H), 7.68 (d, $J$ = 7.6 Hz, 1 H), 7.63 (d, $J$ = 8.8 Hz, 1 H), 7.47 (d, $J$ = 7.0 Hz, 2 H), 7.40 - 7.27 (m, 4 H), 7.22 (s, 2 H), 7.09 - 7.00 (m, 6 H), 6.95 (t, $J$ = 7.3 Hz, 1 H), 6.66 (d, $J$ = 8.5 Hz, 2 H), 5.50 (dd, $J$ = 7.3, 9.1 Hz, 1 H), 5.17 (dt, $J$ = 3.5, 9.5 Hz, 1 H), 5.01 (s, 2 H), 4.98 - 4.90 (m, 2 H), 4.69 (sxt, $J$ = 6.3 Hz, 1 H), 4.47 (d, $J$ = 2.0 Hz, 2 H), 3.88 (s, 6 H), 3.11 (s, 3 H), 3.07 - 2.96 (m, 2 H), 2.86 - 2.75 (m, 1 H), 2.70 - 2.54 (m, 4 H), 2.17 (dd, $J$ = 11.7, 14.0 Hz, 1 H), 1.85 (spt, $J$ = 7.0 Hz, 2 H), 1.76 (d, $J$ = 13.4 Hz, 1 H), 1.49 (s, 3 H), 1.56 - 1.44 (m, 1 H), 1.43 - 1.32 (m, 1 H), 1.14 (d, $J$ = 6.1 Hz, 3 H), 0.94 (d, $J$ = 6.7 Hz, 3 H); $^{13}$C-NMR, HSQC/HMBC (100 MHz, DMSO-$d_6$): $\delta$ = 175.2, 170.8, 170.2, 169.7, 167.8, 160.3, 156.6, 153.8, 148.1, 146.7, 139.0, 137.8, 136.4, 133.4, 132.9, 128.4, 128.2, 127.3, 124.7, 123.8, 123.5, 121.1, 119.1, 118.3, 115.7, 115.3, 111.4, 109.9, 100.3, 74.5, 71.1, 67.3, 56.3, 55.2, 49.2, 48.2, 43.2, 41.6, 38.2, 35.1, 30.8, 29.1, 25.4, 23.9, 19.7, 19.3, 17.5; UV-VIS (PBS/MeCN, 2:1): $\lambda_{max}$ ($\varepsilon$) = 364 nm (18.2 × $10^3$ l·mol$^{-1}$·cm$^{-1}$); HRMS (ESI): *m/Z* calcd for $C_{58}H_{66}N_7O_{11}$ $[M+H]^+$: 1036.482, found: 1036.482.

*cyclo*-[(2*S*,4*E*,8*S*)-Hdn-L-Lys(Glu(2-(4´-((3´´,4´´,5´´-trimethoxyphenyl)diazenyl)-phenoxy)acetyl))-D-*N*MeTrp-L-*β*Tyr]; (**S44**)

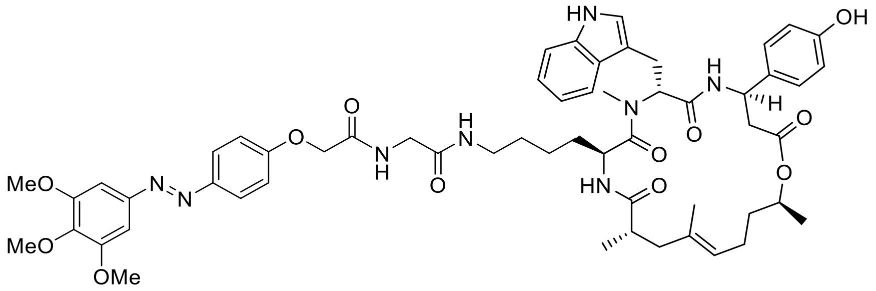

Standard Procedure **SP4** with cyclodepsipeptide **S69** (10.0 mg, 10.1 µmol, 1.0 equiv.), 2-(4´-((3´´,4´´,5´´-trimethoxyphenyl)diazenyl)phenoxy)acetic acid (3.5 mg, 10.1 µmol, 1.0 equiv.), HATU (9.2 mg, 24.3 µmol, 2.4 equiv.) and DIPEA (13.8 µl, 81.0 µmol, 8.0 equiv.) gave conjugate **S44** (7.7 mg, 72%) as a yellow solid after purification by silica-gel chromatography ($CH_2Cl_2$/petroleum ether/acetone, 3:1:1 → 1:1:3). $R_f$ = 0.33 ($CH_2Cl_2$/MeOH, 14:1); $[\alpha]_D^{24}$ = 88.3 (c = 0.05 in MeCN); $^1$H-NMR (600 MHz, DMSO-$d_6$): $\delta$ = 10.82 (br s, 1 H), 9.38 (br s, 1 H), 8.65 (d, $J$ = 8.8 Hz, 1 H), 8.37 (t, $J$ = 5.6 Hz, 1 H), 7.90 (d, $J$ = 9.0 Hz, 2 H), 7.85 (t, $J$ = 5.7 Hz, 1 H), 7.71 (d, $J$ = 8.6 Hz, 1 H), 7.67 (d, $J$ = 7.9 Hz, 1 H), 7.31 (d, $J$ = 8.1 Hz, 1 H), 7.23 (s, 2 H), 7.18 (d, $J$ = 9.0 Hz, 2 H), 7.13 (d, $J$ = 8.6 Hz, 2 H), 7.07 (d, $J$ = 2.0 Hz, 1 H), 7.04 (t, $J$ = 7.6 Hz, 1 H), 6.95 (t, $J$ = 7.0 Hz, 1 H), 6.70 (d, $J$ = 8.4 Hz, 2 H), 5.52 (dd, $J$ = 5.0, 11.6 Hz, 1 H), 5.19 (ddd, $J$ = 2.9, 9.0, 11.7 Hz, 1 H), 4.93 (t, $J$ = 6.6 Hz, 1 H), 4.71 - 4.64 (m, 3 H), 4.59 - 4.51 (m, 1 H), 3.89 (s, 6 H), 3.76 (s, 3 H), 3.79 - 3.72 (m, 2 H), 3.04 (s, 3 H), 3.10 - 2.98 (m, 1 H), 2.96 - 2.88 (m, 2 H), 2.84 (qd, $J$ = 6.6, 13.3 Hz, 1 H), 2.68 (dd, $J$ = 11.6, 14.6 Hz, 1 H), 2.59 (dd, $J$ = 2.9, 14.5 Hz, 1 H), 2.53 - 2.52 (m, 2 H), 2.18 (dd, $J$ = 11.9, 14.5 Hz, 1 H), 1.91 - 1.77 (m, 2 H), 1.73 (d, $J$ = 14.5 Hz, 1 H), 1.49 (s, 3 H), 1.53 - 1.45 (m, 1 H), 1.43 - 1.32 (m, 1 H), 1.16 (d, $J$ = 6.2 Hz, 3 H), 1.14 - 1.04 (m, 2 H), 0.93 (d, $J$ = 6.8 Hz, 3 H), 0.88 - 0.78 (m, 2 H), 0.73 - 0.64 (m, 1 H); $^{13}$C-NMR (151 MHz, DMSO-$d_6$): $\delta$ = 174.5, 172.3, 170.3, 170.1, 168.2, 167.6, 160.2, 156.3, 153.3, 147.8, 146.4, 139.9, 136.1, 133.1, 133.0, 127.1, 127.0, 124.4, 123.4, 123.0, 120.8, 118.6, 118.0, 115.4, 115.1, 111.2, 109.5, 100.0, 71.0, 67.0, 60.2, 56.0, 55.9, 54.6, 49.0, 47.6, 42.5, 41.7, 40.1, 38.2, 37.6, 34.9, 30.6, 28.6, 25.6, 23.7, 22.0, 19.7, 19.5, 16.9; IR: $\tilde{\nu}_{max}$ = 3383, 3290, 2937, 1732 1651, 1504, 1458, 1230, 1126, 1002, 837, 744, 652; UV-VIS (PBS/MeCN, 2:1): $\lambda_{max}$ ($\varepsilon$) = 361 nm (19.9 × 10$^3$ l·mol$^{-1}$·cm$^{-1}$); HRMS (ESI): *m/Z* calcd for $C_{57}H_{71}N_8O_{12}$ [M+H]$^+$: 1059.519, found: 1059.519.

*cyclo*-[(2*S*,4*E*,8*S*)-Hdn-L-Lys(Eaca(2-(4′-((3′′,4′′,5′′-trimethoxyphenyl)diazenyl)-phenoxy)acetyl))-D-*N*MeTrp-L-*β*Tyr]; (**S45**)

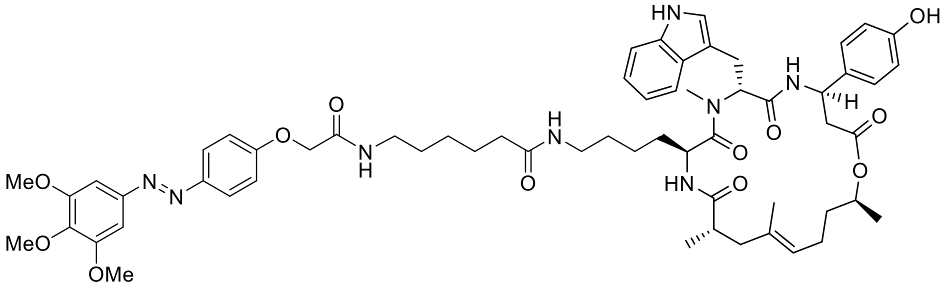

Standard Procedure **SP4** with cyclodepsipeptide **S70** (10.0 mg, 9.6 µmol, 1.0 equiv.), 2-(4′-((3′′,4′′,5′′-trimethoxyphenyl)diazenyl)phenoxy)acetic acid (3.3 mg, 9.6 µmol, 1.0 equiv.), HATU (8.8 mg, 23.0 µmol, 2.4 equiv.) and DIPEA (13.0 µl, 76.7 µmol, 8.0 equiv.) gave conjugate **S45** (6.0 mg, 56%) as a yellow solid after purification by silica-gel chromatography ($CH_2Cl_2$/petroleum ether/acetone, 3:1:1 → 1:1:3). $R_f$ = 0.38 ($CH_2Cl_2$/MeOH, 14:1); $[\alpha]_D^{24}$ = 69.6 (c = 0.06 in MeCN); $^1$H-NMR (500 MHz, DMSO-$d_6$): $\delta$ = 10.82 (d, $J$ = 1.8 Hz, 1 H), 9.31 (s, 1 H), 8.65 (d, $J$ = 8.9 Hz, 1 H), 8.16 (t, $J$ = 5.6 Hz, 1 H), 7.89 (d, $J$ = 8.9 Hz, 2 H), 7.72 - 7.64 (m, 3 H), 7.29 (d, $J$ = 8.2 Hz, 1 H), 7.23 (s, 2 H), 7.16 - 7.11 (m, 4 H), 7.06 (d, $J$ = 1.8 Hz, 1 H), 7.03 (ddd, $J$ = 0.9, 7.0, 7.9 Hz, 1 H), 6.95 (ddd, $J$ = 0.9, 7.0, 7.9 Hz, 1 H), 6.70 (d, $J$ = 8.5 Hz, 2 H), 5.52 (dd, $J$ = 5.0, 11.4 Hz, 1 H), 5.19 (ddd, $J$ = 2.7, 8.9, 11.3 Hz, 1 H), 4.92 (t, $J$ = 6.6 Hz, 1 H), 4.72 - 4.63 (m, 1 H), 4.59 (s, 2 H), 4.58 - 4.52 (m, 1 H), 3.89 (s, 6 H), 3.76 (s, 3 H), 3.13 (q, $J$ = 6.7 Hz, 2 H), 3.03 (s, 3 H), 3.06 - 2.99 (m, 1 H), 2.93 (dd, $J$ = 4.9, 15.0 Hz, 1 H), 2.87 (td, $J$ = 6.7, 13.4 Hz, 1 H), 2.83 - 2.75 (m, 1 H), 2.68 (dd, $J$ = 11.3, 14.6 Hz, 1 H), 2.59 (dd, $J$ = 3.1, 14.6 Hz, 1 H), 2.53 - 2.52 (m, 2 H), 2.17 (dd, $J$ = 11.9, 14.6 Hz, 1 H), 2.03 (t, $J$ = 7.3 Hz, 2 H), 1.90 - 1.77 (m, 2 H), 1.73 (d, $J$ = 15.0 Hz, 1 H), 1.48 (s, 3 H), 1.53 - 1.40 (m, 5 H), 1.40 - 1.33 (m, 1 H), 1.27 - 1.19 (m, 2 H), 1.16 (d, $J$ = 6.4 Hz, 3 H), 1.14 - 1.03 (m, 2 H), 0.92 (d, $J$ = 6.7 Hz, 3 H), 0.85 - 0.77 (m, 2 H), 0.70 - 0.64 (m, 1 H); $^{13}$C-NMR (126 MHz, DMSO-$d_6$): $\delta$ = 174.9, 172.8, 172.4, 170.8, 170.5, 167.4, 160.7, 156.7, 153.8, 148.3, 146.8, 140.3, 136.6, 133.6, 133.5, 127.6, 127.4, 124.8, 123.8, 123.4, 121.3, 119.1, 118.5, 115.8, 115.5, 111.6, 110.0, 100.4, 71.4, 67.6, 60.7, 56.4, 55.0, 49.5, 48.1, 43.0, 42.1, 38.7, 38.5, 38.1, 35.8, 35.3, 31.0, 29.4, 29.1, 26.5, 26.0, 26.0, 25.5, 24.2, 22.5, 20.2, 19.9, 17.4; IR: $\tilde{\nu}_{max}$ = 3290, 2940, 1732, 1651, 1504, 1454, 1369, 1231, 1130, 1001, 840, 744, 652; UV-VIS (PBS/MeCN, 2:1): $\lambda_{max}$ ($\varepsilon$) = 361 nm (18.1 × 10$^3$ l·mol$^{-1}$·cm$^{-1}$); HRMS (ESI): *m/Z* calcd for $C_{61}H_{79}N_8O_{12}$ $[M+H]^+$: 1115.581, found: 1115.580.

*cyclo*-[(2*S*,4*E*,8*S*)-Hdn-L-Lys(Glu(4-((5′-methoxy-2′-((3′′,4′′,5′′-trimethoxyphenyl)-diazenyl)phenyl)amino)-4-oxobutyryl))-D-*N*MeTrp-L-*β*Tyr]; (**S46**)

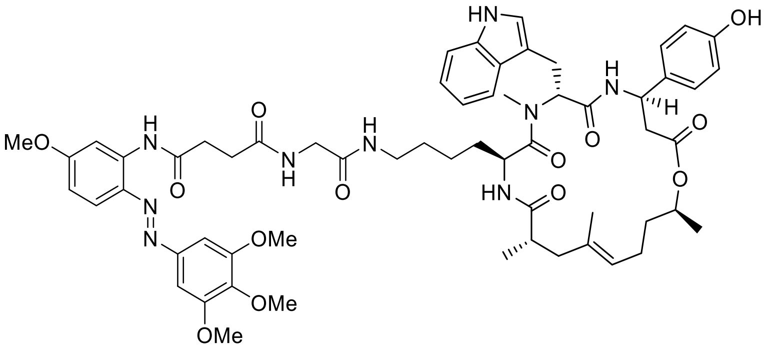

Standard Procedure **SP4** with cyclodepsipeptide **S69** (10.0 mg, 10.1 µmol, 1.0 equiv.), *N*-(5′-Methoxy-2′-((3′′,4′′,5′′-trimethoxyphenyl)diazenyl)phenyl)succinamic acid (4.2 mg, 10.1 µmol, 1.0 equiv.), HATU (9.2 mg, 24.3 µmol, 2.4 equiv.) and DIPEA (13.8 µl, 81.0 µmol, 8.0 equiv.) gave conjugate **S46** (5.9 mg, 52%) as an orange solid after purification by silica-gel chromatography ($CH_2Cl_2$/petroleum ether/acetone, 3:1:1 → 1:1:3). $R_f$ = 0.33 ($CH_2Cl_2$/MeOH, 14:1); $[\alpha]_D^{24}$ = 64.6 (c = 0.06 in MeCN); $^1$H-NMR (600 MHz, DMSO-$d_6$): $\delta$ = 10.77 (s, 1 H), 10.29 (s, 1 H), 9.31 (s, 1 H), 8.64 (d, *J* = 8.8 Hz, 1 H), 8.20 (t, *J* = 5.8 Hz, 1 H), 7.99 (d, *J* = 2.6 Hz, 1 H), 7.74 (d, *J* = 9.0 Hz, 1 H), 7.71 - 7.67 (m, 2 H), 7.65 (d, *J* = 7.9 Hz, 1 H), 7.37 (s, 2 H), 7.28 (d, *J* = 7.9 Hz, 1 H), 7.12 (d, *J* = 8.6 Hz, 2 H), 7.04 (d, *J* = 2.2 Hz, 1 H), 7.02 (t, *J* = 7.8 Hz, 1 H), 6.93 (t, *J* = 7.6 Hz, 1 H), 6.78 (dd, *J* = 2.6, 9.0 Hz, 1 H), 6.69 (d, *J* = 8.6 Hz, 2 H), 5.50 (dd, *J* = 5.0, 11.3 Hz, 1 H), 5.18 (ddd, *J* = 2.9, 8.9, 11.2 Hz, 1 H), 4.91 (t, *J* = 6.6 Hz, 1 H), 4.66 (sxt, *J* = 6.4 Hz, 1 H), 4.54 (dt, *J* = 3.5, 8.3 Hz, 1 H), 3.89 (s, 6 H), 3.82 (s, 3 H), 3.75 (s, 3 H), 3.64 (d, *J* = 5.7 Hz, 2 H), 3.02 (s, 3 H), 3.07 - 2.98 (m, 1 H), 2.91 (dd, *J* = 4.5, 14.9 Hz, 1 H), 2.86 (td, *J* = 6.8, 13.6 Hz, 1 H), 2.83 - 2.79 (m, 1 H), 2.77 (t, *J* = 6.8 Hz, 2 H), 2.67 (dd, *J* = 11.6, 14.6 Hz, 1 H), 2.58 (dd, *J* = 2.9, 14.7 Hz, 1 H), 2.54 (t, *J* = 6.6 Hz, 2 H), 2.52 - 2.51 (m, 2 H), 2.16 (dd, *J* = 11.7, 14.5 Hz, 1 H), 1.89 - 1.76 (m, 2 H), 1.71 (d, *J* = 14.5 Hz, 1 H), 1.47 (s, 3 H), 1.52 - 1.45 (m, 1 H), 1.41 - 1.32 (m, 1 H), 1.15 (d, *J* = 6.4 Hz, 3 H), 1.12 - 1.02 (m, 2 H), 0.91 (d, *J* = 6.8 Hz, 3 H), 0.83 - 0.75 (m, 2 H), 0.70 - 0.62 (m, 1 H); $^{13}$C-NMR, HSQC (150 MHz, DMSO-$d_6$): $\delta$ = 127.3, 123.7, 123.7, 121.2, 118.9, 118.9, 118.5, 115.4, 111.4, 110.4, 106.1, 100.8, 71.2, 60.6, 56.5, 55.9, 54.9, 49.5, 47.9, 42.8, 42.4, 42.1, 38.6, 37.9, 35.1, 32.5, 31.0, 30.9, 30.7, 29.0, 25.9, 23.9, 22.3, 19.9, 19.6, 17.1; IR: $\tilde{\nu}_{max}$ = 3312, 2936, 1651, 1520, 1458, 1288, 1234, 1126, 1003, 837, 744, 651; UV-VIS (PBS/MeCN, 2:1): $\lambda_{max}$ ($\varepsilon$) = 385 nm (19.7 × $10^3$ l·mol$^{-1}$·cm$^{-1}$); HRMS (ESI): *m/Z* calcd for $C_{60}H_{76}N_9O_{13}$ [M+H]$^+$: 1130.556, found: 1130.556.

*cyclo*-[(2*S*,4*E*,8*S*)-Hdn-L-Lys(Eaca(4-((5′-methoxy-2′-((3′′,4′′,5′′-trimethoxyphenyl)-diazenyl)phenyl)amino)-4-oxobutyryl))-D-*N*MeTrp-L-*β*Tyr]; (**S47**)

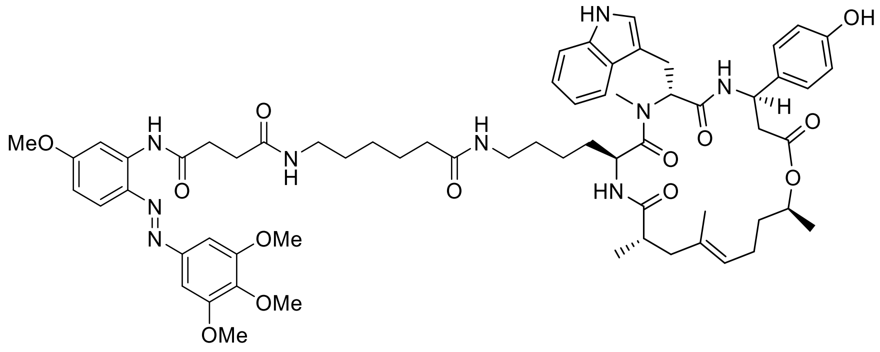

Standard Procedure **SP4** with cyclodepsipeptide **S70** (5.0 mg, 4.8 µmol, 1.0 equiv.), *N*-(5′-Methoxy-2′-((3′′,4′′,5′′-trimethoxyphenyl)diazenyl)phenyl)succinamic acid (2.0 mg, 4.8 µmol, 1.0 equiv.), HATU (4.4 mg, 11.5 µmol, 2.4 equiv.) and DIPEA (6.5 µl, 38.3 µmol, 8.0 equiv.) gave conjugate **S47** (3.2 mg, 57%) as a yellow solid after purification by silica-gel chromatography ($CH_2Cl_2$/petroleum ether/acetone, 3:1:1 → 1:1:3). $R_f$ = 0.25 ($CH_2Cl_2$/MeOH, 14:1); $[\alpha]_D^{24}$ = 94.1 (c = 0.06 in MeCN); $^1$H-NMR (600 MHz, DMSO-$d_6$): $\delta$ = 10.83 (br s, 1 H), 10.27 (s, 1 H), 9.33 (br s, 1 H), 8.66 (d, *J* = 8.8 Hz, 1 H), 8.05 (d, *J* = 2.9 Hz, 1 H), 7.92 (t, *J* = 5.6 Hz, 1 H), 7.75 (d, *J* = 9.0 Hz, 1 H), 7.70 (d, *J* = 8.8 Hz, 1 H), 7.67 (d, *J* = 7.7 Hz, 1 H), 7.64 (t, *J* = 5.6 Hz, 1 H), 7.41 (s, 2 H), 7.29 (d, *J* = 8.1 Hz, 1 H), 7.13 (d, *J* = 8.6 Hz, 2 H), 7.06 (d, *J* = 2.0 Hz, 1 H), 7.03 (t, *J* = 7.2 Hz, 1 H), 6.95 (t, *J* = 7.4 Hz, 1 H), 6.78 (dd, *J* = 2.9, 9.0 Hz, 1 H), 6.70 (d, *J* = 8.6 Hz, 2 H), 5.52 (dd, *J* = 5.0, 11.6 Hz, 1 H), 5.19 (ddd, *J* = 2.6, 9.2, 11.4 Hz, 1 H), 4.92 (t, *J* = 6.5 Hz, 1 H), 4.67 (qd, *J* = 6.4, 12.8 Hz, 1 H), 4.59 - 4.51 (m, 1 H), 3.92 (s, 6 H), 3.84 (s, 3 H), 3.76 (s, 3 H), 3.03 (s, 3 H), 3.07 - 3.01 (m, 1 H), 2.99 (q, *J* = 6.6 Hz, 2 H), 2.92 (dd, *J* = 4.6, 15.0 Hz, 1 H), 2.86 (tt, *J* = 6.7, 13.5 Hz, 1 H), 2.78 (dt, *J* = 6.6, 13.3 Hz, 1 H), 2.73 (t, *J* = 6.8 Hz, 2 H), 2.68 (dd, *J* = 11.6, 14.6 Hz, 1 H), 2.59 (dd, *J* = 2.6, 14.3 Hz, 1 H), 2.54 - 2.52 (m, 2 H), 2.47 (t, *J* = 7.3 Hz, 2 H), 2.17 (dd, *J* = 11.8, 14.6 Hz, 1 H), 1.97 (t, *J* = 7.4 Hz, 2 H), 1.91 - 1.77 (m, 2 H), 1.73 (d, *J* = 14.5 Hz, 1 H), 1.48 (s, 3 H), 1.53 - 1.44 (m, 1 H), 1.43 - 1.35 (m, 4 H), 1.32 (qu, *J* = 7.0 Hz, 2 H), 1.16 (d, *J* = 6.2 Hz, 3 H), 1.15 - 1.04 (m, 3 H), 0.92 (d, *J* = 6.8 Hz, 3 H), 0.87 - 0.76 (m, 2 H), 0.70 - 0.62 (m, 1 H); $^{13}$C-NMR, HSQC (150 MHz, DMSO-$d_6$): $\delta$ = 127.5, 123.6, 123.3, 121.1, 118.9, 18.4, 118.2, 115.3, 111.4, 110.2, 105.7, 100.8, 71.4, 60.5, 56.3, 55.8, 55.0, 49.2, 47.8, 42.7, 42.0, 40.1, 38.9, 38.4, 35.7, 35.2, 32.9, 30.8, 30.8, 29.3, 29.2, 29.1, 28.5, 26.3, 25.4, 23.9, 22.3, 19.8, 19.6, 17.2; IR: $\tilde{\nu}_{max}$ = 3313, 2936, 1651, 1520, 1458, 1373, 1288, 1234, 1126, 1002, 837, 744, 652; UV-VIS (PBS/MeCN, 2:1): $\lambda_{max}$ ($\varepsilon$) = 386 nm (21.8 × 10$^3$ l·mol$^{-1}$·cm$^{-1}$); HRMS (ESI): *m/Z* calcd for $C_{64}H_{84}N_9O_{13}$ $[M+H]^+$: 1186.618, found: 1186.617.

**Synthesis of Jasplakinolide analogs:**

*cyclo*-[(2*S*,4*E*)-Hdo-L-Dap(Boc)-D-*N*MeTrp-L-*β*Tyr]; (**S4**)

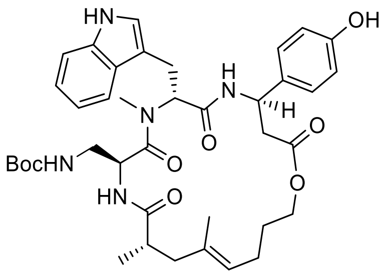

Standard Procedure **SP3** with cyclodepsipeptide **S64** (7.0 mg, 7.98 µmol, 1.0 equiv.) gave cyclodepsipeptide **S4** (3.0 mg, 52%) as a colorless solid after purification by preparative HPLC ($H_2O$/MeCN, 70:30 → 0:100). $R_f$ = 0.24 ($CH_2Cl_2$/MeOH, 19:1); $[\alpha]_D^{24}$ = 27.7 (c = 0.20 in MeCN); $^1$H-NMR (500 MHz, DMSO-$d_6$): $\delta$ = 10.79 (d, $J$ = 0.9 Hz, 1 H), 9.30 (s, 1 H), 8.56 (d, $J$ = 8.5 Hz, 1 H), 7.66 (d, $J$ = 7.9 Hz, 1 H), 7.47 (d, $J$ = 8.9 Hz, 1 H), 7.31 (d, $J$ = 8.2 Hz, 1 H), 7.09 - 7.00 (m, 4 H), 6.95 (dt, $J$ = 0.9, 7.3 Hz, 1 H), 6.68 (d, $J$ = 8.5 Hz, 2 H), 6.41 (t, $J$ = 6.1 Hz, 1 H), 5.48 (dd, $J$ = 6.1, 10.1 Hz, 1 H), 5.20 (ddd, $J$ = 3.1, 8.2, 11.9 Hz, 1 H), 4.95 (t, $J$ = 7.0 Hz, 1 H), 4.88 (dt, $J$ = 4.4, 8.6 Hz, 1 H), 4.06 (td, $J$ = 7.2, 10.7 Hz, 1 H), 3.85 (td, $J$ = 7.1, 10.9 Hz, 1 H), 3.04 (s, 3 H), 3.14 - 2.97 (m, 2 H), 2.75 - 2.54 (m, 4 H), 2.48 - 2.42 (m, 1 H), 2.16 (dd, $J$ = 11.6, 14.6 Hz, 1 H), 1.85 (q, $J$ = 7.6 Hz, 2 H), 1.78 (d, $J$ = 14.3 Hz, 1 H), 1.52 (s, 3 H), 1.46 (dt, $J$ = 6.6, 14.0 Hz, 2 H), 1.30 (s, 9 H), 0.97 (d, $J$ = 7.0 Hz, 3 H); $^{13}$C-NMR (126 MHz, DMSO-$d_6$): $\delta$ = 175.1, 170.8, 170.1, 156.7, 155.9, 136.5, 133.6, 133.2, 127.5, 127.4, 123.8, 123.6, 121.4, 119.2, 118.5, 115.5, 111.6, 110.1, 78.2, 64.3, 55.3, 49.1, 48.9, 43.4, 42.1, 41.7, 38.3, 31.1, 28.6, 28.4, 25.8, 24.3, 19.7, 17.1; IR: $\tilde{\nu}_{max}$ = 3314, 2974, 2932, 1709, 1639, 1516, 1454, 1269, 1250, 1169, 1053, 833, 745, 652; HRMS (ESI): *m/Z* calcd for $C_{42}H_{58}N_5O_8$ $[M+H]^+$: 718.3810, found: 718.3811.

*cyclo*-[(2*S*,4*E*)-Hdo-L-Dap(Boc)-D-*N*MeTrp-L-*β*Tyr(TIPS)]; (**S64**)

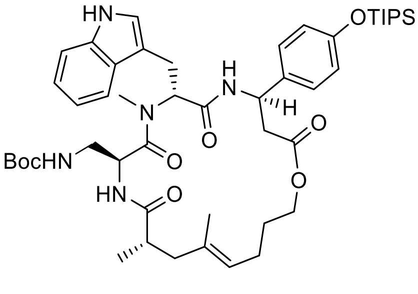

Standard Procedure **SP2** with diene **S71** (80.0 mg, 88.7 µmol, 1.0 equiv.) and 2nd generation Grubbs catalyst (9.0 mg, 11.0 µmol, 0.12 equiv) gave cyclodepsipeptide **S64** (44 mg, 57%) as a colorless solid after purification by silica-gel chromatography (EtOAc/Petrolether, 1:2 → 2:1). $R_f$ = 0.27 ($CH_2Cl_2$/MeOH, 19:1); $[\alpha]_D^{24}$ = 44.1 (c = 1.0 in MeCN); $^1$H-NMR (400 MHz, DMSO-$d_6$): $\delta$ = 10.80 (br s, 1 H), 8.58 (d, $J$ = 8.8 Hz, 1 H), 7.65 (d, $J$ = 7.9 Hz, 1 H), 7.48 (d, $J$ = 8.5 Hz, 1 H), 7.30 (d, $J$ = 8.2 Hz, 1 H), 7.11 (d, $J$ = 8.5 Hz, 2 H), 7.07 - 7.00 (m, 2 H), 6.95 (t, $J$ = 7.6 Hz, 1 H), 6.76 (d, $J$ = 8.5 Hz, 2 H), 6.42 (t, $J$ = 6.0 Hz, 1 H), 5.49 (dd, $J$ = 6.3, 9.8 Hz, 1 H), 5.24 (ddd, $J$ = 2.6, 8.5, 11.1 Hz, 1 H), 4.95 (t, $J$ = 7.0 Hz, 1 H), 4.89 (dt, $J$ = 5.0, 8.3 Hz, 1 H), 4.12 - 3.99 (m, 1 H), 3.91 - 3.81 (m, 1 H), 3.06 (s, 3 H), 3.15 - 2.91 (m, 2 H), 2.80 - 2.54 (m, 4 H), 2.16 (dd, $J$ = 12.3, 14.0 Hz, 1 H), 1.85 (sxt, $J$ = 7.6 Hz, 2 H), 1.79 (d, $J$ = 14.3 Hz, 1 H), 1.52 (s, 3 H), 1.49 - 1.37 (m, 2 H), 1.30 (s, 9 H), 1.28 - 1.16 (m, 4 H), 1.05 (d, $J$ = 7.3 Hz, 18 H), 0.97 (d, $J$ = 6.7 Hz, 3 H), $^{13}$C-NMR (126 MHz, DMSO-$d_6$): $\delta$ = 175.1, 170.8, 170.7, 170.2, 155.9, 154.7, 136.5, 135.4, 133.6, 127.7, 127.4, 123.7, 123.6, 121.3, 119.7, 119.2, 118.5, 111.6, 110.1, 78.2, 64.3, 60.2, 55.3, 49.0, 48.9, 43.4, 41.7, 38.3, 31.1, 28.6, 28.6, 25.8, 24.3, 19.7, 18.2, 17.0, 12.5; IR: $\tilde{\nu}_{max}$ = 3325, 2943, 2866, 2362, 2745, 1639, 1512, 1458, 1265, 1169, 1099, 914, 883, 740, 683; HRMS (ESI): *m/Z* calcd for $C_{48}H_{72}N_5O_8Si$ $[M+H]^+$: 874.5145, found: 874.5157.

*cyclo*-[(2*S*,4*E*)-Hdo-L-Lys(Boc)-D-*N*MeTrp-L-*β*Tyr(TIPS)]; (**S65**)

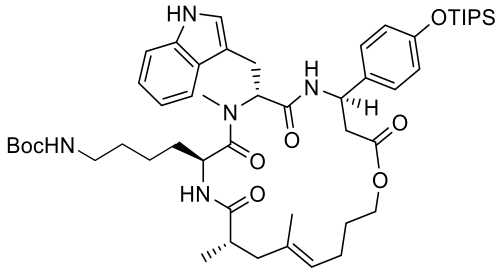

Standard Procedure **SP2** with diene **S72** (74.0 mg, 78.4 µmol, 1.0 equiv.) and 2nd generation Grubbs catalyst (98.0 mg, 9.8 µmol, 0.12 equiv) gave cyclodepsipeptide **S65** (57.3 mg, 80%) as a colorless solid after purification by silica-gel chromatography ($CH_2Cl_2$/MeOH, 98:2). $R_f$ = 0.30 ($CH_2Cl_2$/MeOH, 19:1); $[\alpha]_D^{24}$ = 56.4 (c = 3.0 in MeCN); $^1$H-NMR (300 MHz, DMSO-$d_6$): $\delta$ = 10.77 (d, $J$ = 1.4 Hz, 1 H), 8.65 (d, $J$ = 8.8 Hz, 1 H), 7.72 - 7.60 (m, 2 H), 7.28 (d, $J$ = 7.9 Hz, 1 H), 7.21 (d, $J$ = 8.6 Hz, 2 H), 7.07 (d, $J$ = 1.9 Hz, 1 H), 7.02 (ddd, $J$ = 0.7, 7.0, 8.0 Hz, 1 H), 6.93 (ddd, $J$ = 0.7, 7.0, 8.0 Hz, 1 H), 6.80 (d, $J$ = 8.6 Hz, 2 H), 6.65 (t, $J$ = 5.6 Hz, 1 H), 5.51 (dd, $J$ = 7.4, 8.9 Hz, 1 H), 5.26 (ddd, $J$ = 3.6, 8.4, 11.0 Hz, 1 H), 4.92 (t, $J$ = 6.8 Hz, 1 H), 4.61 - 4.50 (m, 1 H), 4.12 - 4.01 (m, 1 H), 3.89 - 3.77 (m, 1 H), 2.99 (s, 3 H), 3.07 - 2.95 (m, 2

H), 2.82 - 2.58 (m, 5 H), 2.13 (dd, *J* = 11.7, 14.4 Hz, 1 H), 1.85 - 1.75 (m, 2 H), 1.68 (d, *J* = 22.4 Hz, 1 H), 1.49 (s, 3 H), 1.49 - 1.37 (m, 1 H), 1.36 (s, 9 H), 1.27 - 1.17 (m, 3 H), 1.04 (d, *J* = 7.1 Hz, 21 H), 0.91 (d, *J* = 6.7 Hz, 3 H), 0.75 (br s, 2 H), 0.66 - 0.58 (m, 1 H); $^{13}$C-NMR (101 MHz, DMSO-$d_6$): $\delta$ = 174.8, 172.8, 170.7, 170.6, 156.8, 156.0, 136.6, 135.7, 133.5, 127.5, 127.4, 123.8, 123.5, 121.3, 119.8, 119.2, 118.5, 111.6, 110.1, 77.8, 64.5, 55.2, 49.2, 48.0, 43.2, 42.3, 42.0, 38.0, 31.3, 31.2, 29.5, 28.7, 28.6, 26.1, 24.4, 22.4, 20.0, 18.3, 17.2, 12.5, IR: $\tilde{\nu}_{max}$ = 3321, 2943, 2866, 2361, 1732, 1686, 1635, 1512, 1458, 1365, 1265, 1173, 1103, 914, 883, 741, 683, 741, 683; HRMS (ESI): *m/Z* calcd for $C_{51}H_{78}N_5O_8Si$ [M+H]$^+$: 916.5614, found: 916. 5618.

Jasplakinolide-analogs **S66** and **S67**:

Standard Procedure **SP2** with diene **S73** (200 mg, 0.2 mmol, 1 equiv.) and 2$^{nd}$ generation Grubbs catalyst (83.2 mg, 9.8 µmol, 0.05 equiv) gave *E*-cyclodepsipeptide **S66** (68.0 mg, 36%) and *Z*-cyclodepsipeptide **S67** (51.0 mg, 27%) as brown solids after purification by silica-gel chromatography (petroleum ether/EtOAc, 1:1).

*cyclo*-[(2*S*,4*E*,6*R*,8*S*)Htn-L-Lys(Boc)-D-*N*-Me-Trp-L-β-Tyr(TIPS)]; ()

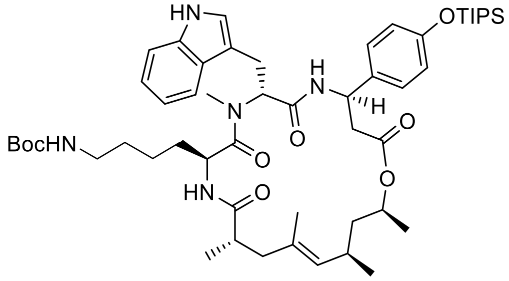

$R_f$ = 0.18 (petroleum ether/EtOAc, 1:1); $[\alpha]_D^{24}$ = 34.9 (c = 1.0 in $CHCl_3$/MeOH, 1:1); $^1$H-NMR (600 MHz, $CDCl_3$): $\delta$ = 9.92 (s, 1H), 7.61 (d, *J* = 7.8 Hz, 2H), 7.36 (d, *J* = 8.1 Hz, 1H), 7.25 – 7.06 (m, 6H), 6.95 (d, *J* = 2.3 Hz, 2H), 6.91 – 6.74 (m, 3H), 6.65 (d, *J* = 6.5 Hz, 1H), 5.66 (dd, *J* = 12.1, 4.8 Hz, 1H), 5.42 – 5.07 (m, 2H), 4.90 – 4.66 (m, 4H), 4.59 (s, 1H), 3.32 (d, *J* = 4.9 Hz, 2H), 3.10 – 2.97 (m, 1H), 2.91 (s, 3H), 2.84 (s, 2H), 2.69 (dd, *J* = 14.8, 4.8 Hz, 1H), 2.61 (dd, *J* = 14.7, 5.6 Hz, 2H), 2.50 – 2.36 (m, 2H), 2.26 – 2.15 (m, 1H), 1.92 – 1.78 (m, 3H), 1.58 (d, *J* = 1.3 Hz, 3H), 1.52 (s, 11H), 1.50 – 1.39 (m, 6H), 1.28 – 1.14 (m, 18H), 1.09 (dd, *J* = 7.5, 3.8 Hz, 29H), 0.93 – 0.72 (m, 17H); $^{13}$C-NMR (126 MHz, $CDCl_3$): $\delta$ = 175.0, 173.4, 170.6, 169.2, 156.7, 155.5, 136.5, 133.8, 132.3, 127.8, 127.2, 127.2, 127.2, 121.7, 121.6, 119.9, 119.9, 119.1, 118.3, 111.5, 79.7, 70.3, 56.4, 49.9, 48.8, 43.7, 40.8, 40.5, 40.3, 39.7, 30.8, 30.2, 29.7, 29.2, 28.5, 22.4, 22.0, 21.1, 20.9, 19.0, 18.6, 17.9, 12.6; IR: $\tilde{\nu}_{max}$ = 3053, 2933, 2867, 2727,

2619, 2454, 2344, 2226, 2110, 2037, 1910, 1831, 1638, 1510, 1457, 1412, 1365, 1264, 1170, 1101, 1012, 912, 835, 740, 685; HRMS (ESI): *m/Z* calcd for $C_{53}H_{82}N_5O_8Si$ $[M+H]^+$: 944.5927, found: 944.5932.

*cyclo*-[(2*S*,4*Z*,6*R*,8*S*)Htn-L-Lys(Boc)-D-*N*-Me-Trp-L-β-Tyr(TIPS)]; ()

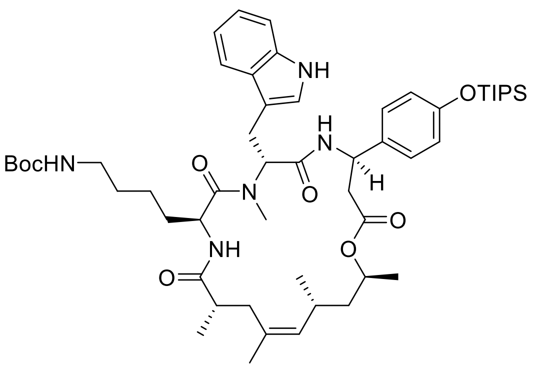

$R_f$ = 0.25 (petroleum ether/EtOAc, 1:1); $[\alpha]_D^{24}$ = 14.6 (c = 1.0 in $CHCl_3$/MeOH, 1:1); $^1$H-NMR (600 MHz, $CDCl_3$): $\delta$ = 9.67 (s, 1H), 7.63 (d, *J* = 7.9 Hz, 1H), 7.37 (d, *J* = 8.0 Hz, 1H), 7.27 (d, *J* = 8.2 Hz, 1H), 7.14 (ddd, *J* = 21.2, 13.6, 7.2 Hz, 4H), 6.99 – 6.95 (m, 1H), 6.87 – 6.77 (m, 2H), 5.66 (dd, *J* = 12.1, 4.8 Hz, 1H), 5.23 (d, *J* = 4.3 Hz, 1H), 5.01 (d, *J* = 9.1 Hz, 1H), 4.81 (d, *J* = 6.2 Hz, 1H), 4.72 (s, 2H), 3.47 – 3.35 (m, 1H), 3.27 (dd, *J* = 16.4, 4.8 Hz, 1H), 3.10 – 2.99 (m, 1H), 2.96 (s, 3H), 2.94 – 2.83 (m, 2H), 2.70 (d, *J* = 4.4 Hz, 1H), 2.61 (d, *J* = 2.7 Hz, 4H), 2.04 – 1.78 (m, 3H), 1.65 (d, *J* = 1.4 Hz, 3H), 1.54 (s, 10H), 1.47 (qt, *J* = 14.7, 8.3 Hz, 4H), 1.42 – 1.34 (m, 2H), 1.31 – 1.21 (m, 10H), 1.17 (d, *J* = 6.8 Hz, 5H), 1.12 (d, *J* = 7.5 Hz, 25H), 0.86 (d, *J* = 6.7 Hz, 9H); $^{13}$C-NMR (126 MHz, $CDCl_3$): $\delta$ = 175.0, 174.1, 169.9, 169.3, 156.5, 155.4, 136.5, 133.5, 133.5, 130.7, 127.3, 127.3, 121.7, 121.7, 119.8, 119.1, 118.5, 111.4, 110.4, 79.7, 71.0, 55.9, 49.4, 49.1, 43.9, 40.5, 40.3, 39.2, 35.6, 31.6, 30.5, 30.2, 30.0, 29.7, 28.5, 23.3, 22.3, 21.5, 21.5, 21.3, 19.1, 17.9, 12.7; IR: $\tilde{\nu}_{max}$ = 3061, 2932, 2866, 2652, 2597, 2533, 2225, 2143, 2106, 2007, 1944, 1892, 1828, 1683, 1644, 1510, 1457, 1366, 1266, 1171, 1102, 1011, 914, 836, 740, 683; HRMS (ESI): *m/Z* calcd for $C_{53}H_{82}N_5O_8Si$ $[M+H]^+$: 944.5927, found: 944.5926.

*cyclo*-[(2*S*,4*E*,8*S*)-Hdn-L-Lys(Gly-Boc)-D-*N*MeTrp-L-*β*Tyr(TIPS)]; (**S69**)

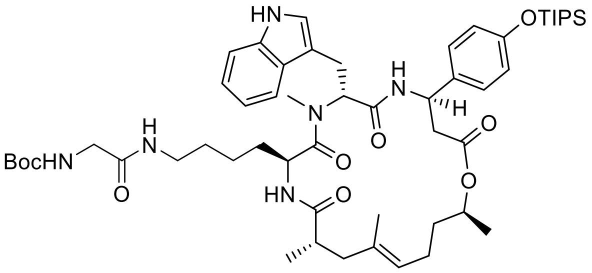

Under an atmosphere of argon, cyclodepsipeptide **S51** (36.6 mg, 39.3 µmol, 1.0 equiv.) was dissolved in dehydrated $CH_2Cl_2$ (2.0 mL) and cooled to 0 °C (ice). TFA (1.0 mL) was added, and the solution was stirred for 1 h at 0 °C. The solution was diluted with toluene (0.5 mL). The solvent was removed under reduced pressure and the residue was dried in fine vacuum. The residue was mixed with Boc-Gly-OH (7.6 mg, 43.2 µmol, 1.1 equiv.) and HATU (35.8 mg, 94.3 µmol, 2.4 equiv.). Under an atmosphere of argon, the mixture was suspended in THF (3 mL) followed by addition of DIPEA (53.4 µL, 314 µmol, 8.0 equiv.). The resulting solution was stirred for 16 h at room temperature. The solvent was removed under reduced pressure and purified by silica-gel chromatography ($CH_2Cl_2$/MeOH, 97:3 → 95:5) to provide cyclopeptide **S69** (29 mg, 75%) as an off-white solid. $R_f$ = 0.31 ($CH_2Cl_2$/MeOH, 19:1); $[\alpha]_D^{24}$ = 29.4 (c = 5.0 in MeCN); $^1$H-NMR (400 MHz, $CDCl_3$): $\delta$ = 9.78 (br s, 1 H), 7.63 (d, $J$ = 7.9 Hz, 1 H), 7.36 (t, $J$ = 7.6 Hz, 2 H), 7.19 - 7.08 (m, 5 H), 6.98 (d, $J$ = 1.8 Hz, 1 H), 6.84 (d, $J$ = 8.8 Hz, 2 H), 6.62 (d, $J$ = 6.4 Hz, 1 H), 6.45 (t, $J$ = 5.7 Hz, 1 H), 5.71 (dd, $J$ = 5.3, 11.4 Hz, 1 H), 5.31 (t, $J$ = 6.0 Hz, 1 H), 5.27 (dt, $J$ = 3.8, 7.9 Hz, 1 H), 5.01 (t, $J$ = 6.7 Hz, 1 H), 4.86 - 4.80 (m, 1 H), 3.86 (d, $J$ = 5.6 Hz, 2 H), 3.38 - 3.18 (m, 3 H), 2.98 (t, $J$ = 5.3 Hz, 1 H), 2.93 (s, 3 H), 2.95 - 2.84 (m, 1 H), 2.79 (dd, $J$ = 4.1, 15.5 Hz, 1 H), 2.60 (dd, $J$ = 7.7, 15.3 Hz, 1 H), 2.49 - 2.42 (m, 2 H), 1.94 - 1.78 (m, 3 H), 1.49 (s, 9 H), 1.48 (s, 3 H), 1.57 - 1.40 (m, 5 H), 1.35 - 1.20 (m, 5 H), 1.11 (d, $J$ = 7.3 Hz, 21 H), 1.07 - 0.97 (m, 2 H), 0.66 - 0.55 (m, 1 H); $^{13}$C-NMR (101 MHz, $CDCl_3$): $\delta$ = 174.7, 173.7, 170.5, 170.1, 169.3, 156.2, 155.5, 136.6, 133.9, 132.9, 127.6, 127.1, 124.5, 121.8, 121.8, 119.9, 119.1, 118.4, 111.6, 110.2, 80.5, 69.6, 56.1, 55.4, 49.5, 49.0, 43.2, 39.8, 39.7, 39.7, 35.6, 31.0, 30.3, 29.5, 28.3, 23.2, 22.6, 21.0, 20.4, 20.3, 17.9, 16.2, 12.7; IR: $\tilde{\nu}_{max}$ = 3310, 2940, 2866, 2361, 1717, 1639, 1508, 1458, 1265, 1169, 841, 741, 679; HRMS (ESI): *m/Z* calcd for $C_{54}H_{83}N_6O_9Si$ $[M+H]^+$: 987.5985, found: 987.5988.

*cyclo*-[(2*S*,4*E*,8*S*)-Hdn-L-Lys(Eaca-Boc)-D-*N*MeTrp-L-*β*Tyr(TIPS)]; (**S70**)

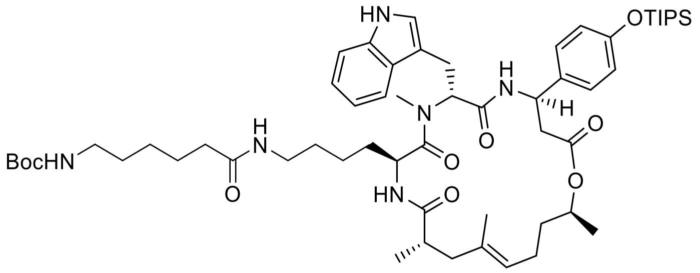

Under an atmosphere of argon, cyclodepsipeptide **S51** (35.0 mg, 37.6 µmol, 1.0 equiv.) was dissolved in dehydrated $CH_2Cl_2$ (2.0 mL) and cooled to 0 °C (ice). TFA (1.0 mL) was added, and the solution was stirred for 1 h at 0 °C. The solution was diluted with toluene (0.5 mL). The solvent was removed under reduced pressure and the residue was dried in fine vacuum. The residue was mixed with oc-6-aminohexanoic acid (9.6 mg, 41.4 µmol, 1.1 equiv.) and HATU (34.3 mg, 90.2 µmol, 2.4 equiv.). Under an atmosphere of argon, the mixture was suspended in THF (3 mL) followed by addition of DIPEA (51.1 µL, 301 µmol, 8.0 equiv.). The resulting solution was stirred for 16 h at room temperature. The solvent was removed under reduced pressure and purified by silica-gel chromatography ($CH_2Cl_2$/MeOH, 97:3 → 95:5) to provide cyclopeptide **S70** (26.7 mg, 68%) as an off-white solid. $R_f$ = 0.21 ($CH_2Cl_2$/MeOH, 19:1); $[\alpha]_D^{24}$ = 32.5 (c = 2.0 in MeCN); $^1$H-NMR (400 MHz, DMSO-$d_6$): $\delta$ = 10.83 (br s, 1 H), 9.31 (s, 1 H), 8.66 (d, $J$ = 8.8 Hz, 1 H), 7.74 - 7.63 (m, 3 H), 7.29 (d, $J$ = 7.9 Hz, 1 H), 7.13 (d, $J$ = 8.5 Hz, 2 H), 7.07 (d, $J$ = 2.0 Hz, 1 H), 7.04 (ddd, $J$ = 1.0, 7.2, 7.9 Hz, 1 H), 6.95 (ddd, $J$ = 0.9, 7.2, 7.9 Hz, 1 H), 6.77 (t, $J$ = 5.7 Hz, 1 H), 6.70 (d, $J$ = 8.5 Hz, 2 H), 5.52 (dd, $J$ = 5.0, 11.1 Hz, 1 H), 5.19 (ddd, $J$ = 2.3, 8.5, 11.4 Hz, 1 H), 4.93 (t, $J$ = 6.6 Hz, 1 H), 4.68 (qd, $J$ = 6.3, 12.8 Hz, 1 H), 4.61 - 4.50 (m, 1 H), 3.04 (s, 3 H), 3.11 - 2.98 (m, 1 H), 2.97 - 2.74 (m, 5 H), 2.73 - 2.63 (m, 2 H), 2.59 (dd, $J$ = 3.1, 14.8 Hz, 1 H), 2.18 (dd, $J$ = 11.7, 14.6 Hz, 1 H), 2.02 (t, $J$ = 7.5 Hz, 2 H), 1.84 (tt, $J$ = 7.1, 13.9 Hz, 2 H), 1.74 (d, $J$ = 14.3 Hz, 1 H), 1.49 (s, 3 H), 1.53 - 1.44 (m, 3 H), 1.37 (s, 9 H), 1.41 - 1.30 (m, 2 H), 1.20 (dd, $J$ = 2.8, 7.2 Hz, 1 H), 1.16 (d, $J$ = 6.1 Hz, 3 H), 1.08 - 1.02 (m, 21 H), 0.93 (d, $J$ = 6.7 Hz, 3 H), 0.87 - 0.78 (m, 3 H), 0.72 - 0.65 (m, 1 H); $^{13}$C-NMR (101 MHz, DMSO-$d_6$): $\delta$ = 174.9, 172.8, 172.4, 170.8, 170.5, 156.7, 156.0, 136.6, 133.6, 133.5, 127.7, 127.6, 127.4, 123.8, 123.4, 121.3, 119.8, 119.1, 118.5, 115.5, 111.6, 110.0, 77.8, 71.4, 55.1, 49.5, 48.1, 43.0, 42.1, 38.5, 38.1, 35.9, 35.3, 31.0, 29.8, 29.5, 29.1, 28.7, 26.5, 26.0, 25.6, 24.2, 22.5, 20.2, 19.9, 18.3, 18.2, 17.4, 12.5; IR: $\tilde{\nu}_{max}$ = 3275, 2932, 2866, 2361, 1639, 1516, 1458, 1366, 1269, 1250, 1173, 1007, 745; HRMS (ESI): *m/Z* calcd for $C_{58}H_{91}N_6O_9Si$ $[M+H]^+$: 1043.661, found: 1043.661.

1-L-Pea-L-Dap(Boc)-D-*N*MeTrp-L-*β*Tyr(TIPS)-*O*-4-penten; (**S71**)

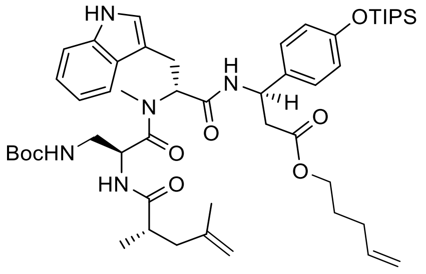

Standard Procedure **SP1** with peptide acid **S50** (100 mg, 120 µmol, 1.0 equiv.), 4-penten-1-ol (73.2 µL, 719 µmol, 6.0 equiv.), DMAP (58.6 mg, 480 µmol, 4.0 equiv.), DIPEA (81.6 µL, 480 µmol, 4.0 equiv.) and EDCI (46.0 mg, 240 µmol, 2.0 equiv.) gave diene **S71** (99 mg, 92%) as a colorless solid after purification by silica-gel chromatography ($CH_2Cl_2$/MeOH, 98:2). $R_f$ = 0.42 ($CH_2Cl_2$/MeOH, 19:1); $[\alpha]_D^{24}$ = 38.7 (c = 2.0 in MeCN); $^1$H-NMR (400 MHz, DMSO-$d_6$): $\delta$ = 10.58 (br s, 1 H), 7.78 (d, $J$ = 5.0 Hz, 1 H), 7.71 (d, $J$ = 8.2 Hz, 1 H), 7.54 (d, $J$ = 7.9 Hz, 1 H), 7.32 (d, $J$ = 8.2 Hz, 1 H), 7.15 (d, $J$ = 8.2 Hz, 2 H), 7.08 - 6.99 (m, 2 H), 6.95 (t, $J$ = 7.0 Hz, 1 H), 6.76 (d, $J$ = 8.2 Hz, 2 H), 6.20 (br s, 1 H), 5.85 - 5.70 (m, 1 H), 5.32 (dd, $J$ = 5.3, 8.8 Hz, 1 H), 5.24 (q, $J$ = 7.6 Hz, 1 H), 5.06 - 4.91 (m, 2 H), 4.72 (br s, 1 H), 4.66 (br s, 1 H), 4.69 - 4.57 (m, 1 H), 3.97 (t, $J$ = 6.1 Hz, 2 H), 3.35 (dd, $J$ = 5.6, 15.2 Hz, 1 H), 3.22 - 3.13 (m, 1 H), 2.97 (s, 3 H), 3.03 - 2.85 (m, 2 H), 2.84 - 2.73 (m, 2 H), 2.27 (dd, $J$ = 5.7, 13.9 Hz, 1 H), 2.09 - 1.96 (m, 2 H), 1.92 (dd, $J$ = 8.6, 13.9 Hz, 1 H), 1.66 (s, 3 H), 1.62 - 1.48 (m, 2 H), 1.36 (s, 9 H), 1.30 - 1.14 (m, 4 H), 1.07 (d, $J$ = 7.3 Hz, 18 H), 1.00 (d, $J$ = 6.7 Hz, 3 H); $^{13}$C-NMR (126 MHz, DMSO-$d_6$): $\delta$ = 176.5, 171.4, 170.6, 169.1, 154.9, 143.4, 138.1, 136.7, 134.7, 128.1, 127.6, 123.4, 121.3, 119.6, 118.7, 118.6, 115.5, 114.9, 112.2, 111.7, 110.7, 78.7, 63.8, 60.7, 57.2, 50.3, 49.5, 41.9, 41.0, 37.9, 31.3, 29.8, 28.6, 27.8, 24.0, 22.5, 18.2, 17.3, 12.6; IR: $\tilde{\nu}_{max}$ = 3310, 2943, 2866, 2361, 1743, 1643, 1508, 1458, 1366, 1265, 1169, 1103, 995, 914, 883, 837, 741, 683; HRMS (ESI): *m/Z* calcd for $C_{50}H_{76}N_5O_8Si$ $[M+H]^+$: 902.5458, found: 902.5470.

1-L-Pea-L-Lys(Boc)-D-*N*MeTrp-L-*β*Tyr(TIPS)-*O*-4-penten; (**S72**)

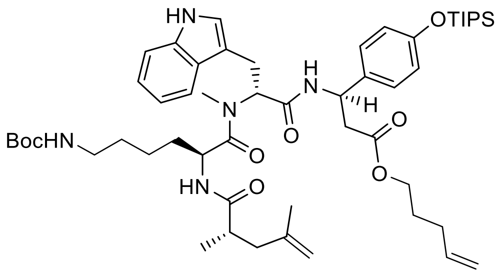

Standard Procedure **SP1** with peptide acid **S49** (80.0 mg, 91.3 µmol, 1.0 equiv.), 4-penten-1-ol (74.4 µL, 730 µmol, 8.0 equiv.), DMAP (44.6 mg, 365 µmol, 4.0 equiv.), DIPEA (62.1 µL,

365 µmol, 4.0 equiv.) and EDCI (35.0 mg, 183 µmol, 2.0 equiv.) gave diene **S72** (82 mg, 95%) as a colorless solid after purification by silica-gel chromatography ($CH_2Cl_2$/MeOH, 98:2). $R_f$ = 0.32 ($CH_2Cl_2$/MeOH, 19:1); $[\alpha]_D^{24}$ = 22.4 (c = 2.0 in MeCN); $^1$H-NMR (300 MHz, $CDCl_3$): $\delta$ = 9.59 (br s, 1 H), 7.61 (d, $J$ = 7.6 Hz, 1 H), 7.34 (d, $J$ = 7.8 Hz, 1 H), 7.22 (d, $J$ = 8.3 Hz, 1 H), 7.16 (d, $J$ = 8.6 Hz, 2 H), 7.19 - 7.12 (m, 1 H), 7.12 - 7.06 (m, 1 H), 6.98 (d, $J$ = 1.5 Hz, 1 H), 6.81 (d, $J$ = 8.5 Hz, 2 H), 6.26 (d, $J$ = 6.1 Hz, 1 H), 5.87 - 5.70 (m, 1 H), 5.40 (q, $J$ = 7.4 Hz, 1 H), 5.11 - 4.94 (m, 3 H), 4.76 (s, 1 H), 4.69 (s, 1 H), 4.63 - 4.53 (m, 1 H), 4.00 (t, $J$ = 6.6 Hz, 1 H), 3.68 (t, $J$ = 6.5 Hz, 1 H), 3.44 (dd, $J$ = 5.3, 15.7 Hz, 1 H), 3.30 (dd, $J$ = 11.9, 16.3 Hz, 1 H), 2.95 (s, 3 H), 2.99 - 2.86 (m, 2 H), 2.84 - 2.72 (m, 1 H), 2.46 - 2.30 (m, 2 H), 2.16 (q, $J$ = 7.7 Hz, 1 H), 2.09 - 1.97 (m, 3 H), 1.68 (s, 3 H), 1.74 - 1.58 (m, 2 H), 1.53 (s, 9 H), 1.46 - 1.36 (m, 1 H), 1.33 - 1.17 (m, 6 H), 1.09 (d, $J$ = 6.9 Hz, 21 H), 0.89 - 0.76 (m, 2 H); $^{13}$C-NMR (101 MHz, $CDCl_3$): $\delta$ = 176.1, 173.3, 170.9, 169.2, 156.5, 155.4, 142.9, 138.3, 137.4, 133.0, 127.5, 127.3, 121.8, 121.8, 119.9, 119.1, 118.5, 115.3, 112.4, 111.4, 110.5, 79.8, 64.0, 62.5, 56.6, 49.7, 49.4, 41.8, 40.7, 40.3, 38.8, 31.8, 30.7, 30.0, 29.9, 28.5, 27.7, 23.0, 22.3, 21.5, 17.9, 17.3, 12.6; IR: $\tilde{v}_{max}$ = 3302, 2943, 2866, 2361, 1735, 1643, 1512, 1458, 1269, 1169, 995, 914, 883, 740, 683; HRMS (ESI): *m/Z* calcd for $C_{53}H_{82}N_5O_8Si$ $[M+H]^+$: 944.5927, found: 944.5938.

(2*S*,4*R*)-2-[L-Pea-L-Lys(Boc)-D-*N*-Me-Trp-L-β-Tyr(TIPS)-*O*-]-4-Me-hex-5-en; ()

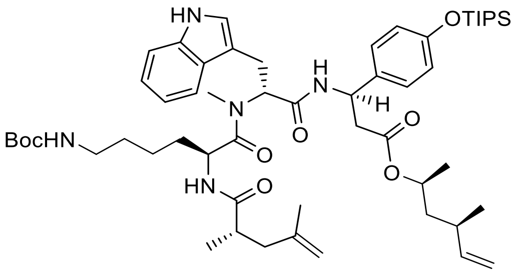

Peptide acid **S49** (280.0 mg, 91.3 µmol, 1.0 equiv.) was mixed with MNBA (31.4 mg, 91.3 µmol, 1.0 equiv.) and DIPEA (31 µl, 183 µmol, 2 equiv.) in $CH_2Cl_2$ (7 mL) at 25 °C and stirred for 30 min. DMAP (11 mg, 91.3 µmol, 1 equiv.) was added, followed by the addition of alcohol **S48** (40 mg, 0.34 mmol, 1.3 equiv.). The mixture was heated to reflux for 16 h, cooled to 25 °C, diluted with $CH_2Cl_2$ (30 mL), and washed with water (10 mL). The organic layer was dried with $Na_2SO_4$, evaporated under reduced pressure, and purified by silica-gel chromatography (petroleum ether/EtOAc 1:1) to provide cyclopeptide **S73** (234 mg, 75%) as a yellow solid. $R_f$ = 0.42 (petroleum ether/EtOAc 1:1); $[\alpha]_D^{24}$ = 19.0 (c = 1.0 in $CHCl_3$/MeOH, 1:1); $^1$H-NMR (400 MHz, $CDCl_3$): $\delta$ = 9.58 (s, 1H), 7.60 (d, $J$ = 7.7 Hz, 1H), 7.33 (d, $J$ = 8.0 Hz,

1H), 7.23 (d, *J* = 8.5 Hz, 1H), 7.18 – 7.05 (m, 4H), 6.96 (d, *J* = 2.3 Hz, 1H), 6.79 (d, *J* = 8.6 Hz, 2H), 6.26 (d, *J* = 6.2 Hz, 1H), 5.74 – 5.57 (m, 2H), 5.38 (d, *J* = 7.7 Hz, 1H), 5.03 – 4.79 (m, 3H), 4.75 (t, *J* = 1.9 Hz, 1H), 4.72 – 4.61 (m, 2H), 4.56 (q, *J* = 6.2 Hz, 1H), 3.41 (d, *J* = 4.7 Hz, 1H), 3.30 (d, *J* = 12.2 Hz, 1H), 2.94 (s, 6H), 2.76 (d, *J* = 6.2 Hz, 1H), 2.49 – 2.23 (m, 2H), 2.03 (ddd, *J* = 16.6, 13.8, 7.5 Hz, 2H), 1.66 (d, *J* = 1.3 Hz, 3H), 1.51 (s, 8H), 1.46 – 1.34 (m, 2H), 1.22 (d, *J* = 7.8 Hz, 6H), 1.19 – 1.00 (m, 24H), 0.90 (d, *J* = 6.7 Hz, 3H), 0.81 (qt, *J* = 9.6, 4.7 Hz, 2H); $^{13}$C-NMR (101 MHz, $CDCl_3$): $\delta$ = 176.1, 173.3, 170.9, 169.2, 156.5, 155.4, 142.9, 138.3, 137.4, 133.0, 127.5, 127.3, 121.8, 121.8, 119.9, 119.1, 118.5, 115.3, 112.4, 111.4, 110.5, 79.8, 64.0, 62.5, 56.6, 49.7, 49.4, 41.8, 40.7, 40.3, 38.8, 31.8, 30.7, 30.0, 29.9, 28.5, 27.7, 23.0, 22.3, 21.5, 17.9, 17.3, 12.6; IR: $\tilde{\nu}_{max}$ = 2943, 2867, 2250, 1791, 1719, 1640, 1609, 1532, 1510, 1266, 1170, 1124, 1104, 1070, 1011, 995, 908, 883, 836, 812, 785, 73, 675, 646; HRMS (ESI): *m/Z* calcd for $C_{55}H_{86}N_5O_8Si$ [M+H]$^+$: 972.6240, found: 972.6245.

**Cyclopeptide analogs:**

*cyclo*-[(2*S*,4*E*,8*S*)Hdn-L-Dap(Boc)-D-*N*-Me-Trp-L-β-Tyr(TIPS)-NH]; )

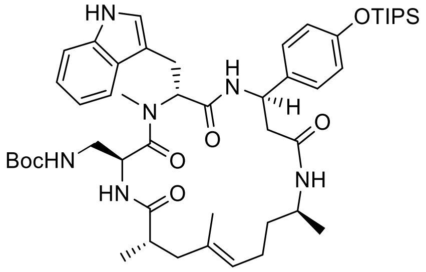

Standard Procedure **SP2** with diene **S74** (77.0 mg, 85.4 µmol, 1.0 equiv.) and 2nd generation Grubbs catalyst (8.7 mg, 11.0 µmol, 0.12 equiv) gave cyclodepsipeptide **S61** (56 mg, 74%) as a colorless solid after purification by silica-gel chromatography. $R_f$ = 0.18 (EtOAc); $[\alpha]_D^{24}$ = 28.4 (c = 1.0 in $CHCl_3$/MeOH, 1:1); $^1$H-NMR (400 MHz, $CDCl_3$): $\delta$ = 8.26 (s, 1 H), 8.12 (d, *J* = 8.1 Hz, 1 H), 7.63 (d, *J* = 7.9 Hz, 1 H), 7.34 (s, 1 H), 7.25 – 6.95 (m, 6 H), 6.92 – 6.75 (m, 3 H), 5.88 – 5.73 (m, 1 H), 5.20 (dt, *J* = 20.3, 7.5 Hz, 2 H), 5.05 (s, 1 H), 4.91 (d, *J* = 5.4 Hz, 1 H), 4.60 (s, 1 H), 3.86 (s, 1 H), 3.51 (d, *J* = 5.1 Hz, 1 H), 3.42 – 3.10 (m, 3 H), 2.97 (s, 3 H), 2.62 (dd, *J* = 14.1, 4.5 Hz, 2 H), 2.57 – 2.30 (m, 4 H), 2.14 – 1.76 (m, 4 H), 1.76 – 1.60 (m, 1 H), 1.54 (s, 3 H), 1.43 (s, 11 H), 1.33 – 1.15 (m, 11 H), 1.11 (d, *J* = 7.3 Hz, 21 H), 1.01 (d, *J* = 6.6 Hz, 3 H); $^{13}$C-NMR (101 MHz, $CDCl_3$): $\delta$ = 175.8, 171.54, 169.38, 168.83, 156, 155.3, 136.3, 133.7, 133.2, 127.0, 126.9, 124.8, 122.3, 122.2, 119.9, 119.5, 118.7, 111.3, 111.9, 79.2, 77.2, 56.6, 50.4, 49.8, 43.5, 43.3, 42.8, 42.4, 39.8, 36.7, 30.6, 28.4, 23.5, 23.2, 21.1, 19.8, 17.9,

16.4, 12.6; IR: $\tilde{\nu}_{max}$ = 2938, 2633, 2608, 2247, 2206, 2183, 2147, 2122, 2009, 1978, 1942, 1918, 1509, 1457, 1389, 1365, 1260, 1168, 1097, 1012, 912, 884, 835, 782, 739, 683; HRMS (ESI): *m/Z* calcd for $C_{49}H_{75}N_6O_7Si$ $[M+H]^+$: 887.5461, found: 887.5460.

*cyclo*-[(2*S*,4*E*,8*S*)Hdn-L-Dap(Boc)-D-*N*-Me-Trp-L-β-Tyr(TIPS)-N(Me)]; ()

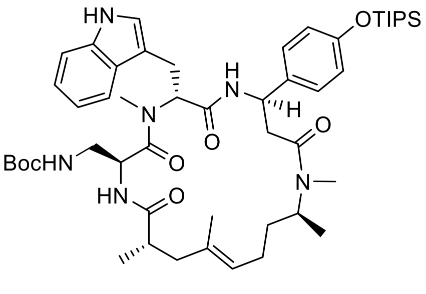

Standard Procedure **SP2** with diene **S75** (70.0 mg, 77.7 µmol, 1.0 equiv.) and 2nd generation Grubbs catalyst (7.9 mg, 11.0 µmol, 0.12 equiv) gave cyclodepsipeptide **S62** (42 mg, 62%) as a colorless solid after purification by silica-gel chromatography. $R_f$ = 0.24 (EtOAc); $[\alpha]_D^{24}$ = 14.6 (c = 0.5 in $CHCl_3$/MeOH, 1:1); $^1$H-NMR (400 MHz, $CDCl_3$): $\delta$ = 8.42 (s, 1 H), 7.98 (d, *J* = 8.0 Hz, 1 H), 7.62 (d, *J* = 7.7 Hz, 1 H), 7.33 (d, *J* = 8.0 Hz, 1 H), 7.21 – 7.05 (m, 5 H), 6.97 (s, 2 H), 6.81 (d, *J* = 7.9 Hz, 3 H), 5.87 – 5.55 (m, 1 H), 5.21 (s, 1 H), 5.06 (s, 1 H), 4.98 – 4.72 (m, 2 H), 4.59 (d, *J* = 7.1 Hz, 1 H), 3.55 – 3.08 (m, 4 H), 2.92 (s, 4 H), 2.82 (s, 3 H), 2.71 – 2.42 (m, 7 H), 1.94 – 1.58 (m, 5 H), 1.51 (s, 4 H), 1.43 (d, *J* = 8.4 Hz, 13 H), 1.25 (d, *J* = 7.5 Hz, 8 H), 1.10 (t, *J* = 7.2 Hz, 28 H), 1.03 – 0.65 (m, 6H); $^{13}$C-NMR (101 MHz, $CDCl_3$): $\delta$ = 176.1, 171.7, 170.0, 168.8, 155.9, 155.4, 136.3, 134.0, 133.6, 127.1, 127.0, 126.9, 125.1, 122.2, 119.9, 119.6, 119.4, 118.6, 111.4, 111.3, 110.9, 79.1, 77.3, 64.4, 56.2, 50.4, 49.2, 45.7, 43.5, 43.0, 39.9, 39.4, 38.6, 32.6, 30.3, 29.7, 28.7, 28.4, 23.7, 22.9, 20.2, 18.1, 17.9, 16.4, 12.6; IR: $\tilde{\nu}_{max}$ = 2928, 2865, 2721, 2694, 2626, 2591, 2562, 2532, 2227, 2160, 2037, 1991, 1919, 1828, 1683, 1649, 1510, 1458, 1418, 1339, 1056, 917, 796, 740, 687, 614; HRMS (ESI): *m/Z* calcd for $C_{50}H_{77}N_6O_7Si$ $[M+H]^+$: 901.5618, found: 901.5635.

(2*S*)-2-[L-Pea-L-Dap(Boc)-D-*N*-Me-Trp-L-β-Tyr(TIPS)-NH-]-hex-5-en; ()

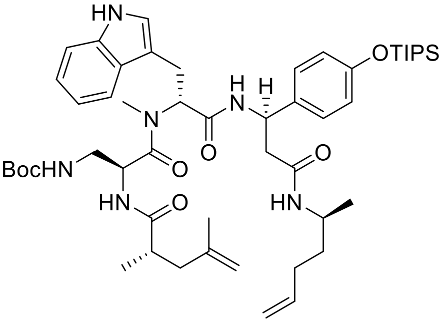

Boc-protected amine **S76** (30 mg, 0.15 mmol, 1.5 equiv.) was dissolved in $CH_2Cl_2$ (5 mL) at 0 °C and TFA (2 mL) was added dropwise. The reaction mixture was stirred for 1 h, at the same temperature. The volatiles were removed under reduced pressure at 25 °C. Tripeptide **S50** (83.4 mg, 0.1 mmol, 1.0 equiv.) was dissolved in DMF (10 mL), and HATU (38 mg, 0.1 mmol, 1.0 equiv.), and DIPEA (70.0 µL, 0.4 mmol, 4.0 equiv.) were added at 25 °C. After gentle stirring for 5 min, this solution was added to previously deprotected carbamate **S76**. The reaction mixture was stirred for 16 h at the same temperature. The mixture was diluted with EtOAc (50 mL), washed with brine (2 x 20 mL), and dried with $Na_2SO_4$. The mixture was concentrated under reduced pressure and purified by silica-gel chromatography (petroleum ether/EtOAc 2:8), which provided peptide diene **S74** as a white solid (81 mg, 88%). $R_f$ = 0.53 (EtOAc); $[\alpha]_D^{24}$ = 22.9 (c = 1.0 in $CHCl_3$/MeOH, 1:1); $^1$H-NMR (300 MHz, $CDCl_3$): $\delta$ = 8.77 (d, $J$ = 19.0 Hz, 1 H), 7.60 – 7.28 (m, 3 H), 7.22 – 6.83 (m, 5 H), 6.77 (dd, $J$ = 8.6, 6.8 Hz, 2 H), 6.38 (t, $J$ = 5.9 Hz, 1 H), 5.83 – 5.58 (m, 2 H), 5.45 – 5.23 (m, 1 H), 5.14 – 4.85 (m, 2 H), 4.84 – 4.56 (m, 3 H), 4.56 – 4.19 (m, 1 H), 3.91 (dh, $J$ = 12.1, 6.0, 5.4 Hz, 1 H), 3.56 (ddd, $J$ = 32.4, 15.6, 5.0 Hz, 1 H), 3.30 – 3.12 (m, 1 H), 3.07 (dt, $J$ = 14.2, 5.7 Hz, 1 H), 2.96 – 2.76 (m, 4 H), 2.75 – 2.21 (m, 4 H), 2.11 – 1.78 (m, 3 H), 1.64 (d, $J$ = 6.5 Hz, 3 H), 1.51 – 1.32 (m, 11 H), 1.32 – 1.14 (m, 5 H), 1.08 (dd, $J$ = 6.8, 2.8 Hz, 24 H), 0.98 – 0.79 (m, 2 H); $^{13}$C-NMR (75 MHz, $CDCl_3$): $\delta$ = 177.5, 174.9, 171.2, 169.8, 169.3, 169.2, 168.3, 157.1, 156.8, 155.4, 155.2, 143.5, 142.6, 137.8, 136.2, 136.1, 134.1, 133.6, 127.7, 127.1, 126.8, 123.2, 122.4, 122.1, 121.9, 119.8, 119.2, 118.8, 114.9, 114.9, 112.3, 112.1, 111.3, 111.1, 110.3, 80.3, 79.8, 56.8, 51.2, 50.7, 44.0, 44.2, 41.7, 40.8, 38.5, 35.8, 31.2, 30.3, 30.2, 30.1, 29.7, 28.3, 28.3, 23.2, 22.4, 22.2, 20.6, 20.4, 17.9, 17.4, 17.4, 12.6; IR: $\tilde{\nu}_{max}$ = 2964, 2733, 2680, 2633, 2599, 2561, 2234, 2205, 1973, 1864, 1824, 1637, 1510, 1366, 1265, 1171, 1099, 1074, 997, 912, 884, 836, 781, 739, 680; HRMS (ESI): *m/Z* calcd for $C_{51}H_{79}N_6O_7Si$ $[M+H]^+$: 915.5774, found: 915.5779.

(2*S*)-2-[L-Pea-L-Dap(Boc)-D-*N*-Me-Trp-L-β-Tyr(TIPS)-N(Me)-]-hex-5-en; ()

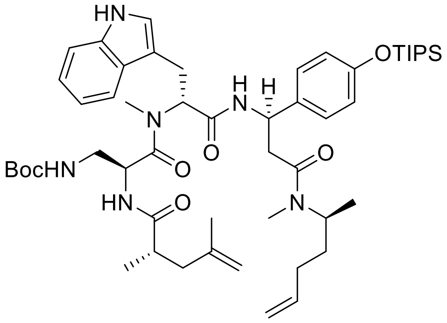

Boc-protected amine **S77** (30 mg, 0.14 mmol, 1.3 equiv.) was dissolved in $CH_2Cl_2$ (5 mL) at 0 °C. TFA (2 mL) was added dropwise. The reaction mixture was stirred for 1 hour at the same temperature, and then the volatiles were removed under reduced pressure at 25 °C. Tripeptide **S50** (91 mg, 0.109 mmol, 1.0 equiv.) was dissolved in DMF (10 mL) and HATU (41 mg, 0.109 mmol, 1.0 equiv.), HOAt (15 mg, 0.109 mmol, 1.0 equiv.) and DIPEA (72 µL, 0.41 mmol, 4.0 equiv.) were added at 25 °C. After gentle stirring for 5 min, this solution was added to previously deprotected carbamate **S77**. The reaction mixture was stirred for 16 h at the same temperature. The mixture was diluted with EtOAc (50 mL), washed with brine (2 x 20 mL), and dried with $Na_2SO_4$. The mixture was concentrated under reduced pressure and purified by silica-gel chromatography (petroleum ether/EtOAc 2:8), which provided peptide diene **S75** as a light brown solid (72 mg, 71%). $R_f$ = 0.70 (EtOAc); $[\alpha]_D^{24}$ = 19.3 (c = 0.5 in $CHCl_3$/MeOH, 1:1); $^1$H-NMR (400 MHz, $CDCl_3$): $\delta$ = 8.59 (s, 1 H), 8.01 (s, 1 H), 7.75 (t, *J* = 8.1 Hz, 1 H), 7.59 (d, *J* = 7.8 Hz, 1 H), 7.52 – 7.37 (m, 1 H), 7.28 (s, 2 H), 7.10 (d, *J* = 20.1 Hz, 5 H), 6.96 (d, *J* = 7.1 Hz, 1 H), 6.87 – 6.72 (m, 2 H), 5.86 – 5.56 (m, 2 H), 5.54 – 5.18 (m, 1 H), 4.95 (d, *J* = 1.6 Hz, 3 H), 4.71 (d, *J* = 26.6 Hz, 5 H), 3.58 (d, *J* = 77.5 Hz, 2 H), 3.21 (dddd, *J* = 31.1, 24.7, 16.5, 11.6 Hz, 3 H), 3.09 – 2.96 (m, 1 H), 2.94 (d, *J* = 3.7 Hz, 2 H), 2.88 (d, *J* = 5.8 Hz, 4 H), 2.81 (s, 3 H), 2.65 (d, *J* = 10.7 Hz, 3 H), 2.57 (s, 2 H), 2.42 (s, 2 H), 2.33 – 2.08 (m, 1 H), 1.95 (dd, *J* = 43.7, 6.7 Hz, 3 H), 1.67 (d, *J* = 4.5 Hz, 4 H), 1.41 (s, 12 H), 1.31 – 1.17 (m, 5 H), 1.09 (dd, *J* = 7.2, 3.2 Hz, 24 H), 0.99 (d, *J* = 6.7 Hz, 5 H); $^{13}$C-NMR (101 MHz, $CDCl_3$): $\delta$ = 176.7, 176.5, 174.8, 171.3, 171.1, 170.6, 170.1, 168.9, 168.8, 167.8, 162.6, 156.6, 155.2, 143.6, 143.1, 143.0, 137.9, 137.4, 137.3, 136.3, 136.1, 134.0, 127.8, 127.4, 123.2, 122.5, 122.1, 122.0, 119.8, 119.3, 118.8, 118.2, 115.6, 114.8, 112.2, 112.1, 111.3, 111.2, 80.0, 79.6, 77.4, 62.4, 57.1, 50.1, 50.0, 47.9, 42.0, 41.7, 38.8, 38.7, 38.6, 36.5, 33.4, 32.9, 32.8, 31.5, 30.7, 30.6, 30.5, 28.6, 28.4, 26.2, 23.2, 22.4, 22.3, 18.6, 18.0, 17.8, 17.4, 12.7; IR: $\tilde{\nu}_{max}$ = 2964, 2615, 2574, 2549, 2234, 2182, 2152, 2107, 2074, 1974, 1940, 1901, 1639, 1509, 1457, 1263, 1169, 1125, 1101, 996, 911, 886, 838, 740, 683, 600; HRMS (ESI): *m/Z* calcd for $C_{52}H_{81}N_6O_7Si$ $[M+H]^+$: 929.5931, found: 929.5934.

(2*S*)-Hex-5-en-2'-yl-carbamic acid *t*-Bu-ester ()

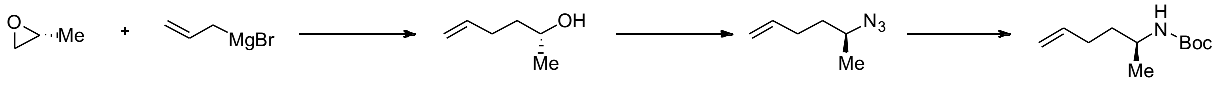

**S78** **S79** **S76**

(*R*)-Hex-5-en-2-ol (**S78**) was synthesized according to the procedure described for (*S*)-hex-5-en-2-ol (**3**). The analytical data were in accordance with previously reported data.[13] Alcohol **S78** (870 mg, 8.7 mmol, 1.0 equiv.) was dissolved in $CH_2Cl_2$ (50 mL), and the reaction mixture was cooled to 0 °C. DPPA (2.1 mL, 9.6 mmol, 1.1 equiv.), $Ph_3P$ (2.5 g, 9.6 mmol, 1.1 equiv.) and DIAD (1.8 mL, 9.6 mmol, 1.1 equiv.) were added, and the reaction mixture was let to warm to 25 °C over 16 h. The reaction mixture was diluted with PE (~30 mL), and the mixture was directly applied to a preequilibrated silica gel flash column (PE/$CH_2Cl_2$ 9:1) to provide (*S*)-5-azidohex-1-ene (**S79**) as a yellow volatile liquid (590 mg, 54% yield).

Azide **S79** (510 mg, 4.08 mmol, 1 equiv.) was dissolved in THF (15 mL), and the solution cooled to 0 °C. Tri-*n*-butylphosphine (1.3 mL, 5.3 mmol, 1.3 equiv.) was added dropwise, over 15 min, while the addition evolution of gas was observed. After the reaction mixture temperature had reached 25 °C (~30 min), $H_2O$ (1.4 mL, 81.6 mmol, 20 equiv.) was added, and the reaction mixture was heated to reflux with stirring, over 16 h. After cooling to 25 °C, 1 M NaOH aqueous solution (5 mL), 1,4-dioxane (5 mL), and $Boc_2O$ (1.3 g, 6.12 mmol, 1.5 equiv.) were consecutively added. The reaction mixture was stirred for 16 h at 25 °C. The organic solvents were evaporated under reduced pressure. The remaining aqueous solution was diluted with water (~50 mL), and acidified with 2 M aqueous solution of $KHSO_4$ to pH 2-3. The mixture was was extracted with $CH_2Cl_2$ (3 x 50 mL), the combined organic extracts were dried with $Na_2SO_4$, and concentrated under reduced pressure. The residue was purified by silica-gel chromatography (petroleum ether/EtOAc 95:5) to provide the carbamate **S76** (64% yield, 516 mg) as a colorless oil that solidified at low temperature. Higher yields (55% over 3 steps) were achieved when volatiles were not completely removed, indicating high volatility of alcohol **S78** and azide **S79**. $R_f$ = 0.68 (petroleum ether/EtOAc, 9:1); $[\alpha]_D^{24}$ = -0.48 (c = 1.0 in $CHCl_3$/MeOH, 1:1); $^1$H-NMR (300 MHz, $CDCl_3$): $\delta$ = 5.79 (d, $J$ = 6.7 Hz, 1 H), 5.06 – 4.90 (m, 2 H), 4.34 (s, 1 H), 3.63 (d, $J$ = 6.8 Hz, 1H), 2.06 (s, 2 H), 1.42 (s, 11 H), 1.10 (d, $J$ = 6.6 Hz, 3 H); $^{13}$C-NMR (75 MHz, $CDCl_3$): $\delta$ = 155.0, 137.8, 114.5, 78.6, 45.8, 36.2, 30.0, 28.1, 20.9; IR: $\tilde{\nu}_{max}$ = 3153, 3118, 3032, 2951, 2750, 2700, 2328, 2250, 2189, 2156, 2046, 2021, 1577, 1467, 1118, 1035, 962, 929, 860, 806, 761, 713; HRMS (ESI): *m/Z* calcd for $C_{11}H_{22}NO_2$ $[M+H]^+$: 200.1645, found: 200.1647.

[13] V. Nasufović, F. Küllmer, J. Bößneck, H.-M. Dahse, H. Görls, P. Bellstedt, P. Stallforth, H.-D. Arndt, *Chemistry – A European Journal* **2021**, *27*, 11633-11642.

(*S*)-Hex-5-en-2-yl(methyl)carbamic acid *tert*-butyl ester; ()

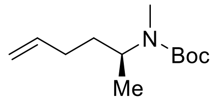

Carbamate **S76** (135 mg, 0.67 mmol, 1.0 equiv.) was dissolved in THF (10 mL) and cooled to 0 °C. NaH (54.0 mg, 1.35 mmol, 2.0 equiv., 60% suspended in mineral oil) was added, and the mixture was stirred at the same temperature for 30 min. MeI (167 µL, 2.7 mmol, 4.0 equiv.) was added and stirred for 16 h, allowing it to reach 25 °C. Water was added (10 mL) and the mixture was acidified with aqueous HCl solution (0.1 M) to pH 2-3. The mixture was diluted with water (~50 mL) and extracted with EtOAc (3 x 50 mL). The organic extracts were combined, dried with $Na_2SO_4$, and concentrated under reduced pressure. The residue was purified by silica-gel chromatography (petroleum ether/EtOAc 95:5) to provide carbamate **S77** as a viscous yellow oil (121 mg, 85% yield). $R_f$ = 0.75 (petroleoum ether/EtOAc, 9:1); $[\alpha]_D^{24}$ = 13.0 (c = 0.1 in $CHCl_3$/MeOH, 1:1); $^1$H-NMR (400 MHz, $CDCl_3$): $\delta$ = 5.91 – 5.67 (m, 1 H), 5.06 – 4.82 (m, 2 H), 4.16 (d, $J$ = 77.9 Hz, 1 H), 2.74 – 2.58 (m, 3 H), 1.98 (t, $J$ = 6.9 Hz, 2 H), 1.61 – 1.52 (m, 1 H), 1.45 (s, 11 H), 1.25 (s, 2 H), 1.07 (d, $J$ = 6.8 Hz, 3 H), 0.87 (s, 2 H); $^{13}$C-NMR (101 MHz, $CDCl_3$): $\delta$ = 156.0, 138.3, 114.8, 79.2, 50.5, 49.5, 33.5, 30.8, 29.8, 28.6, 27.2, 18.6, 18.2; IR: $\tilde{\nu}_{max}$ = 2982, 2582, 2272, 2208, 2182, 2151, 2129, 2102, 2044, 2008, 1966, 1932, 1908, 1693, 1482, 1428, 1393, 1186, 1145, 1119, 1029, 890, 804, 765, 717, 681; HRMS (ESI): *m/Z* calcd for $C_{12}H_{24}NO_2$ $[M+H]^+$: 214.1802, found: 214.1804.

**Synthesis of azobenzenes:**

2-(4´-((4´´-(4´´´-Methoxybenzyl)-3´´,4´´-dihydro-2´´*H*-benzo[*b*][1´´,4´´]oxazin-7´´-yl)-diazenyl)phenoxy)acetic acid; (**S55**)

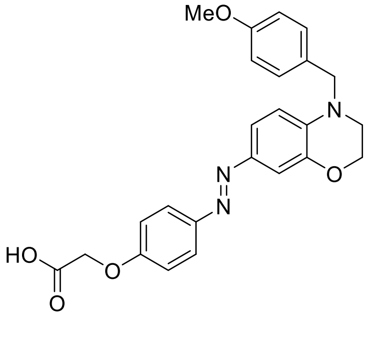

Standard Procedure **SP6** with azobenzene **S80** (8.0 mg, 0.02 mmol, 1.0 equiv.) gave acid **S55** (8.0 mg, quant.) as a red solid which was used for the next step without further purification. LRMS: *m/Z* (%) = 434.5 (100) $[M+H]^+$.

2-(4´-((4´´-(4´´´-chlorbenzyl)-3´´,4´´-dihydro-2´´*H*-benzo[*b*][1´´,4´´]oxazin-7´´-yl)-diazenyl)phenoxy)acetic acid; (**S56**)

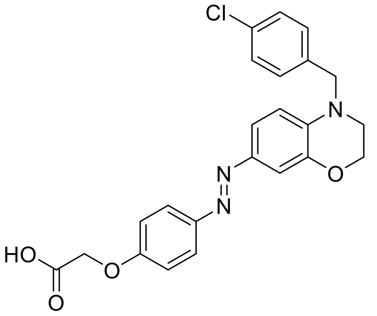

Standard Procedure **SP6** with azobenzene **S81** (10.0 mg, 0.02 mmol, 1.0 equiv.) gave acid **S56** (10 mg, quant.) as an orange solid which was used for the next step without further purification. LRMS: *m/Z* (%) = 438.9 (100) [M+H]$^+$.

2-(4´-((4´´-(3´´´,4´´´-dichlorobenzyl)-3´´,4´´-dihydro-2´´*H*-benzo[*b*][1´´,4´´]oxazin-7´´-yl)-diazenyl)phenoxy)acetic acid; (**S57**)

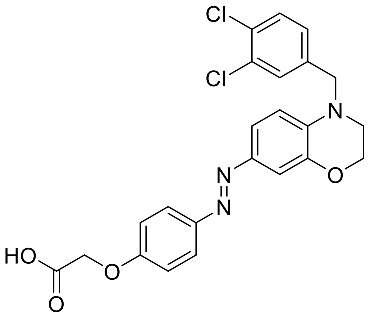

Standard Procedure **SP6** with azobenzene **S82** (50.0 mg, 0.10 mmol, 1.0 equiv.) gave acid **S57** (44 mg, 94) as a yellow solid which was used for the next step without further purification. LRMS: *m/Z* (%) = 473.4 (100) [M+H]$^+$.

2-(4´-((4´´-(4´´´-methylbenzyl)-3´´,4´´-dihydro-2´´*H*-benzo[*b*][1´´,4´´]oxazin-7´´-yl)-diazenyl)phenoxy)acetic acid; (**S58**)

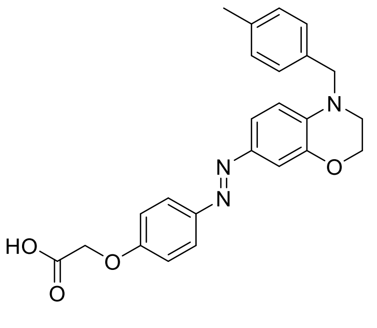

Standard Procedure **SP6** with azobenzene **S83** (12.0 mg, 0.03 mmol, 1.0 equiv.) gave acid **S58** (12 mg, quant.) as an orange solid which was used for the next step without further purification. LRMS: *m/Z* (%) = 418.4 (100) [M+H]⁺.

2-(4´-((4´´-(4´´´-(trifluoromethyl)benzyl)-3´´,4´´-dihydro-2´´*H*-benzo[*b*][1´´,4´´]-oxazin-7´´-yl)diazenyl)phenoxy)acetic acid; (**S59**)

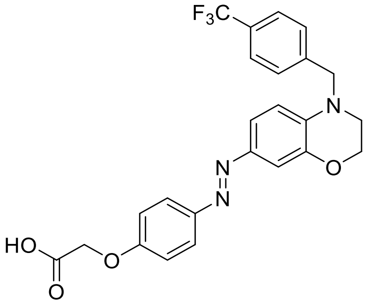

Standard Procedure **SP6** with azobenzene **S84** (30.0 mg, 0.06 mmol, 1.0 equiv.) gave acid **S59** (28 mg, 99%) as an orange solid which was used for the next step without further purification. LRMS: *m/Z* (%) = 472.5 (100) [M+H]⁺.

2-(4´-((4´´-(4´´´-(*tert*-butyl)benzyl)-3´´,4´´-dihydro-2´´*H*-benzo[*b*][1´´,4´´]oxazin-7´´-yl)-diazenyl)-phenoxy)acetic acid; (**S60**)

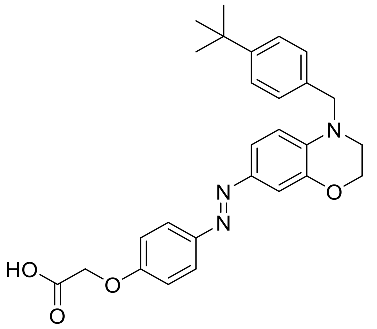

Standard Procedure **SP6** with azobenzene **S85** (41.0 mg, 0.08 mmol, 1.0 equiv.) gave acid **S60** (25 mg, 65%.) as a red solid which was used for the next step without further purification. LRMS: *m/Z* (%) = 460.5 (100) [M+H]⁺.

2-(4´-((4´´-phenyl-3´´,4´´-dihydro-2´´*H*-benzo[*b*][1´´,4´´]oxazin-7´´-yl)diazenyl)-phenoxy)acetic acid; (**S61**)

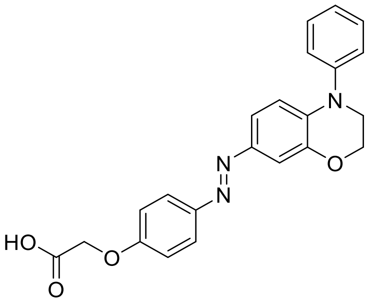

Standard Procedure **SP6** with azobenzene **S86** (30.0 mg, 0.07 mmol, 1.0 equiv.) gave acid **S61** (27.0 mg, 97%) as a black solid which was used for the next step without further purification. LRMS: *m/Z* (%) = 388.5 (100) [M+H]$^+$.

2-(4´-((4´´-phenethyl-3´´,4´´-dihydro-2´´*H*-benzo[*b*][1´´,4´´]oxazin-7´´-yl)diazenyl)-phenoxy)acetic acid; (**S62**)

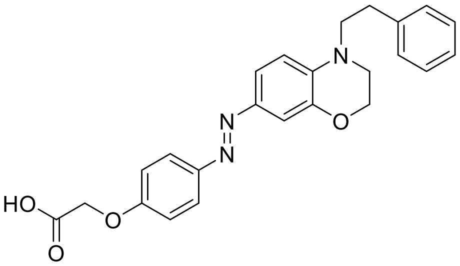

Standard Procedure **SP6** with azobenzene **S87** (12.0 mg, 0.03 mmol, 1.0 equiv.) gave acid **S62** (12 mg, quant.) as an orange solid which was used for the next step without further purification. LRMS: *m/Z* (%) = 418.5 (100) [M+H]$^+$.

ethyl-2-(4´-((4´´-(4´´´-methoxybenzyl)-3´´,4´´-dihydro-2´´*H*-benzo[*b*][1´´,4´´]oxazin-7´´-yl)diazenyl)phenoxy)acetate; (**S80**)

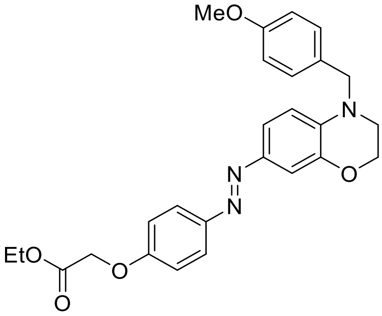

Standard Procedure **SP5** with ethyl-2-(4´-((3´´,4´´-dihydro-2´´*H*-benzo[b][1´´,4´´]oxazin-7´´-yl)diazenyl)phenoxy) acetate (70.0 mg, 0.21 mmol, 1.0 equiv.) and anisaldehyde (75 µl, 0.62 mmol, 3.0 equiv.) gave azobenzene **S80** (10.0 mg, 10%) as a brown solid after purification by silica-gel chromatography (EtOAc/petroleum ether, 1:4 → 1:1). $R_f$ = 0.70 (EtOAc/petroleum ether, 1:2). $^1$H-NMR (250 MHz, $CDCl_3$): $\delta$ = 7.83 (d, $J$ = 9.1 Hz, 2 H), 7.51 - 7.42 (m, 2 H), 7.21 (d, $J$ = 8.8 Hz, 2 H), 7.00 (d, $J$ = 9.0 Hz, 2 H), 6.89 (d, $J$ = 8.6 Hz, 2 H), 6.77 (d, $J$ = 8.2 Hz, 1 H), 4.69 (s, 2 H), 4.52 (s, 2 H), 4.36 - 4.24 (m, 4 H), 3.81 (s, 3 H), 3.50 - 3.42 (m, 2 H), 1.32 (t, $J$ = 7.1 Hz, 3 H); $^{13}$C-NMR (63 MHz, $CDCl_3$): $\delta$ = 168.6, 159.0, 159.0, 148.0, 144.6, 143.8, 138.3, 128.9, 128.2, 123.9, 120.5, 114.8, 114.2, 111.0, 108.4, 65.6, 64.3, 61.5, 55.3, 53.9, 47.0, 14.1; IR: $\tilde{\nu}_{max}$ = 3734, 3630, 2978, 2935, 1600, 1508, 1319, 1246, 1195, 1033; UV-VIS (MeCN + 0.5 % piperidine): $\lambda_{max}$ ($\varepsilon$) = 419 nm (18.4 × $10^3$ l·mol$^{-1}$·cm$^{-1}$); HRMS (ESI): *m/Z* calcd for $C_{26}H_{28}N_3O_5$ [M+H]$^+$: 462.2023, found: 462.2032.

ethyl-2-(4´-((4´´-(4´´´-chlorbenzyl)-3´´,4´´-dihydro-2´´*H*-benzo[*b*][1´´,4´´]oxazin-7´´-yl)-diazenyl)phenoxy)acetate; (**S81**)

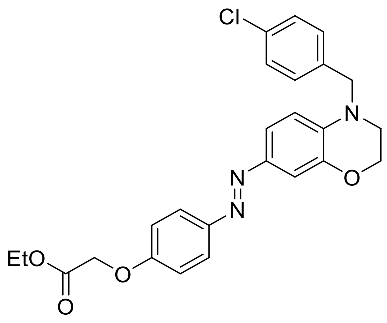

Standard Procedure **SP5** with ethyl-2-(4´-((3´´,4´´-dihydro-2´´*H*-benzo[b][1´´,4´´]oxazin-7´´-yl)diazenyl)phenoxy) acetate (70.0 mg, 0.21 mmol, 1.0 equiv.) and *p*-chlorbenzaldehyde (86.5 mg, 0.62 mmol, 3.0 equiv.) gave azobenzene **S81** (12.0 mg, 13%) as a brown solid after purification by silica-gel chromatography (EtOAc/petroleum ether, 1:4 → 1:1). $R_f$ = 0.70 (EtOAc/petroleum ether, 1:2); $^1$H-NMR (300 MHz, $CDCl_3$): $\delta$ = 7.83 (d, $J$ = 9.0 Hz, 2 H), 7.45 (qd, $J$ = 2.3, 4.6 Hz, 2 H), 7.36 - 7.29 (m, 2 H), 7.25 - 7.18 (m, 2 H), 6.99 (d, $J$ = 9.0 Hz, 2 H), 6.68 (d, $J$ = 9.2 Hz, 1 H), 4.69 (s, 2 H), 4.54 (s, 2 H), 4.35 - 4.25 (m, 4 H), 3.52 - 3.44 (m, 2 H), 1.32 (t, $J$ = 7.1 Hz, 3 H); $^{13}$C-NMR (75 MHz, $CDCl_3$): $\delta$ = 168.6, 159.1, 147.9, 144.8, 143.8, 137.9, 135.6, 133.1, 129.0, 128.2, 124.0, 120.4, 114.8, 111.0, 108.5, 65.5, 64.2, 61.5, 54.0, 47.4, 14.1; IR: $\tilde{\nu}_{max}$ = 3726, 3630, 2978, 2924, 2846, 1597, 1508, 1319, 1250, 1196, 1053, 652; UV-

VIS (MeCN + 0.5 % piperidine): $\lambda_{max}$ ($\varepsilon$) = 415 nm (23.8 × $10^3$ l·mol$^{-1}$·cm$^{-1}$); HRMS (ESI): *m/Z* calcd for $C_{25}H_{25}ClN_3O_4$ $[M+H]^+$: 466.1528, found: 466.1537.

ethyl-2-(4′-((4′′-(3′′′,4′′′-dichlorobenzyl)-3′′,4′′-dihydro-2′′*H*-benzo[*b*][1′′,4′′]oxazin-7′′-yl)diazenyl)phenoxy)acetate; (**S82**)

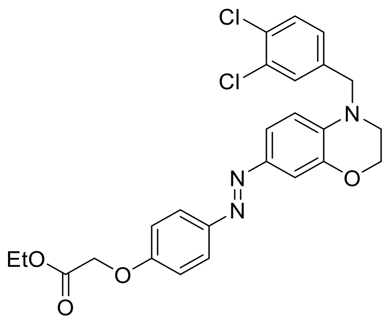

Under a nitrogen atmosphere, ethyl-2-(4′-((3′′,4′′-dihydro-2′′*H*-benzo[b][1′′,4′′]oxazin-7′′-yl)diazenyl)phenoxy) acetate (100 mg, 0.29 mmol, 1.0 equiv.) and $K_2CO_3$ (202 mg, 1.46 mmol, 5.0 equiv.) were dissolved in anhydrous DMF (7.5 mL). 3,4-dichlorobenzyl bromide (44.7 µL, 0.31 mmol, 1.05 equiv.) was added and the mixture was stirred for 6 h at 150 ° C until conversion was complete (TLC check). The solution was diluted with EtOAc (40 mL) and phosphate buffer (30 mL, pH = 7). The organic phase was separated, followed by extraction of the aqueous phase with EtOAc (5 × 15 mL). The combined organic extracts were dried with $Na_2SO_4$, filtered, and concentrated under reduced pressure. Purification by silica-gel chromatography (EtOAc/petroleum ether, 1:4 → 1:1) provided azobenzene **S82** as a brown solid (123 mg, 84%). $R_f$ = 0.70 (EtOAc/petroleum ether, 1:2); $^1$H-NMR (300 MHz, $CDCl_3$): $\delta$ = 7.84 (d, *J* = 9.0 Hz, 2 H), 7.49 - 7.41 (m, 3 H), 7.38 (d, *J* = 2.0 Hz, 1 H), 7.13 (dd, *J* = 2.0, 8.3 Hz, 1 H), 7.00 (d, *J* = 9.0 Hz, 2 H), 6.64 (d, *J* = 9.2 Hz, 1 H), 4.69 (s, 2 H), 4.52 (s, 2 H), 4.36 - 4.25 (m, 4 H), 3.52 - 3.47 (m, 2 H), 1.32 (t, *J* = 7.2 Hz, 3 H); $^{13}$C-NMR (75 MHz, $CDCl_3$): $\delta$ = 168.6, 159.2, 147.8, 145.0, 144.0, 137.7, 137.6, 133.0, 131.4, 130.8, 128.7, 126.1, 124.1, 120.3, 114.8, 111.2, 108.8, 65.5, 64.3, 61.5, 53.9, 47.7, 14.2; ; IR: $\tilde{\nu}_{max}$ = 3726, 3630, 3599, 2970, 2924, 2866, 1508, 1246, 1192, 1053, 1015, 652; UV-VIS (MeCN + 0.5 % piperidine): $\lambda_{max}$ ($\varepsilon$) = 415 nm (12.4 × $10^3$ l·mol$^{-1}$·cm$^{-1}$); HRMS (ESI): *m/Z* calcd for $C_{25}H_{24}Cl_2N_3O_4$ $[M+H]^+$: 500.1138, found: 500.1149.

ethyl-2-(4′-((4′′-(4′′′-methylbenzyl)-3′′,4′′-dihydro-2′′*H*-benzo[*b*][1′′,4′′]oxazin-7′′-yl)-diazenyl)phenoxy)acetate; (**S83**)

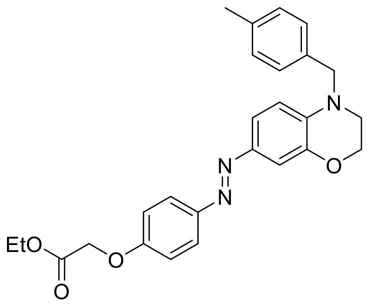

Under a nitrogen atmosphere, ethyl-2-(4´-((3´´,4´´-dihydro-2´´*H*-benzo[b][1´´,4´´]oxazin-7´´-yl)diazenyl)phenoxy) acetate (40.0 mg, 0.12 mmol, 1.0 equiv.) and $K_2CO_3$ (81.0 mg, 0.59 mmol, 5.0 equiv.) were dissolved in anhydrous DMF (3.0 mL). *p*-Methylbenzyl bromide (21.7 mg, 0.12 mmol, 1.0 equiv.) was added and the mixture was stirred for 16 h at 150 ° C until conversion was complete (TLC check). The solution was diluted with EtOAc (20 mL) and phosphate buffer (15 mL, pH = 7). The organic phase was separated, followed by extraction of the aqueous phase with EtOAc (5 × 15 mL). The combined organic extracts were dried with $Na_2SO_4$, filtered, and concentrated under reduced pressure. Purification by silica-gel chromatography (EtOAc/petroleum ether, 1:4 → 1:1) provided azobenzene **S83** as a brown solid (12.0 mg, 22%). $R_f$ = 0.70 (EtOAc/petroleum ether, 1:2); $^1$H-NMR (250 MHz, $CDCl_3$): $\delta$ = 7.83 (d, $J$ = 9.0 Hz, 2 H), 7.51 - 7.42 (m, 2 H), 7.21 - 7.13 (m, 4 H), 7.00 (d, $J$ = 9.1 Hz, 2 H), 6.75 (d, $J$ = 9.3 Hz, 1 H), 4.69 (s, 2 H), 4.54 (s, 2 H), 4.36 - 4.24 (m, 4 H), 3.52 - 3.43 (m, 2 H), 2.35 (s, 3 H), 1.32 (t, $J$ = 7.1 Hz, 3 H); $^{13}$C-NMR (63 MHz, $CDCl_3$): $\delta$ = 168.6, 159.0, 148.0, 144.5, 143.8, 138.3, 137.1, 133.9, 129.5, 126.9, 123.9, 120.5, 114.8, 111.0, 108.4, 65.6, 64.3, 61.5, 54.2, 47.2, 21.1, 14.1; IR: $\tilde{\nu}_{max}$ = 3734, 3630, 2978, 2924, 2870, 1751, 1597, 1508, 1319, 1250, 1196, 1053, 652; UV-VIS (MeCN + 0.5 % piperidine): $\lambda_{max}$ ($\varepsilon$) = 419 nm (17.1 × $10^3$ l·mol$^{-1}$·cm$^{-1}$); HRMS (ESI): *m/Z* calcd for $C_{26}H_{28}N_3O_4$ $[M+H]^+$: 446.2074, found: 446.2089.

ethyl-2-(4´-((4´´-(4´´´-(trifluoromethyl)benzyl)-3´´,4´´-dihydro-2´´*H*-benzo[*b*][1´´,4´´]-oxazin-7´´-yl)diazenyl)phenoxy)acetate; (**S84**)

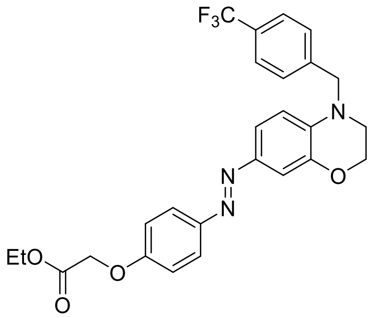

Standard Procedure **SP5** with ethyl-2-(4´-((3´´,4´´-dihydro-2´´*H*-benzo[b][1´´,4´´]oxazin-7´´-yl)diazenyl)phenoxy) acetate (100.0 mg, 0.29 mmol, 1.0 equiv.) and *p*-(trifluoromethyl)benzaldehyde (120 µL, 0.88 mmol, 3.0 equiv.) gave azobenzene **S84** (30.4 mg, 21%) as a yellow solid after purification by silica-gel chromatography (EtOAc/petroleum ether, 1:4 → 1:1). $R_f$ = 0.70 (EtOAc/petroleum ether, 1:2); $^1$H-NMR (300 MHz, $CDCl_3$): $\delta$ = 7.83 (d, $J$ = 9.0 Hz, 2 H), 7.62 (d, $J$ = 8.1 Hz, 2 H), 7.49 - 7.36 (m, 4 H), 7.00 (d, $J$ = 9.0 Hz, 2 H), 6.64 (d, $J$ = 8.4 Hz, 1 H), 4.69 (s, 2 H), 4.62 (s, 2 H), 4.36 - 4.24 (m, 4 H), 3.53 - 3.47 (m, 2 H), 1.31 (t, $J$ = 7.1 Hz, 3 H); $^{13}$C-NMR (101 MHz, $CDCl_3$): $\delta$ = 168.6, 159.2, 147.9, 144.9, 143.9, 141.4, 137.7, 127.0, 125.8, 125.8, 125.8, 125.7, 124.0, 120.3, 114.8, 111.1, 108.7, 65.5, 64.2, 61.5, 54.3, 47.7, 14.1; $^{19}$F-NMR (377 MHz , $CDCl_3$): $\delta$ = -62.5 (s, 3 F); IR: $\tilde{\nu}_{max}$ = 3726, 3630, 2982, 2924, 2847, 1751, 1597, 1508, 1323, 1249, 1199, 1157, 1118; UV-VIS (MeCN + 0.5 % piperidine): $\lambda_{max}$ ($\varepsilon$) = 415 nm (22.8 × $10^3$ l·mol$^{-1}$·cm$^{-1}$); HRMS (ESI): *m/Z* calcd for $C_{26}H_{25}F_3N_3O_4$ [M+H]$^+$: 500.1792, found: 500.1801.

ethyl-2-(4´-((4´´-(4´´´-(*tert*-butyl)benzyl)-3´´,4´´-dihydro-2´´*H*-benzo[*b*][1´´,4´´]oxazin-7´´-yl)diazenyl)phenoxy)acetate; (**S85**)

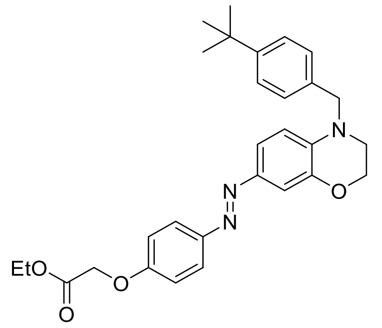

Standard Procedure **SP5** with ethyl-2-(4´-((3´´,4´´-dihydro-2´´*H*-benzo[b][1´´,4´´]oxazin-7´´-yl)diazenyl)phenoxy) acetate (100.0 mg, 0.29 mmol, 1.0 equiv.) and 4-*tert*-butylbenzaldehyde (75 µl, 0.62 mmol, 3.0 equiv.) gave azobenzene **S85** (26.1 mg, 19%) as a brown solid after purification by silica-gel chromatography (EtOAc/petroleum ether, 1:4 → 1:1). $R_f$ = 0.70 (EtOAc/petroleum ether, 1:2); $^1$H-NMR (300 MHz, $CDCl_3$): $\delta$ = 7.83 (d, $J$ = 8.9 Hz, 2 H), 7.51 - 7.44 (m, 2 H), 7.38 (d, $J$ = 8.4 Hz, 2 H), 7.22 (d, $J$ = 8.4 Hz, 2 H), 7.00 (d, $J$ = 9.0 Hz, 2 H), 6.76 (d, $J$ = 9.2 Hz, 1 H), 4.69 (s, 2 H), 4.56 (s, 2 H), 4.35 - 4.26 (m, 4 H), 3.53 - 3.46 (m, 2 H), 1.33 (s, 9 H), 1.32 (t, $J$ = 7.1 Hz, 3 H); $^{13}$C-NMR (75 MHz, $CDCl_3$): $\delta$ = 168.6, 159.0, 150.3, 148.0, 144.5, 143.7, 138.3, 133.9, 126.6, 125.7, 123.9, 120.5, 114.8, 111.0, 108.4, 65.5, 64.3, 61.4, 54.1, 47.2, 34.5, 31.3, 14.1; IR: $\tilde{\nu}_{max}$ = 3726, 3630, 3600, 2963, 2847, 1751, 1597, 1516,

1319, 1250, 1196, 652; UV-VIS (MeCN + 0.5 % piperidine): $\lambda_{max}$ ($\varepsilon$) = 418 nm (22.4 × $10^3$ $l{\cdot}mol^{-1}{\cdot}cm^{-1}$); HRMS (ESI): *m/Z* calcd for $C_{29}H_{34}N_3O_4$ $[M+H]^+$: 488.2544, found: 488.2555.

ethyl 2-(4′-((4′′-phenyl-3′′,4′′-dihydro-2′′*H*-benzo[*b*][1′′,4′′]oxazin-7′′-yl)diazenyl)-phenoxy)acetate; (**S86**)

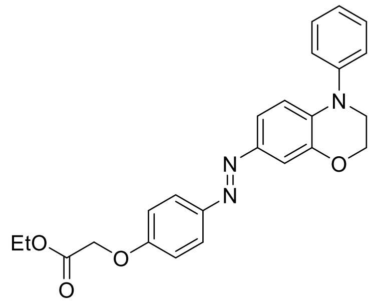

In a microwave rection vial under an atmosphere of nitrogen ethyl-2-(4′-((3′′,4′′-dihydro-2′′*H*-benzo[b][1′′,4′′]oxazin-7′′-yl)diazenyl)phenoxy) acetate (50.0 mg, 0.14 mmol, 1.0 equiv) and $Cs_2CO_3$ (47.7 mg, 0.14 mmol, 1.0 equiv) were dissolved in degassed anhydrous acetonitrile (4 ml). After addition of bromobenzyl (23 mg, 0.14 mmol, 1.0 equiv), $Pd(dpa)_2$ (8.45 mg, 14.7 µmol, 0.1 equiv) and RuPhos (13.7 mg, 29.3 µmol, 0.2 equiv) the vial was sealed and stirred at 100 °C for 19 hours. The pH was adjusted to 7 with phosphate buffer, and $CHCl_3$ was added. The organic layer was separated followed by extraction of the aqueous layer with $CHCl_3$ (3 × 7 mL). The combined organic extracts were dried with $Na_2SO_4$, filtered, and concentrated under reduced pressure. Purification by silica-gel chromatography (petroleum ether/ethyl acetate 4:1 → 1:1) provided azobenzene **S86** as a black solid (45.1 mg, 74%).$R_f$ = 0.70 (EtOAc/petroleum ether, 1:2) ; $^1$H-NMR (400 MHz, $CDCl_3$): $\delta$ = 7.86 (d, *J* = 9.1 Hz, 2 H), 7.50 (d, *J* = 2.3 Hz, 1 H), 7.49 - 7.41 (m, 2 H), 7.39 (dd, *J* = 2.2, 8.6 Hz, 1 H), 7.34 - 7.30 (m, 2 H), 7.21 (tt, *J* = 1.2, 7.3 Hz, 1 H), 7.02 (d, *J* = 9.1 Hz, 2 H), 6.94 (d, *J* = 8.8 Hz, 1 H), 4.71 (s, 2 H), 4.41 - 4.38 (m, 2 H), 4.31 (q, *J* = 7.3 Hz, 2 H), 3.84 - 3.79 (m, 2 H), 1.33 (t, *J* = 7.0 Hz, 3 H); $^{13}$C-NMR (101 MHz, $CDCl_3$): $\delta$ = 168.6, 159.3, 147.9, 146.2, 145.8, 144.5, 135.7, 129.7, 124.9, 124.4, 124.2, 118.5, 114.9, 114.9, 109.8, 65.6, 64.5, 61.5, 48.7, 14.2; IR: $\tilde{\nu}_{max}$ = 3734, 2978, 2924, 2866, 1751, 1589, 1496, 1319, 1253, 1195, 1076; UV-VIS (MeCN + 0.5 % piperidine): $\lambda_{max}$ ($\varepsilon$) = 420 nm (18.1 × $10^3$ $l{\cdot}mol^{-1}{\cdot}cm^{-1}$); HRMS (ESI): *m/Z* calcd for $C_{24}H_{24}N_3O_4$ $[M+H]^+$: 418.1761, found: 418.1772.

ethyl-2-(4´-((4´´-phenethyl-3´´,4´´-dihydro-2´´*H*-benzo[*b*][1,4]oxazin-7´´-yl)diazenyl)phenoxy)acetate; (**S87**)

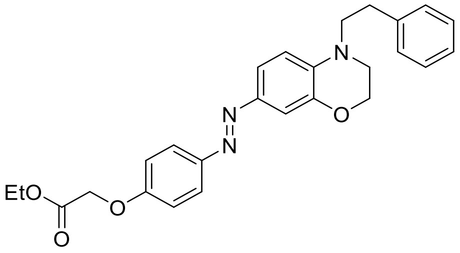

Standard Procedure **SP5** with ethyl-2-(4´-((3´´,4´´-dihydro-2´´*H*-benzo[b][1´´,4´´]oxazin-7´´-yl)diazenyl)phenoxy) acetate (50.0 mg, 0.14 mmol, 1.0 equiv.) and phenylacetaldehyde (51 µl, 0.44 mmol, 3.0 equiv.) gave azobenzene **S87** (12.6 mg, 19%) as a red solid after purification by silica-gel chromatography (EtOAc/petroleum ether, 1:4 → 1:1). $R_f$ = 0.70 (EtOAc/petroleum ether, 1:2); $^1$H-NMR (300 MHz, $CDCl_3$): $\delta$ = 7.84 (d, $J$ = 9.0 Hz, 2 H), 7.54 (dd, $J$ = 2.3, 8.6 Hz, 1 H), 7.43 (d, $J$ = 2.2 Hz, 1 H), 7.37 - 7.29 (m, 2 H), 7.26 - 7.21 (m, 3 H), 7.01 (d, $J$ = 9.0 Hz, 2 H), 6.78 (d, $J$ = 8.8 Hz, 1 H), 4.69 (s, 2 H), 4.30 (q, $J$ = 7.1 Hz, 2 H), 4.18 - 4.11 (m, 2 H), 3.63 (t, $J$ = 7.4 Hz, 2 H), 3.33 - 3.25 (m, 2 H), 2.96 (t, $J$ = 7.4 Hz, 2 H), 1.32 (t, $J$ = 7.1 Hz, 3 H); $^{13}$C-NMR (75 MHz, $CDCl_3$): $\delta$ = 168.7, 159.0, 148.0, 144.2, 143.8, 139.1, 137.5, 128.8, 128.7, 126.6, 123.9, 120.9, 114.8, 110.2, 108.4, 65.5, 64.0, 61.5, 52.8, 47.6, 32.7, 14.2; IR: $\tilde{\nu}_{max}$ = 3734, 3063, 2978, 2932, 2870, 1755, 1597, 1516, 1350, 1319, 1250, 1196, 1153; UV-VIS (MeCN + 0.5 % piperidine): $\lambda_{max}$ ($\varepsilon$) = 427 nm (16.9 × $10^3$ l·mol$^{-1}$·cm$^{-1}$); HRMS (ESI): *m/Z* calcd for $C_{26}H_{28}N_3O_4$ [M+H]$^+$: 446.2074, found: 446.2083.

### Supplementary Note 3: optojasp $^{1}$H- /$^{13}$C-NMRs

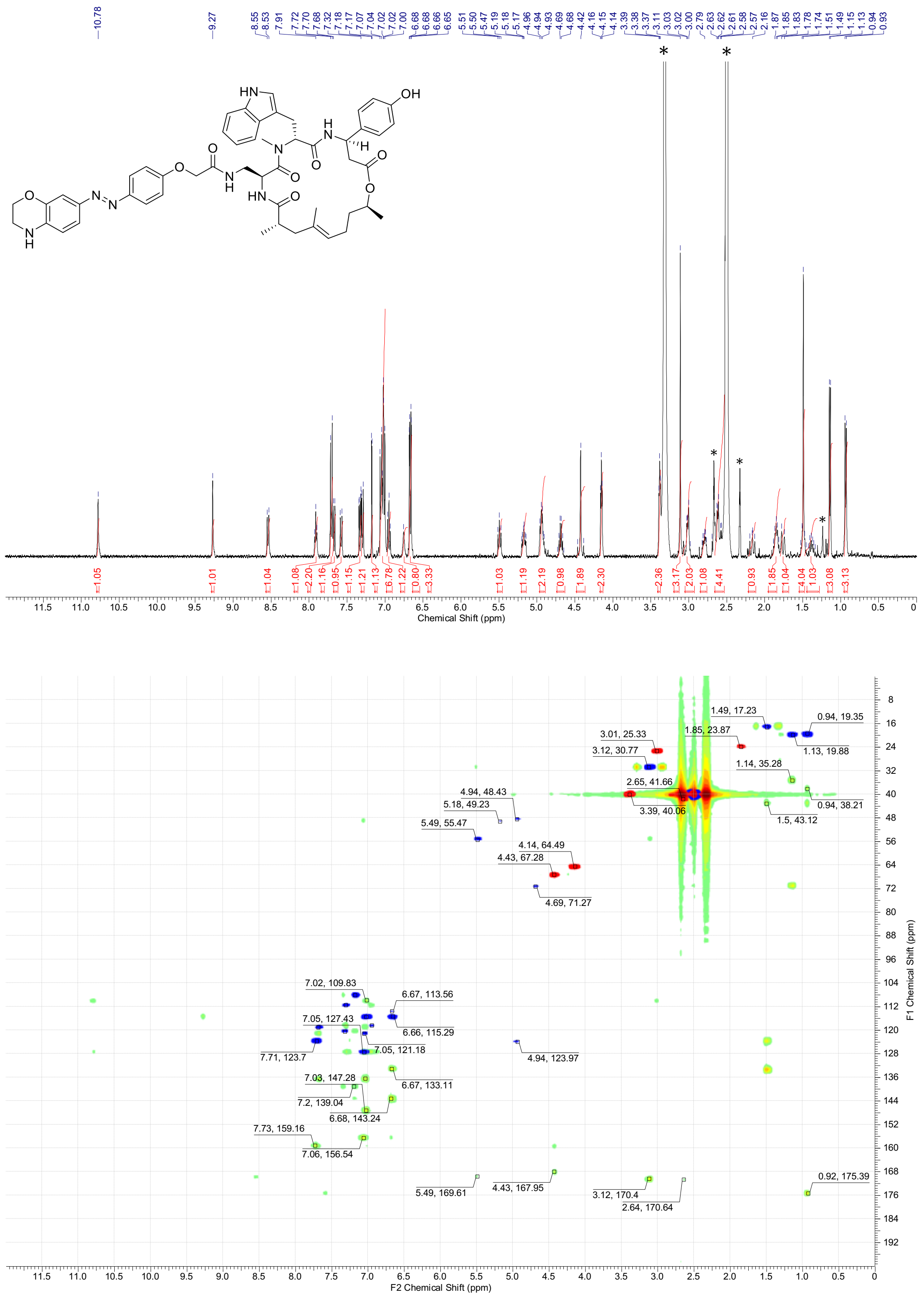

$^{1}$H-NMR spectra (400 MHz, DMSO-$d_6$) and overlay of $^{13}$C-HSQC (red/blue) and $^{13}$C-HMBC (green) spectra (100 MHz, DMSO-$d_6$) of **5**. * = NMR-solvent, $H_2O$, grease

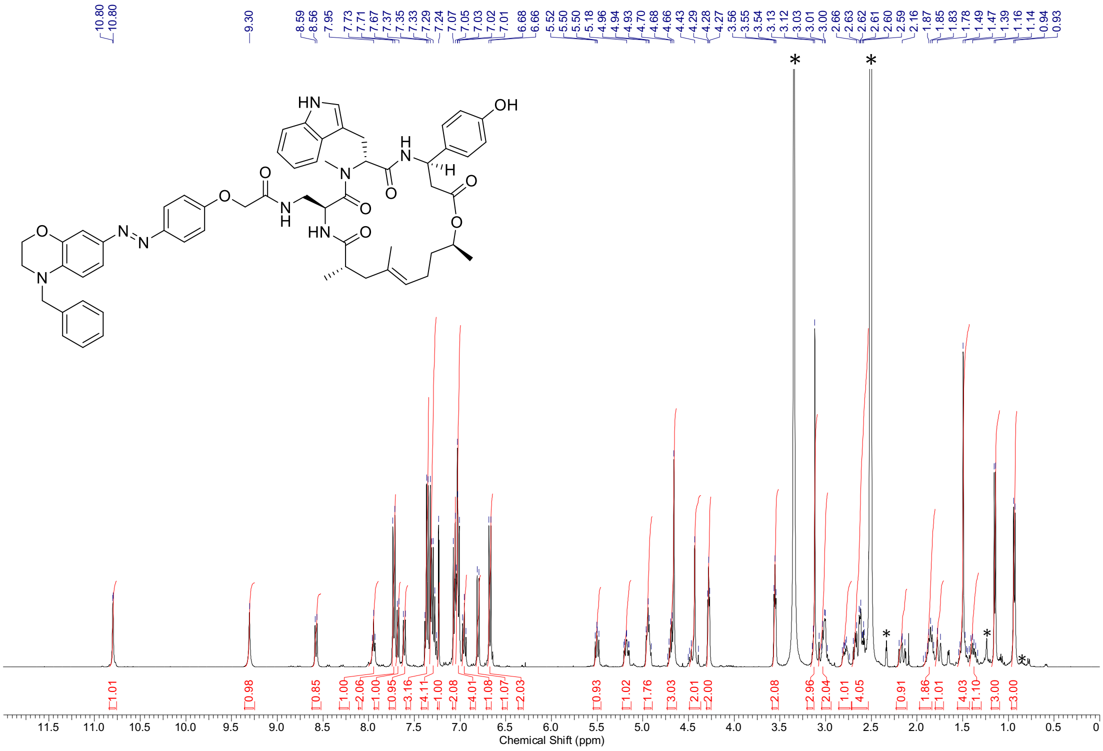

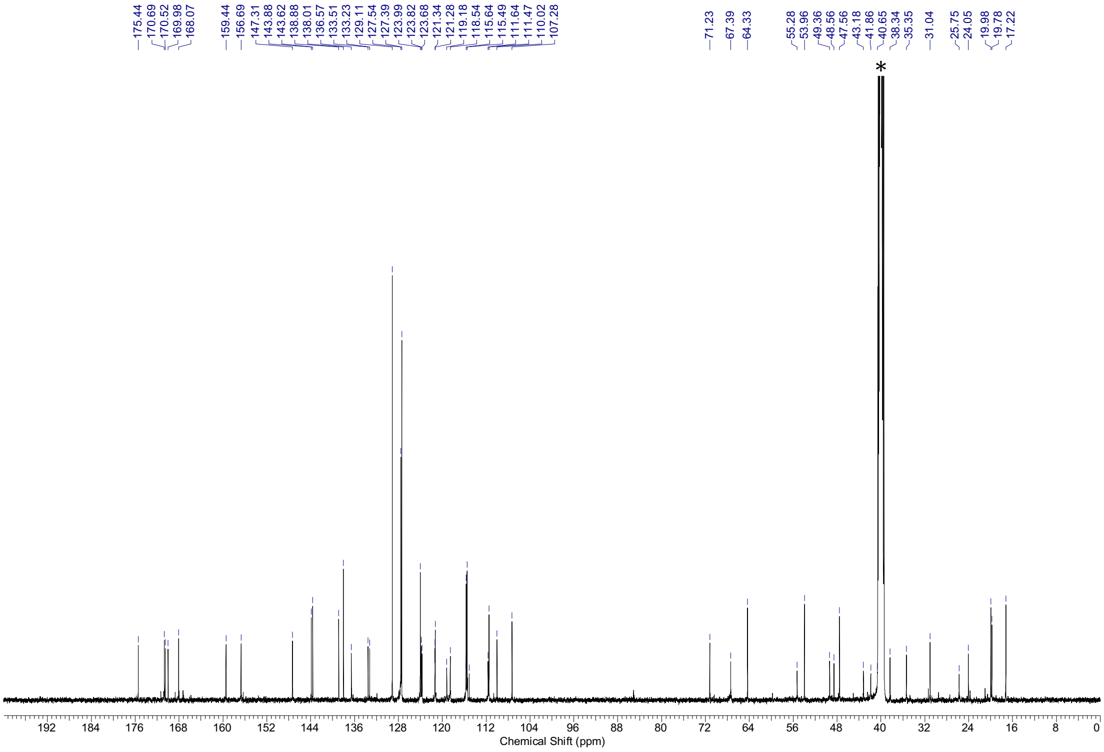

[1]H-NMR spectra (500 MHz, DMSO-*d*$_6$) and [13]C-NMR spectra (126 MHz, DMSO-*d*$_6$) of neo Optojasp (nOJ) **6.** * = NMR-solvent, $H_2O$, grease

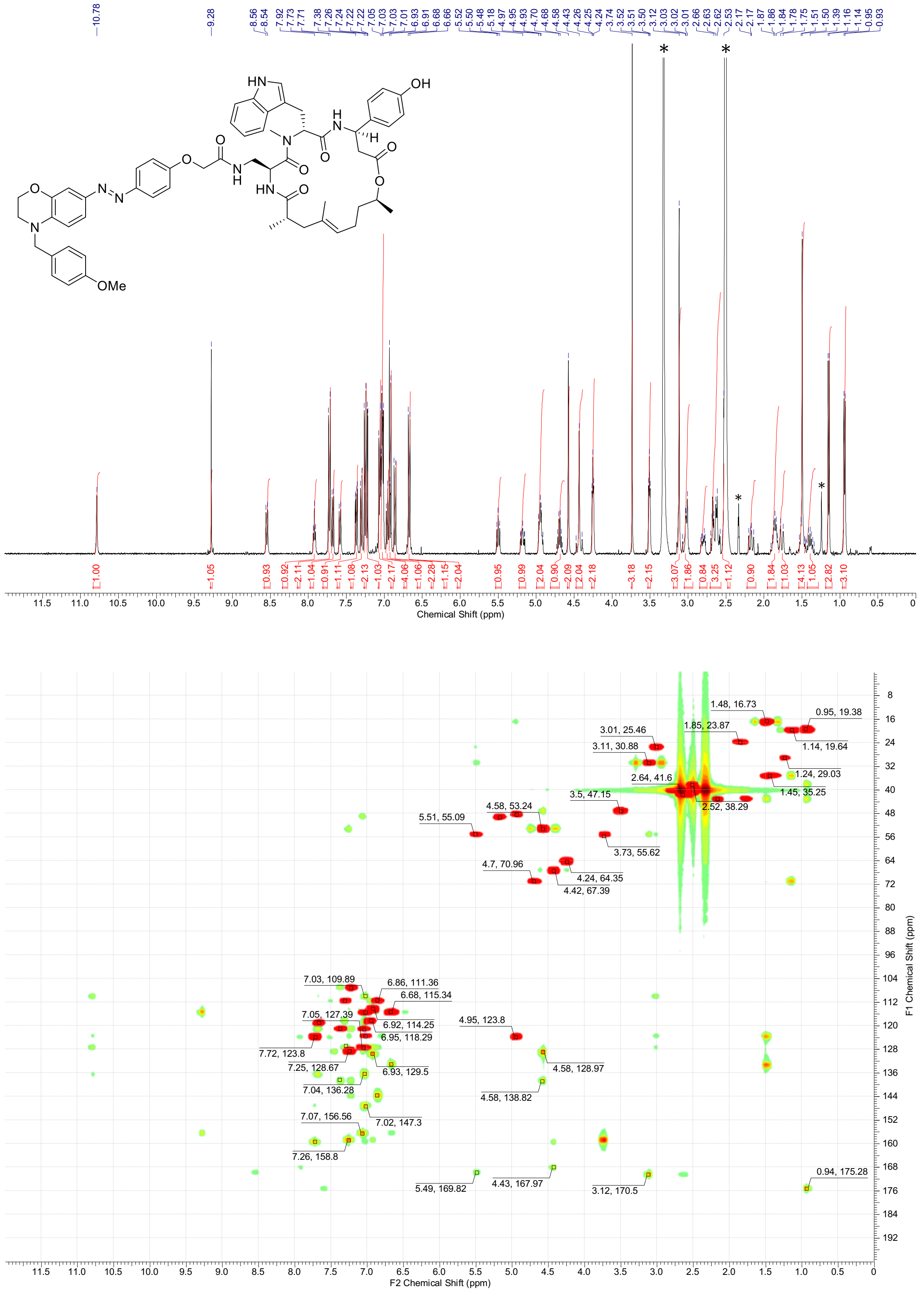

$^1$H-NMR spectra (600 MHz, DMSO-$d_6$) and overlay of $^{13}$C-HSQC (red) and $^{13}$C-HMBC (green) spectra (150 MHz, DMSO-$d_6$) of **7**. * = NMR-solvent, $H_2O$, grease

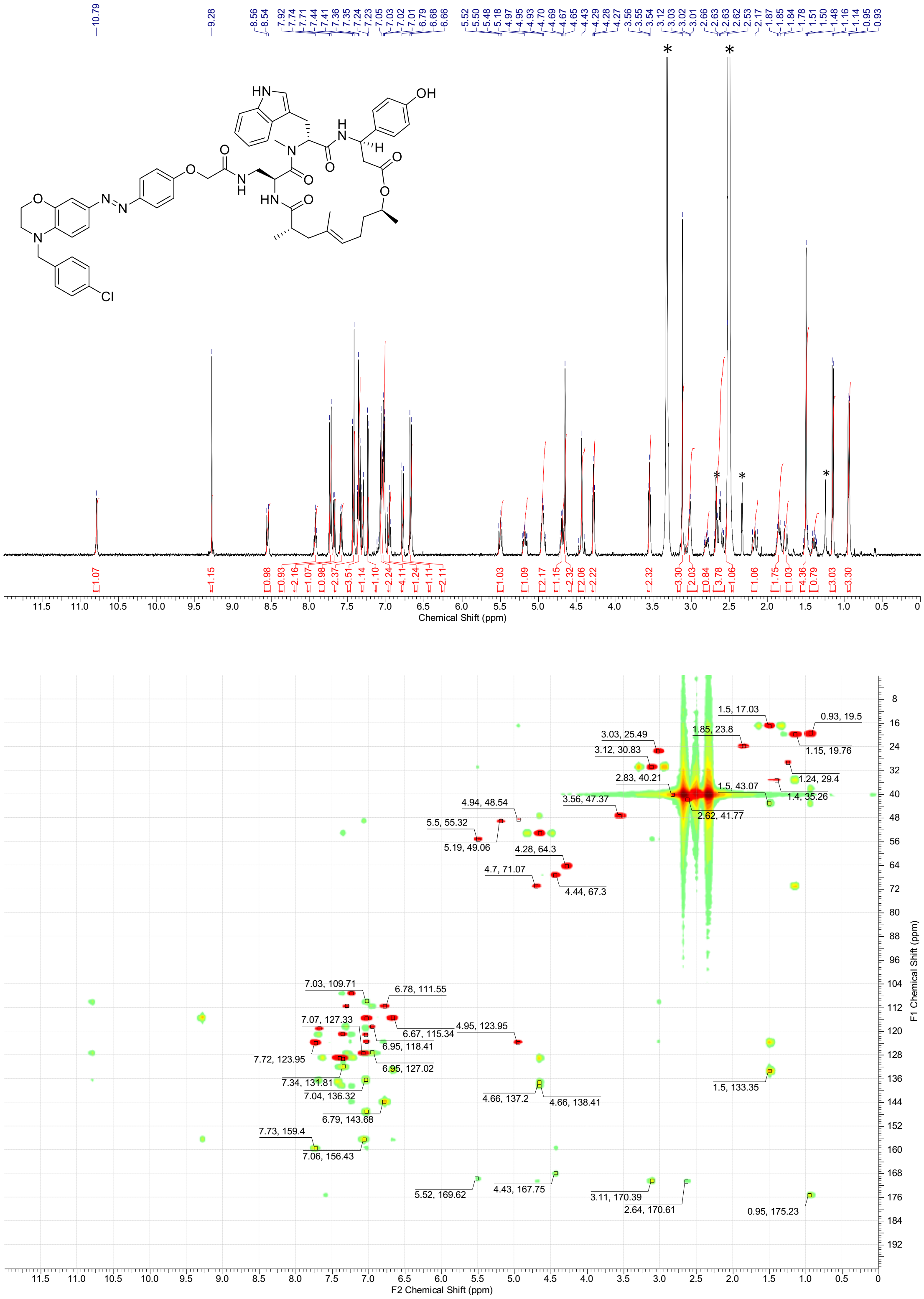

[1]H-NMR spectra (600 MHz, DMSO-$d_6$) and overlay of [13]C-HSQC (red) and [13]C-HMBC (green) spectra (150 MHz, DMSO-$d_6$) of **8.** * = NMR-solvent, $H_2O$, grease

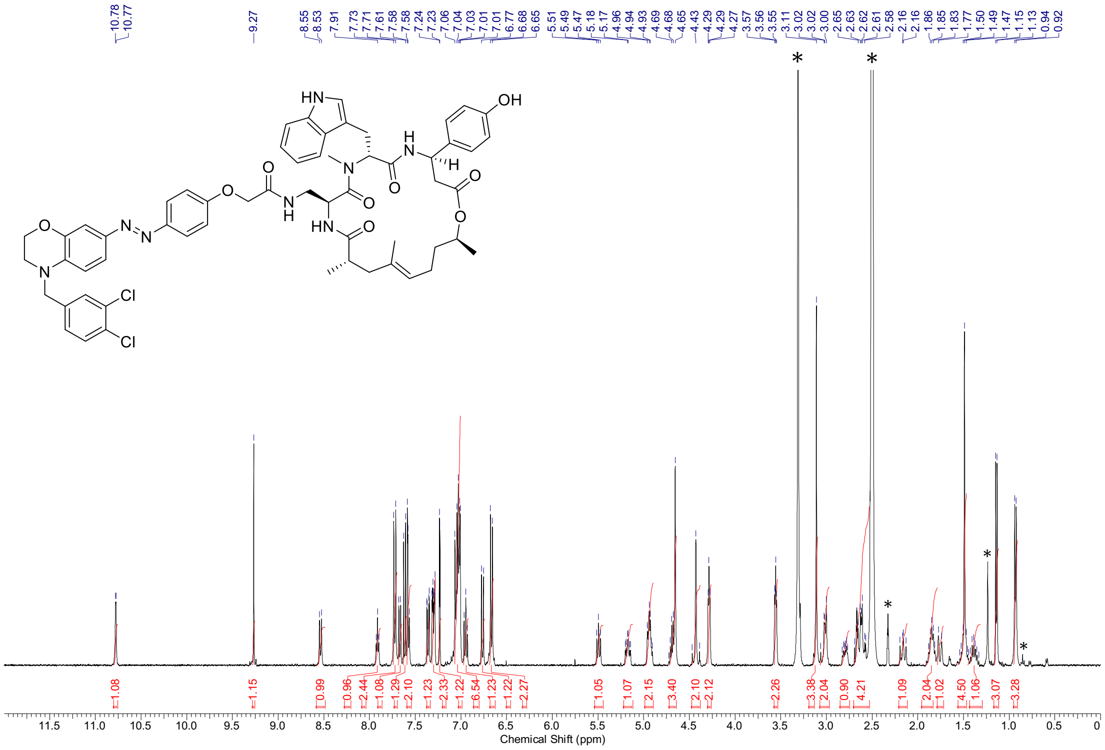

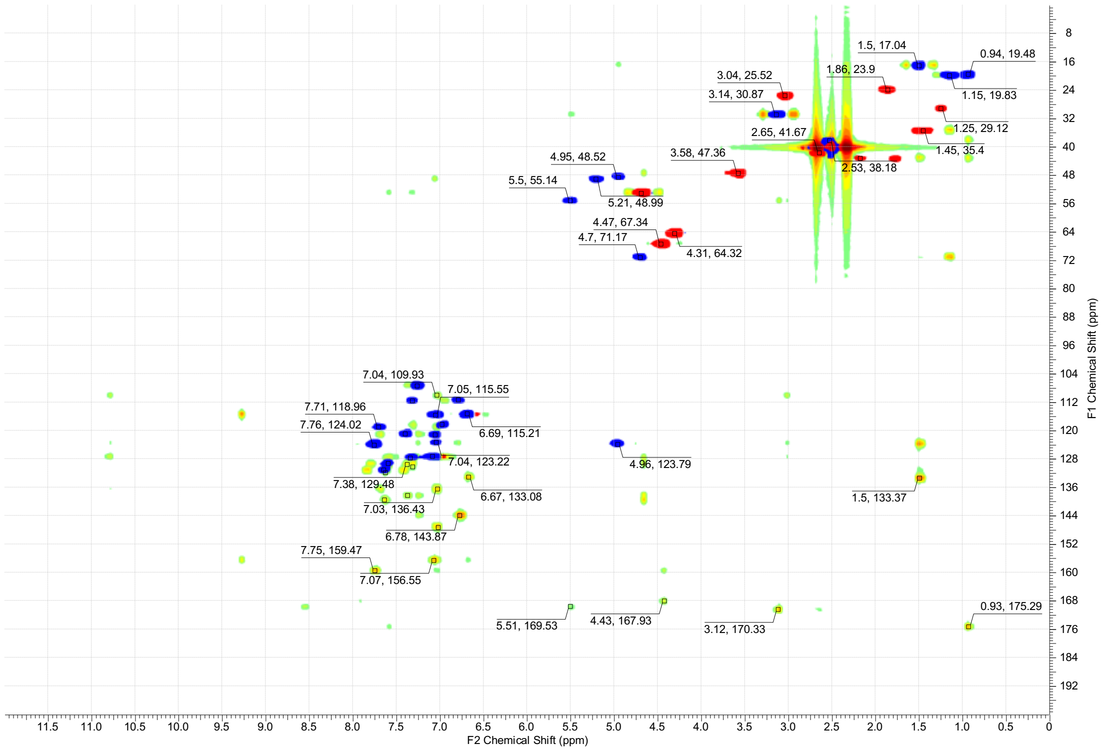

$^{1}$H-NMR spectra (400 MHz, DMSO-$d_6$) and overlay of $^{13}$C-HSQC (red/blue) and $^{13}$C-HMBC (green) spectra (100 MHz, DMSO-$d_6$) of **9.** * = NMR-solvent, $H_2O$, grease

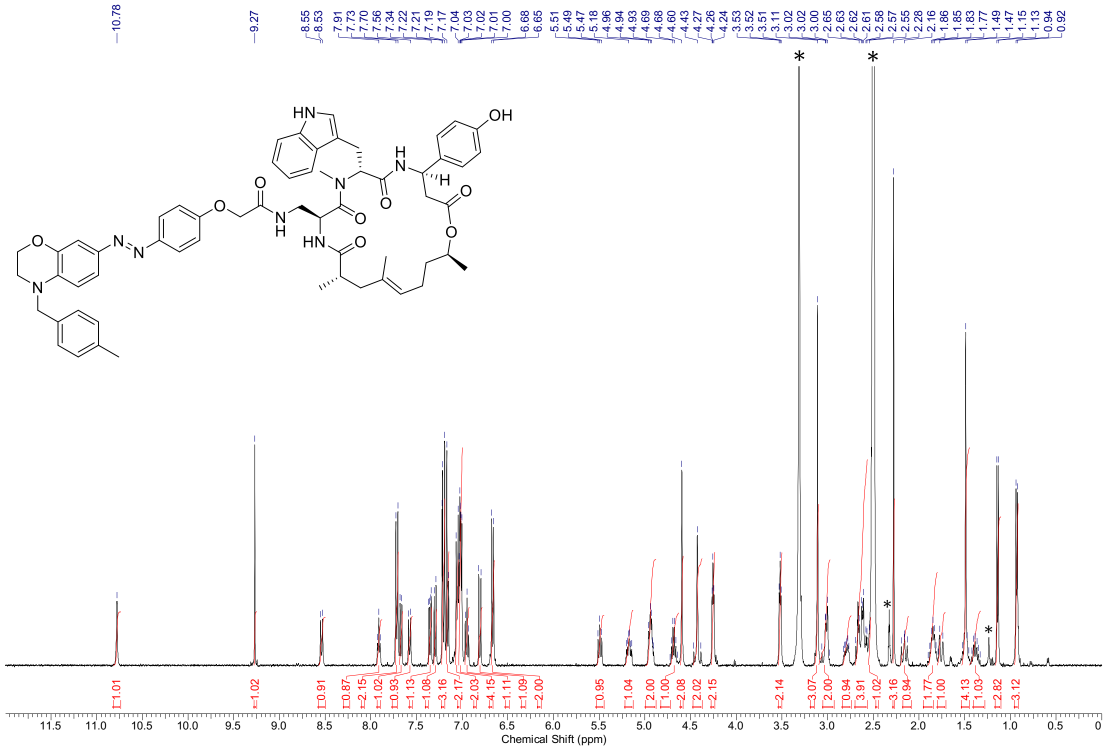

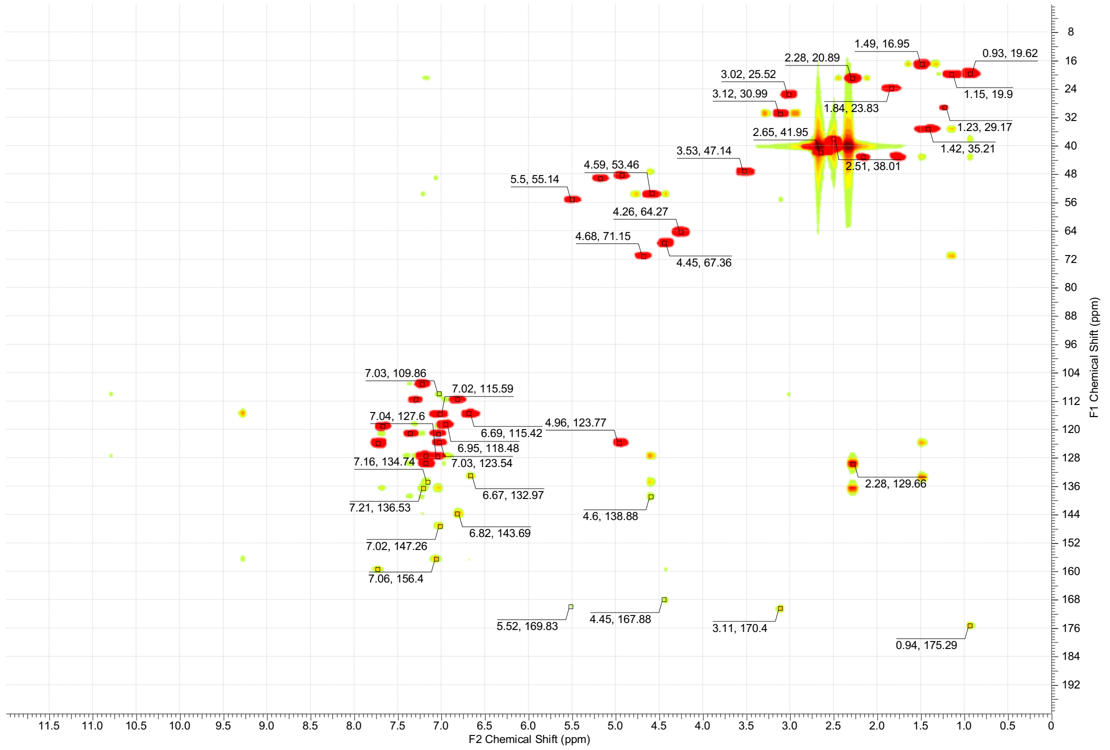

$^{1}$H-NMR spectra (600 MHz, DMSO-$d_6$) and overlay of $^{13}$C-HSQC (red) and $^{13}$C-HMBC (green) spectra (150 MHz, DMSO-$d_6$) of **10.** * = NMR-solvent, $H_2O$, grease

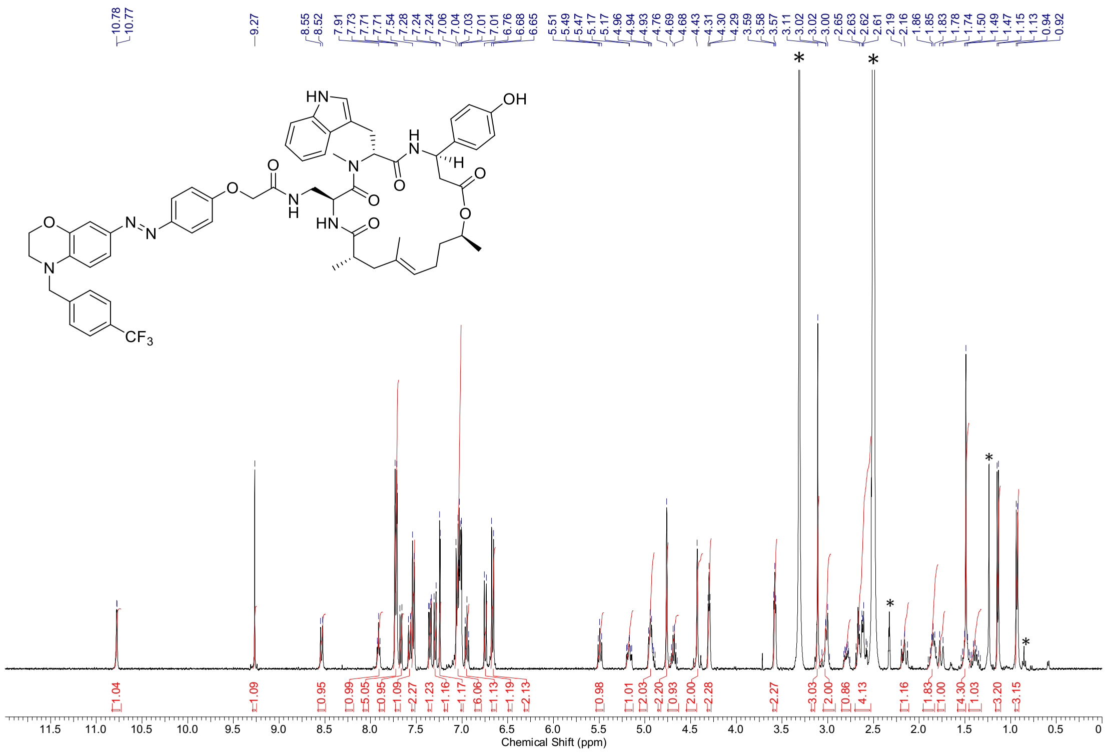

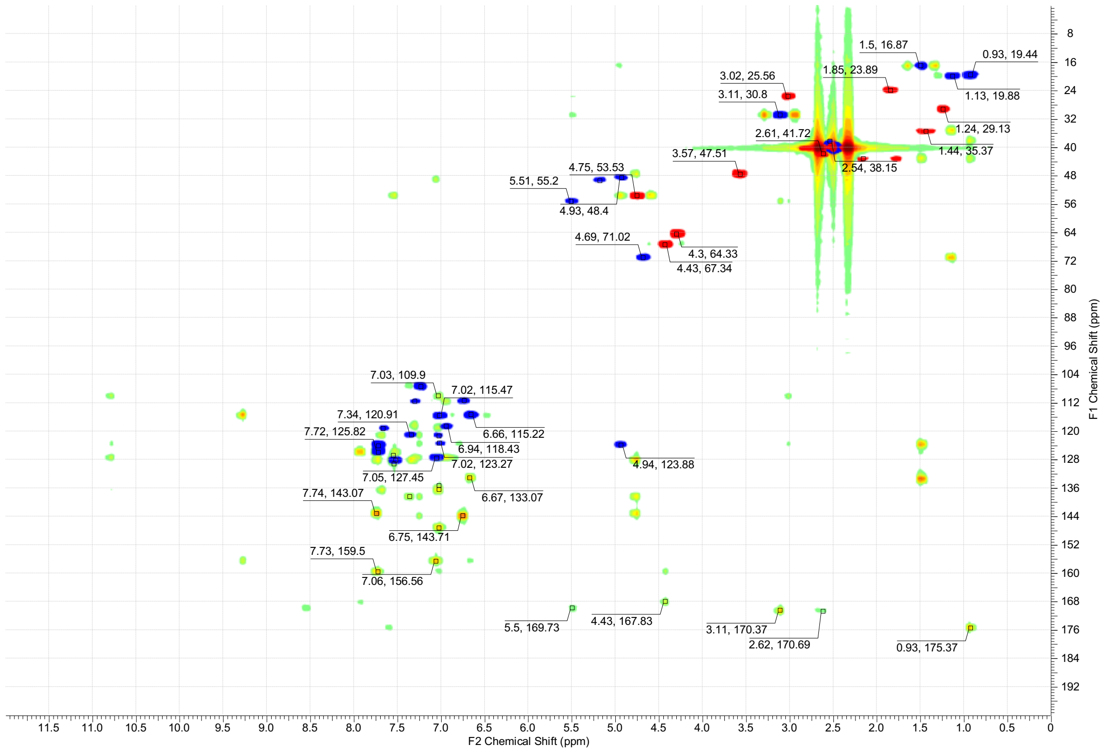

$^{1}$H-NMR spectra (400 MHz, DMSO-$d_6$) and overlay of $^{13}$C-HSQC (red/blue) and $^{13}$C-HMBC (green) spectra (100 MHz, DMSO-$d_6$) of **11.** * = NMR-solvent, $H_2O$, grease

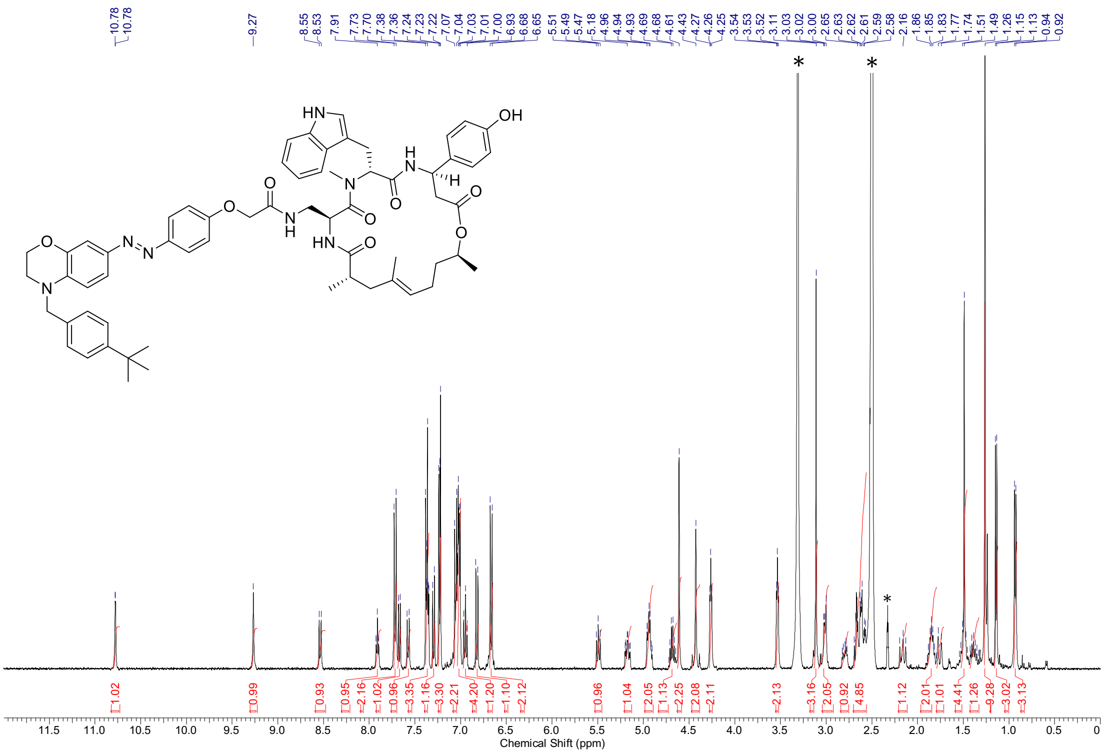

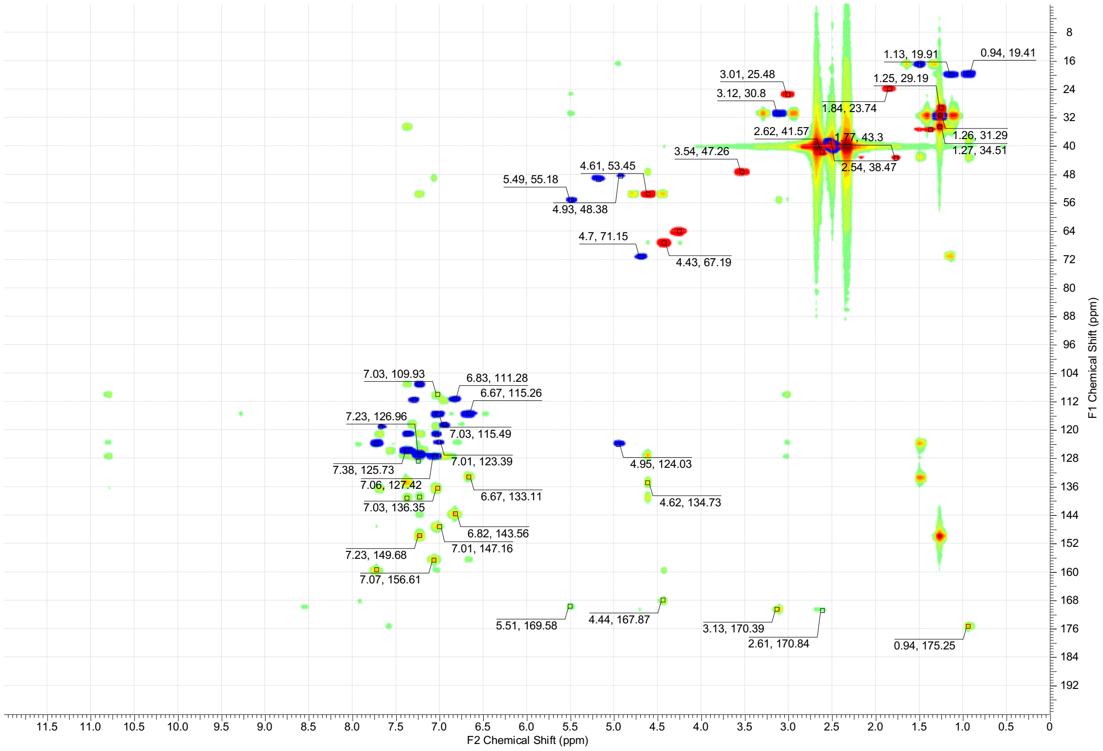

$^{1}$H-NMR spectra (600 MHz, DMSO-$d_6$) and overlay of $^{13}$C-HSQC (red/blue) and $^{13}$C-HMBC (green) spectra (150 MHz, DMSO-$d_6$) of **12.** * = NMR-solvent, $H_2O$, grease

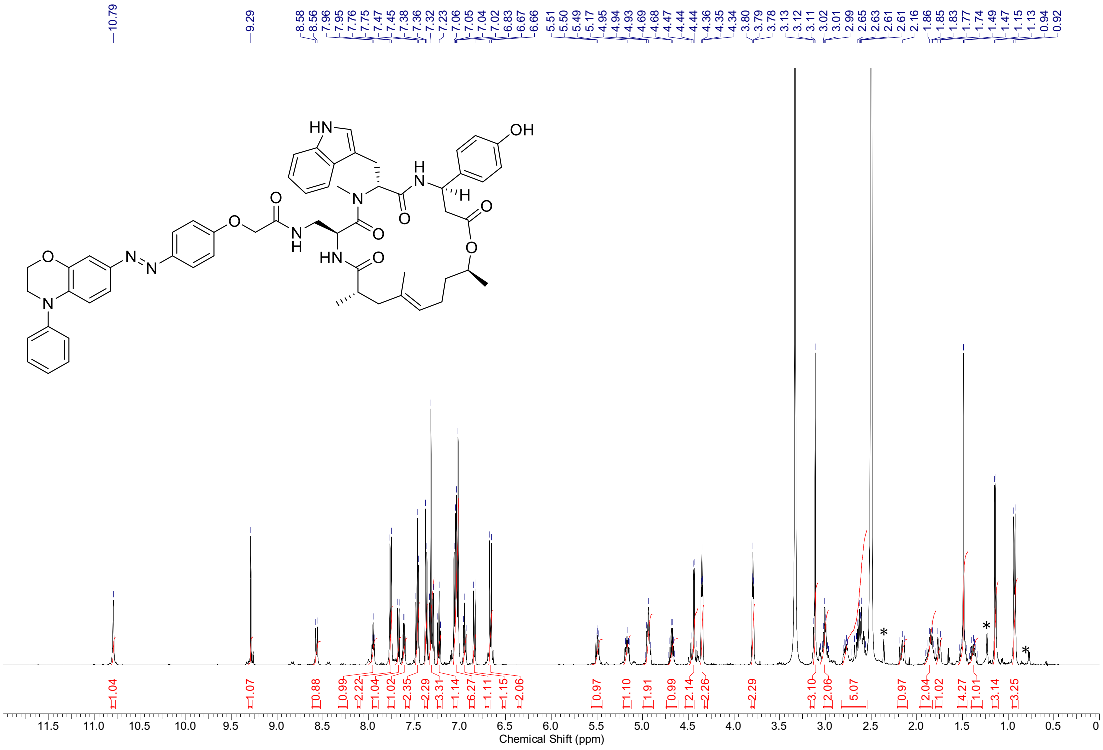

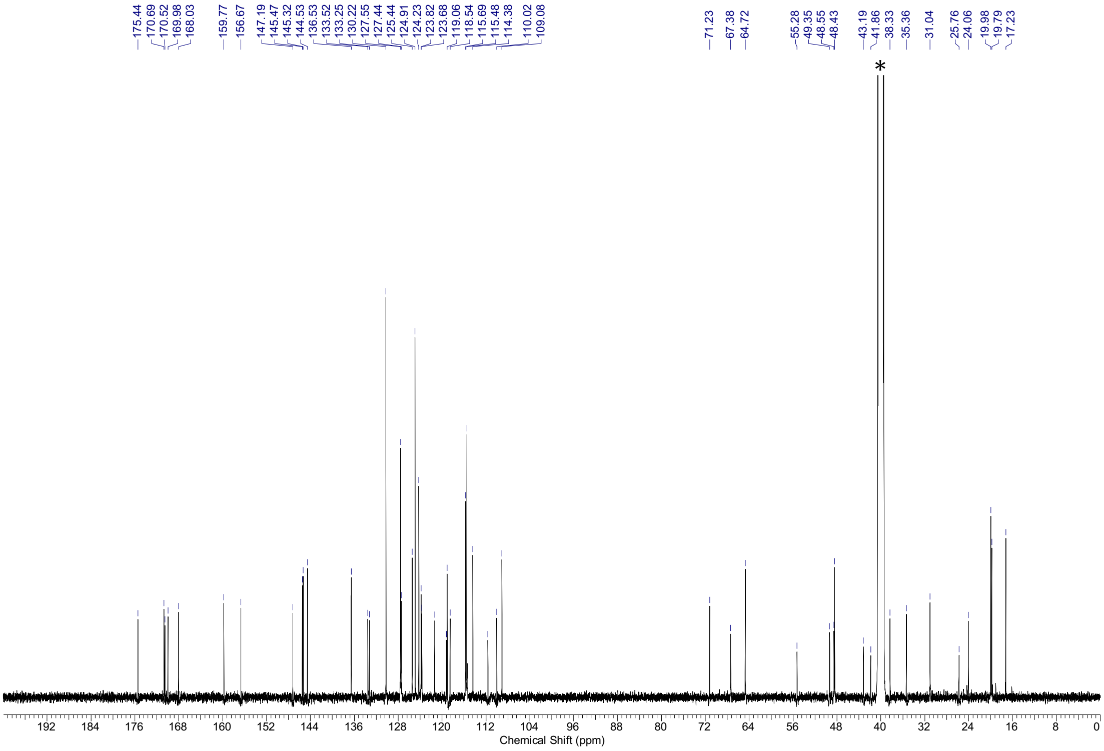

$^1$H-NMR spectra (500 MHz, DMSO-$d_6$) and $^{13}$C-NMR spectra (126 MHz, DMSO-$d_6$) of **13.** * = NMR-solvent, $H_2O$, grease

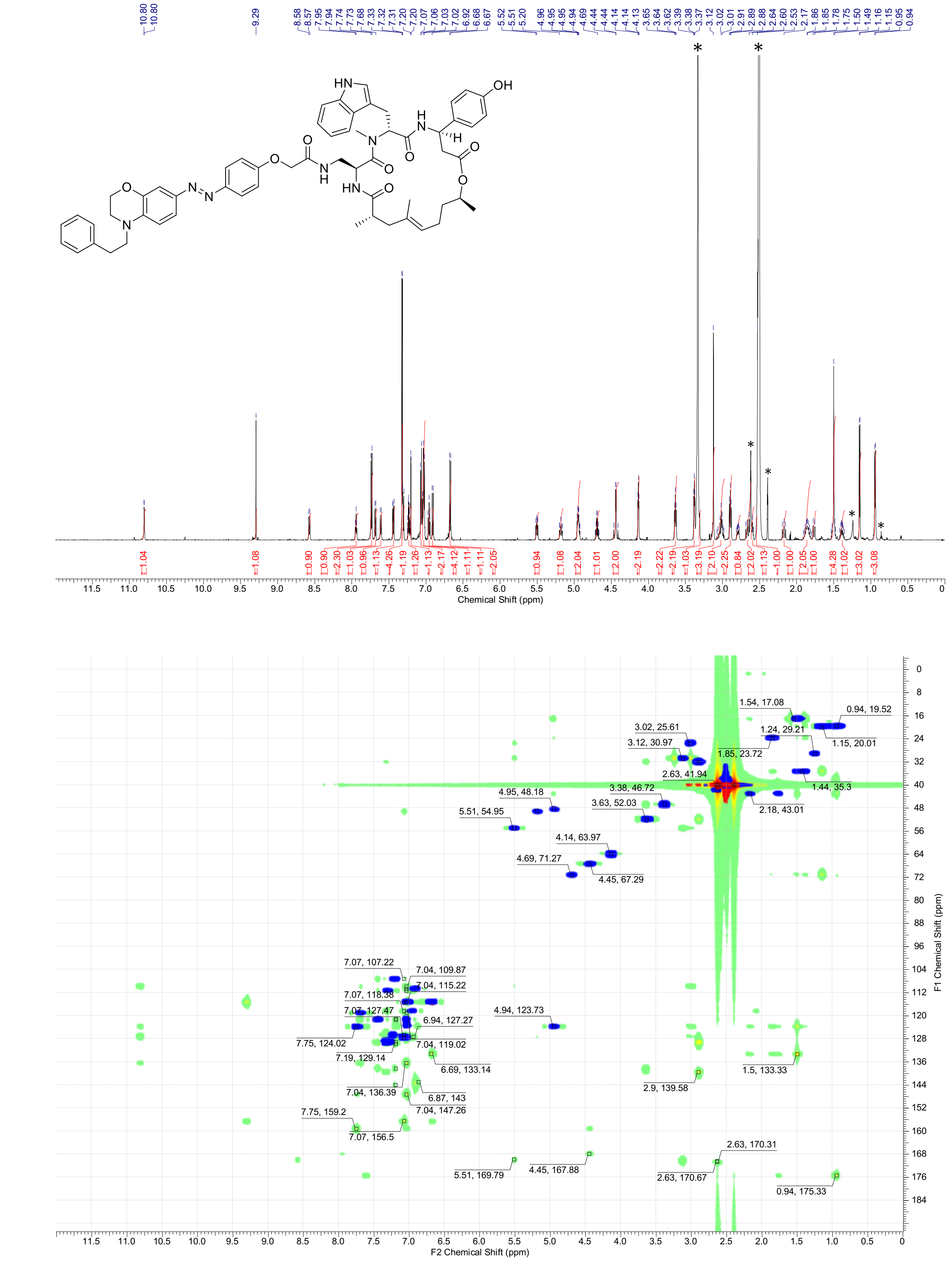

$^{1}$H-NMR spectra (600 MHz, DMSO-$d_6$) and overlay of $^{13}$C-HSQC (blue) and $^{13}$C-HMBC (green) spectra (150 MHz, DMSO-$d_6$) of **14.** * = NMR-solvent, $H_2O$, grease

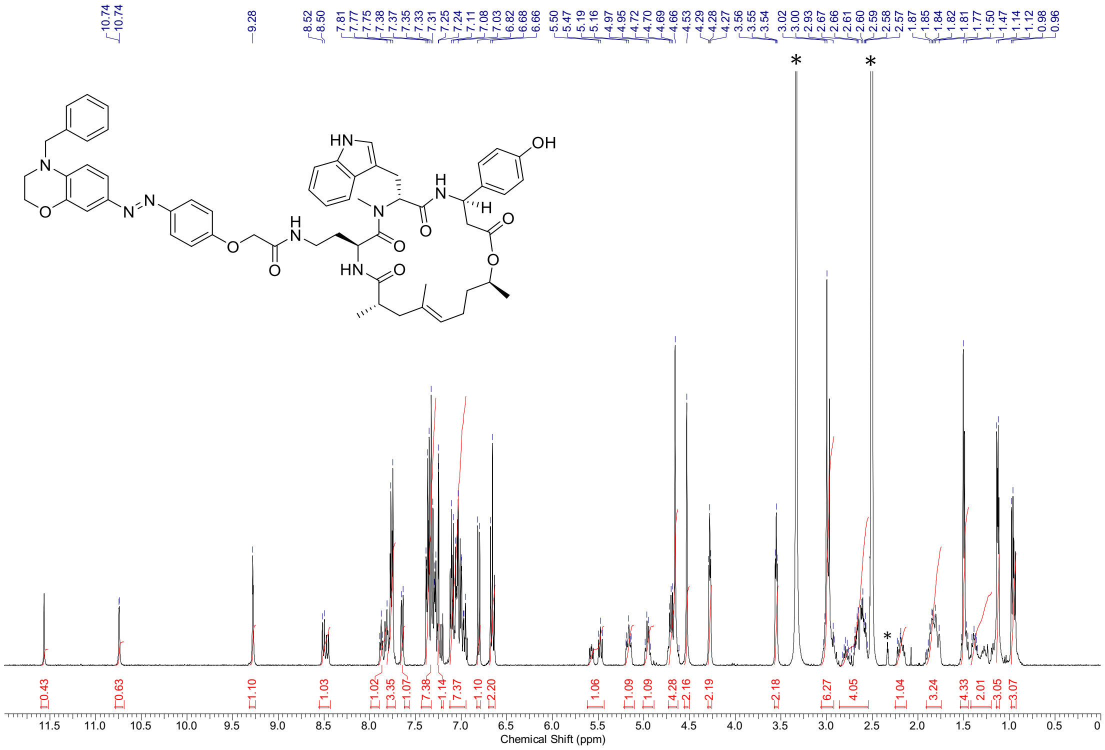

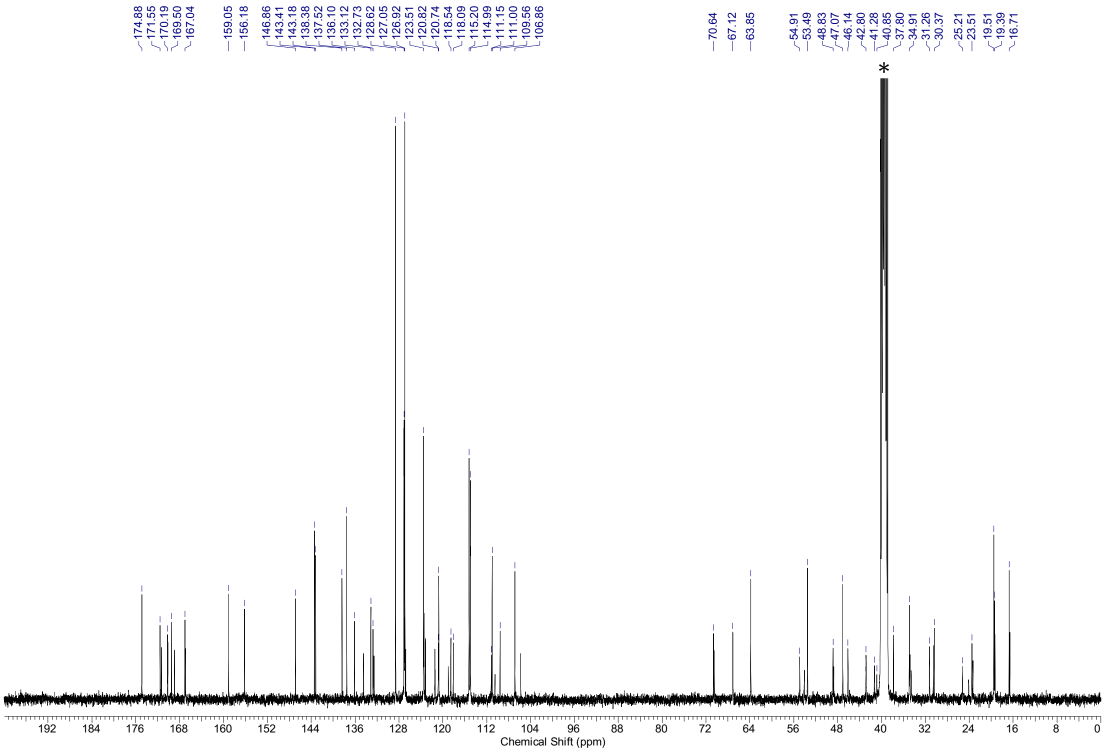

$^{1}H$-NMR spectra (500 MHz, DMSO-$d_6$) and $^{13}C$-NMR spectra (126 MHz, DMSO-$d_6$) of **15.** * = NMR-solvent, $H_2O$

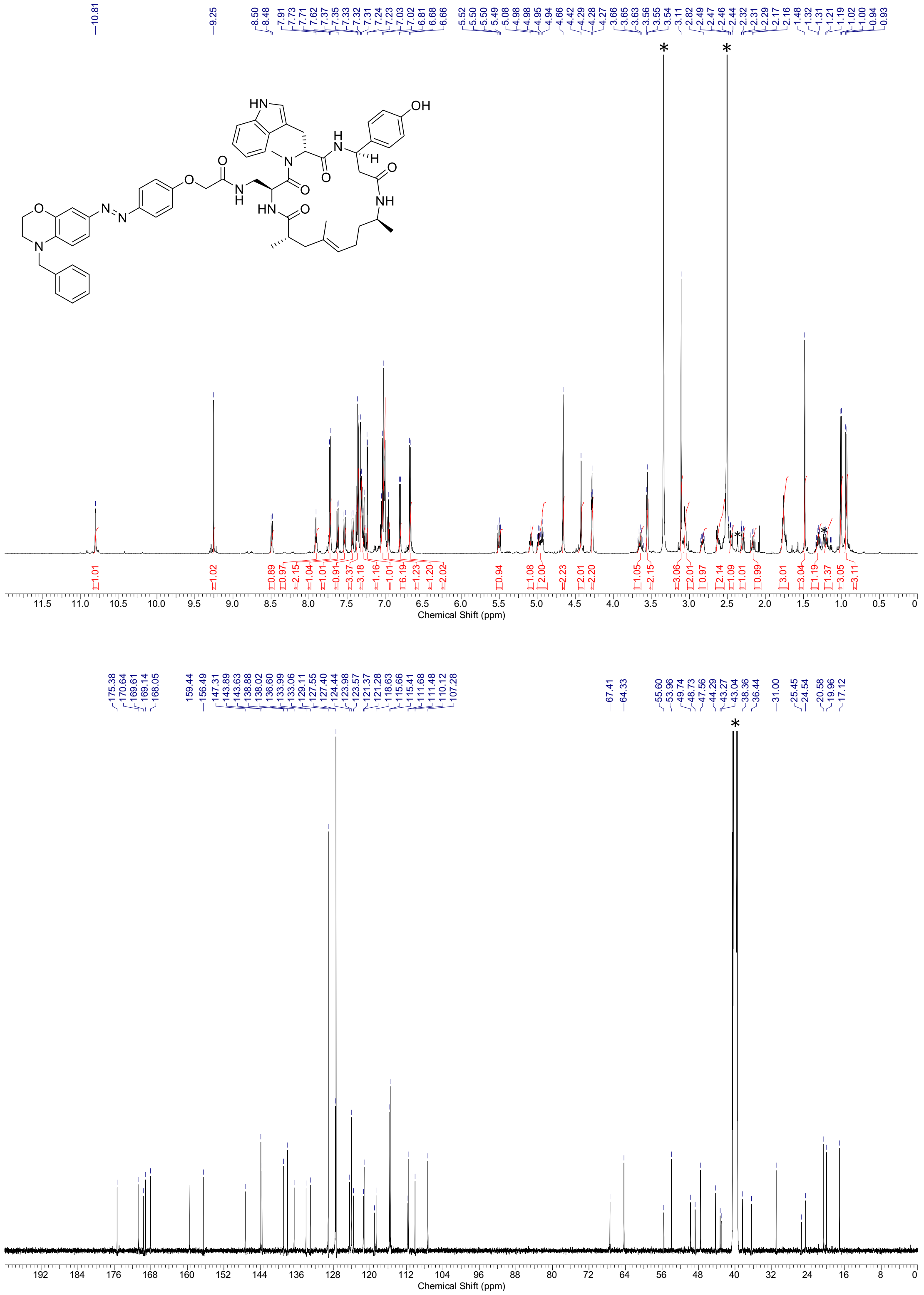

$^{1}$H-NMR spectra (500 MHz, DMSO-$d_6$) and $^{13}$C-NMR spectra (126 MHz, DMSO-$d_6$) of **16.** * = NMR-solvent, $H_2O$, grease

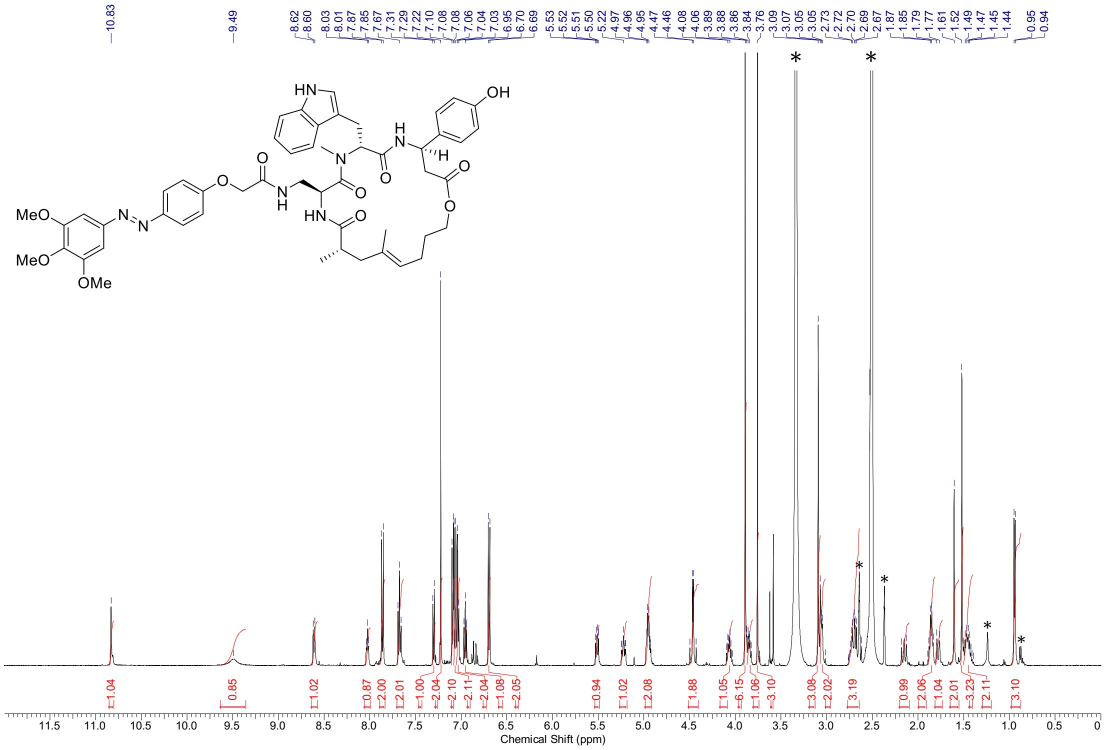

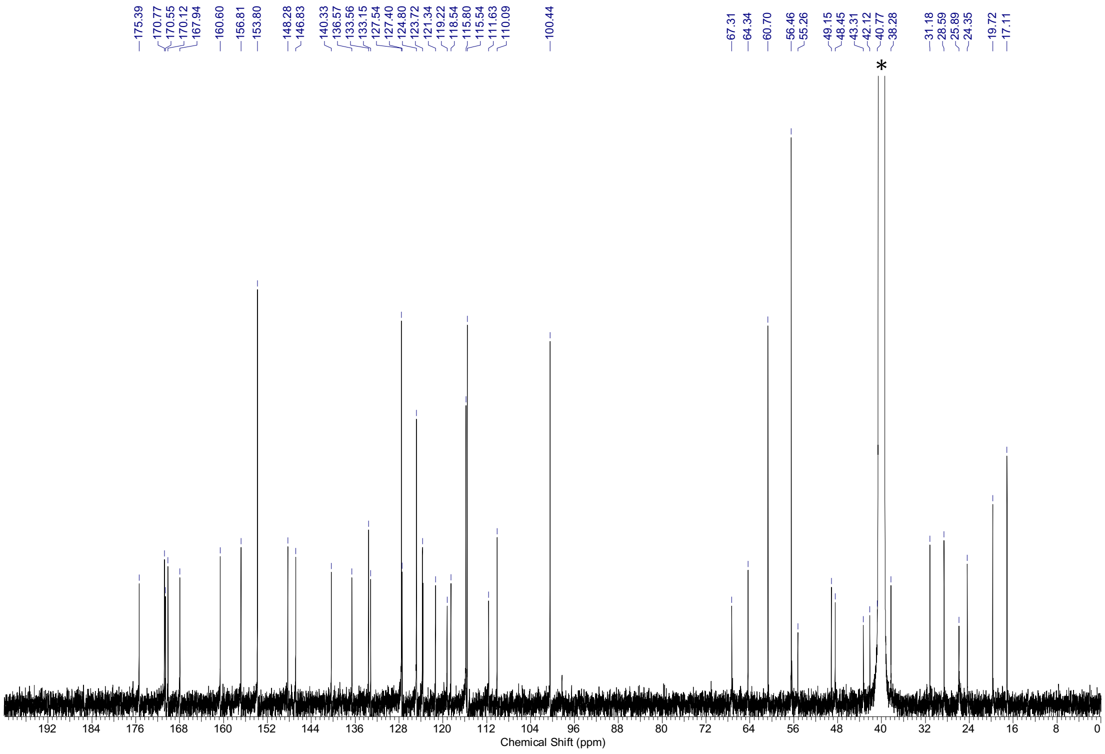

$^{1}$H-NMR spectra (500 MHz, DMSO-$d_6$) and $^{13}$C-NMR spectra (126 MHz, DMSO-$d_6$) of **S1.** * = NMR-solvent, $H_2O$, grease

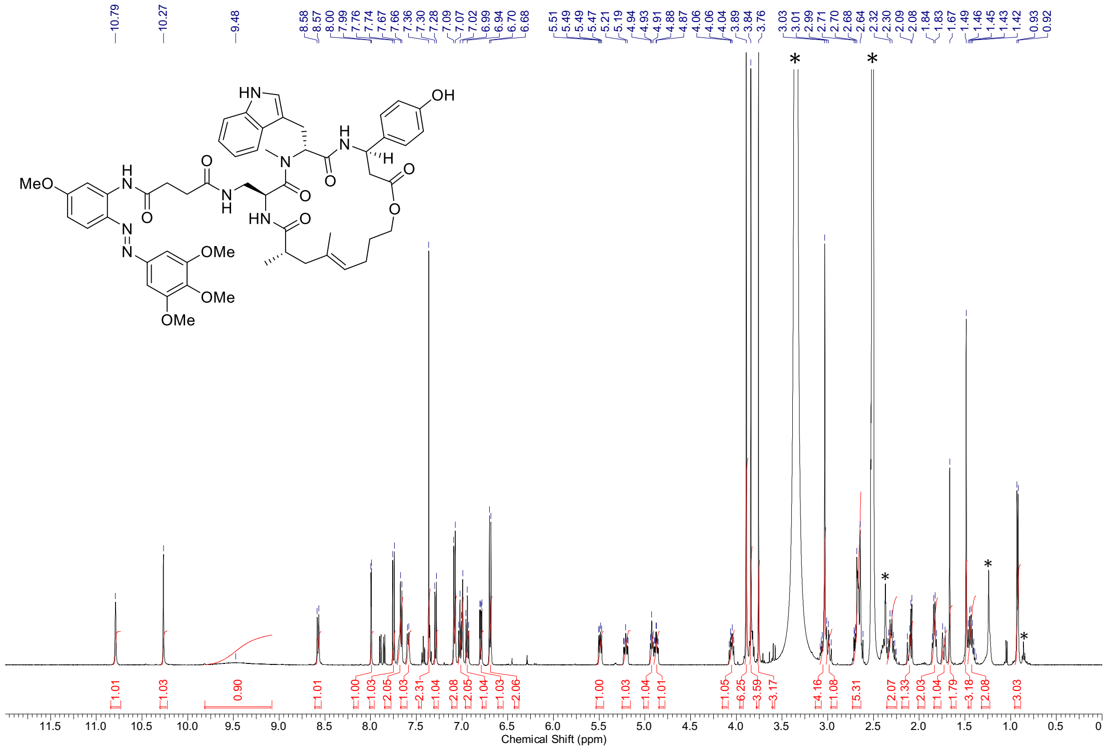

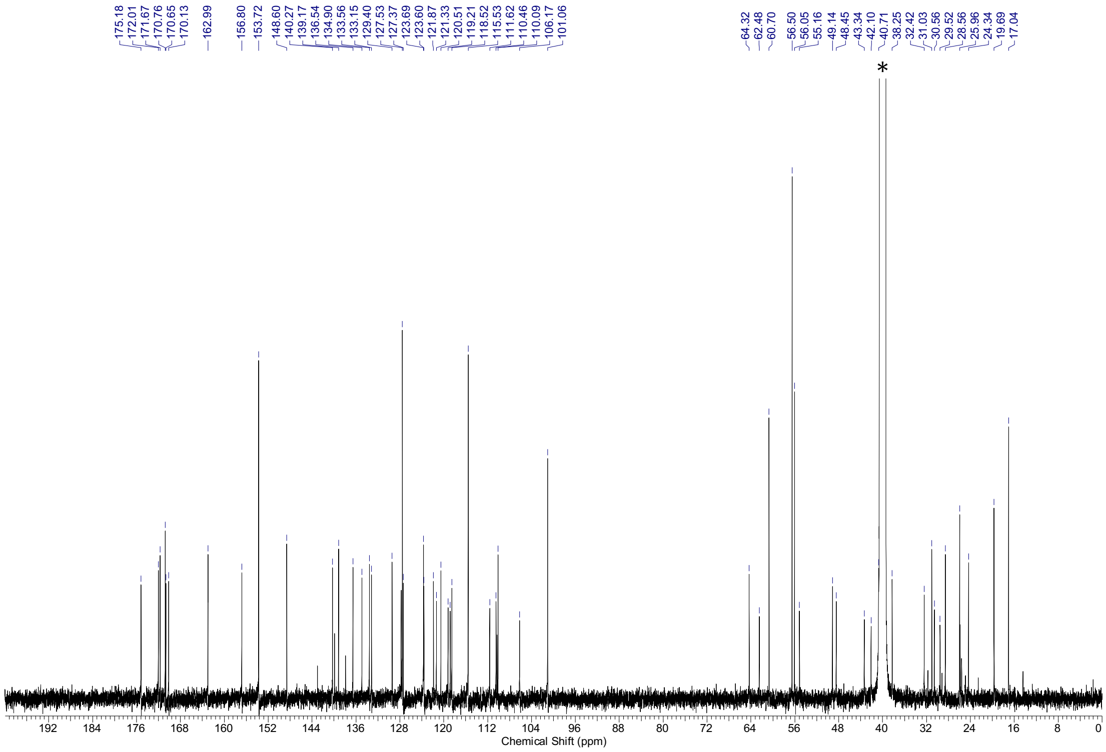

[1]H-NMR spectra (500 MHz, DMSO-$d_6$) and [13]C-NMR spectra (126 MHz, DMSO-$d_6$) of **S2**. * = NMR-solvent, $H_2O$, grease

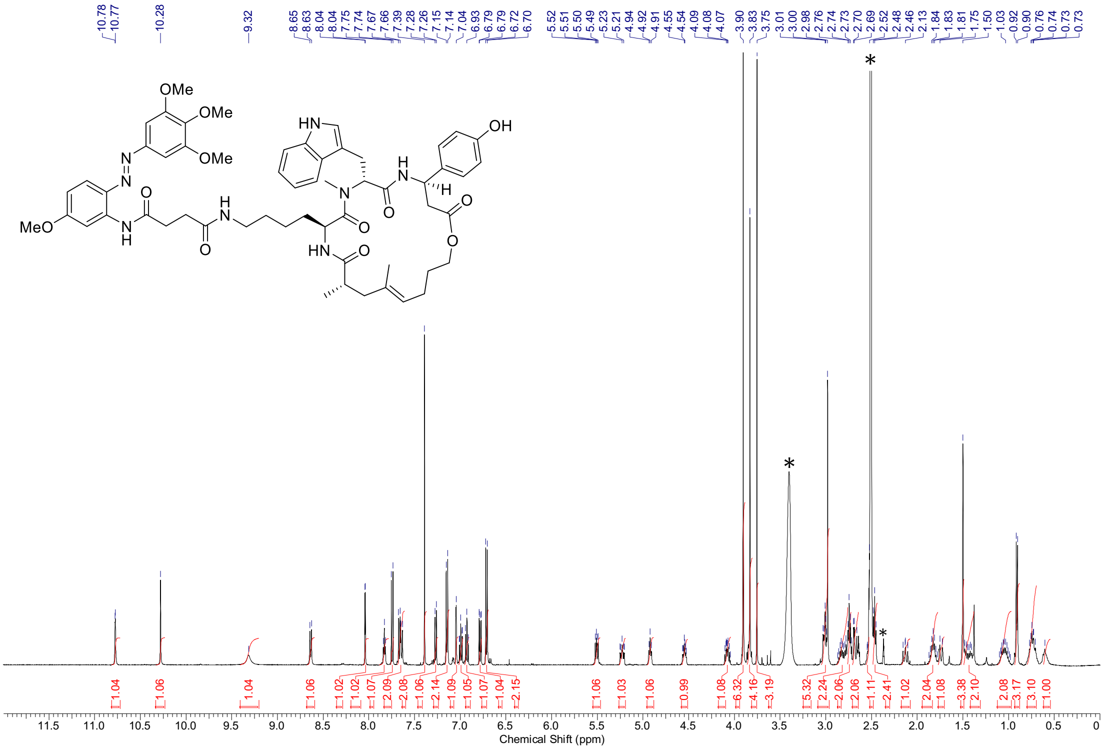

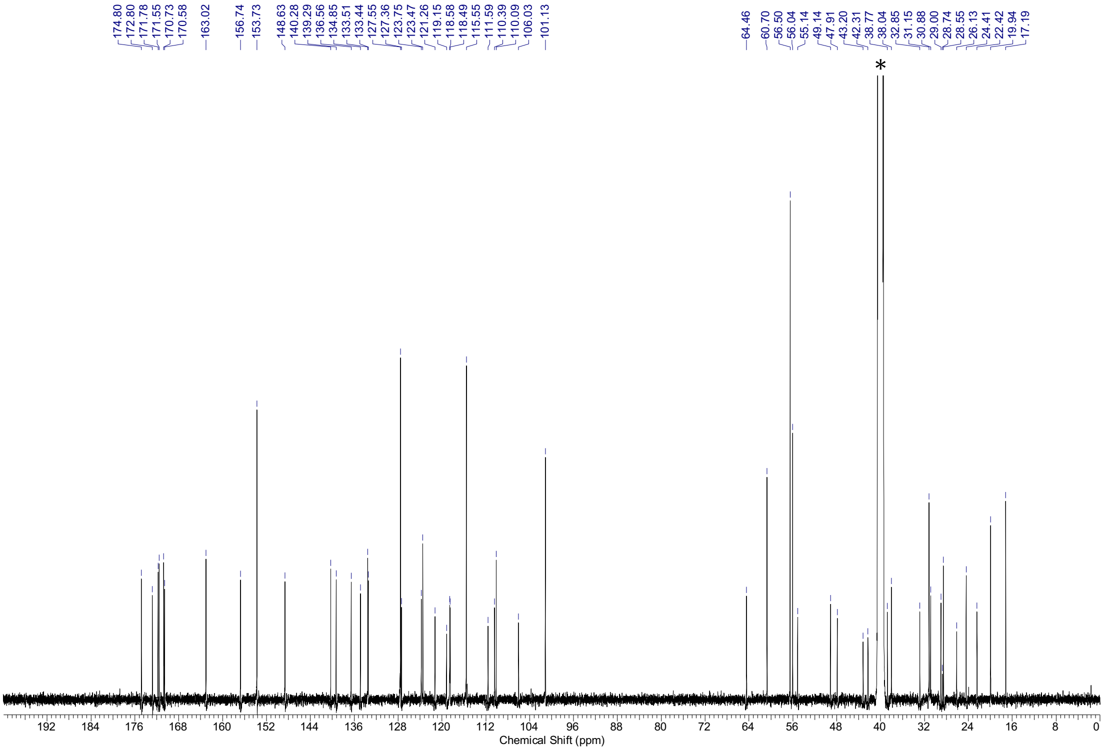

$^{1}H$-NMR spectra (500 MHz, DMSO-*d*$_6$) and $^{13}C$-NMR spectra (126 MHz, DMSO-*d*$_6$) of **S3.** * = NMR-solvent, $H_2O$

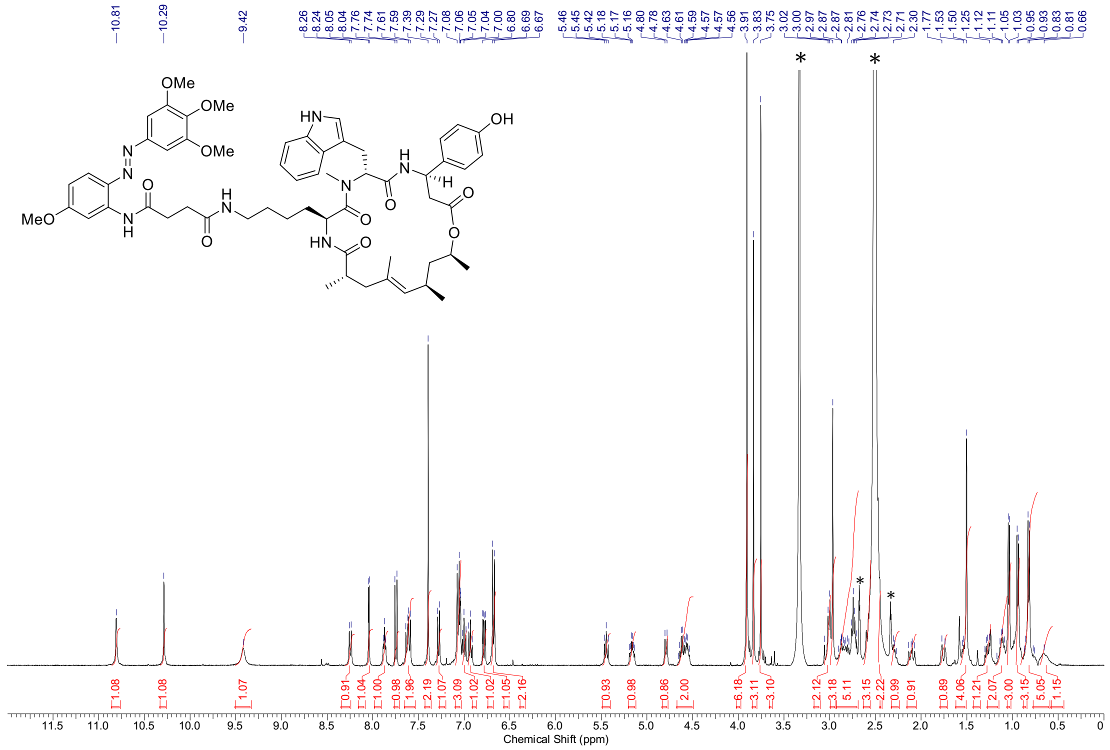

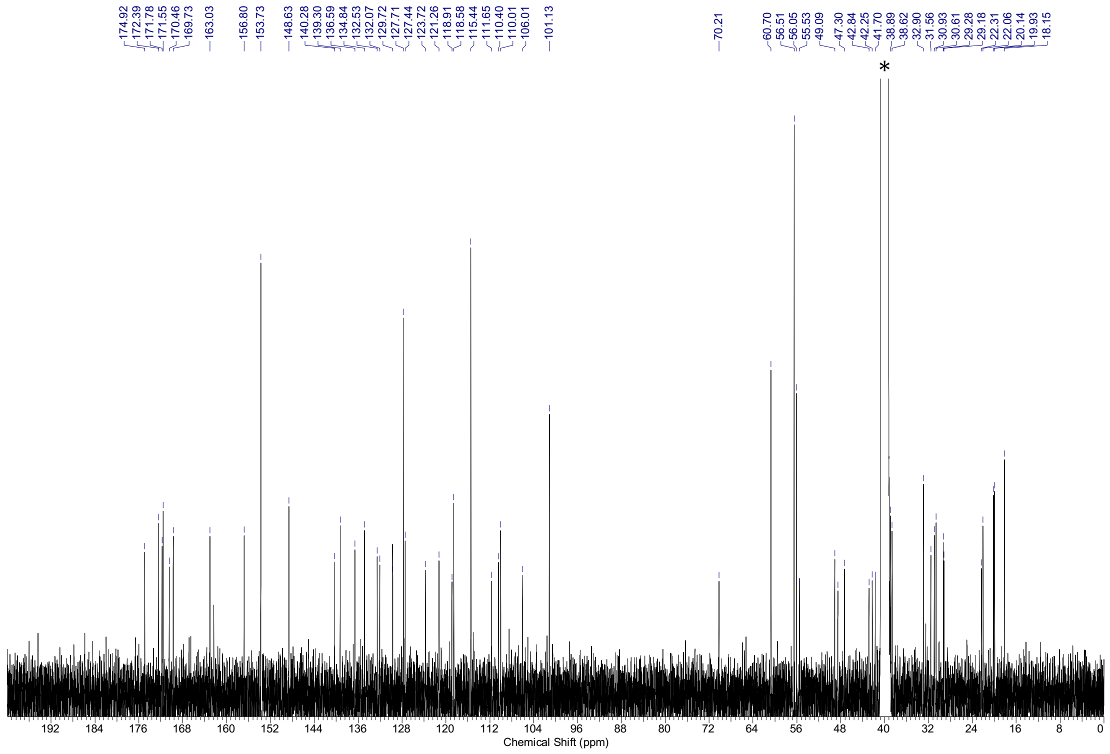

$^1$H-NMR spectra (500 MHz, DMSO-$d_6$) and $^{13}$C-NMR spectra (126 MHz, DMSO-$d_6$) of **S6.** * = NMR-solvent, $H_2O$

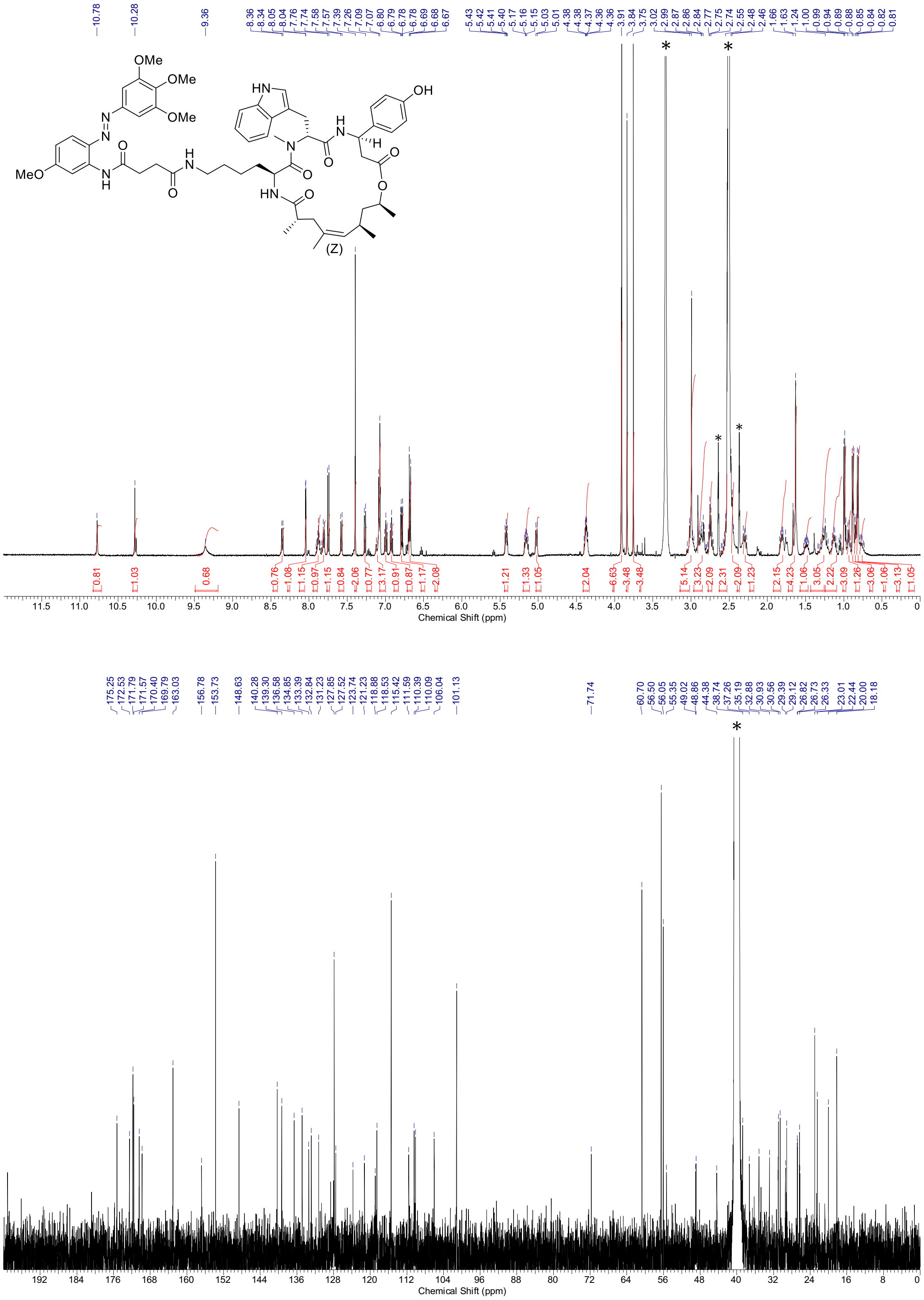

[1]H-NMR spectra (500 MHz, DMSO-$d_6$) and [13]C-NMR spectra (126 MHz, DMSO-$d_6$) of **S7**. * = NMR-solvent, $H_2O$

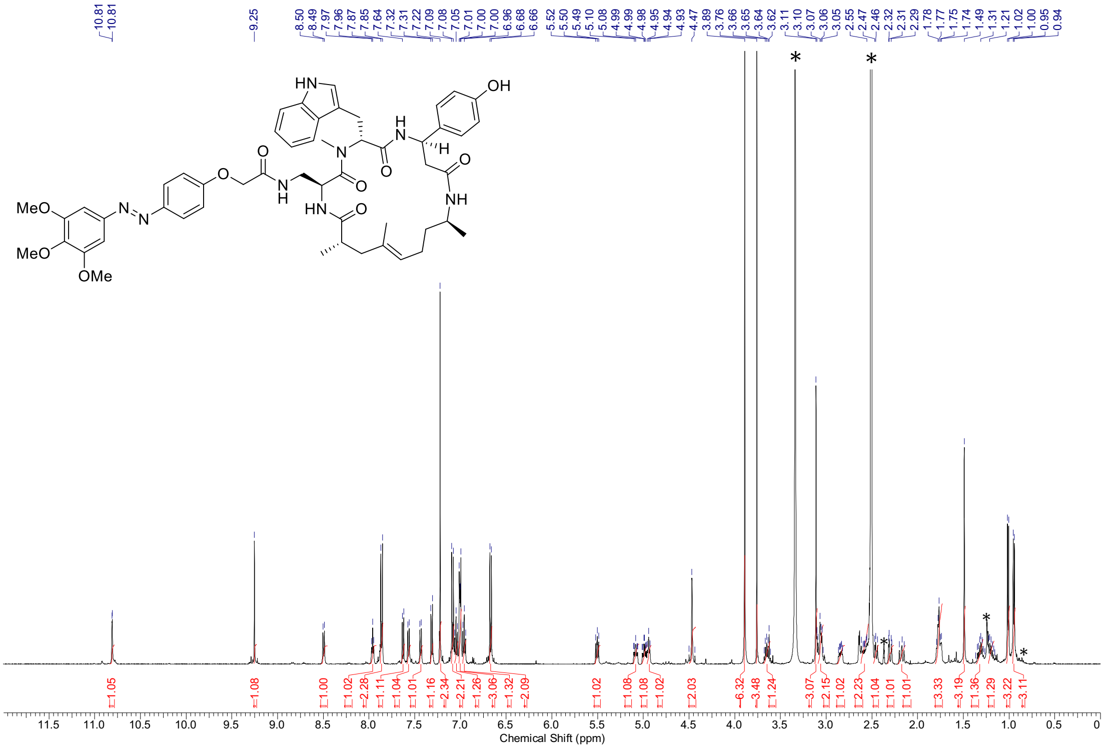

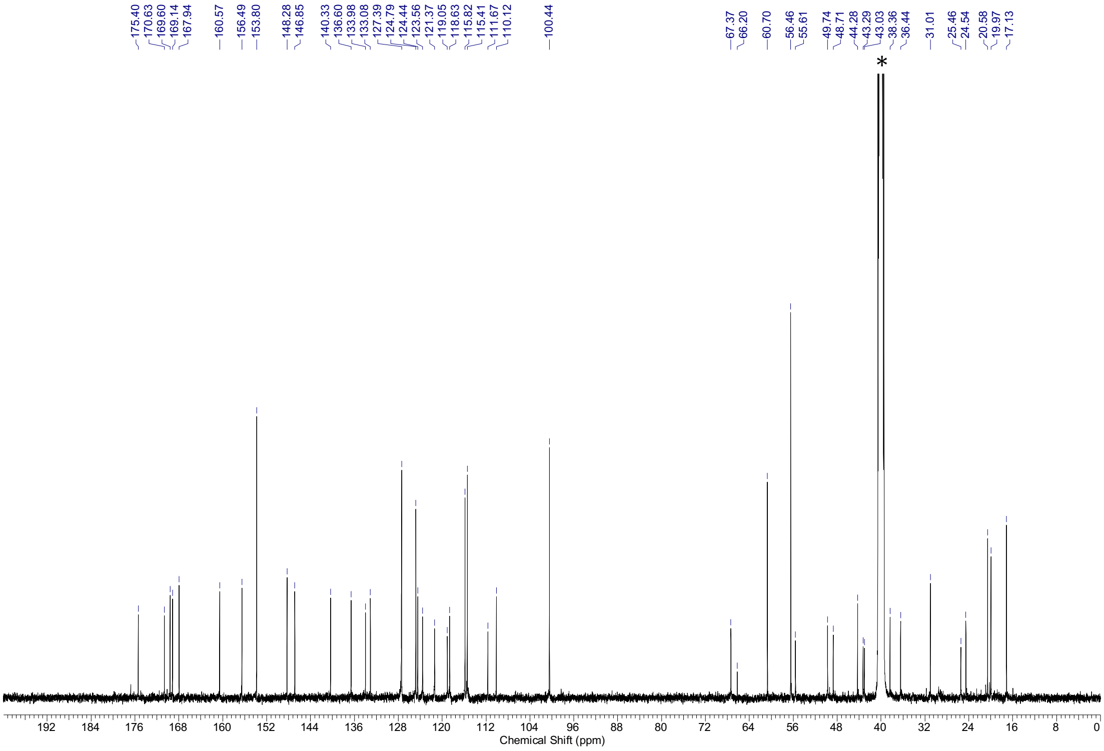

$^{1}$H-NMR spectra (500 MHz, DMSO-$d_6$) and $^{13}$C-NMR spectra (126 MHz, DMSO-$d_6$) of **S8**. * = NMR-solvent, $H_2O$, grease

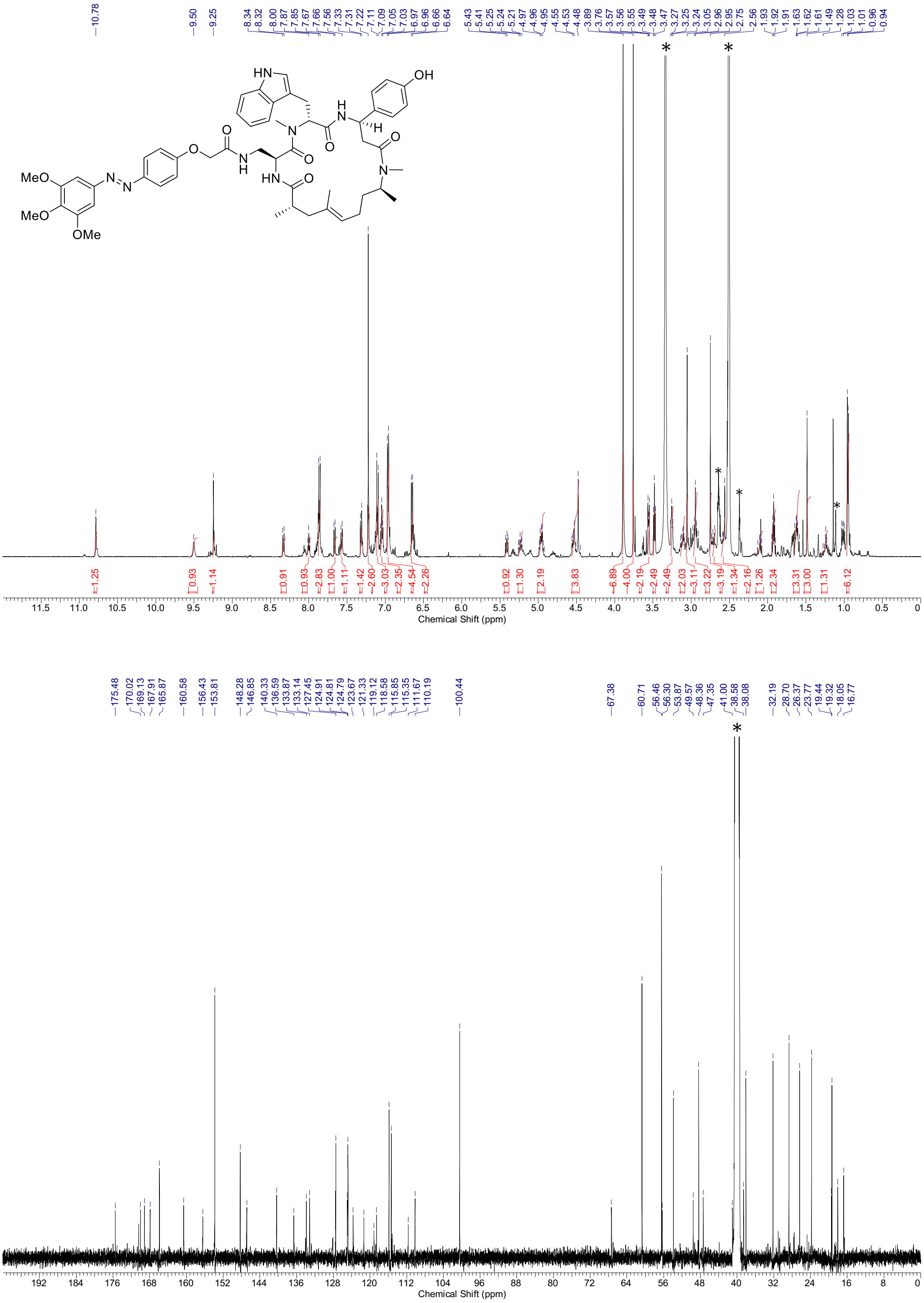

[1]H-NMR spectra (500 MHz, DMSO-$d_6$) and [13]C-NMR spectra (126 MHz, DMSO-$d_6$) of **S9.** * = NMR-solvent, $H_2O$, grease

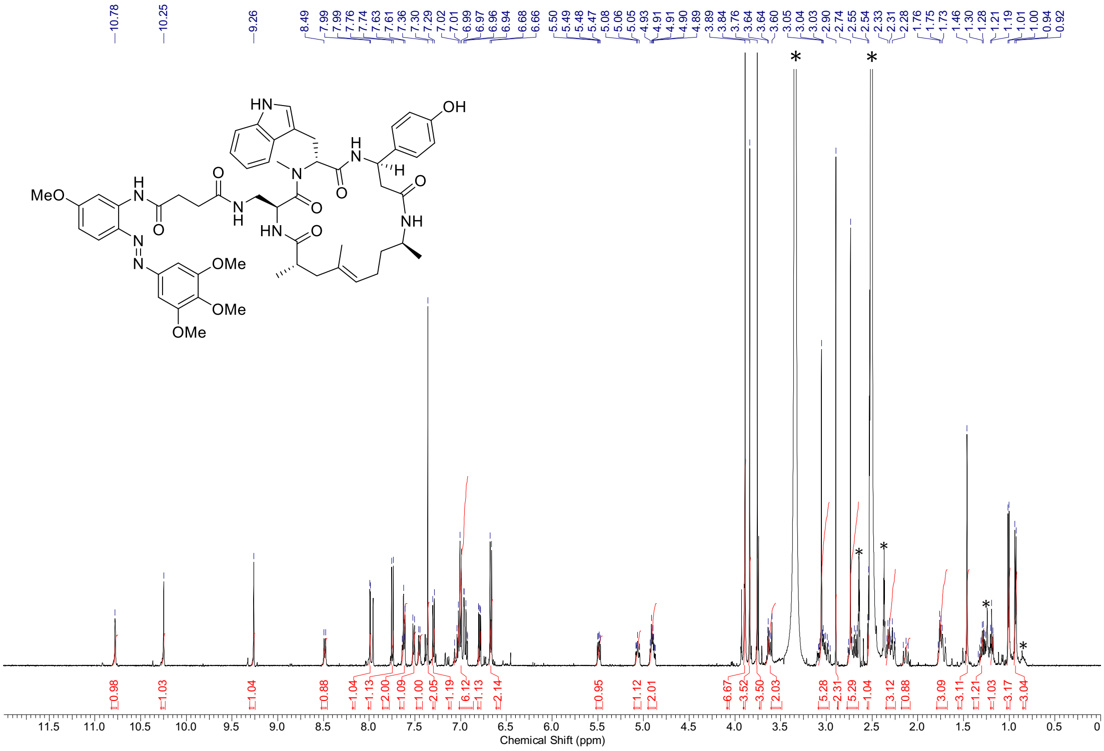

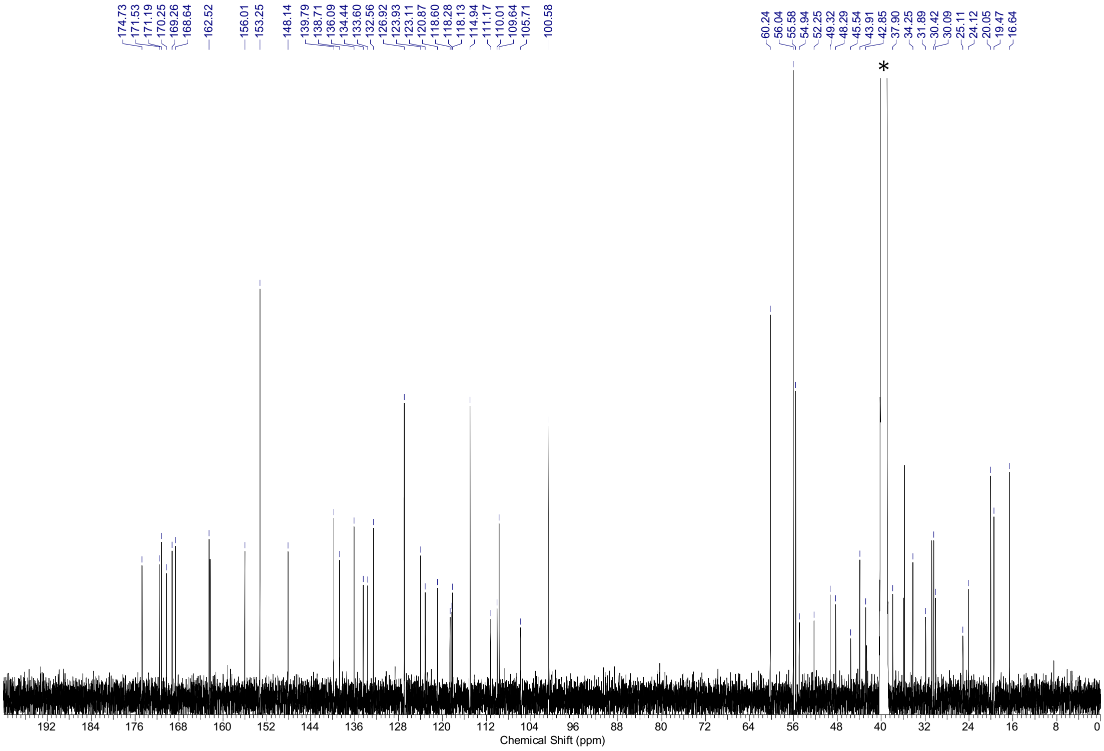

[1]H-NMR spectra (500 MHz, DMSO-$d_6$) and [13]C-NMR spectra (126 MHz, DMSO-$d_6$) of **S10.** * = NMR-solvent, $H_2O$, grease

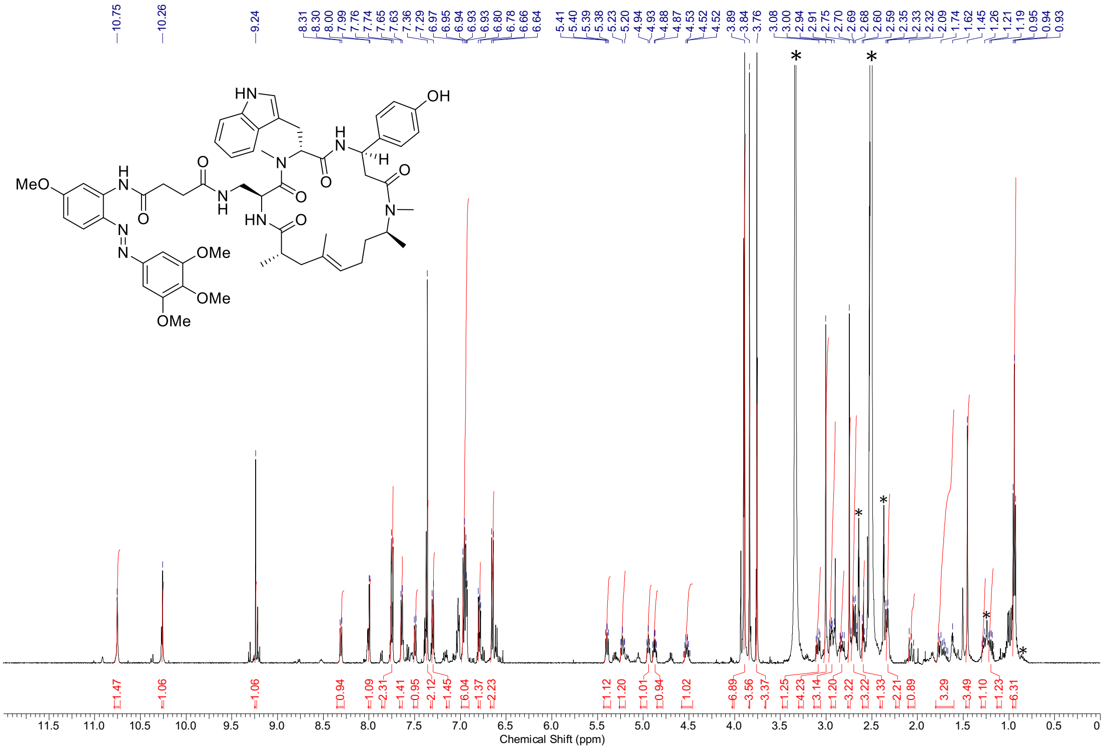

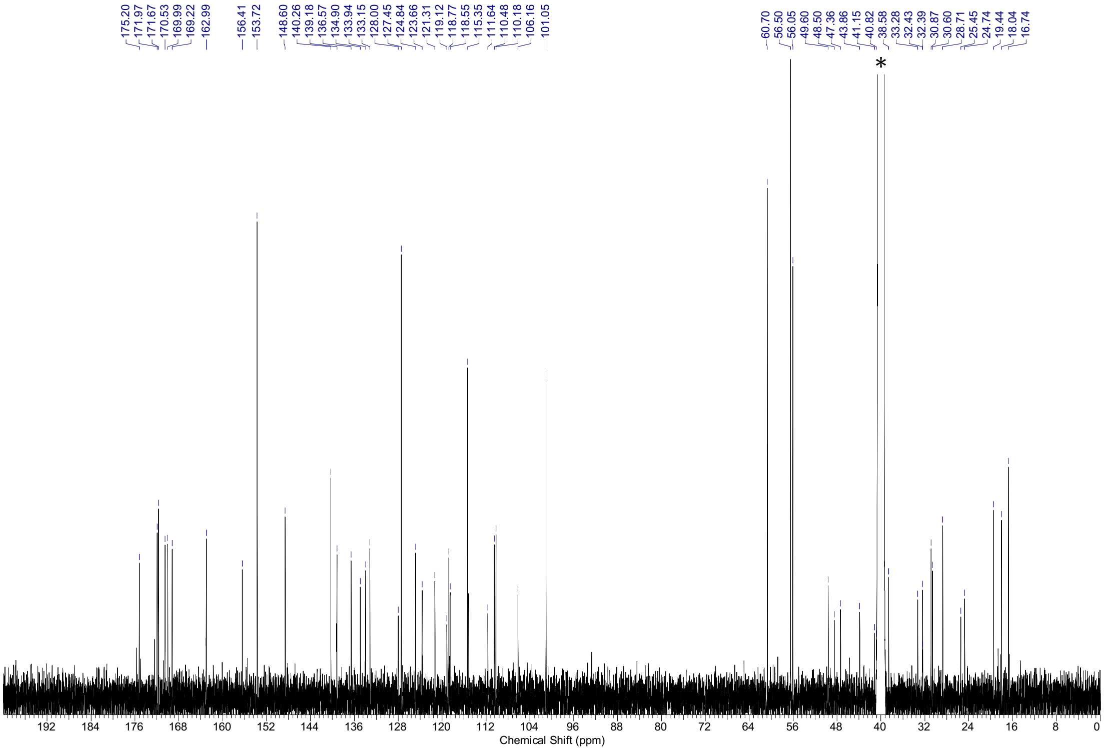

[1]H-NMR spectra (500 MHz, DMSO-*d*$_6$) and [13]C-NMR spectra (126 MHz, DMSO-*d*$_6$) of **S11.** * = NMR-solvent, $H_2O$, grease

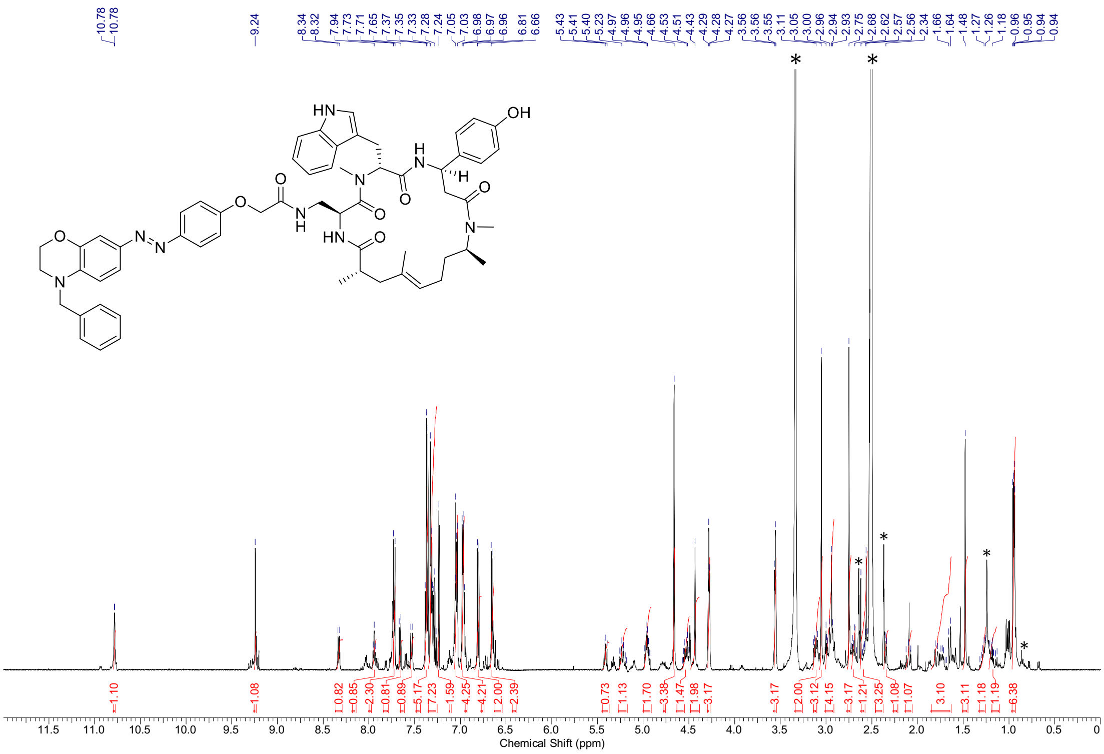

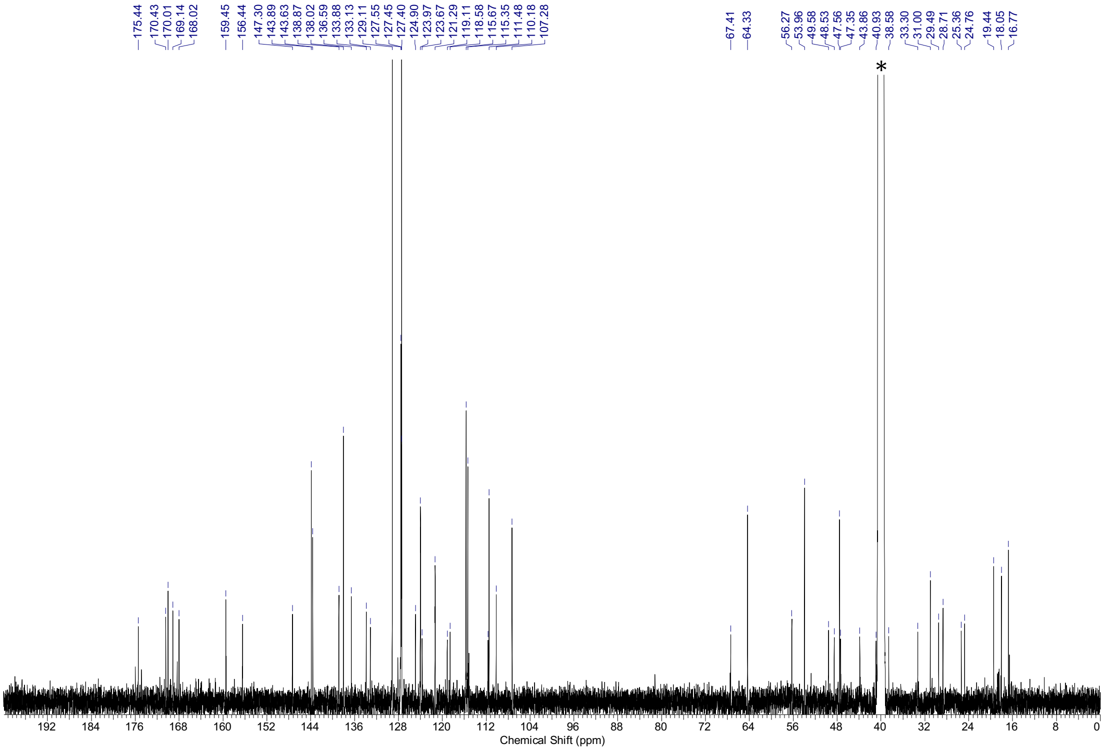

[1]H-NMR spectra (500 MHz, DMSO-$d_6$) and [13]C-NMR spectra (126 MHz, DMSO-$d_6$) of **S12.** * = NMR-solvent, $H_2O$, grease

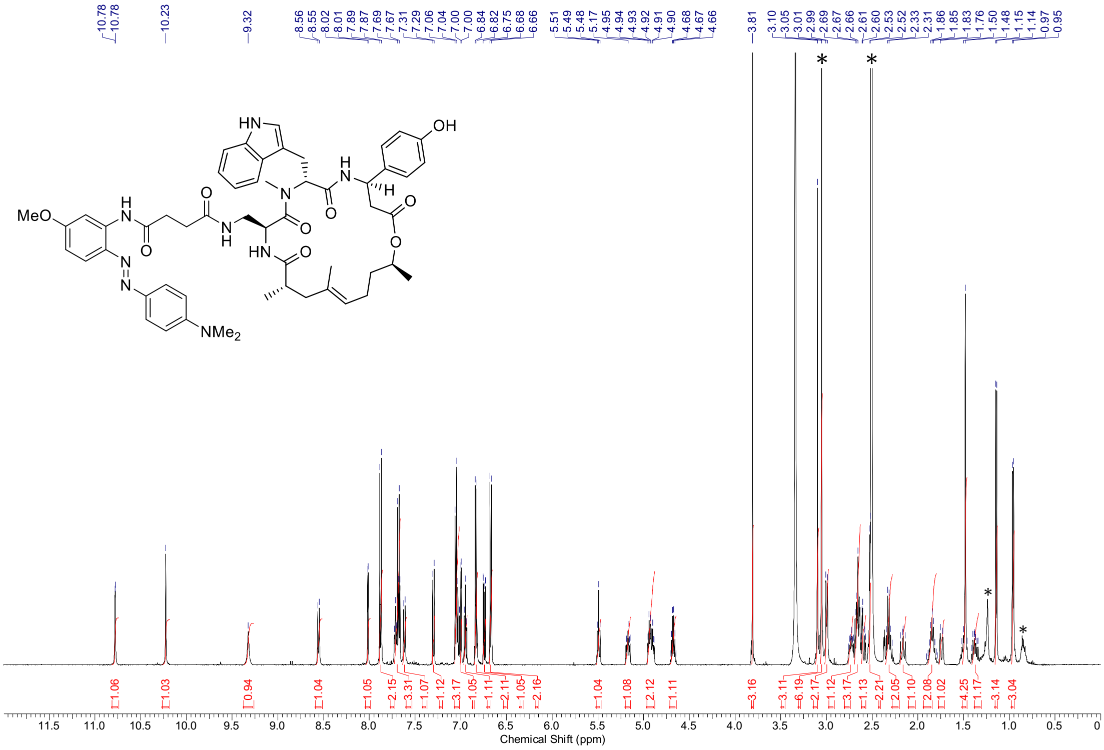

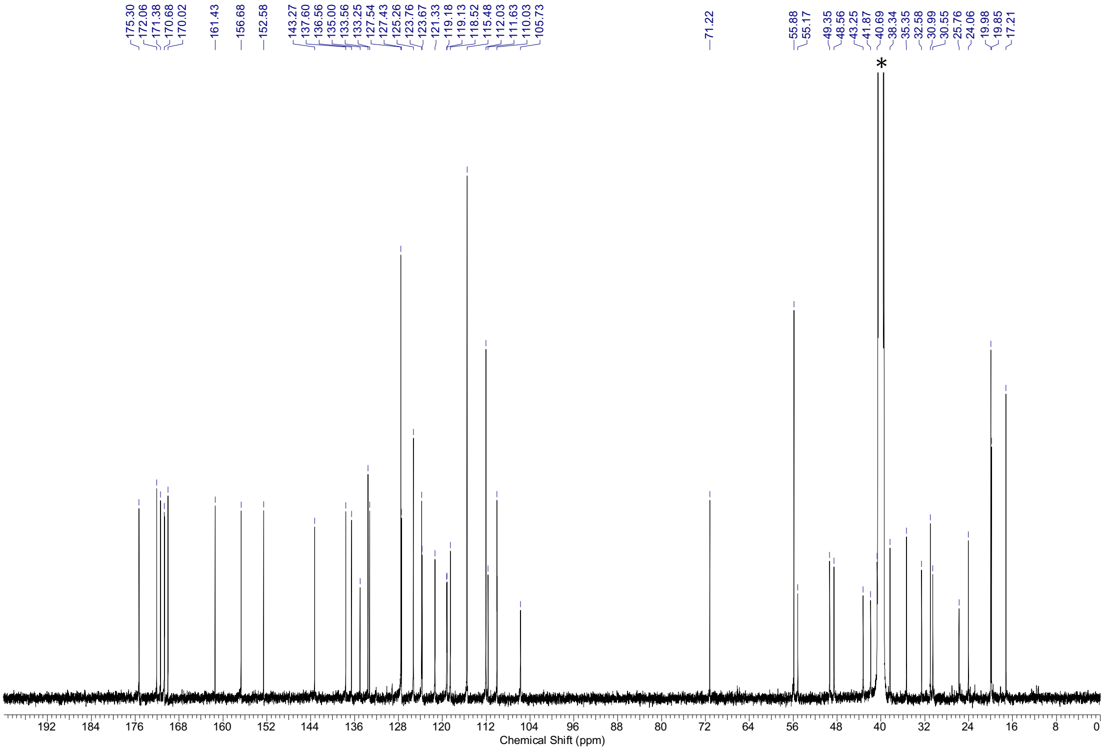

$^{1}$H-NMR spectra (500 MHz, DMSO-$d_6$) and $^{13}$C-NMR spectra (126 MHz, DMSO-$d_6$) of **S17.** * = NMR-solvent, $H_2O$, grease

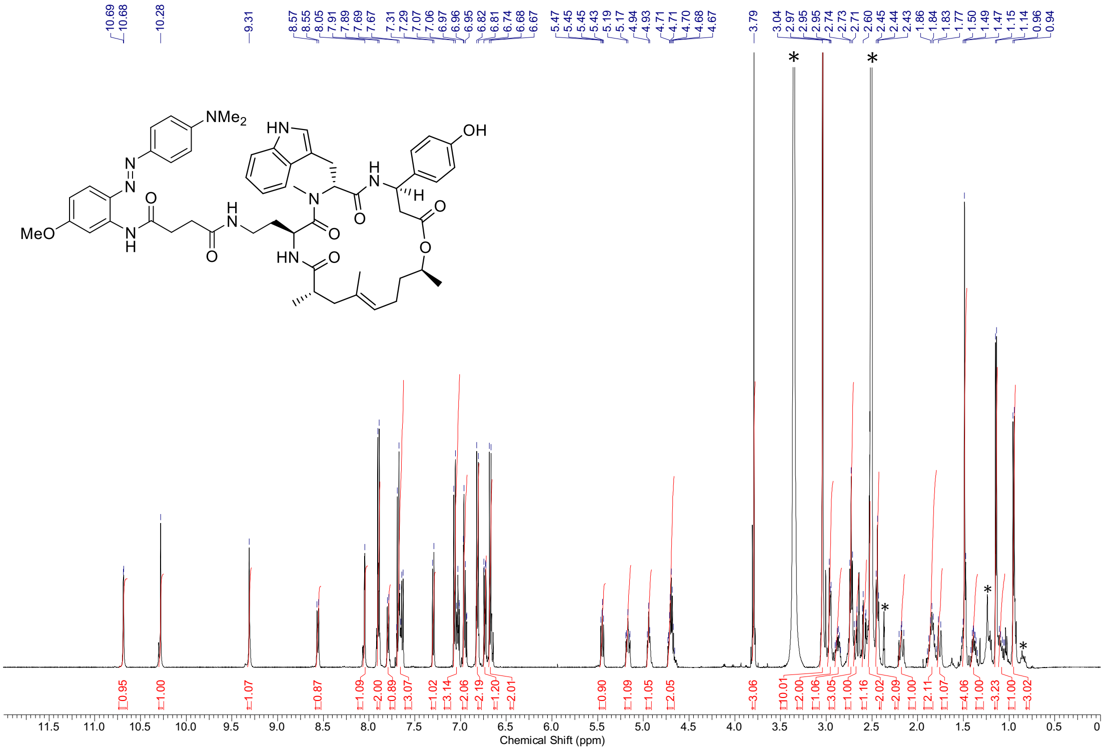

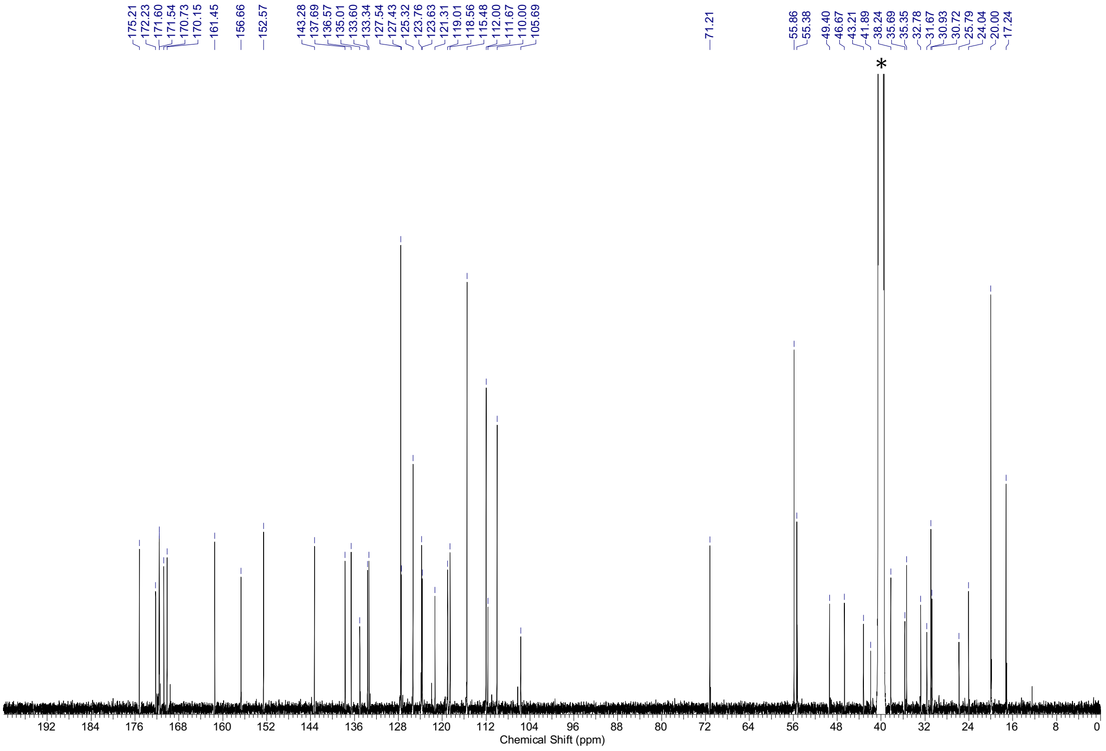

[1]H-NMR spectra (500 MHz, DMSO-$d_6$) and [13]C-NMR spectra (126 MHz, DMSO-$d_6$) of **S18**. * = NMR-solvent, $H_2O$, grease

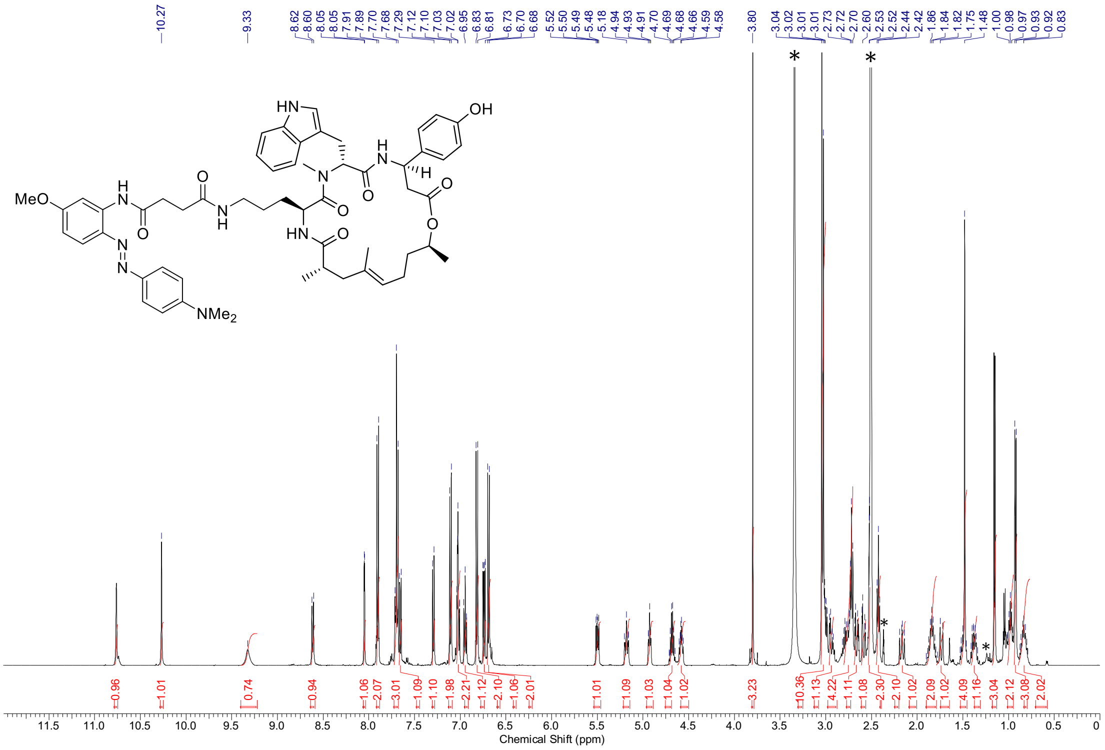

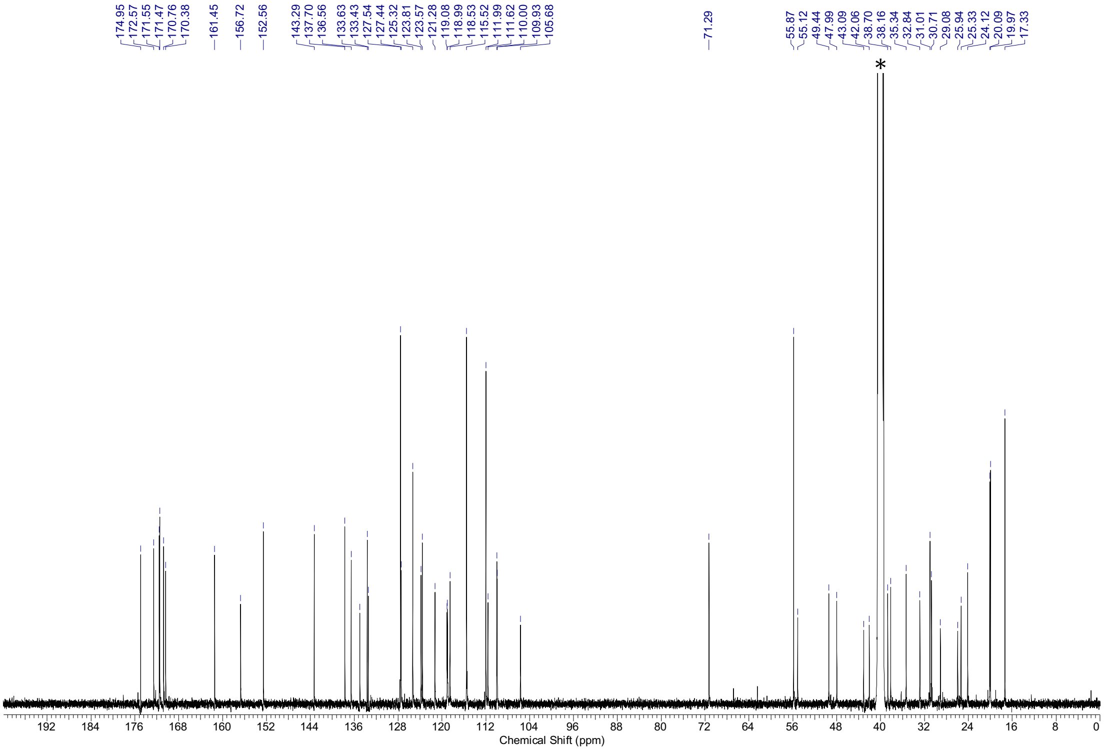

[1]H-NMR spectra (500 MHz, DMSO-*d*6) and [13]C-NMR spectra (126 MHz, DMSO-*d*6) of **S19**. * = NMR-solvent, $H_2O$, grease

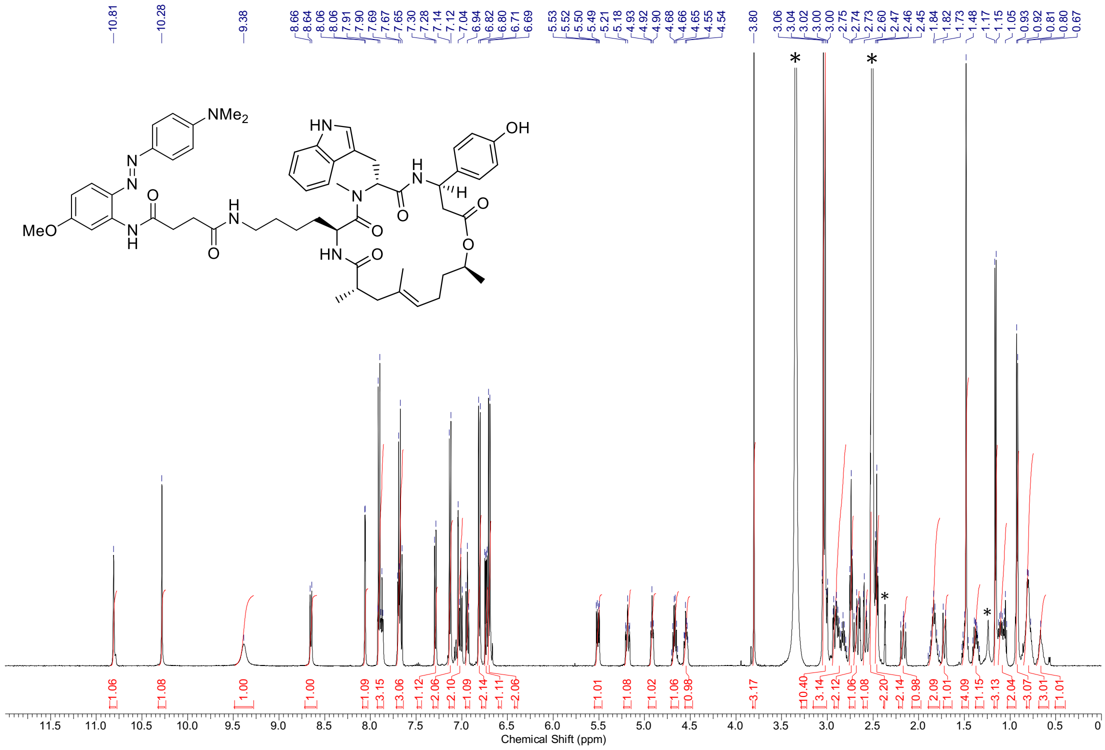

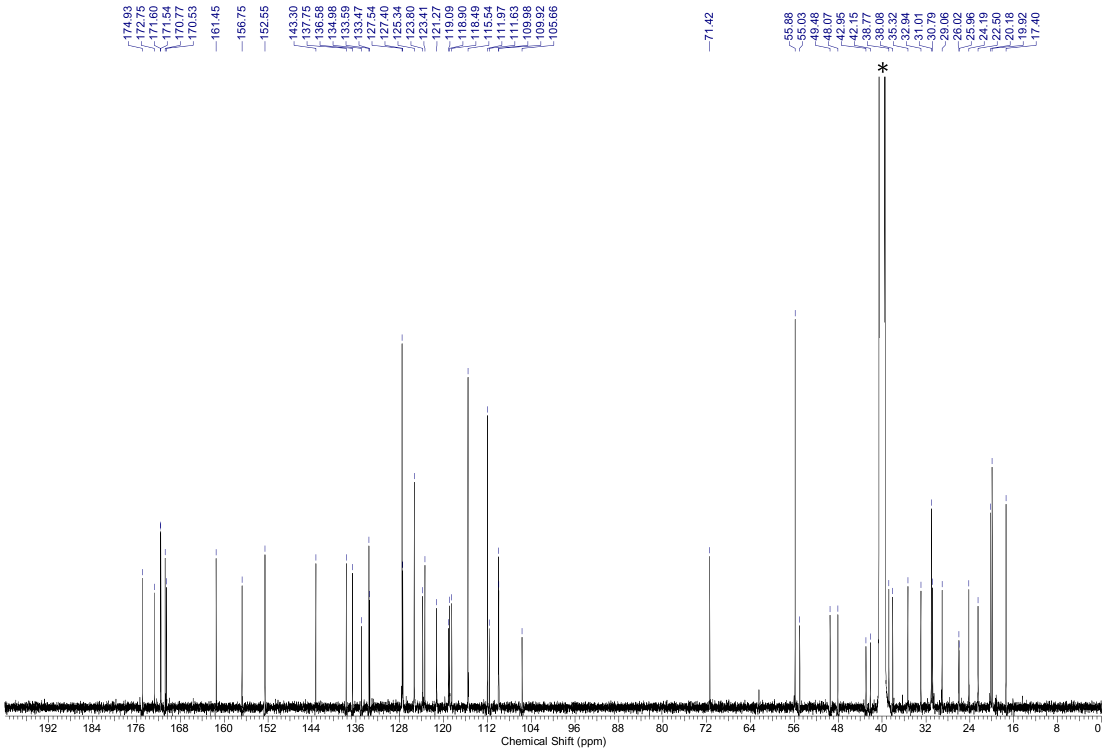

$^1$H-NMR spectra (500 MHz, DMSO-$d_6$) and $^{13}$C-NMR spectra (126 MHz, DMSO-$d_6$) of **S20**. * = NMR-solvent, $H_2O$, grease

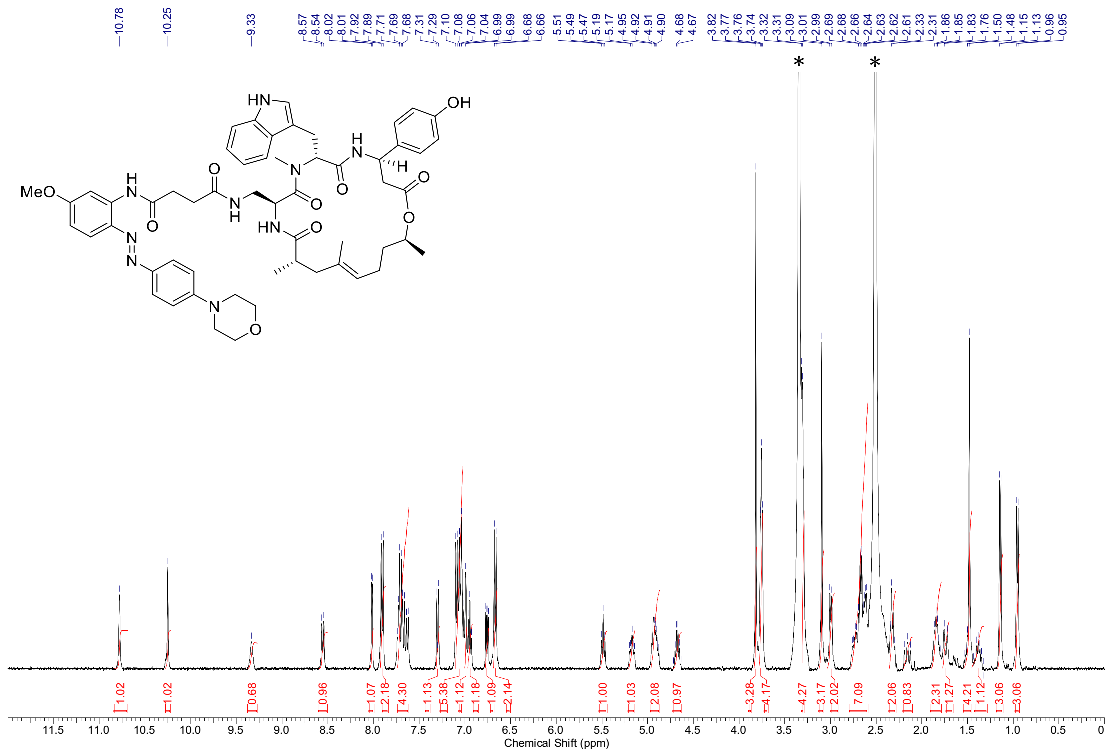

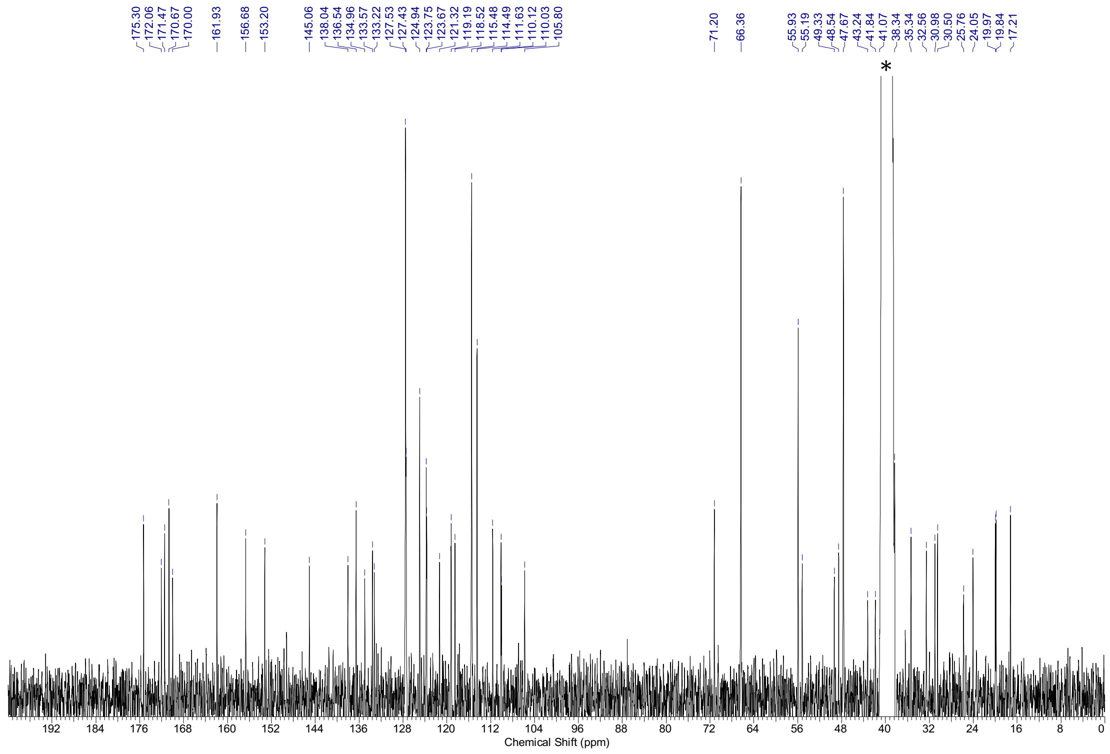

$^1$H-NMR spectra (400 MHz, DMSO-$d_6$) and $^{13}$C-NMR spectra (101 MHz, DMSO-$d_6$) of **S21**. * = NMR-solvent, $H_2O$

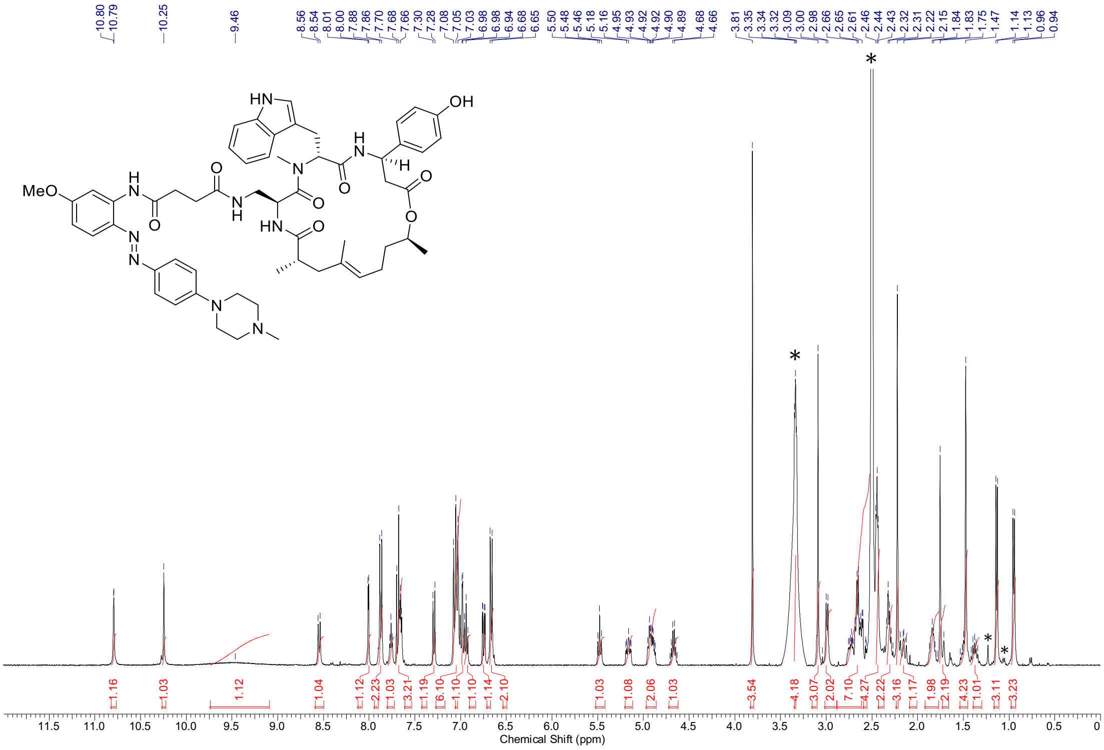

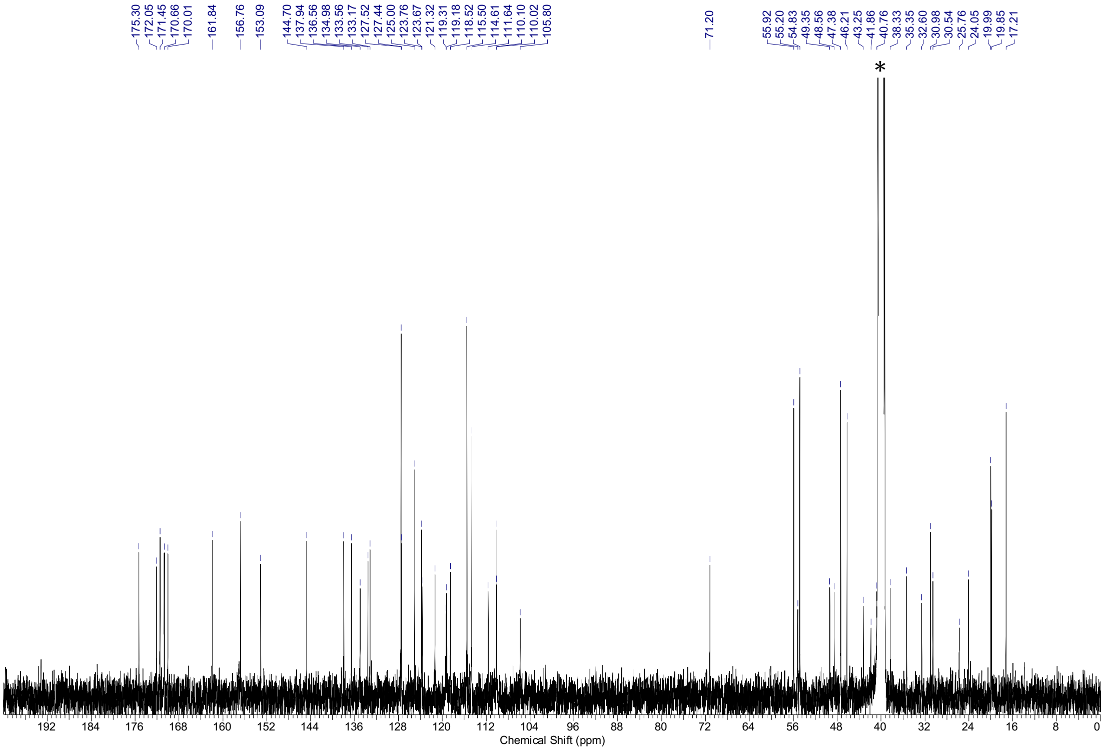

[1]H-NMR spectra (400 MHz, DMSO-$d_6$) and [13]C-NMR spectra (101 MHz, DMSO-$d_6$) of **S22.** * = NMR-solvent, $H_2O$, grease

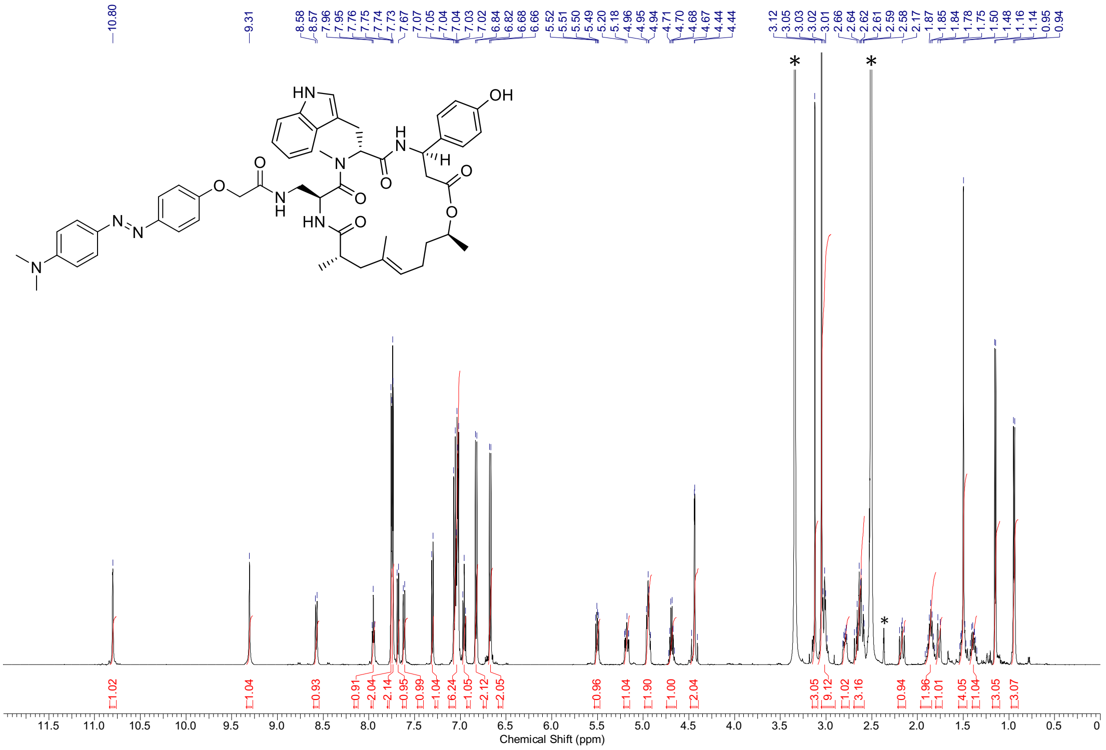

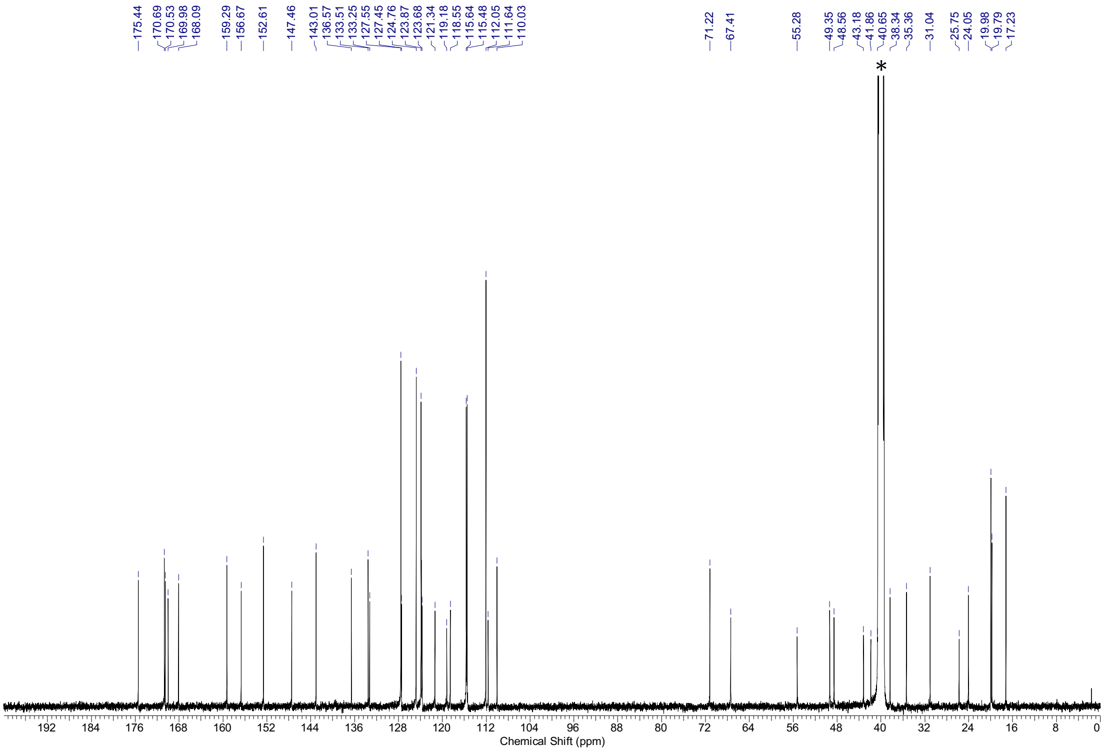

$^{1}H$-NMR spectra (500 MHz, DMSO-$d_6$) and $^{13}C$-NMR spectra (126 MHz, DMSO-$d_6$) of **S23.** * = NMR-solvent, $H_2O$

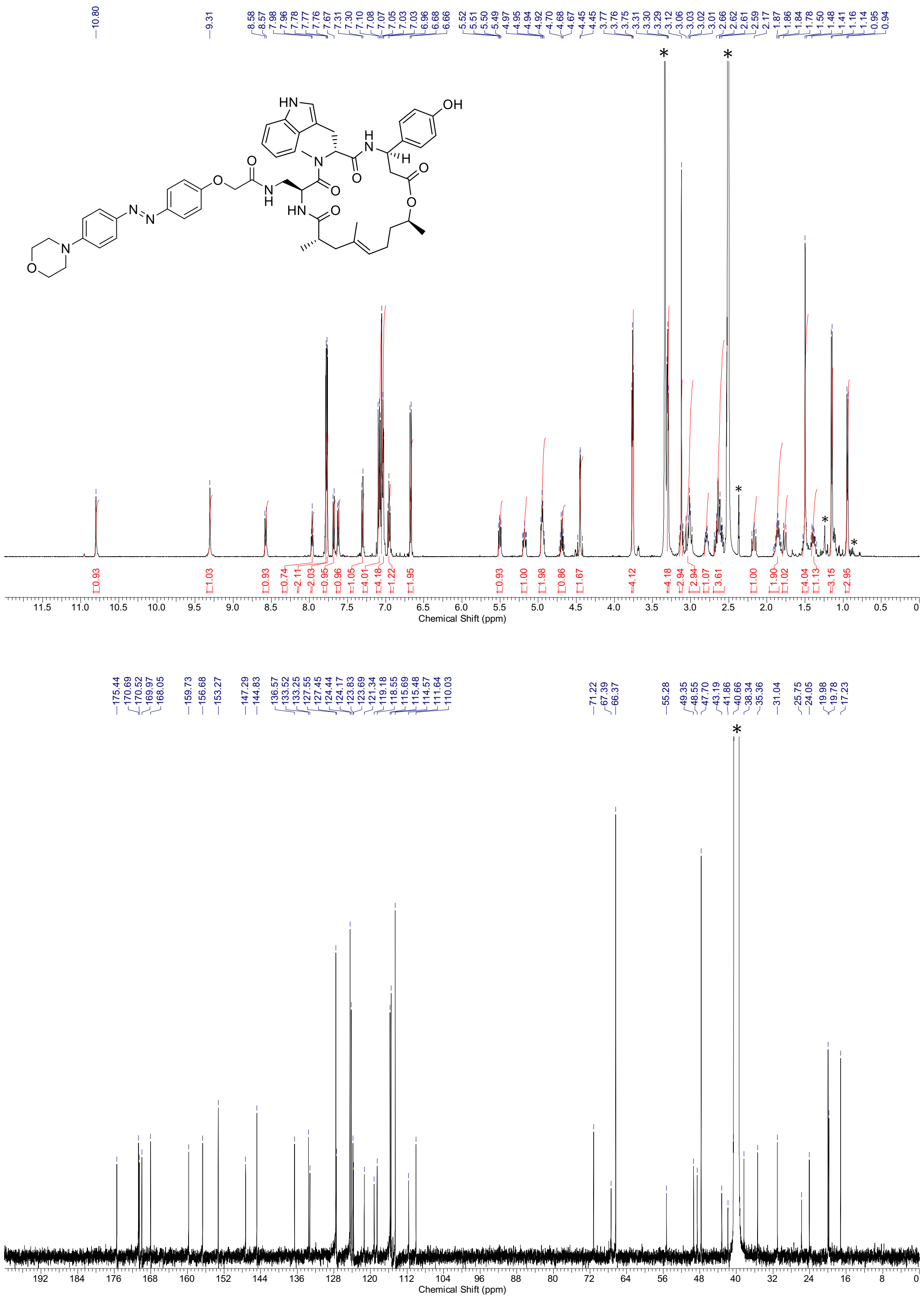

$^{1}H$-NMR spectra (500 MHz, DMSO-$d_6$) and $^{13}C$-NMR spectra (126 MHz, DMSO-$d_6$) of **S24**. * = NMR-solvent, $H_2O$, grease

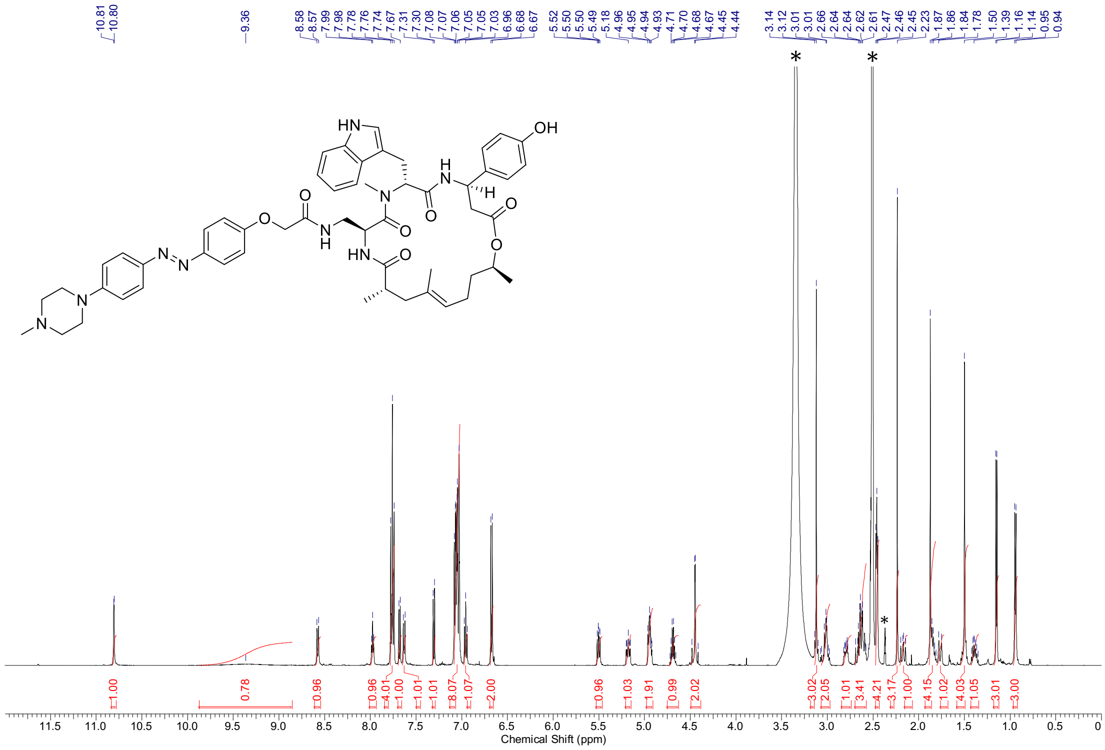

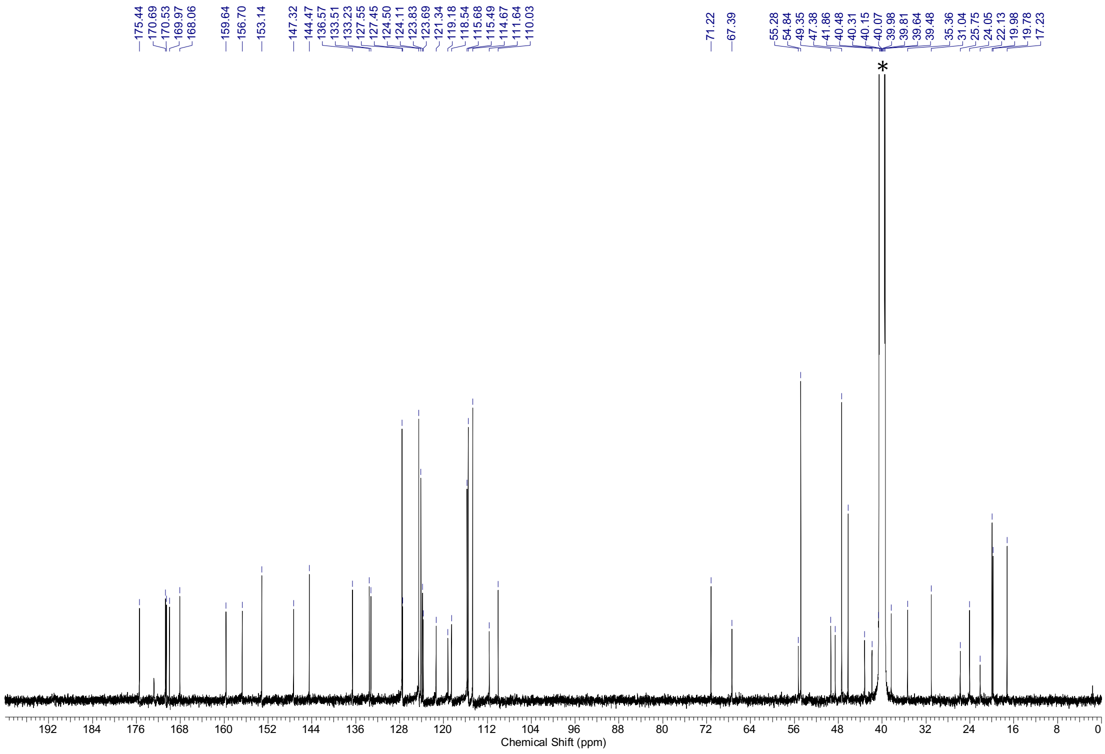

[1]H-NMR spectra (500 MHz, DMSO-$d_6$) and [13]C-NMR spectra (126 MHz, DMSO-$d_6$) of **S25.** * = NMR-solvent, $H_2O$

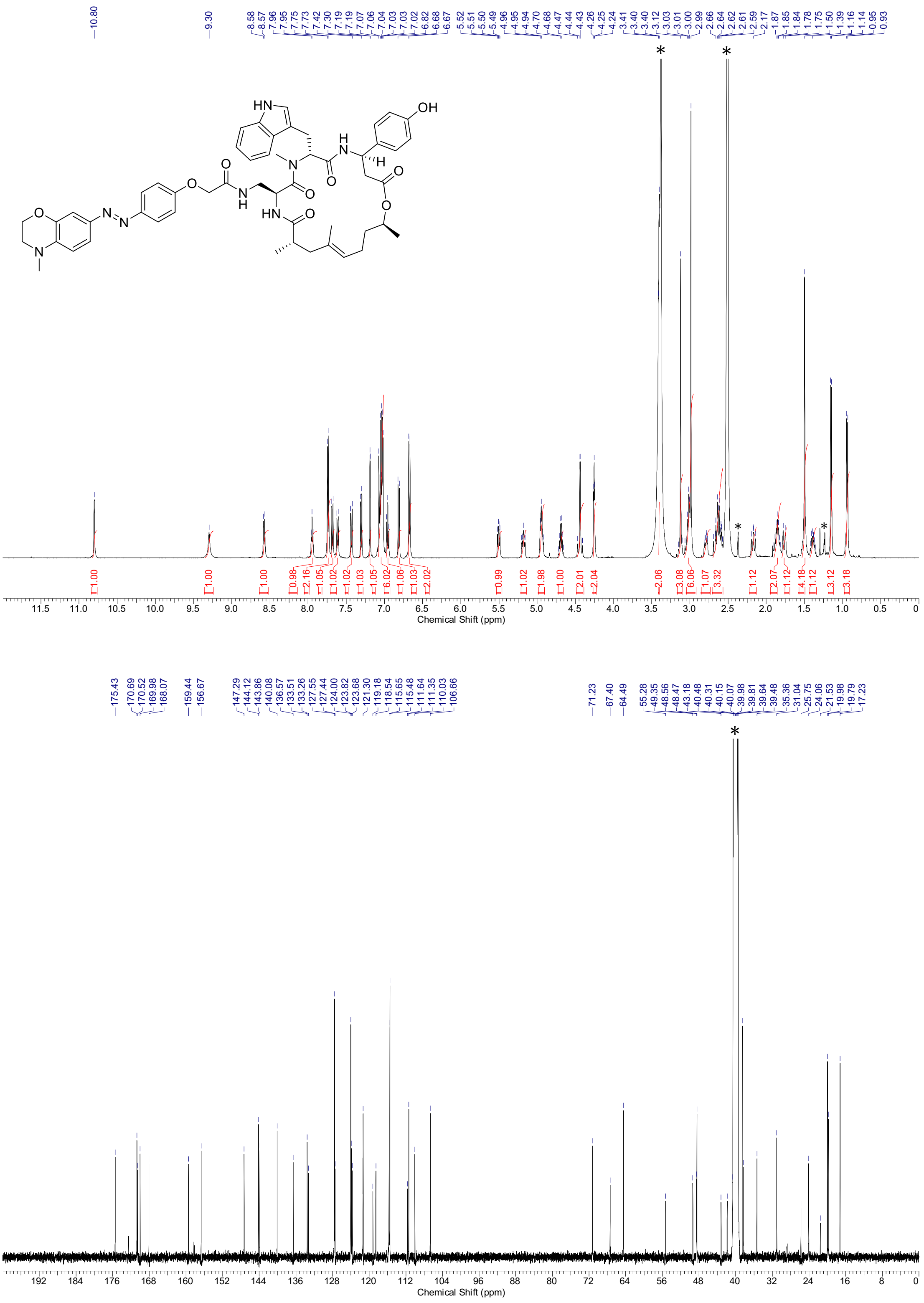

$^{1}$H-NMR spectra (500 MHz, DMSO-$d_6$) and $^{13}$C-NMR spectra (126 MHz, DMSO-$d_6$) of **S26.** * = NMR-solvent, $H_2O$, grease

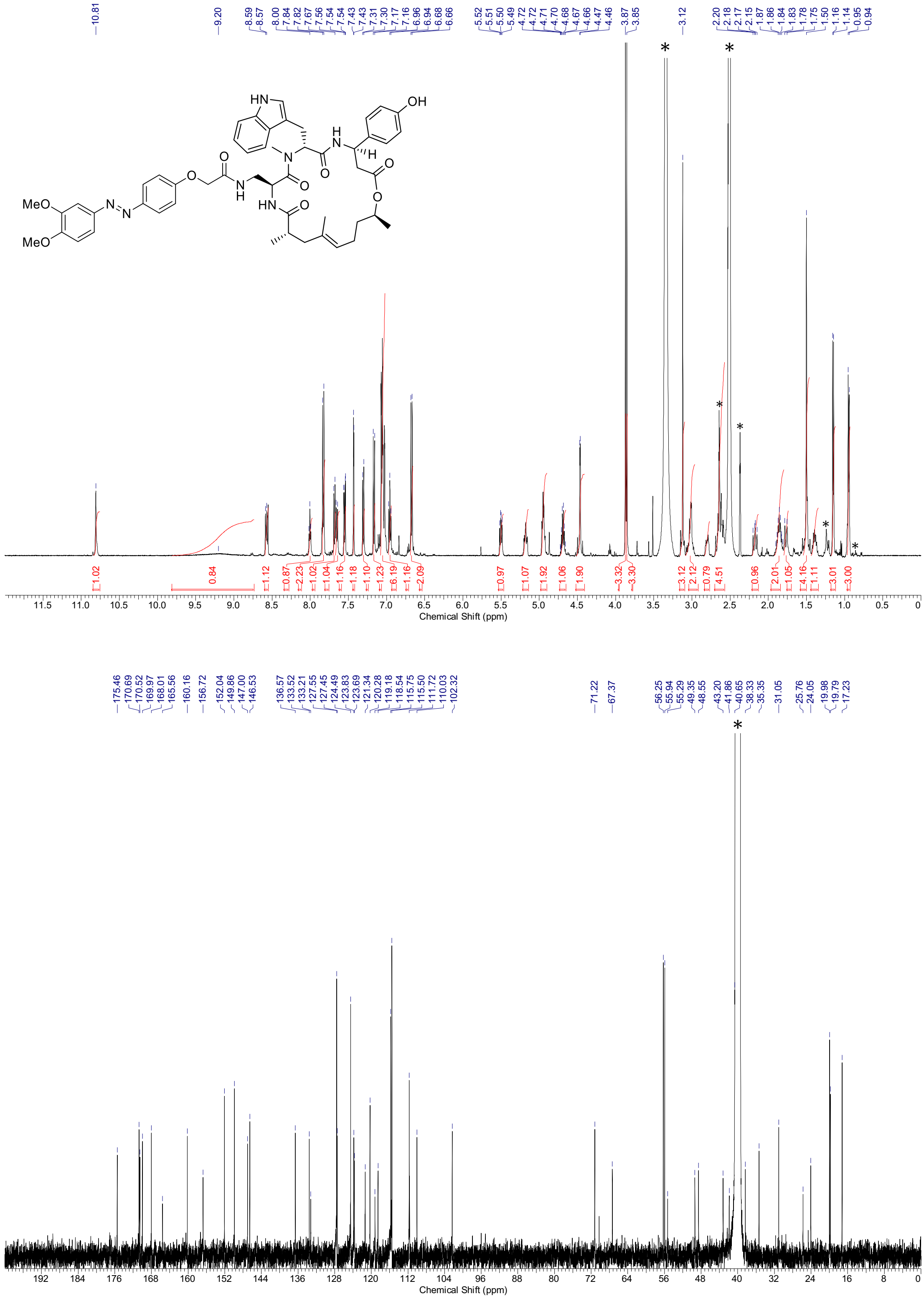

[1]H-NMR spectra (500 MHz, DMSO-$d_6$) and [13]C-NMR spectra (126 MHz, DMSO-$d_6$) of **S27**. * = NMR-solvent, $H_2O$, grease

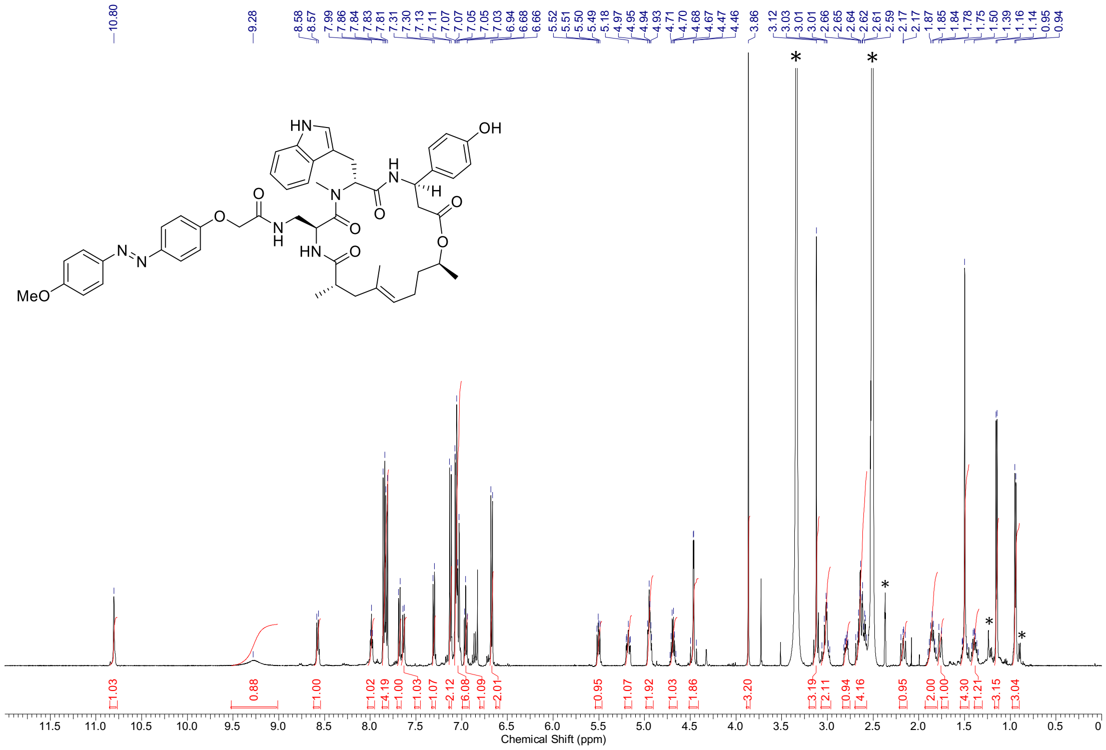

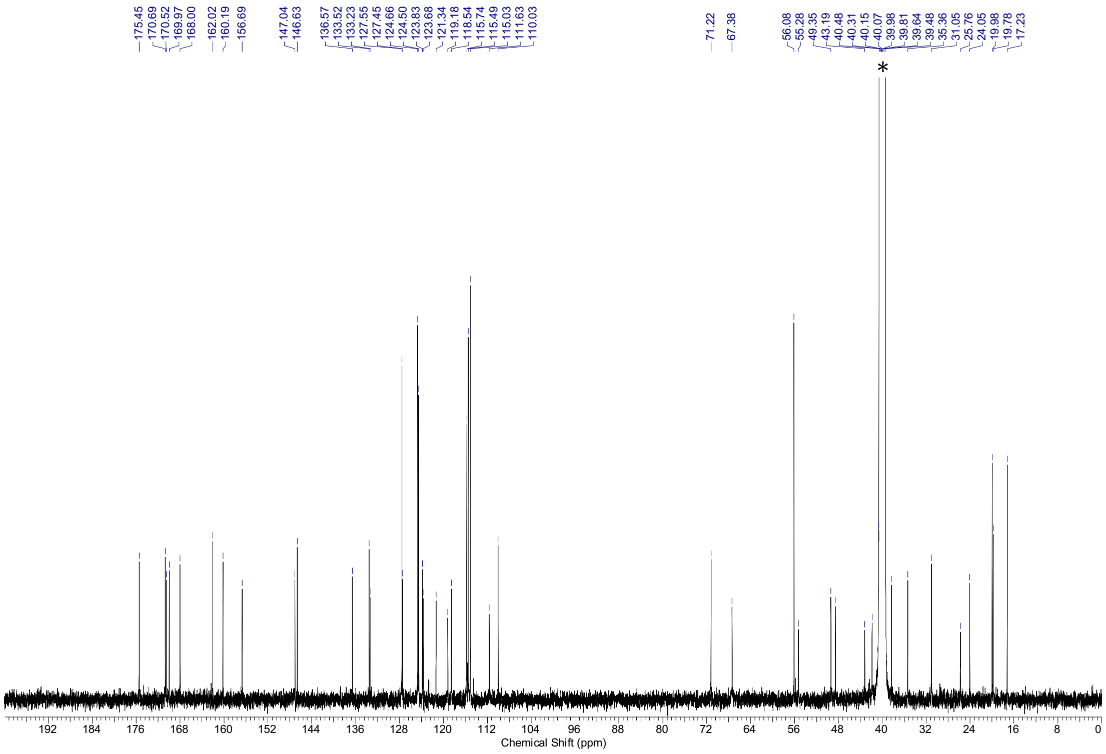

[1]H-NMR spectra (500 MHz, DMSO-*d*6) and [13]C-NMR spectra (126 MHz, DMSO-*d*6) of **S28.** * = NMR-solvent, $H_2O$, grease

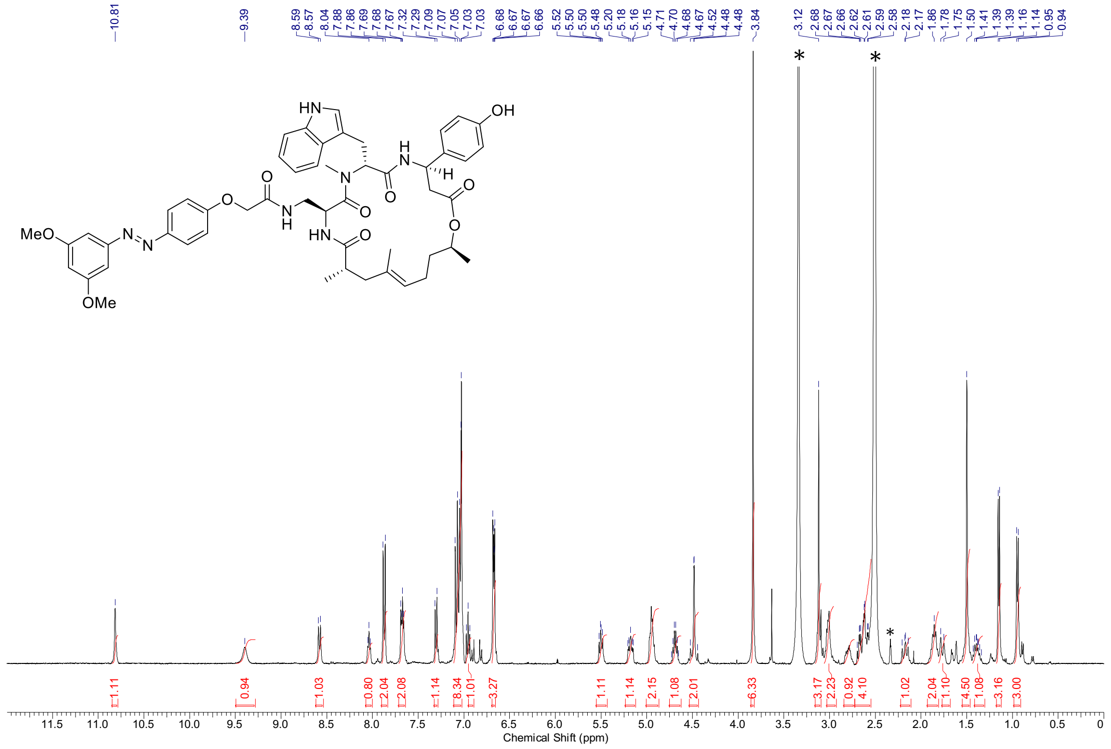

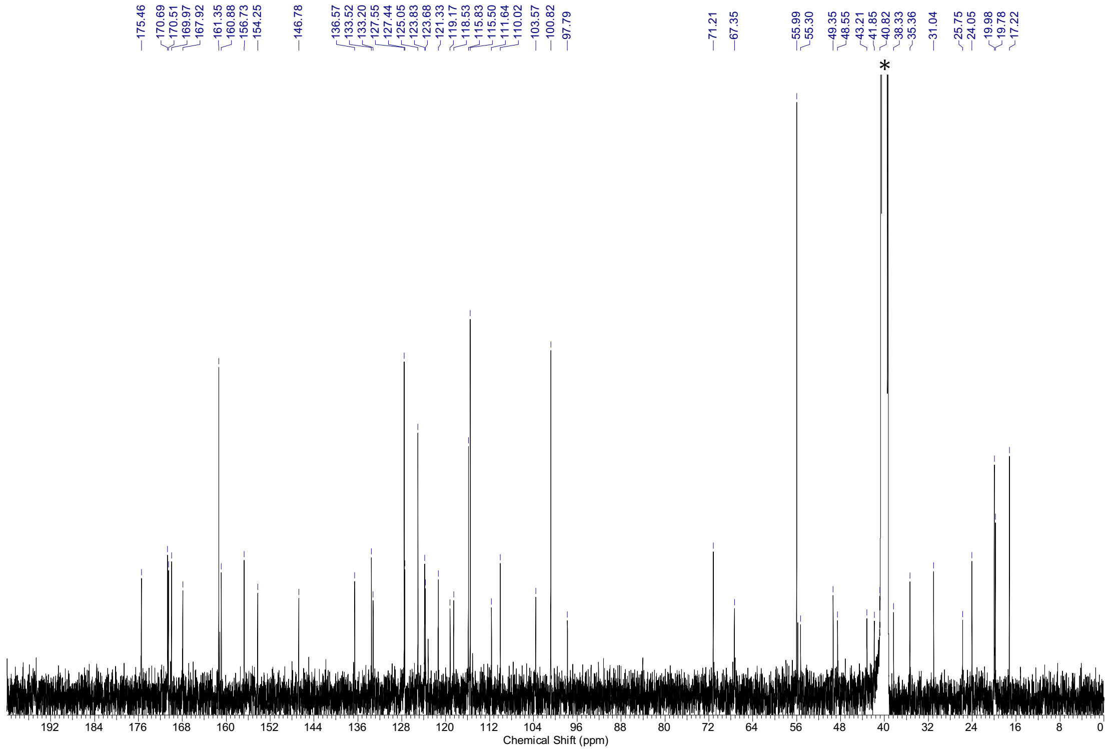

[1]H-NMR spectra (400 MHz, DMSO-$d_6$) and [13]C-NMR spectra (101 MHz, DMSO-$d_6$) of **S29.** * = NMR-solvent, $H_2O$

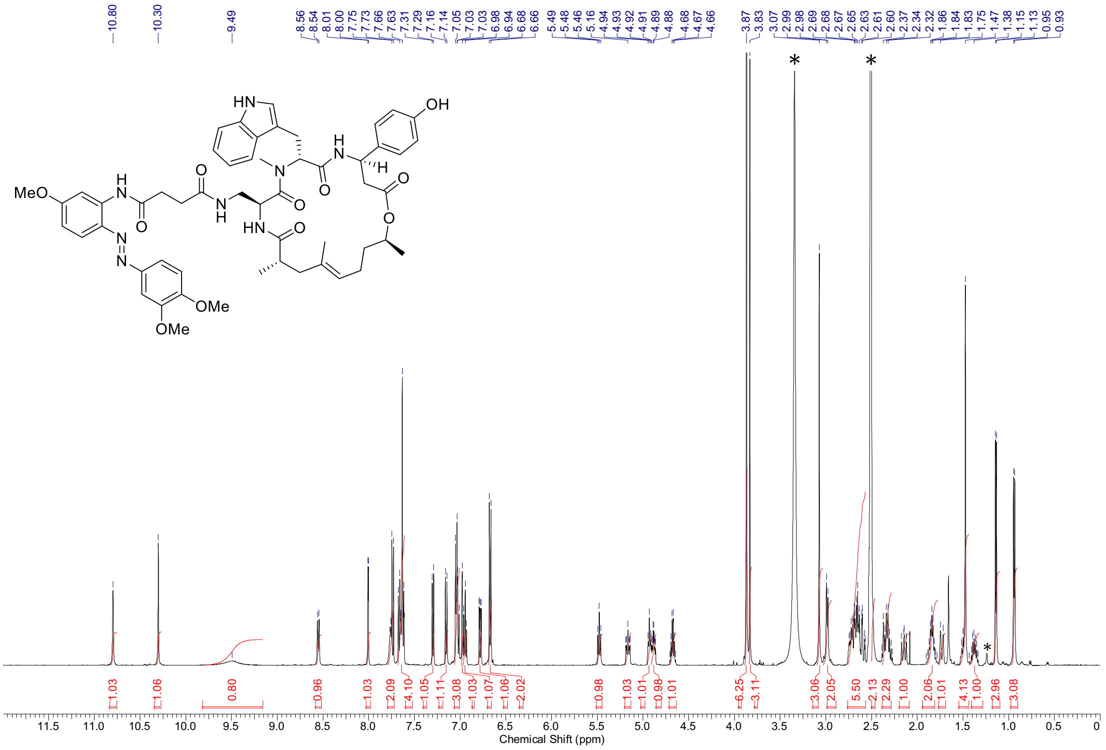

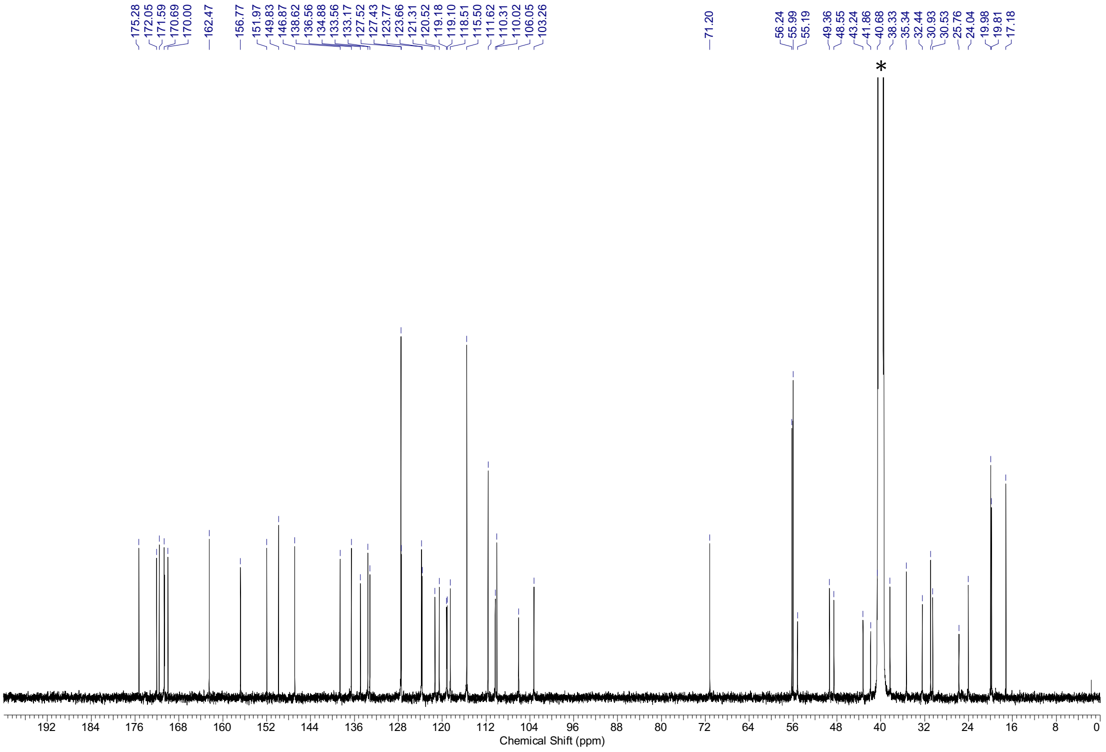

$^{1}H$-NMR spectra (500 MHz, DMSO-$d_6$) and $^{13}C$-NMR spectra (126 MHz, DMSO-$d_6$) of **S30.** * = NMR-solvent, $H_2O$, grease

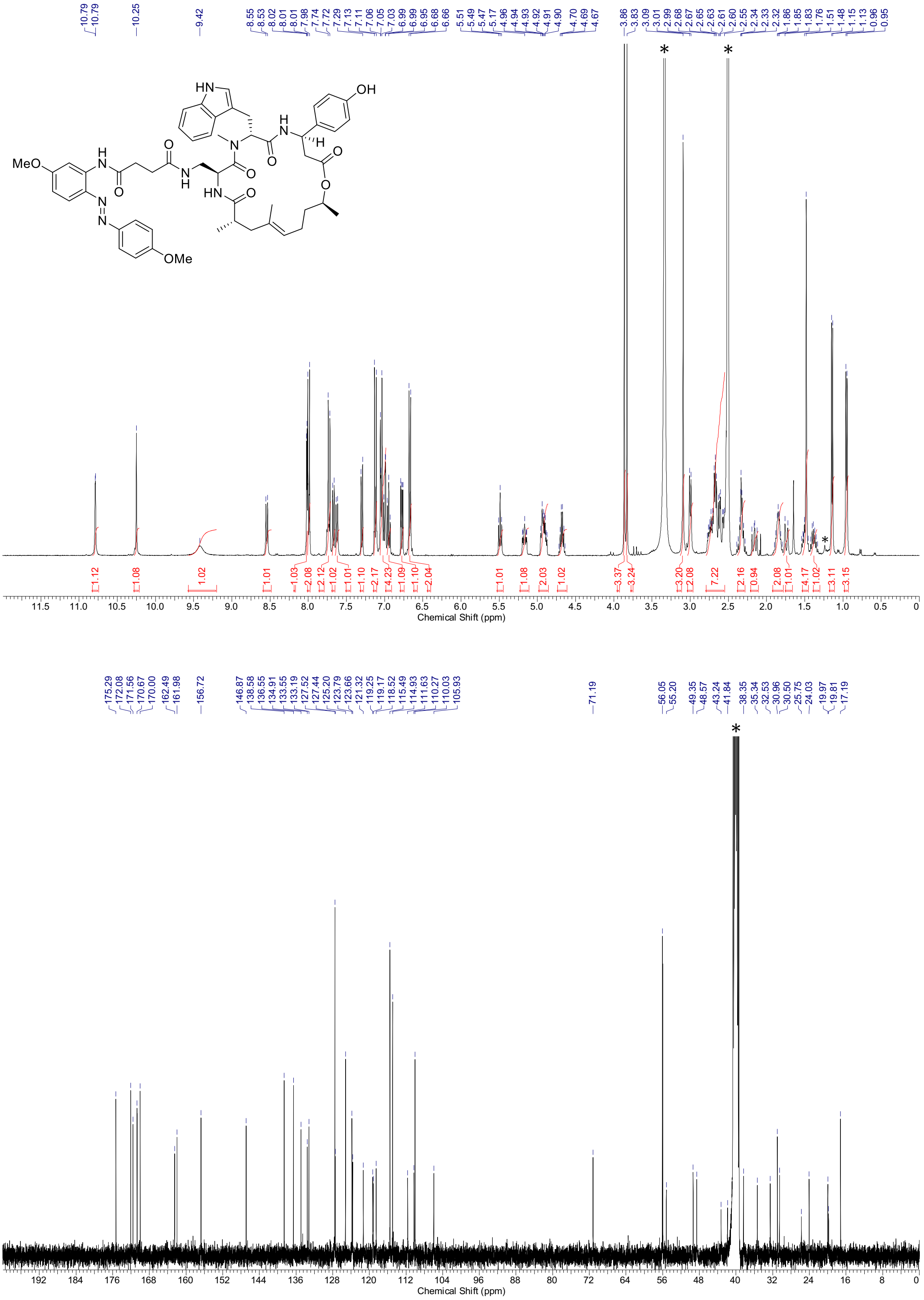

$^{1}$H-NMR spectra (400 MHz, DMSO-$d_6$) and $^{13}$C-NMR spectra (101 MHz, DMSO-$d_6$) of **S31.** * = NMR-solvent, $H_2O$, grease

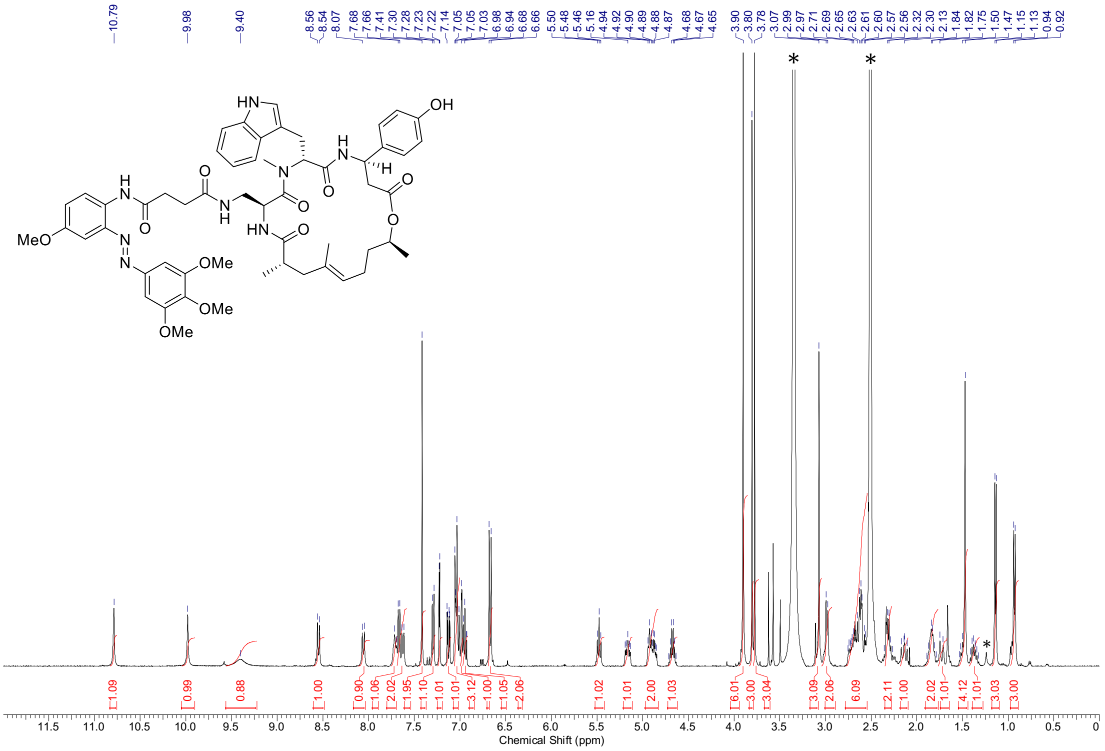

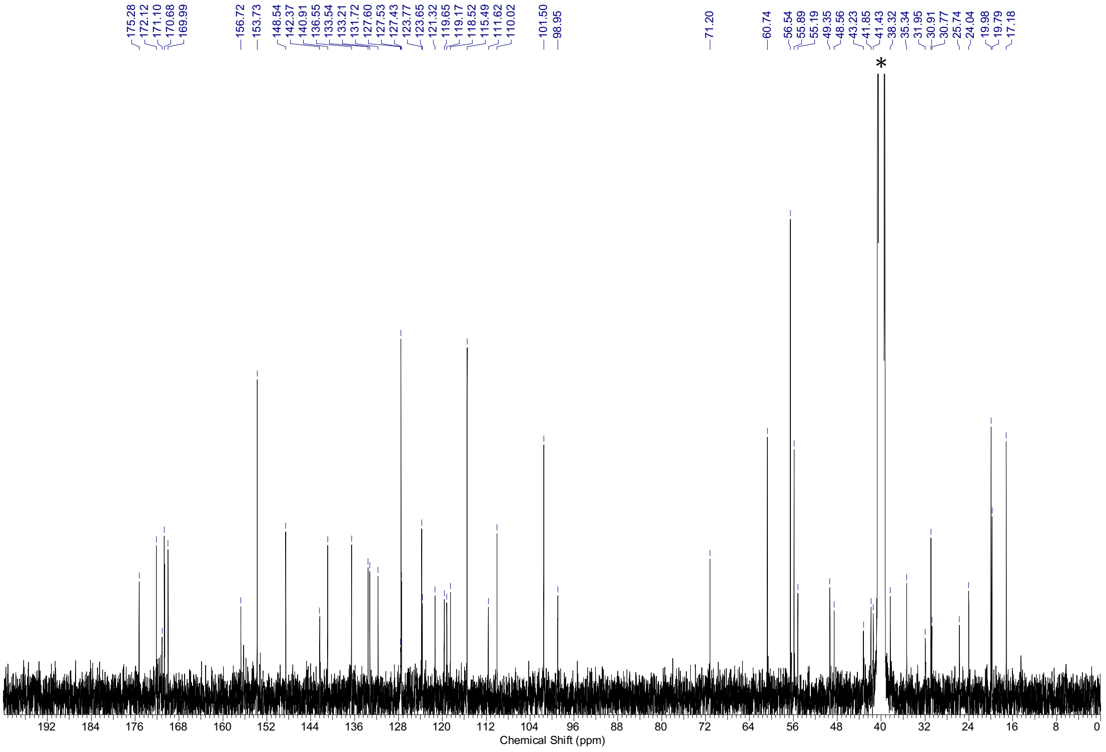

$^{1}$H-NMR spectra (400 MHz, DMSO-$d_6$) and $^{13}$C-NMR spectra (101 MHz, DMSO-$d_6$) of **S32.** * = NMR-solvent, $H_2O$, grease

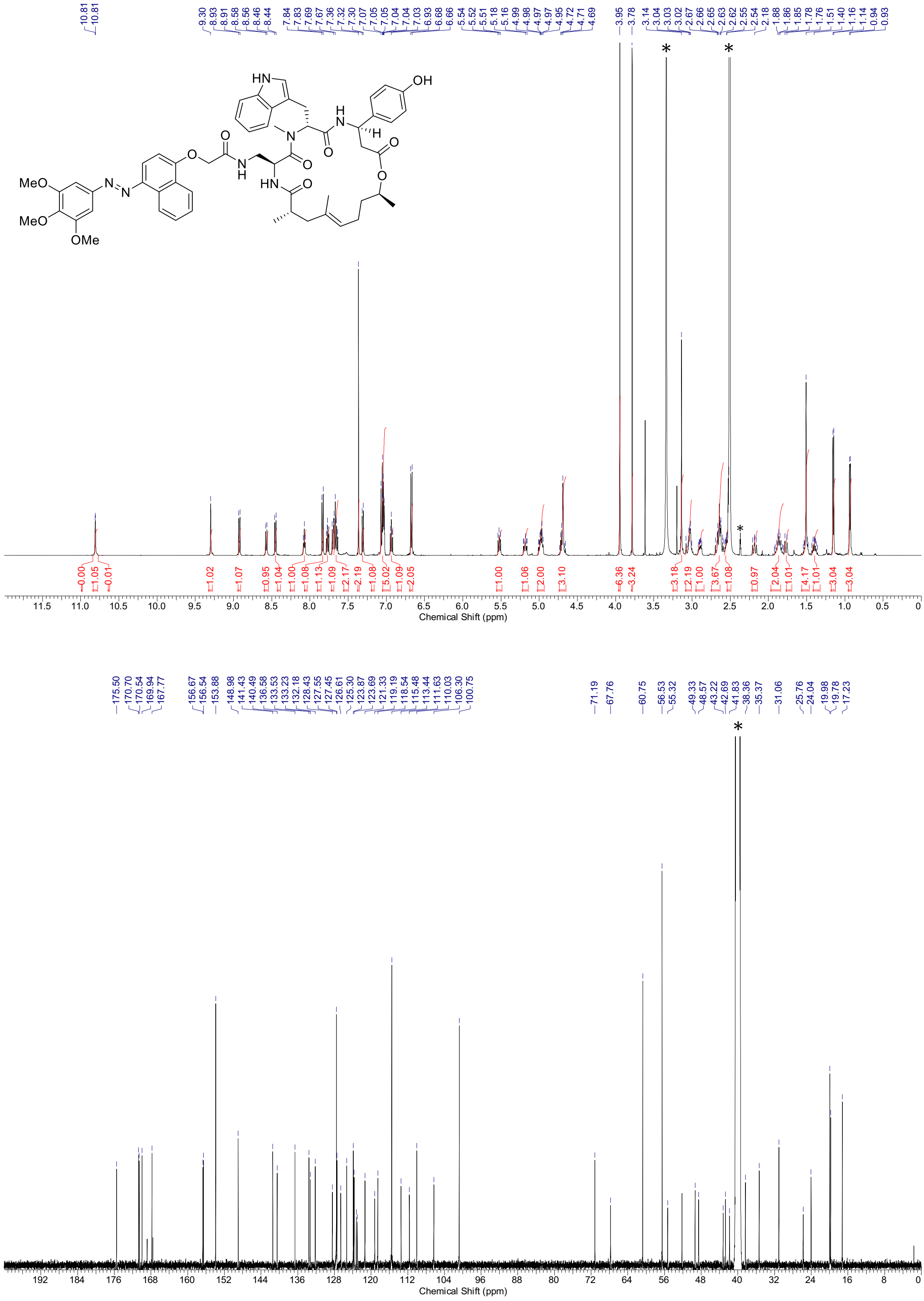

[1]H-NMR spectra (500 MHz, DMSO-*d*6) and [13]C-NMR spectra (126 MHz, DMSO-*d*6) of **S33.** * = NMR-solvent, $H_2O$

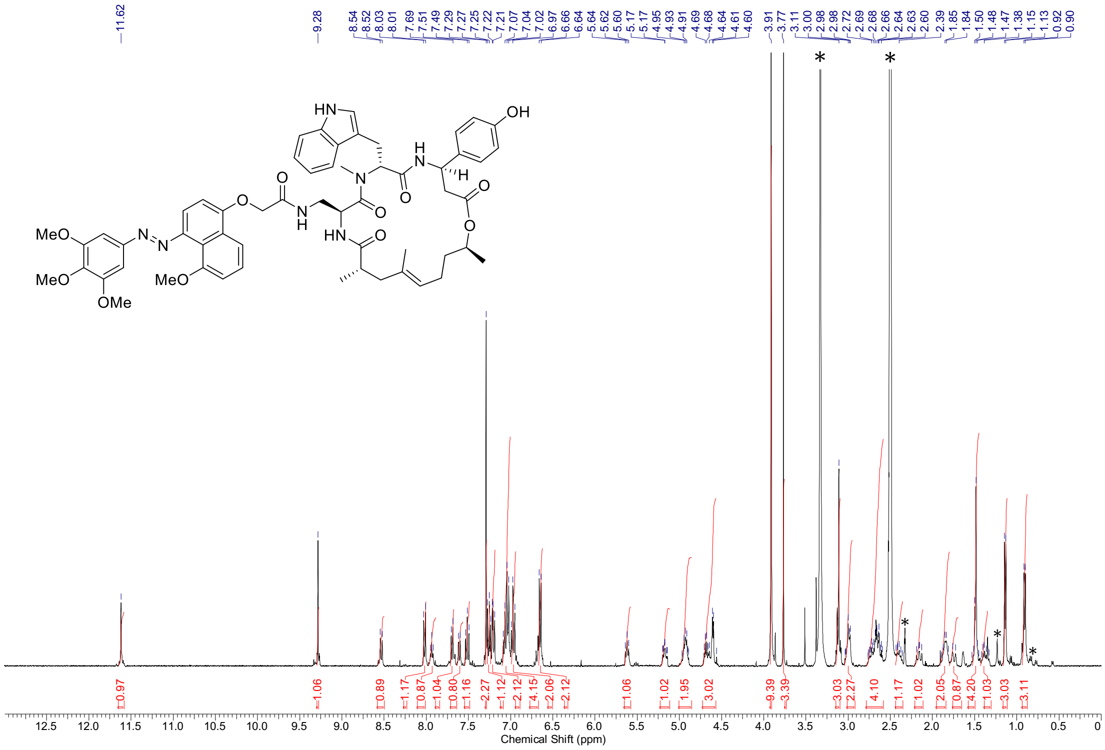

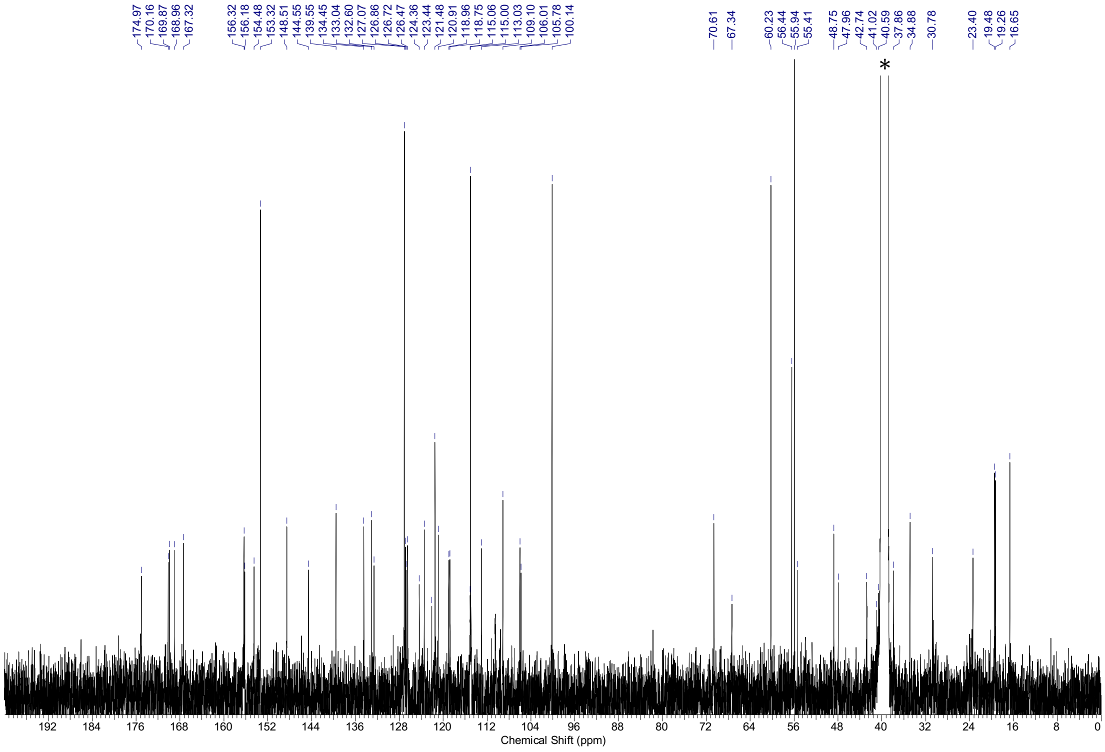

$^{1}$H-NMR spectra (500 MHz, DMSO-$d_6$) and $^{13}$C-NMR spectra (126 MHz, DMSO-$d_6$) of **S34.** * = NMR-solvent, $H_2O$, grease

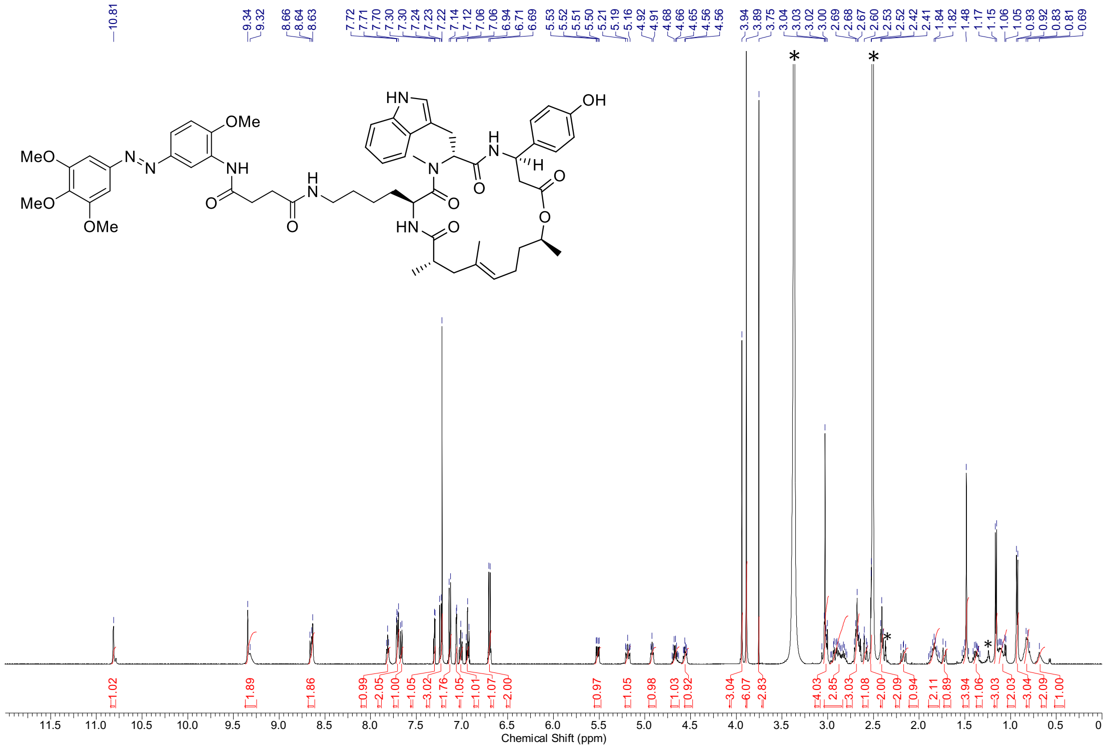

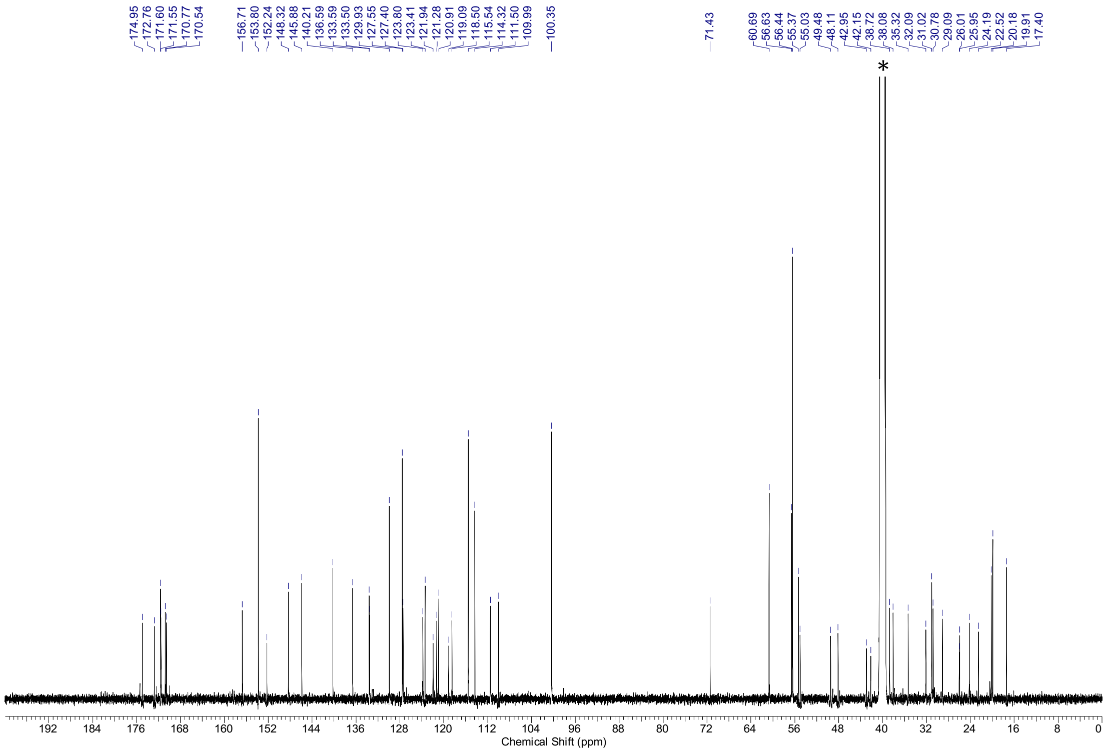

$^1$H-NMR spectra (500 MHz, DMSO-$d_6$) and $^{13}$C-NMR spectra (126 MHz, DMSO-$d_6$) of **S35.** * = NMR-solvent, $H_2O$, grease

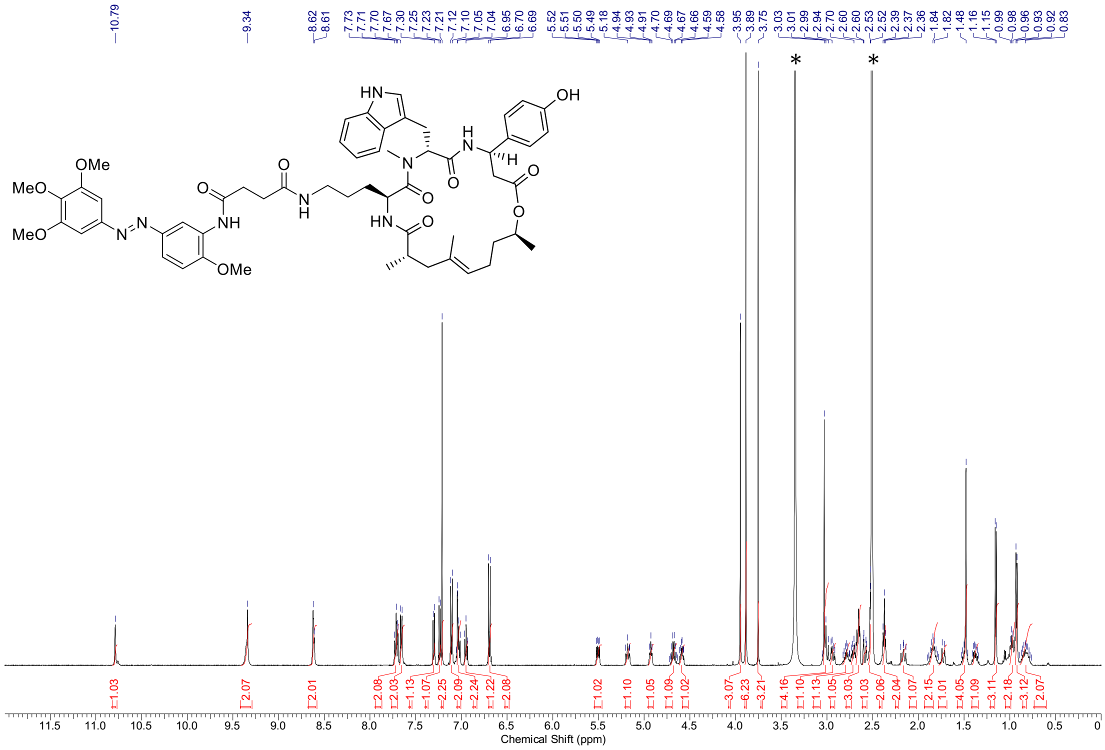

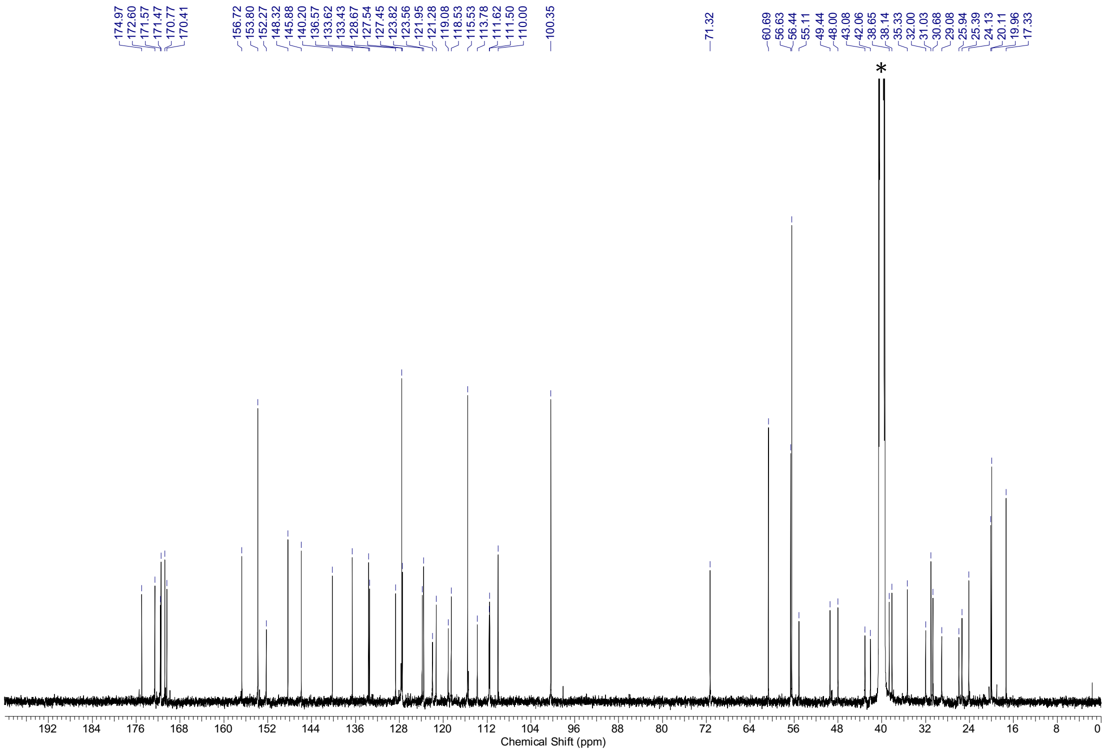

$^{1}$H-NMR spectra (500 MHz, DMSO-*d*$_6$) and $^{13}$C-NMR spectra (126 MHz, DMSO-*d*$_6$) of **S36.** * = NMR-solvent, $H_2O$

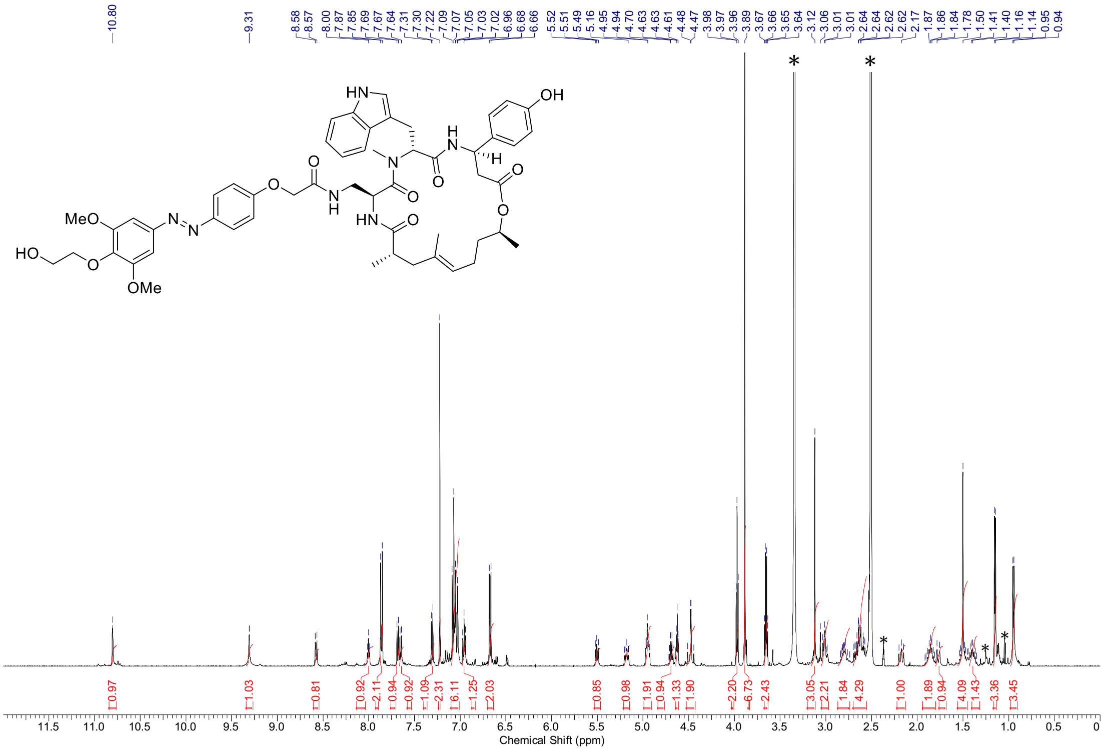

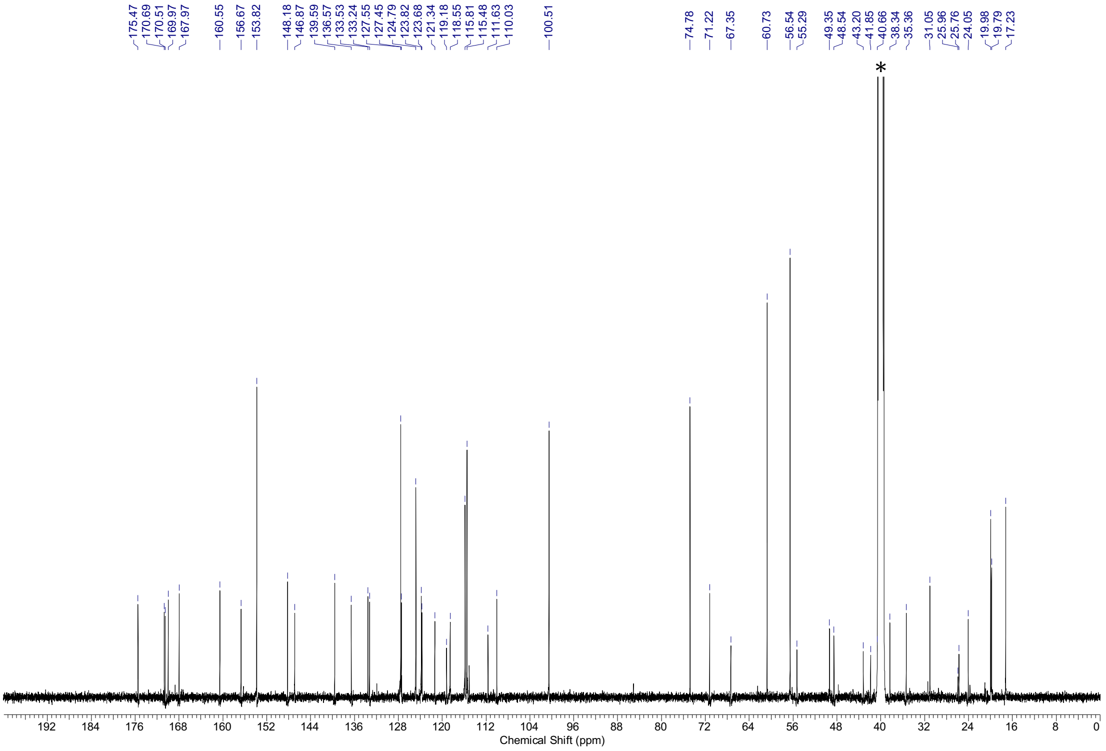

[1]H-NMR spectra (500 MHz, DMSO-$d_6$) and [13]C-NMR spectra (126 MHz, DMSO-$d_6$) of **S37**. * = NMR-solvent, $H_2O$, grease

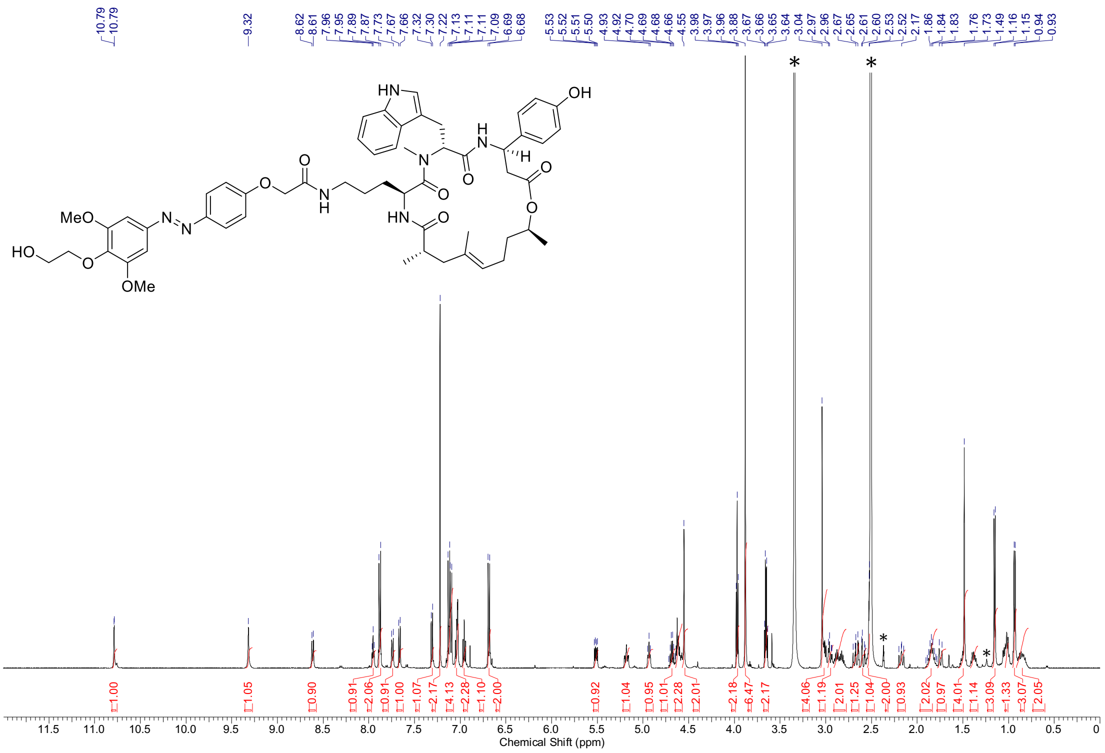

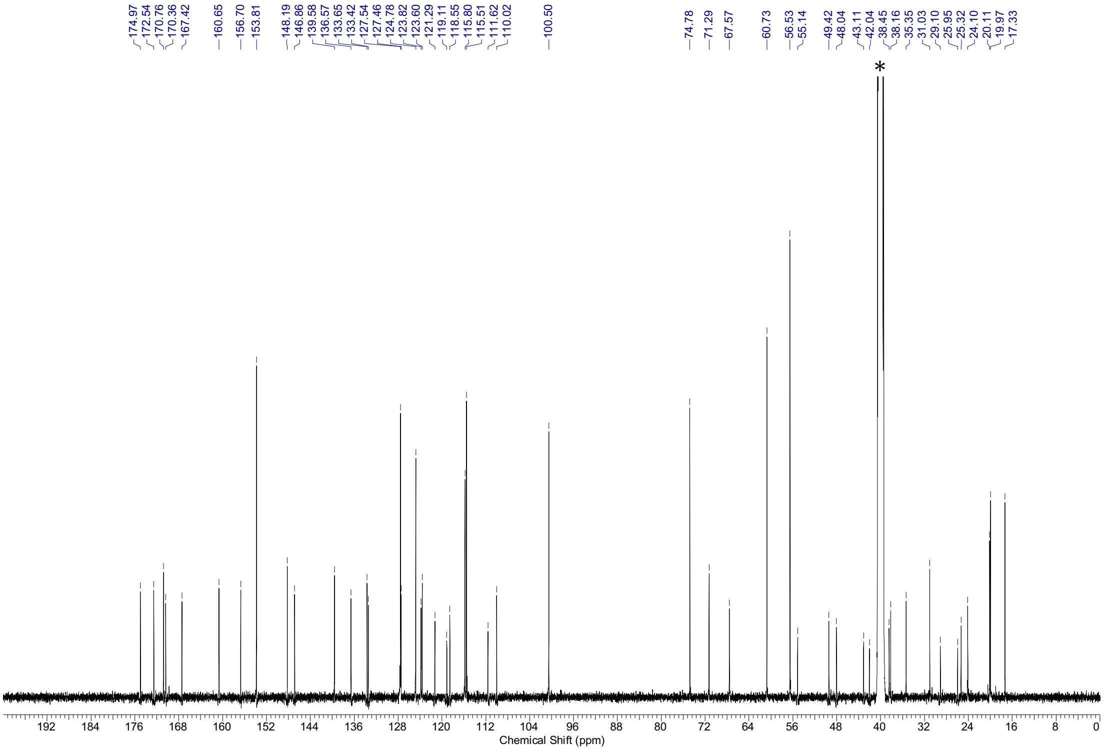

$^{1}$H-NMR spectra (500 MHz, DMSO-*d*$_6$) and $^{13}$C-NMR spectra (126 MHz, DMSO-*d*$_6$) of **S38.** * = NMR-solvent, $H_2O$, grease

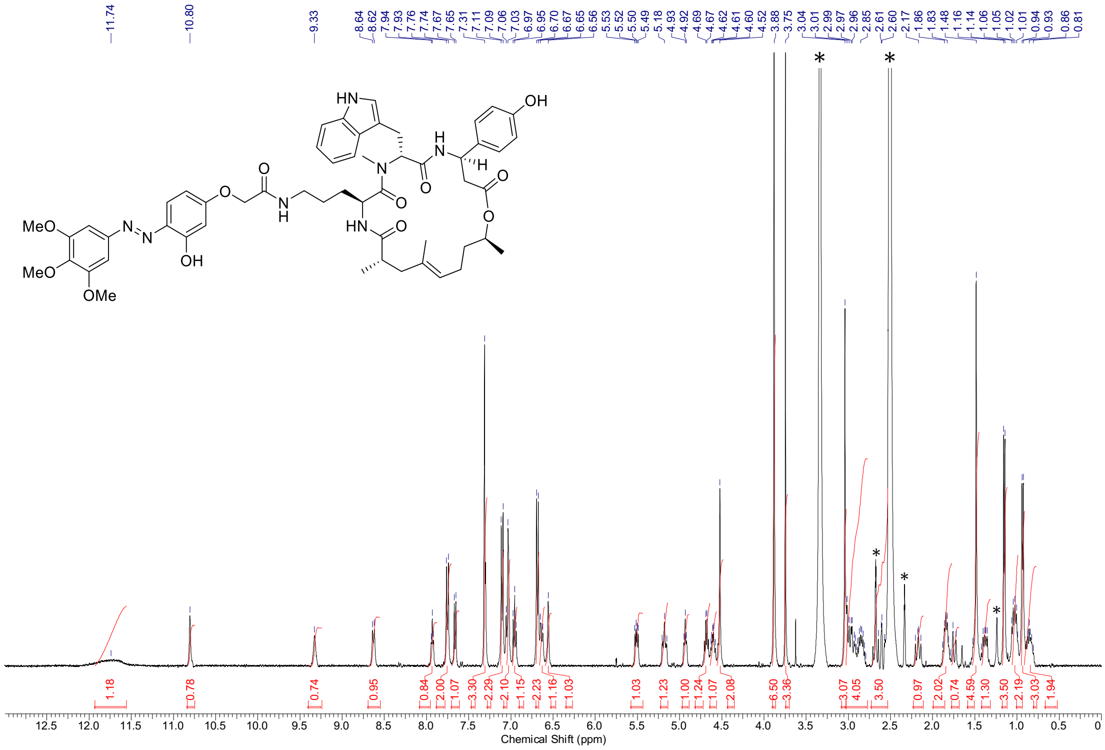

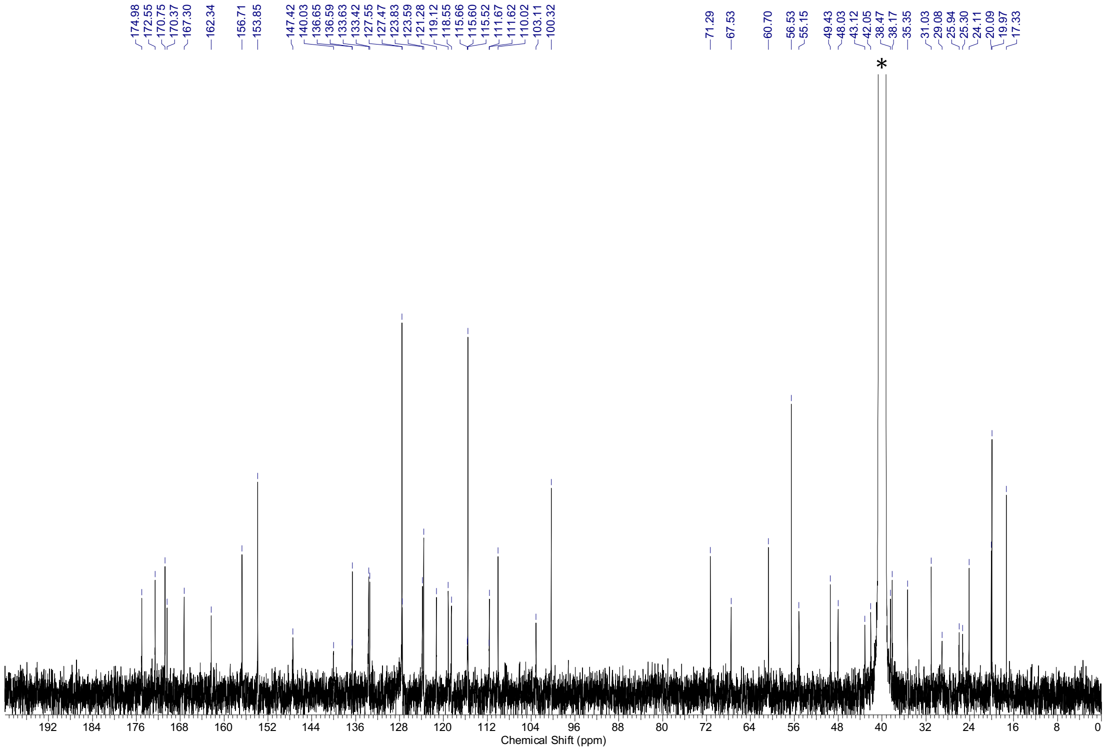

$^{1}H$-NMR spectra (400 MHz, DMSO-$d_6$) and $^{13}C$-NMR spectra (101 MHz, DMSO-$d_6$) of **S39**. * = NMR-solvent, $H_2O$, grease

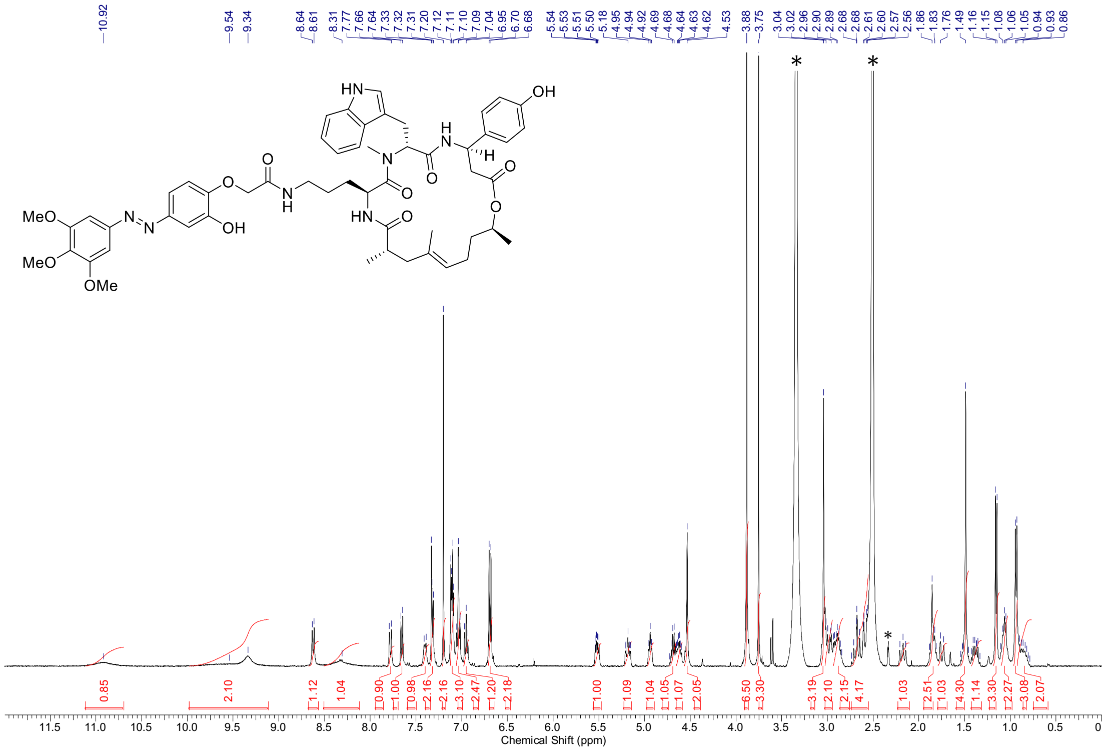

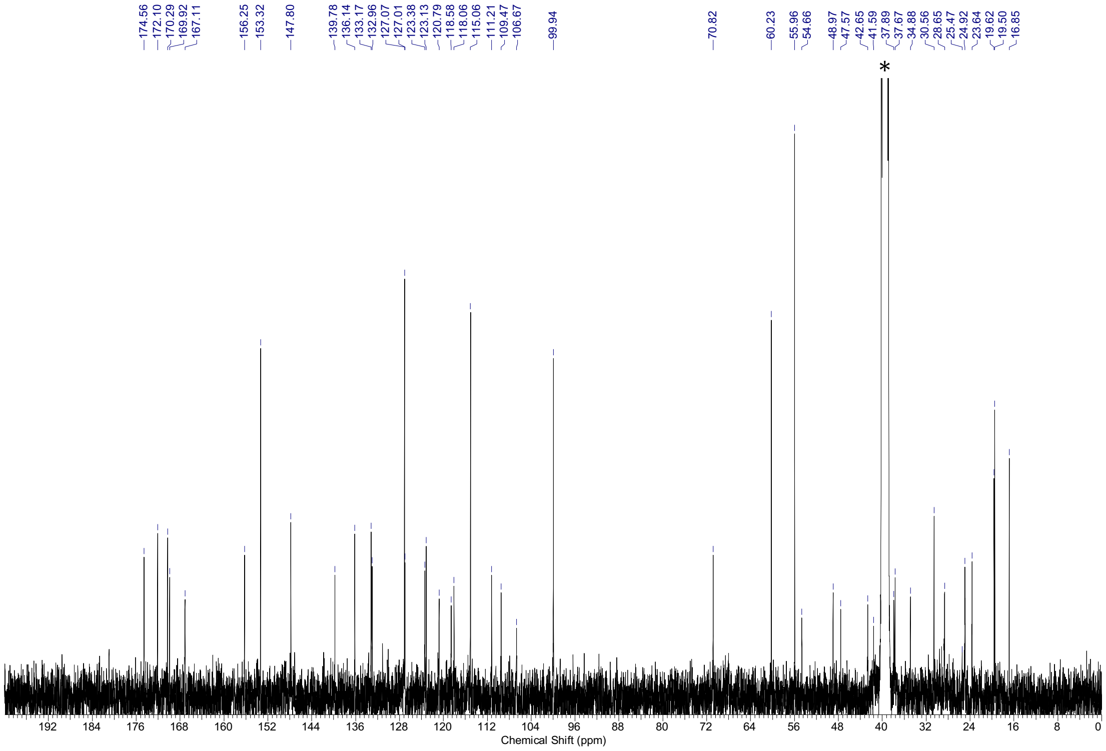

[1]H-NMR spectra (400 MHz, DMSO-$d_6$) and [13]C-NMR spectra (101 MHz, DMSO-$d_6$) of **S40.** * = NMR-solvent, $H_2O$

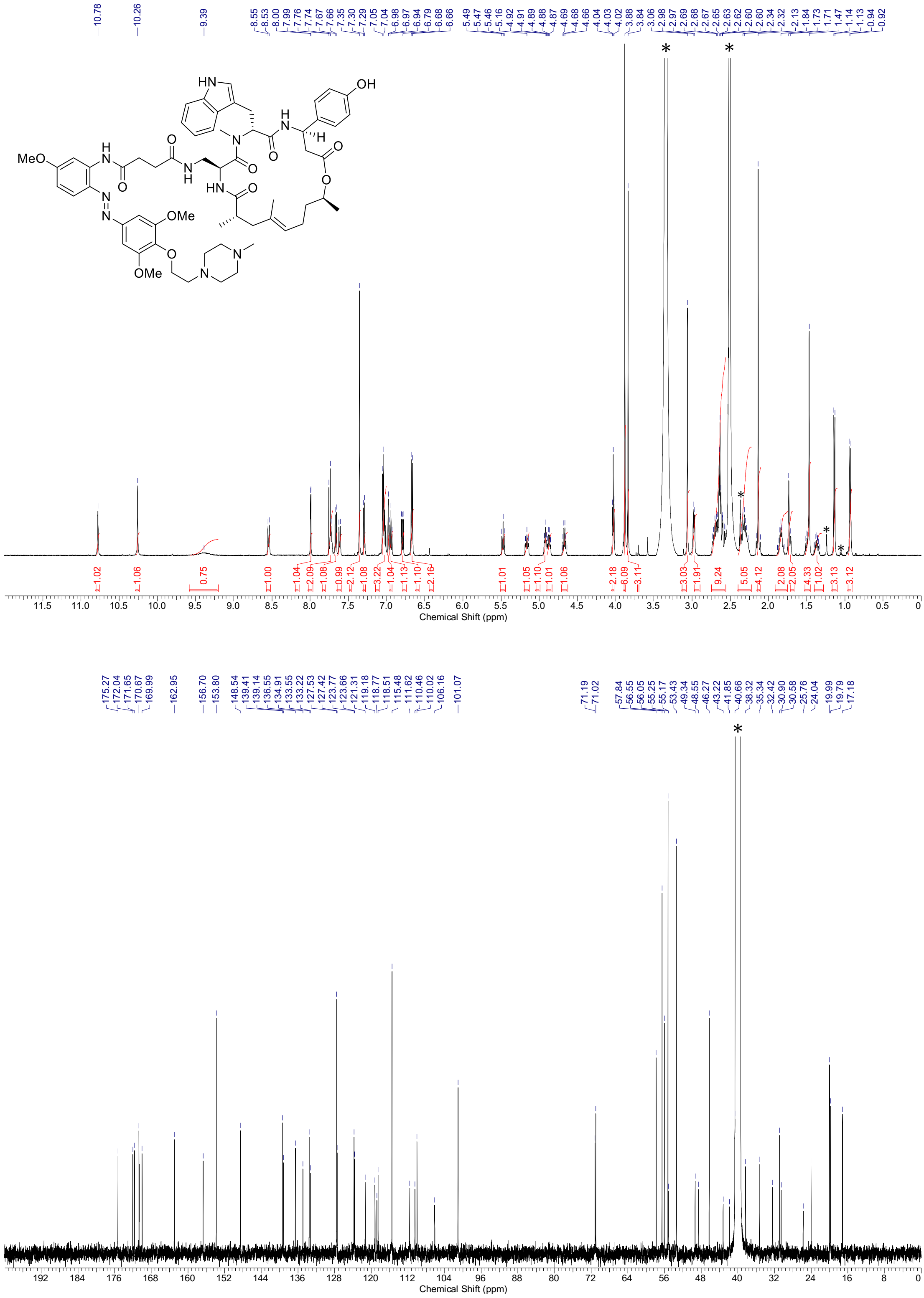

[1]H-NMR spectra (500 MHz, DMSO-$d_6$) and [13]C-NMR spectra (126 MHz, DMSO-$d_6$) of **S41.** * = NMR-solvent, $H_2O$, grease

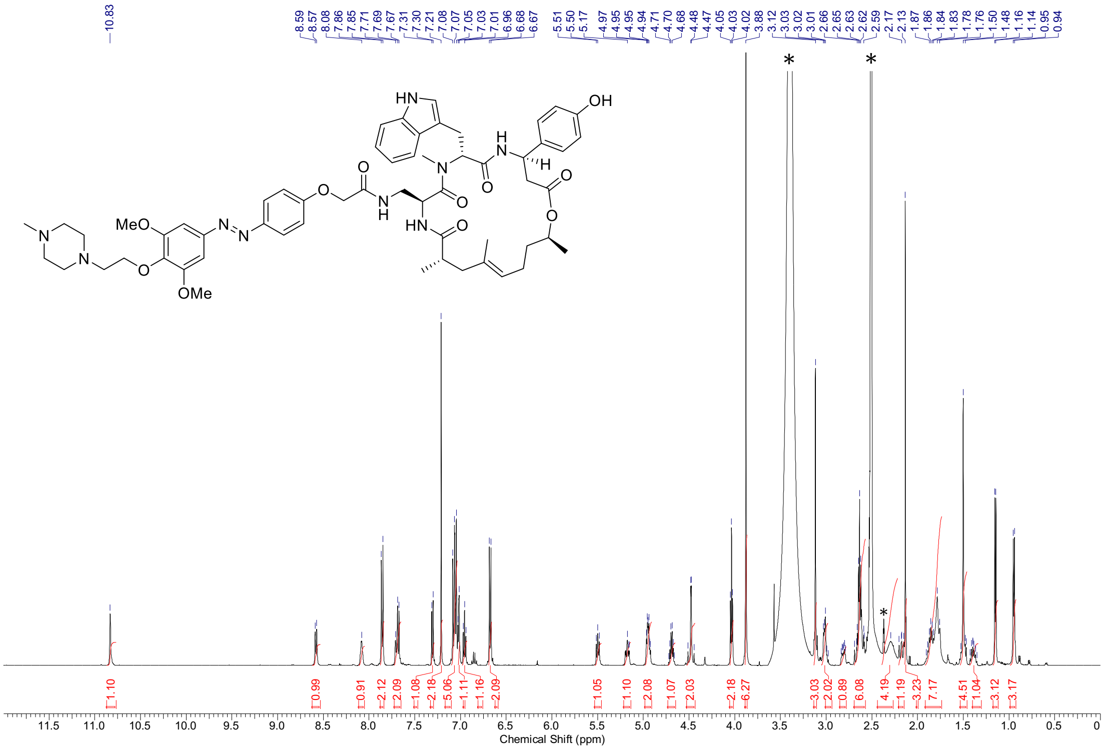

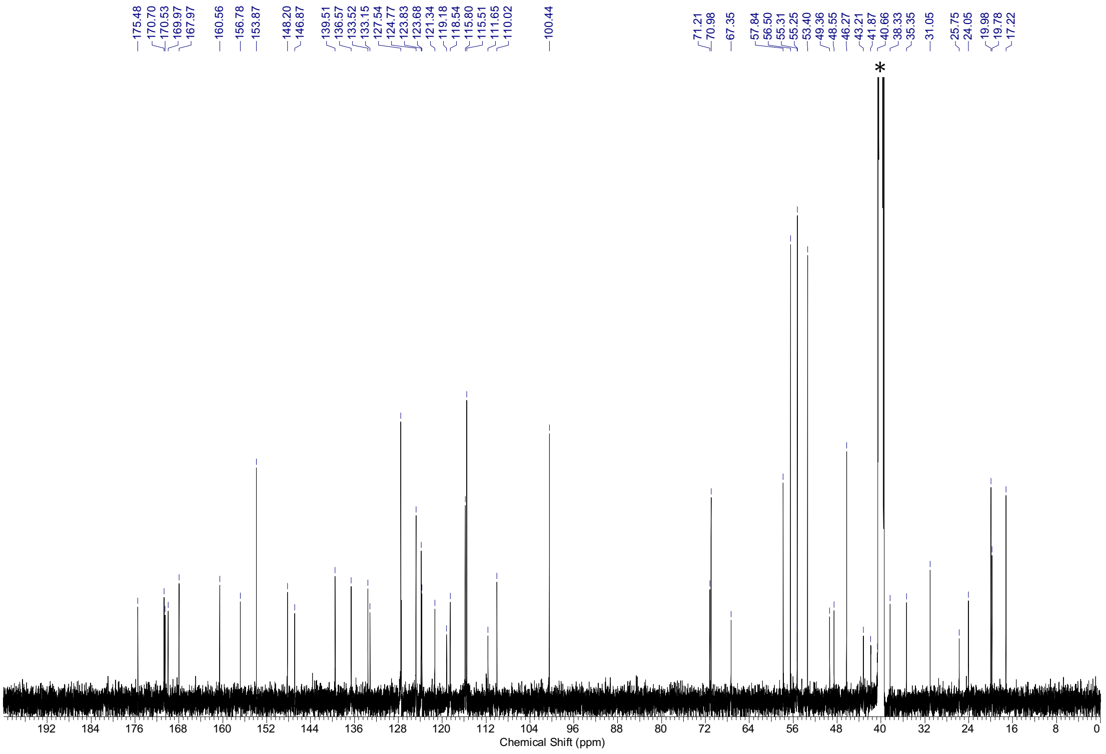

[1]H-NMR spectra (500 MHz, DMSO-$d_6$) and [13]C-NMR spectra (126 MHz, DMSO-$d_6$) of **S42.** * = NMR-solvent, $H_2O$

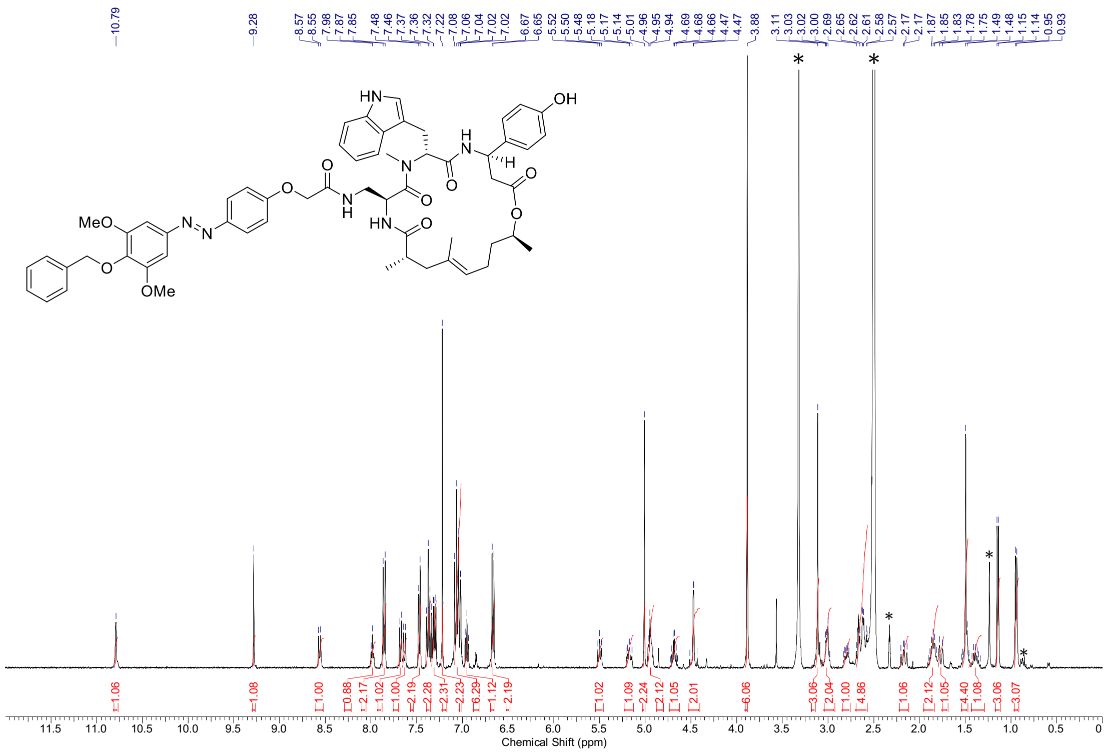

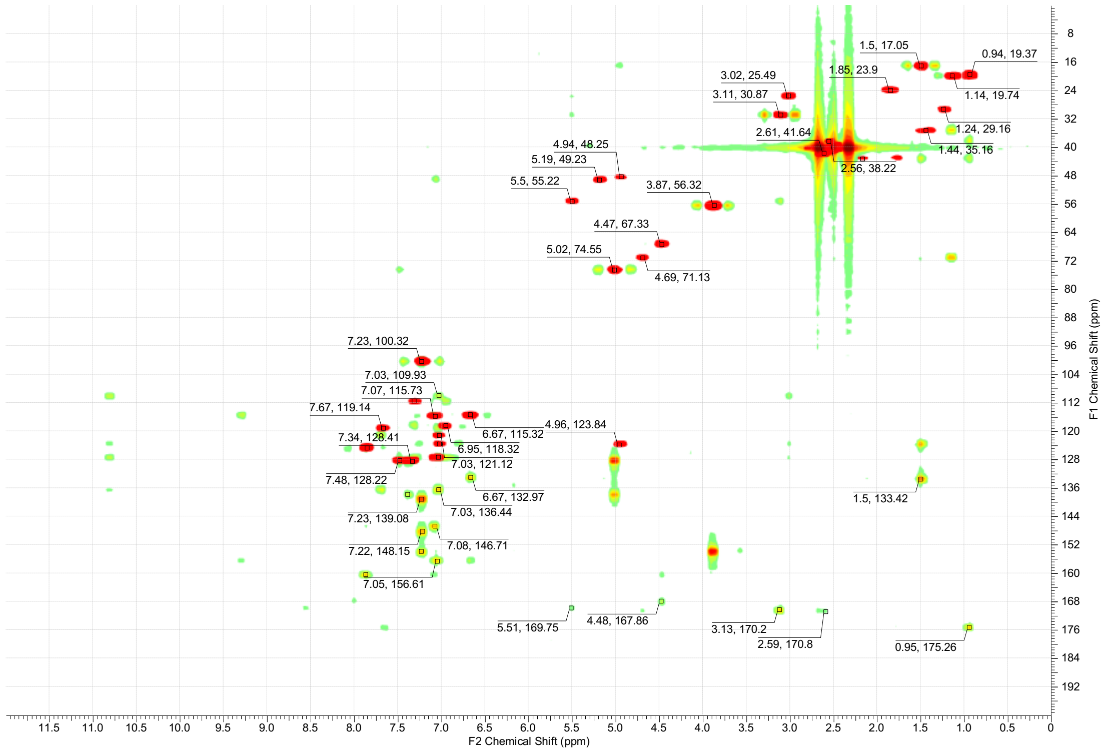

$^{1}$H-NMR spectra (400 MHz, DMSO-$d_6$) and overlay of $^{13}$C-HSQC (red) and $^{13}$C-HMBC (green) spectra (100 MHz, DMSO-$d_6$) of **S43.** * = NMR-solvent, $H_2O$, grease

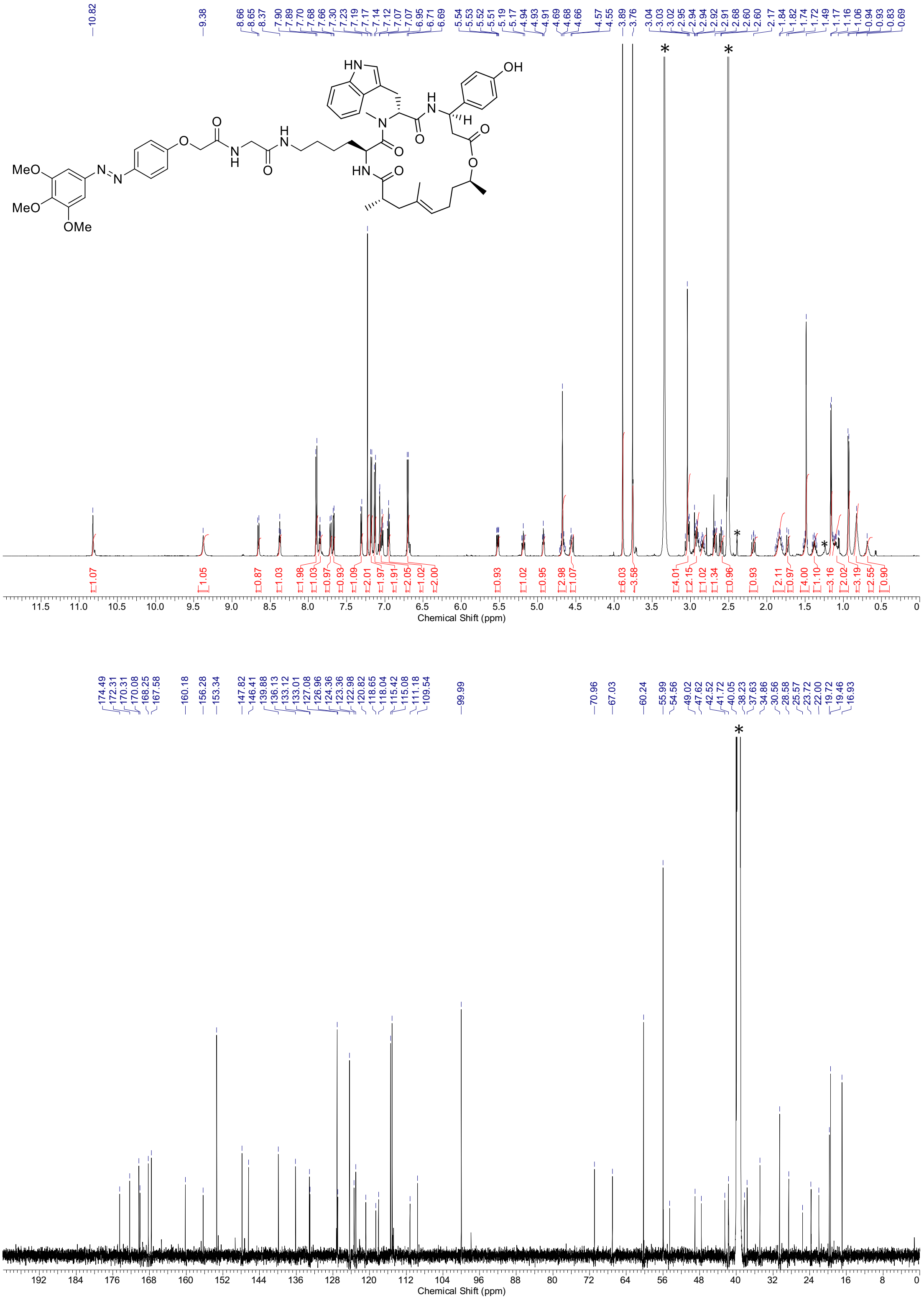

$^{1}$H-NMR spectra (600 MHz, DMSO-$d_6$) and $^{13}$C-NMR spectra (151 MHz, DMSO-$d_6$) of **S44.** * = NMR-solvent, $H_2O$, grease

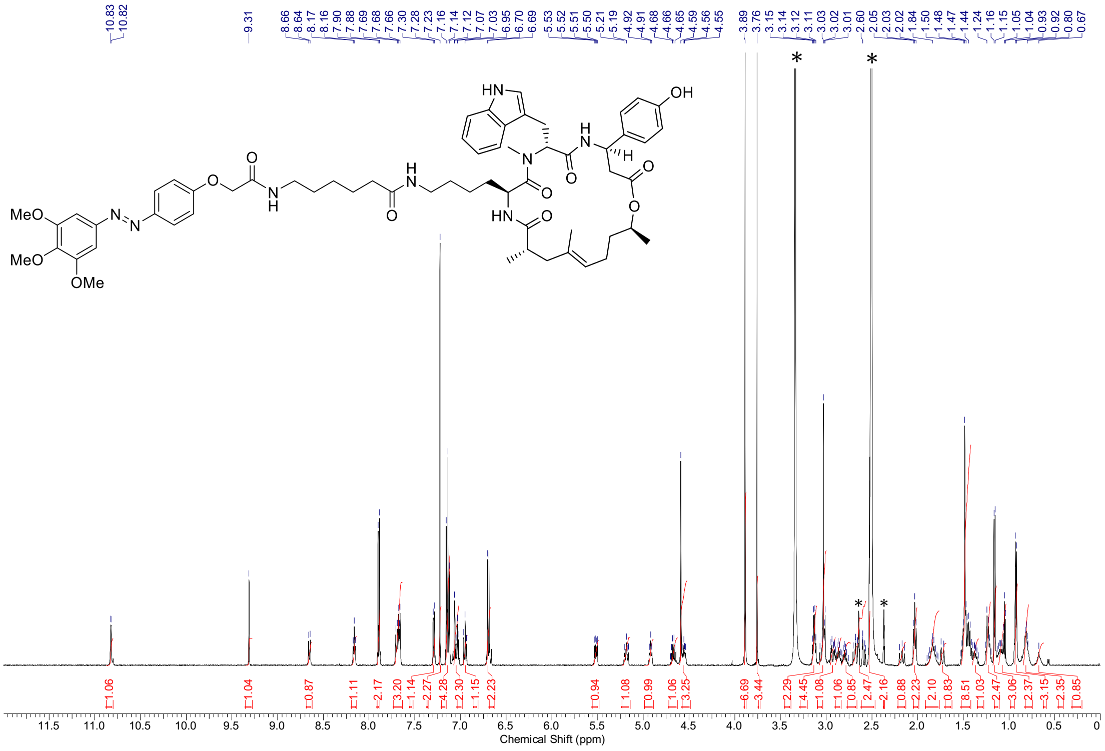

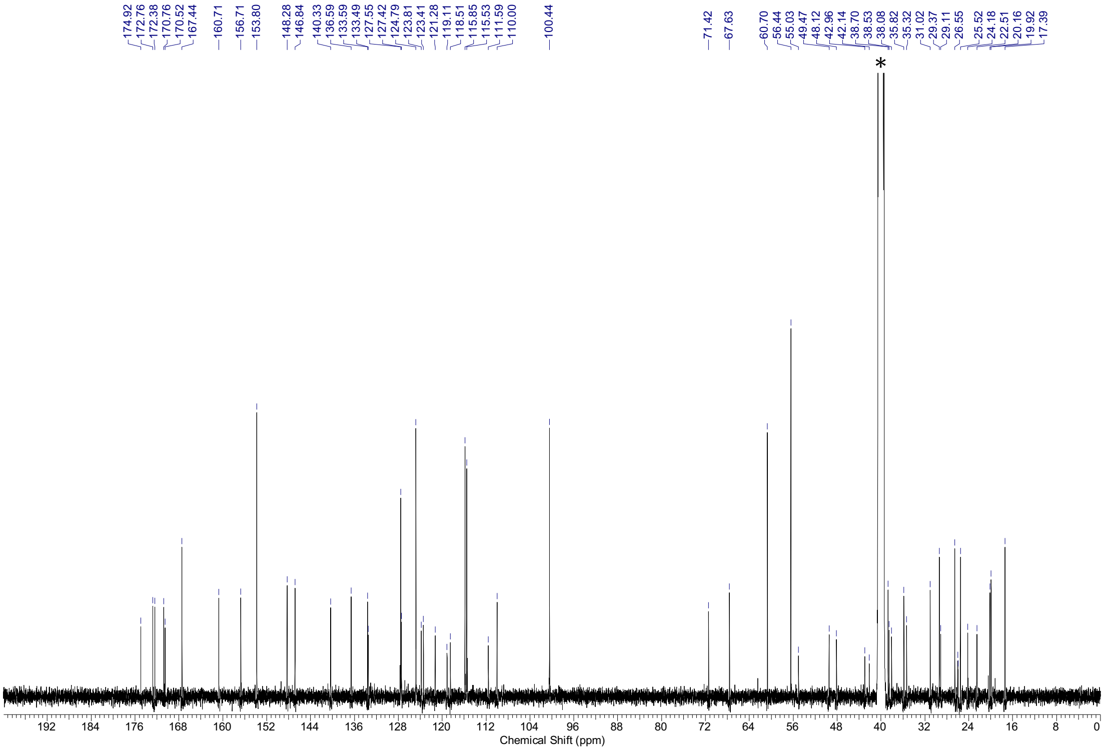

[1]H-NMR spectra (500 MHz, DMSO-*d*6) and [13]C-NMR spectra (126 MHz, DMSO-*d*6) of **S45.** * = NMR-solvent, $H_2O$

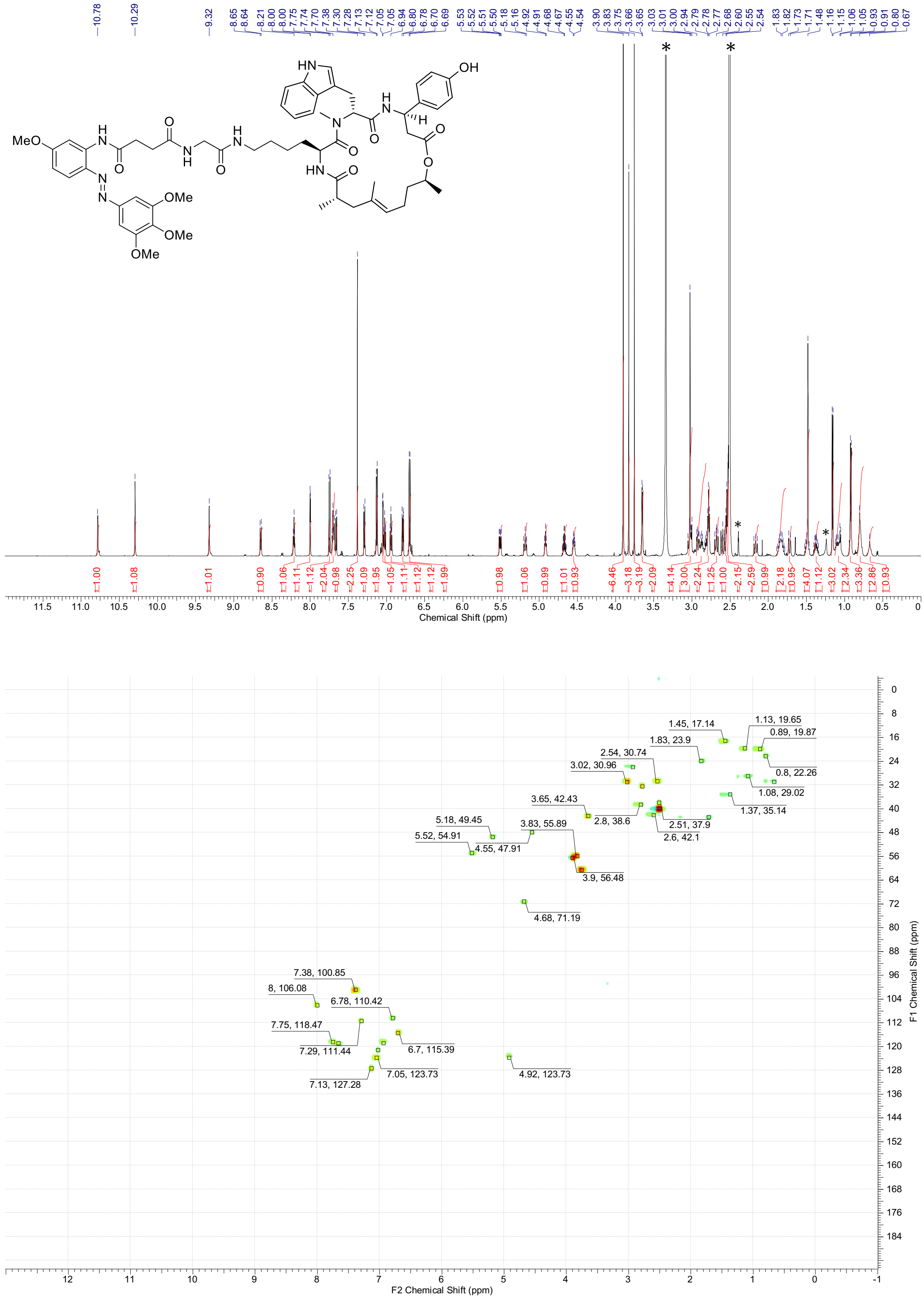

$^{1}$H-NMR spectra (600 MHz, DMSO-$d_6$) and $^{13}$C-HSQC spectra (150 MHz, DMSO-$d_6$) of **S46.** * = NMR-solvent, $H_2O$, grease

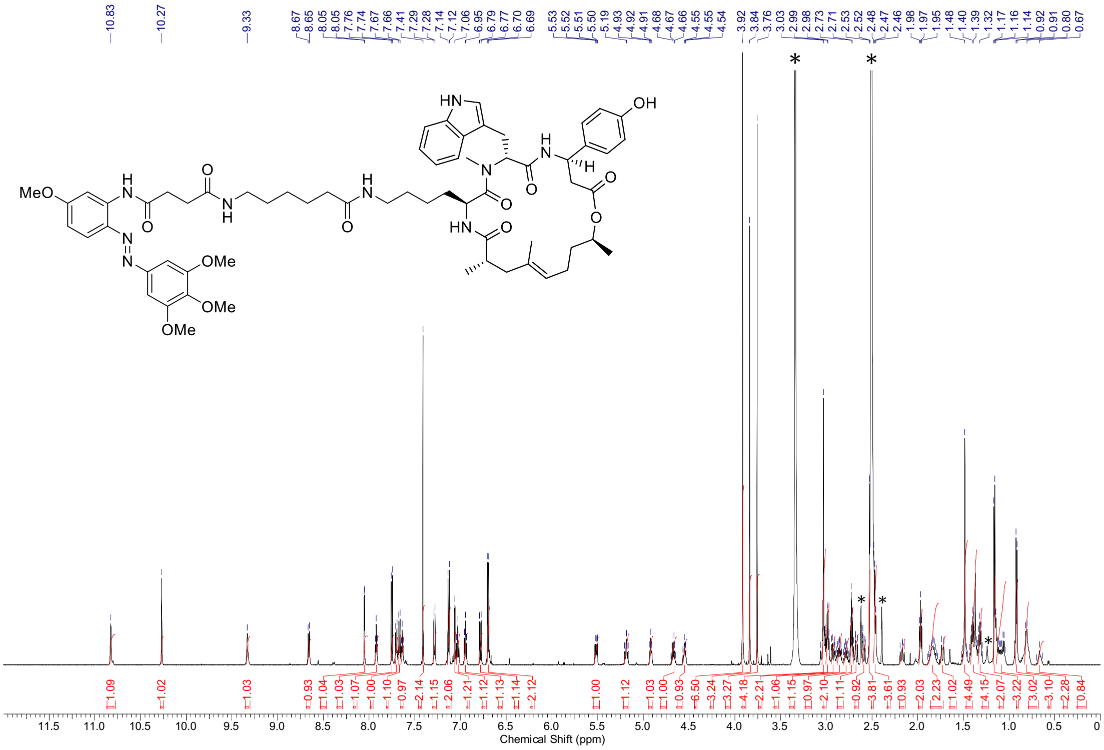

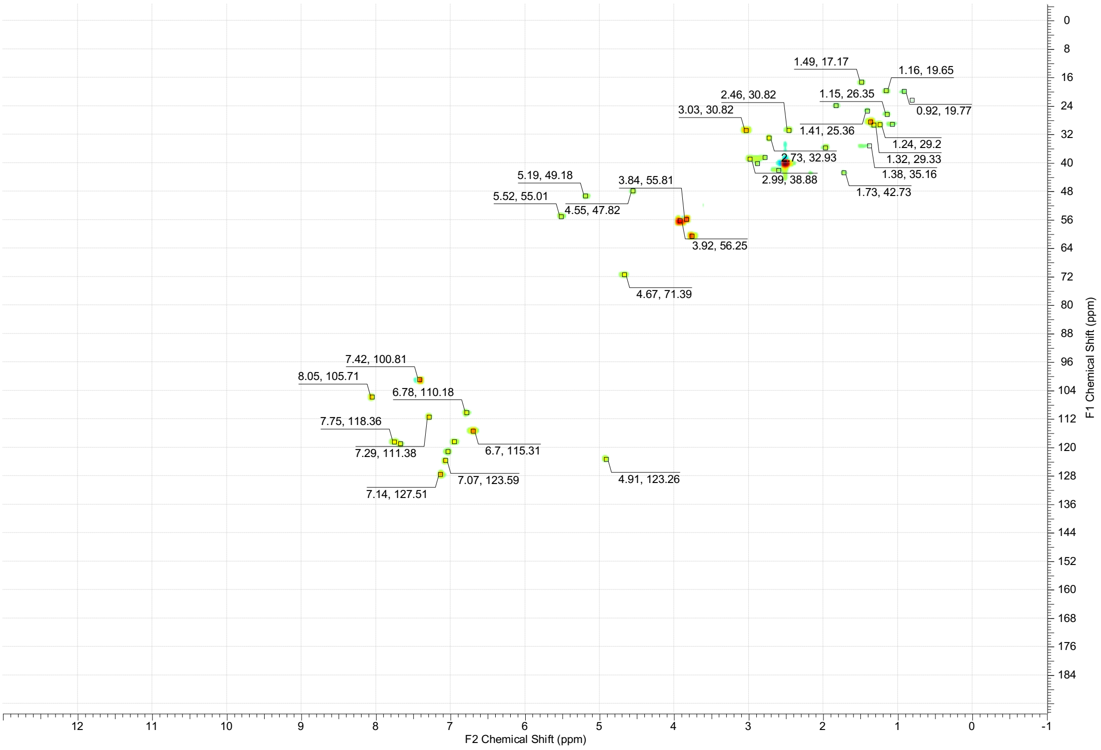

$^{1}$H-NMR spectra (600 MHz, DMSO-$d_6$) and $^{13}$C-HSQC spectra (150 MHz, DMSO-$d_6$) of **S47.** * = NMR-solvent, $H_2O$, grease

### Supplementary Note 4: UV-VIS spectra

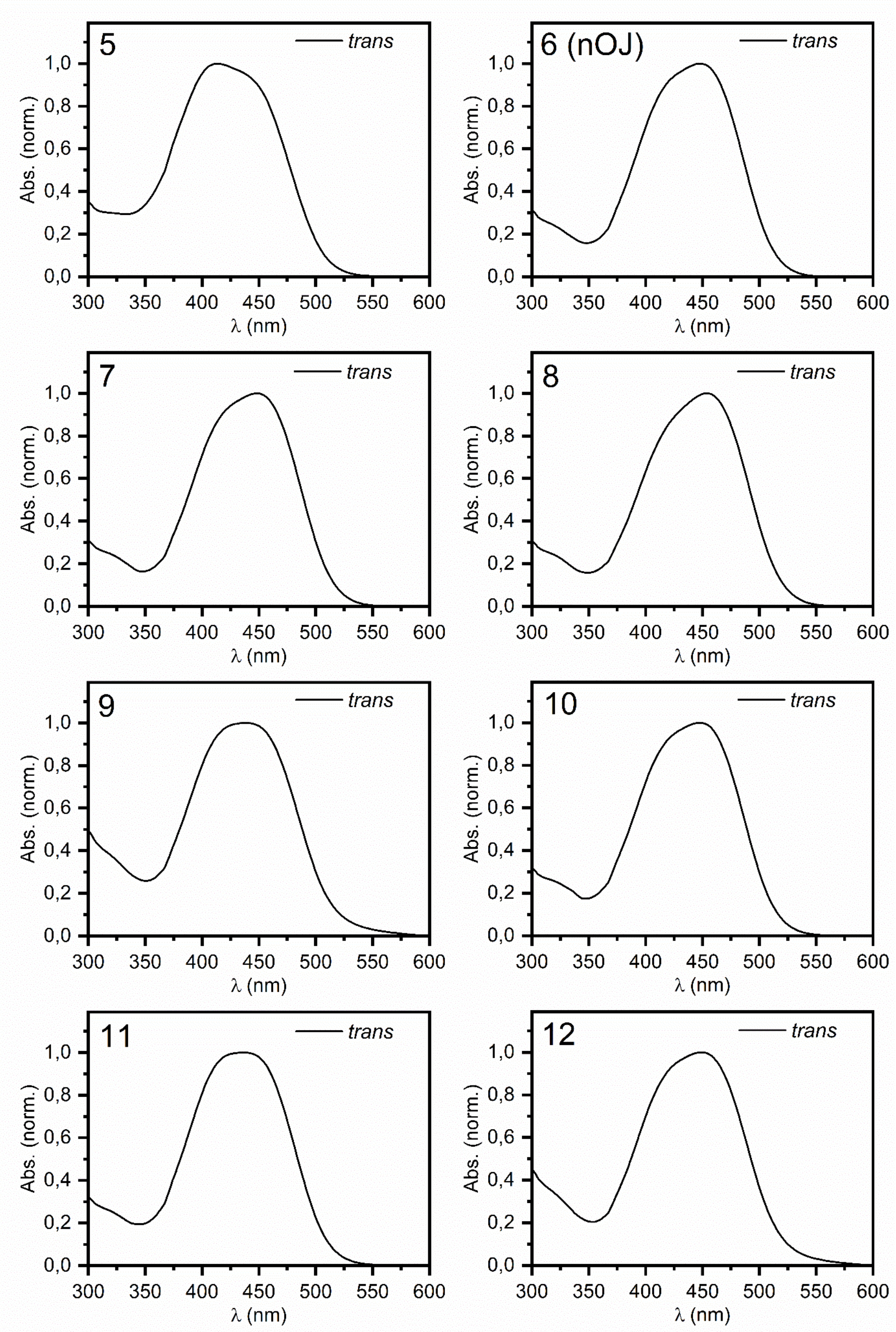

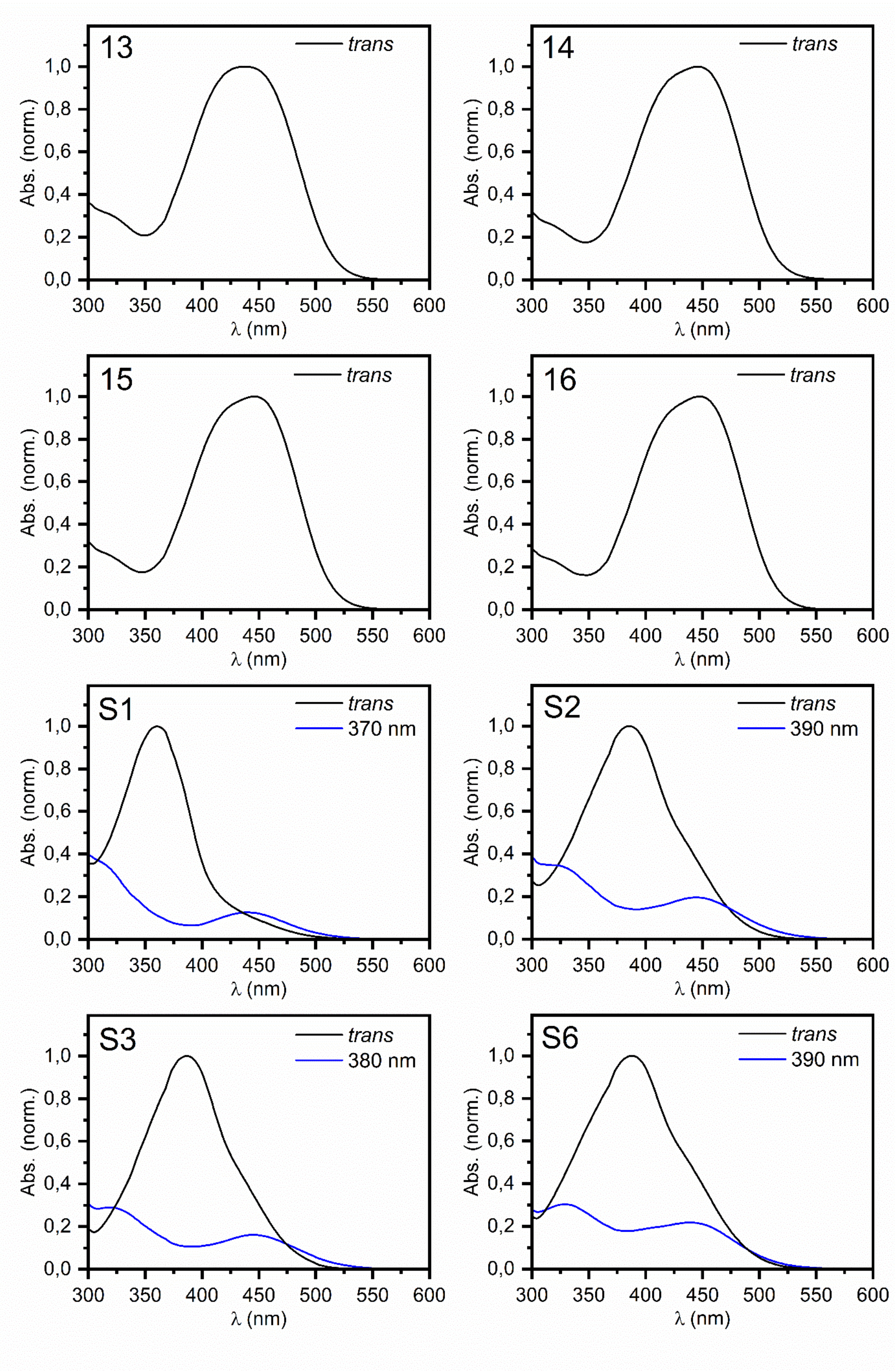
13
trans
14
trans
15
trans
16
trans
S1
trans
370 nm
S2
trans
390 nm
S3
trans
380 nm
S6
trans
390 nm
Abs. (norm.)
1,0
0,8
0,6
0,4
0,2
0,0
300 350 400 450 500 550 600
λ (nm)

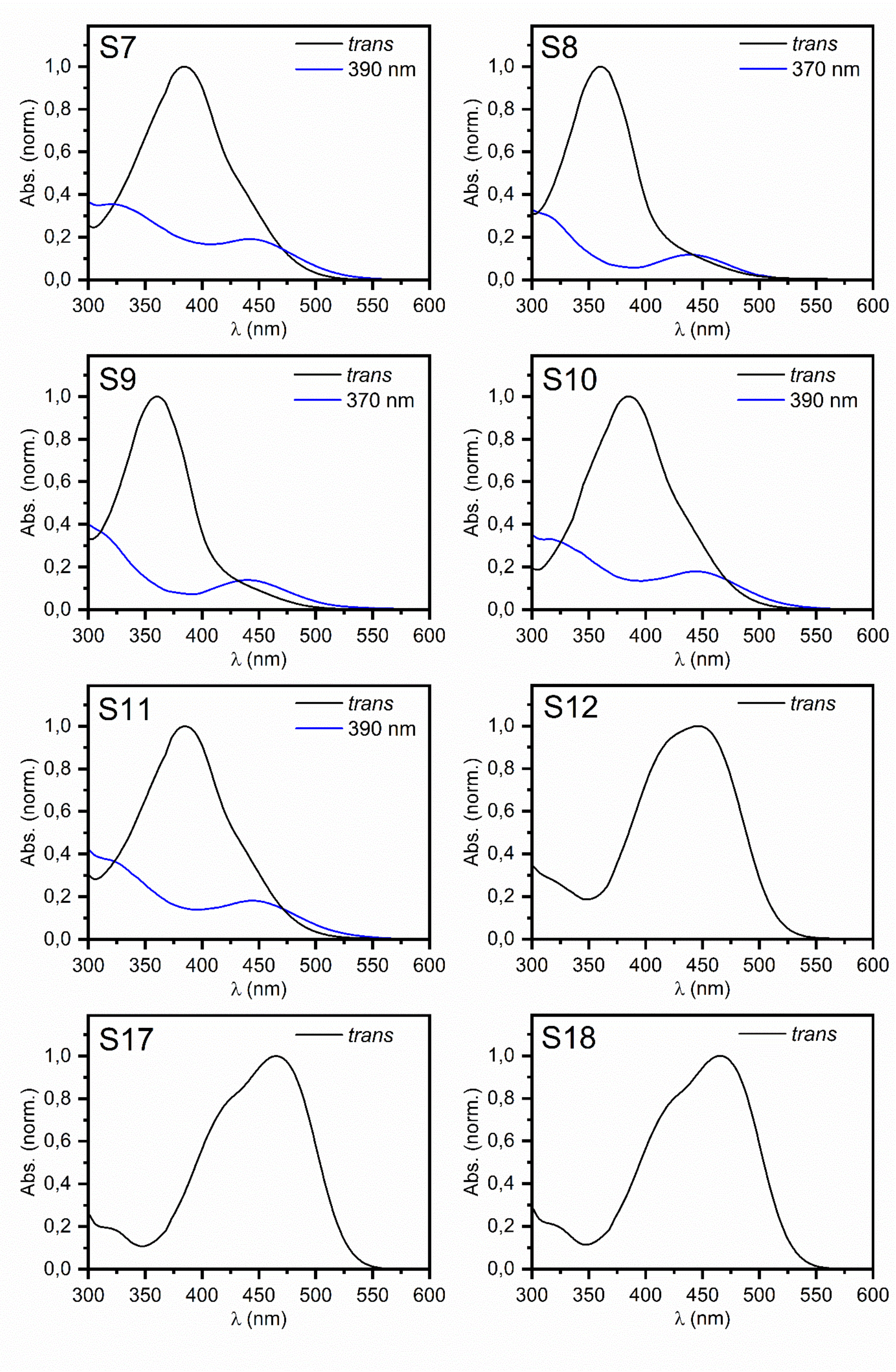
S7
trans
390 nm
S8
trans
370 nm
S9
trans
370 nm
S10
trans
390 nm
S11
trans
390 nm
S12
trans
S17
trans
S18
trans
Abs. (norm.)
λ (nm)
0,0
0,2
0,4
0,6
0,8
1,0
300
350
400
450
500
550
600

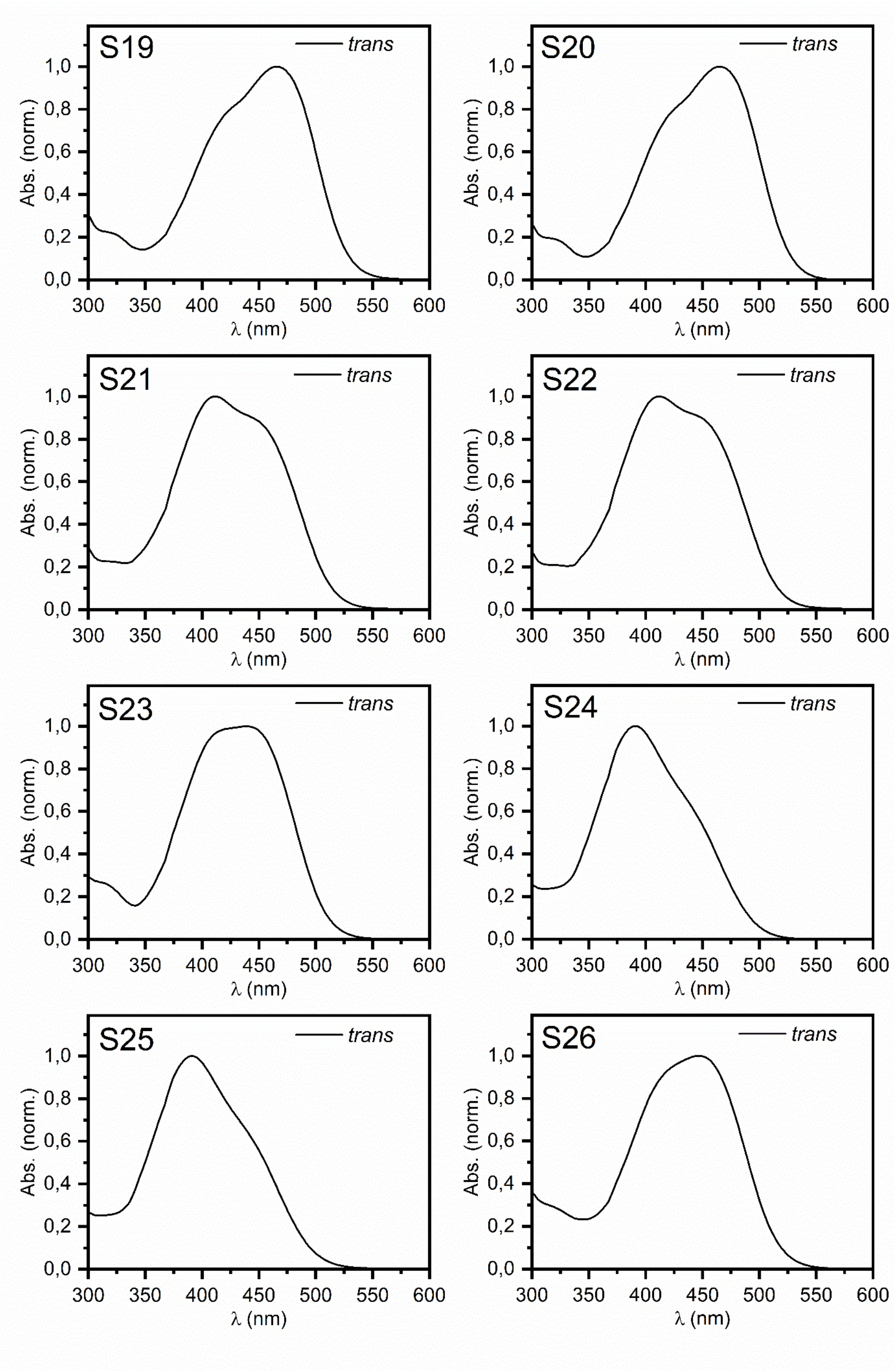
S19
trans
S20
trans
S21
trans
S22
trans
S23
trans
S24
trans
S25
trans
S26
trans
Abs. (norm.)
λ (nm)
0,0
0,2
0,4
0,6
0,8
1,0
300
350
400
450
500
550
600

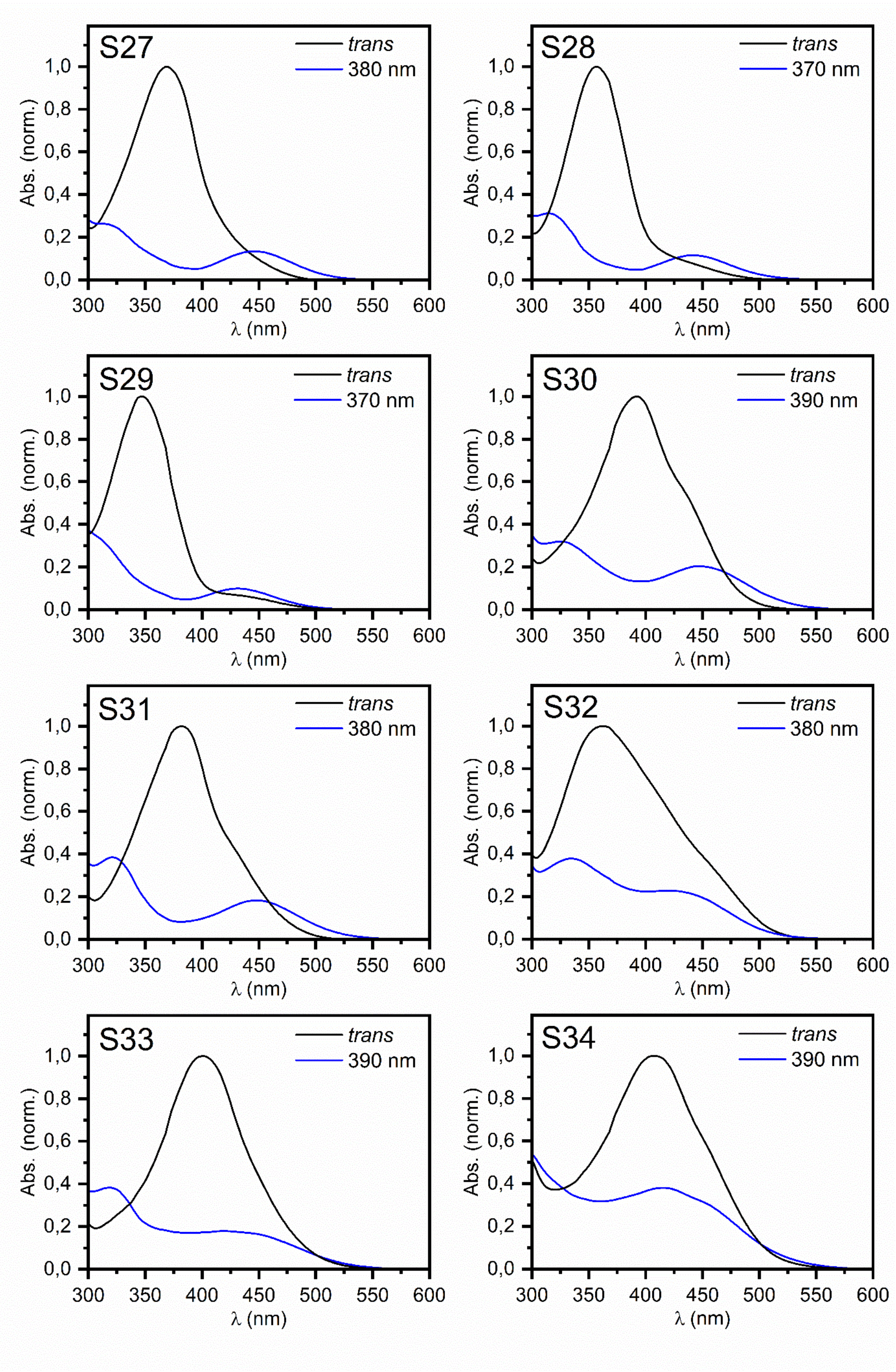
S27
trans
380 nm
S28
trans
370 nm
S29
trans
370 nm
S30
trans
390 nm
S31
trans
380 nm
S32
trans
380 nm
S33
trans
390 nm
S34
trans
390 nm
Abs. (norm.)
λ (nm)

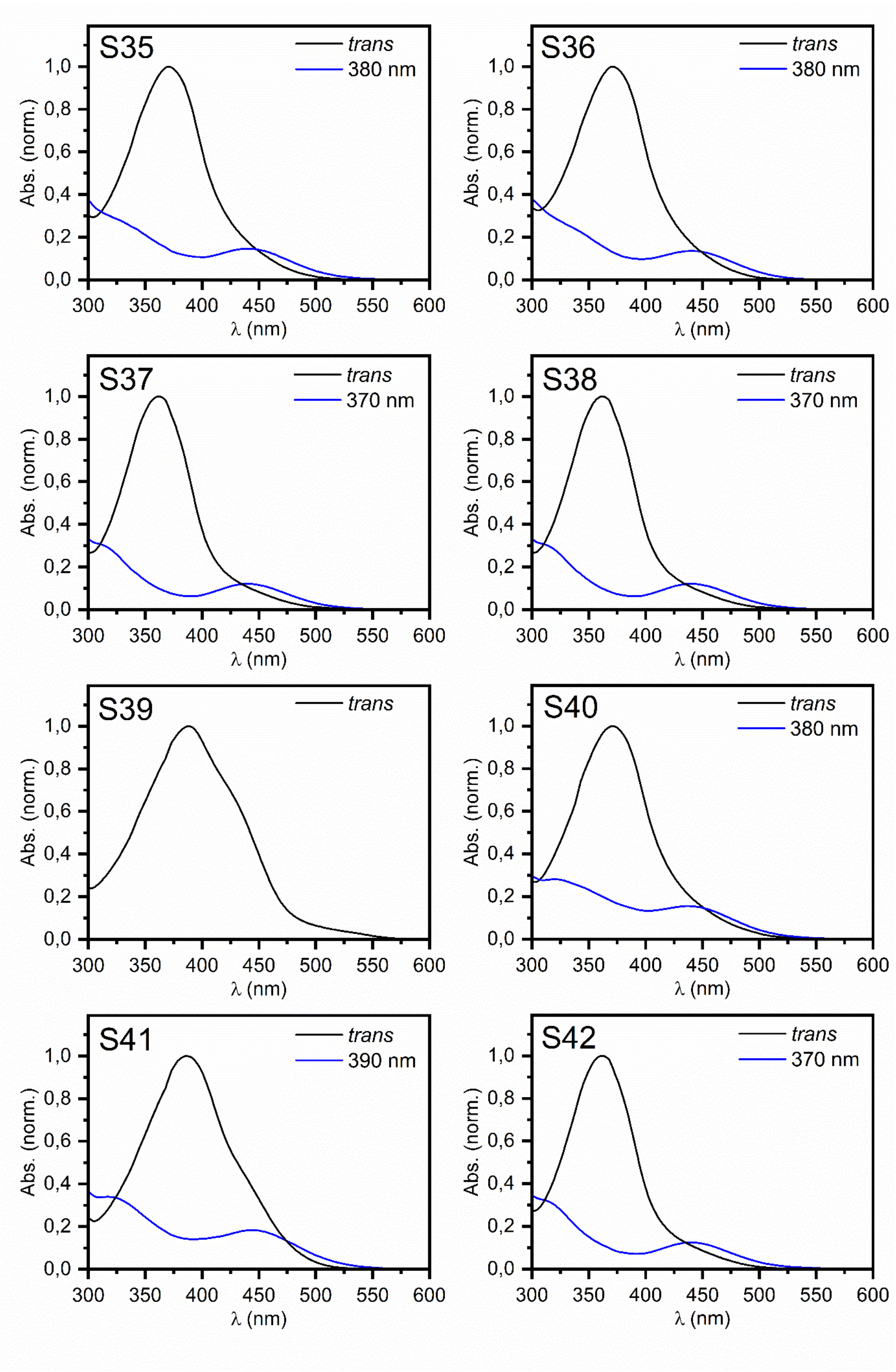
S35
trans
380 nm
S36
trans
380 nm
S37
trans
370 nm
S38
trans
370 nm
S39
trans
S40
trans
380 nm
S41
trans
390 nm
S42
trans
370 nm
Abs. (norm.)
λ (nm)

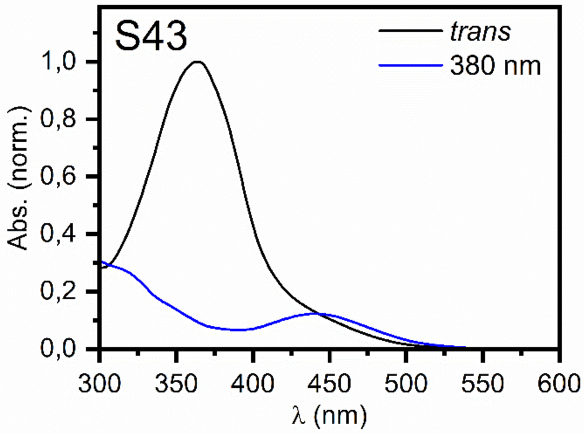
S43
trans
380 nm
Abs. (norm.)
1,0
0,8
0,6
0,4
0,2
0,0
300 350 400 450 500 550 600
λ (nm)

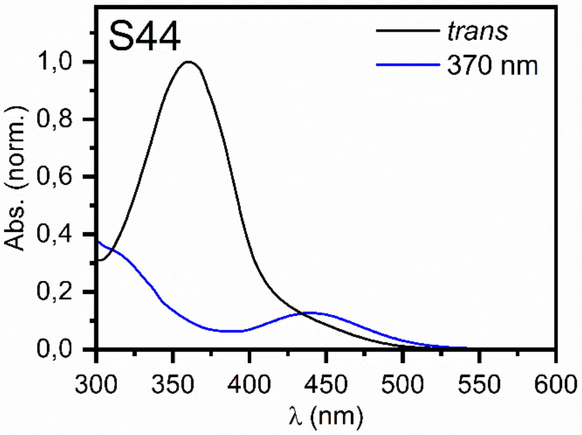
S44
trans
370 nm
Abs. (norm.)
1,0
0,8
0,6
0,4
0,2
0,0
300 350 400 450 500 550 600
λ (nm)

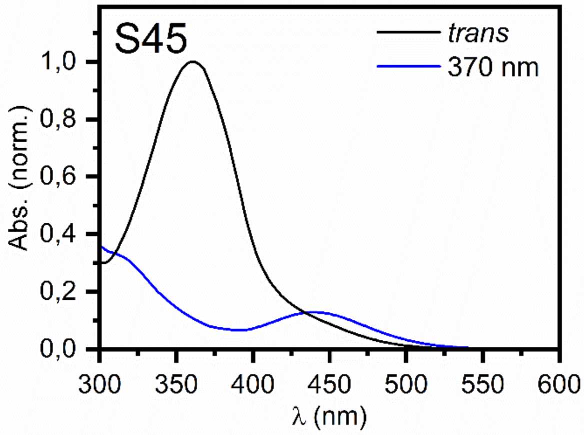
S45
trans
370 nm
Abs. (norm.)
1,0
0,8
0,6
0,4
0,2
0,0
300 350 400 450 500 550 600
λ (nm)

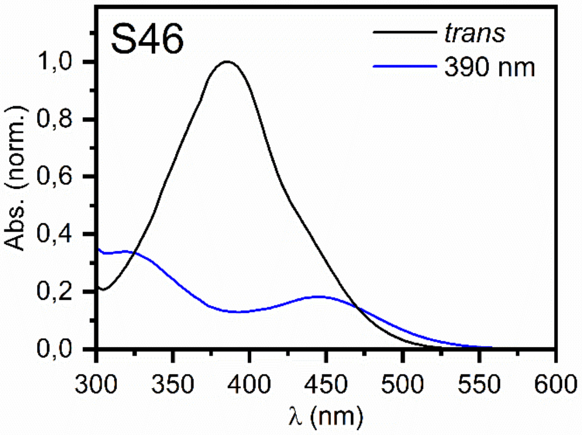
S46
trans
390 nm
Abs. (norm.)
1,0
0,8
0,6
0,4
0,2
0,0
300 350 400 450 500 550 600
λ (nm)

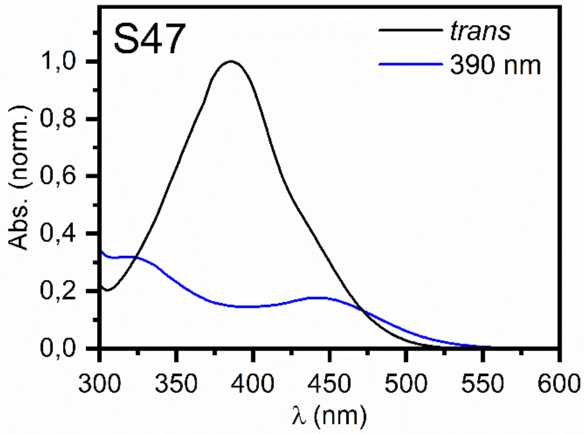
S47
trans
390 nm
Abs. (norm.)
1,0
0,8
0,6
0,4
0,2
0,0
300 350 400 450 500 550 600
λ (nm)
